# Biphasic Response of Protein Kinase A to Cyclic Adenosine Monophosphate Triggers Distinct Epithelial Phenotypes

**DOI:** 10.1101/747030

**Authors:** João Pedro Fonseca, Zara Y. Weinberg, Elham Aslankoohi, Hana El-Samad

## Abstract

Despite the large diversity of the proteins involved in cellular signaling, many intracellular signaling pathways converge onto one of only dozens of small molecule second messengers. Cyclic adenosine monophosphate (cAMP), one of these second messengers, is known to regulate activity of both Protein Kinase A (PKA) and the Extracellular Regulated Kinase (ERK), among other signaling pathways. In its role as an important cellular signaling hub, intracellular cAMP concentration has long been assumed to monotonically regulate its known effectors.

Using an optogenetictool that can introduce precise amounts of cAMP in MDCKI cells, we identify genes whose expression changes biphasically with monotonically increasing cAMP levels. By examining the behavior of PKA and ERK1/2 in the same dose regime, we find that these kinases also respond biphasically to increasing cAMP levels, with opposite phases. We reveal that this behavior results from an elaborate integration by PKA of many cellular signals triggered by cAMP. In addition to the direct activation of PKA, cAMP also modulates the activity of p38 and ERK, which then converge to inhibit PKA. These interactions and their ensuing biphasic PKA profile have important physiological repercussions, influencing the ability of MDCKI cells to proliferate and form acini. Our data, supported by computational modeling, synthesize a set of network interconnections involving PKA and other important signaling pathways into a model that demonstrates how cells can capitalize on signal integration to create a diverse set of responses to cAMP concentration and produce complex input-output relationships.

## Introduction

Cyclic adenosine monophosphate (cAMP) is a second messenger that is an essential cellular currency in all kingdoms of life^1–3^. In mammalian systems, cAMP is the major regulator of Protein Kinase A (PKA), an important cellular kinase. Even small changes in PKA activity are known to exert regulation on a wide spectrum of cellular components, ranging from metabolic enzymes and cytoskeleton components to various transcription factors, such as the cAMP response element binding protein (CREB)^4–7^. PKA is a holoenzyme composed of two regulatory subunits (PKAr) and two catalytic subunits (PKAc). Upon binding of cAMP to the PKAr, all subunits dissociate, allowing PKAc to phosphorylate and regulate its targets^8^. Through this influence on PKA, cAMP controls a large swath of crucial cellular processes, such as proliferation, death and differentiation^3^.

Because of its physiological importance, the level of cAMP is subject to intricate regulation on its production by adenylyl cyclases (AC) and degradation by phosphodiesterases (PDE)^3^. It has been largely assumed that the activity of PKA correlates monotonically with the cAMP concentration. However, this simple view of the functional relationship between cAMP and PKA has recently been challenged. For example, it was demonstrated that the formation of memory in rodents shows a biphasic pattern that depends on cAMP level^9^. When the hippocampus of mice was microinjected with low doses of cAMP analogs, they recalled their learnt behavior better than control mice. However, when treated with high doses of cAMP analogs, mice were worse at retaining the learnt behavior. A similar physiological biphasic relationship was also noted between cAMP and the secretion rate of malpighi tubes of Drosophila^10^. When these organs were stimulated with low levels of cAMP, produced either with cAMP analogs or with an optogenetically-controlled AC, they showed an increase in secretion rate. By contrast, when the dose of cAMP was large, tubes had a secretion rate that was lower than unstimulated organs. Furthermore, recent data have documented that serotonin (5-HT), a hormone known to increase cAMP intracellular concentration^11^, promotes the biphasic activation and inactivation of PKA in mammary epithelial cells, and that these PKA patterns result in an increase and decrease of transepithelial electrical resistance (TEER), respectively^12^. Intriguingly, the same work also demonstrated that the stress related p38 MAPK (p38) is important for an overall inhibition of TEER, therefore suggesting that p38 and perhaps other MAPKs are involved in the regulation of PKA in a cAMP dependent fashion^12,13^. These data give hints of a complex relationship between PKA and cAMP in which multiple cellular pathways modulate the dependence of PKA on cAMP to generate a biphasic relationship that induces distinct phenotypes at different cAMP concentrations.

To quantitatively study the relationship between intracellular cAMP levels and their resulting effect on cellular state, we employed the bacterial light-activated adenylyl cyclase bPAC^14^. bPAC was isolated from the *Beggiatoa* bacterial genus, and is functional when expressed in other species such as yeast, mice, or human cells lines^15–17^, showing low activity in the dark and rapid activation upon exposure to blue light to specifically elevate intracellular cAMP^14^. Unlike environmental and chemical inputs, bPAC specifically controls cAMP levels without interference or crosstalk with other cellular variables. It therefore provided us with an essential tool to unambiguously probe the effects of cAMP dose-dependent transcriptional response and physiological outcomes.

We expressed bPAC in Madin-Darby Canine Kidney Type I (MDCKI) cells^18^ where the cAMP-PKA pathway has been shown to regulate acini formation^19^ so as to establish a meaningful physiological output. Many renal epithelial phenotypes, such as control of cyst size^20^, podocyte differentiation^21^, and fibrogenic programs in response to high glucose^22^ are also controlled by the cAMP-PKA pathway. Therefore, understanding the effects of increasing cAMP concentrations in renal epithelial models, such as MDCK cells, is essential for understanding the biology of normal tissues as well as the pathological tissues in diseases such as diabetic glomerulopathy, nephrotic syndrome or polycystic kidney disease.

Our investigations using bPAC reveal that an increasing cAMP dose leads to complex transcriptional responses where some genes are regulated monotonically by cAMP levels while others see biphasic expression phenotypes at low vs. high doses of intracellular cAMP. We identify transcription factors that bind genes upregulated in both programs, as well as programspecific transcription factors, such as CREB that binds genes upregulated in low cAMP. Biological processes enriched in these two programs suggest that different cAMP doses differentially regulate many important phenotypes, such as proliferation and kidney morphogenesis. In determining the signaling pathways underlying these transcriptional changes, we show that PKA activity is similarly biphasic, increasing and then decreasing as a function of cAMP. ERK activity exhibits the same biphasic behavior as a function of cAMP dose but in the opposite direction, decreasing and then increasing with increasing cAMP. By contrast, p38 shows a monotonic dependence on cAMP. We show that the PKA inhibition at high cAMP dose is not generated by the known feedback loops regulating directly the level of cAMP, for example by regulation through increased PDE activity. Instead, we uncover that biphasic PKA activity is the result of a network of interactions involving cross-regulation with ERK and p38. We further demonstrate that low versus high cAMP doses have important phenotypic consequences for cellular proliferation and the ability of cells to form acini structures.

Together, these findings call for updating our understanding of PKA activity as a faithful monotonically dependent reporter of cAMP dose. Instead, we replace this notion with one in which PKA activity can be finely tuned and modulated by other cAMP-responsive cellular pathways. This modulation has large phenotypic consequences, eliciting phenotypes reminiscent of involvement of these cells in health and disease of the kidney. Notably, our modeling suggests these relationships are heavily dependent on the relative concentrations and catalytic rates of PKA and MAPKs, enabling cell-type specific topologies of these signaling pathways and suggesting how multicellular organisms might develop tissue specific responses to changes in intracellular cAMP levels.

## Results

### bPAC-generated cAMP inputs elicit differential transcriptional responses

To quantitatively examine the effects of increasing cAMP inputs in MDCK cells, we expressed bPAC^14^, a blue light responsive cyclase that we targeted to the cytoplasm (MDCKI + bPAC). To test the ability of bPAC to rapidly produce cAMP in this cell line, we measured cAMP levels by ELISA in the presence and absence of blue light. In the absence of light, both MDCKI and MDCKI+bPAC had similarly low levels of cAMP (~0.1nM/ng protein, Figure 1A). After 20 minutes of continuous blue light exposure at maximum amplitude, only MDCKI+bPAC showed a large increase in intracellular cAMP (1318nM/ng protein, Figure 1A). To explore the range of intracellular cAMP concentrations that can be generated using bPAC, we further exposed MDCKI+bPAC cells to blue light with varying duty cycles for 20 minutes. To minimize light toxicity, we chose a period of 30s for blue light exposure in every cycle. Because of the fast kinetics of bPAC activation and shut-off, the increase in intracellular cAMP over time is continuous and not pulsatile. As expected, cAMP concentration increased with duty cycle (Figure 1B), reaching its maximum at continuous blue light exposure, demonstrating that bPAC can be used to generate a large range of precise cAMP levels spanning more than 3 orders of magnitude.

**Figure 1.**
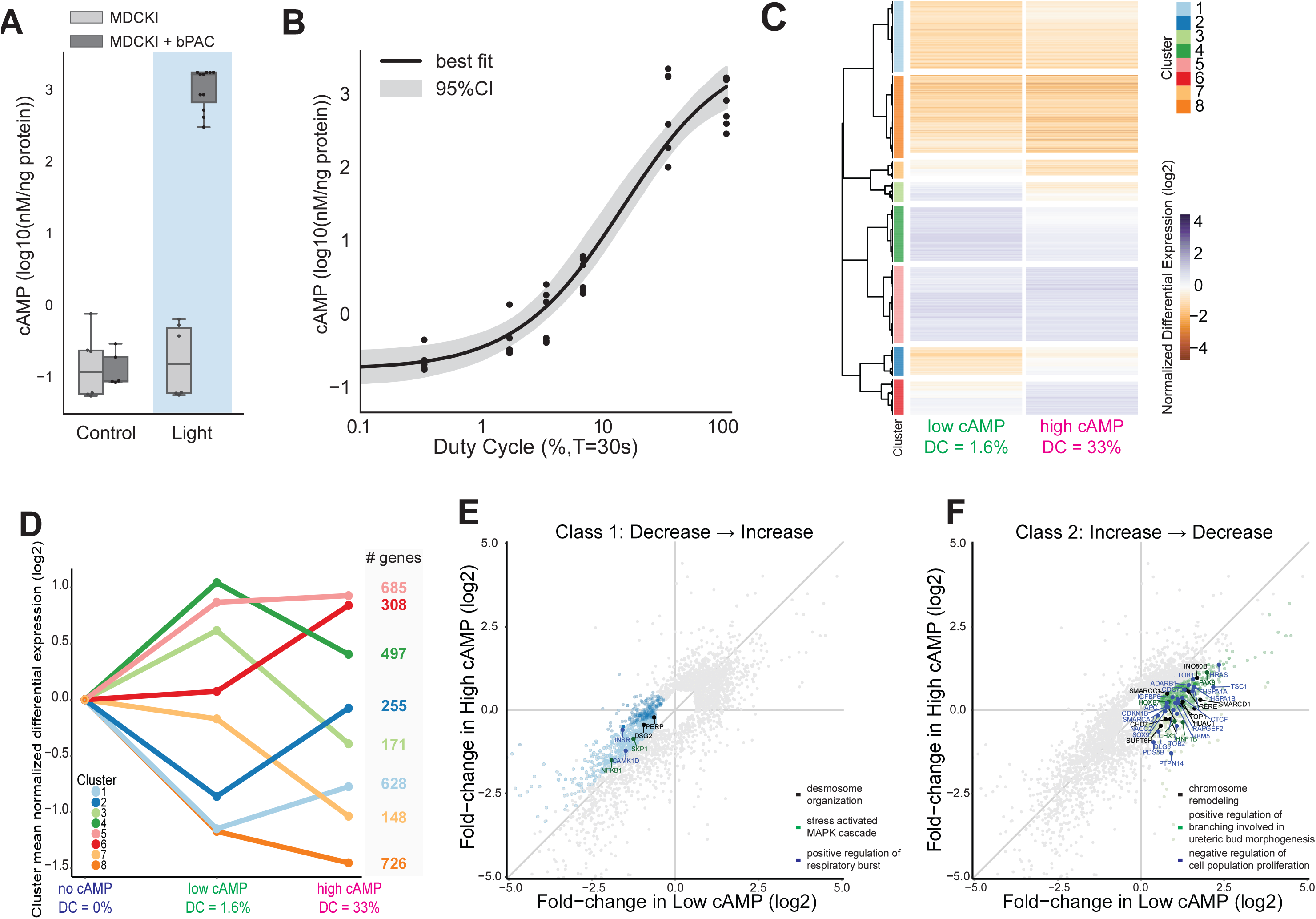
Different cAMP doses produce different transcriptional responses. **(A)** Intracellular cAMP concentration of MDCKI and MDCKI + bPAC in dark (Control) and after 20min of constant blue light exposure (Light). Box plots indicate distribution quartiles and whiskers indicate the rest of distribution. Data points from biological and technical replicates used to generate distributions are plotted. (B) Intracellular cAMP concentration of MDCKI + bPAC cells at 20 minutes of light treatment with different duty cycles for period equal to 30s (%, T=30s). Data points are shown in log scale, together with mean and 95% confidence interval of fit of a hyperbolic equation 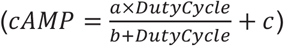 to data. **C**) Normalized expression level of differentially expressed genes (n = 3421) of MDCKI + bPAC cells after 2h of low and high cAMP doses (Duty Cycles of 1.7% and 33%, respectively). Hierarchical clustering of correlated gene expression identifies 8 clusters. (D) Mean expression of genes included in the clusters identified in (A) for different cAMP inputs. Total number of genes included in each cluster is indicated. (E) Scatter plot of all differentially expressed genes with their response in low cAMP dose versus high cAMP dose. Clusters 1 and 2, whose genes activity correlate with ERK1/2 activity are identified with color (light blue, Cluster 1; dark blue, Cluster 2). Selected genes for 3 GO terms enriched in Clusters 1 and 2 are identified: desmosome organization (GO:0002934, black), stress activated MAPK cascade (GO:0051403, green) and positive regulation of respiratory burst (GO:0060267, blue). (F) Scatter plot of all differentially expressed genes with their response in low cAMP dose versus high cAMP dose. Clusters 3 and 4, whose genes activity correlate with PKA activity are identified with color (light green, Cluster 3; dark green, Cluster 3). Selected genes for 4 GO terms enriched in Clusters 3 and 4 are identified: chromatin remodeling (GO:0006338, black), positive regulation of branching involved in ureteric bud morphogenesis and mesonephros development (GO:0090190 and GO:0001823, green) and negative regulation of cell population proliferation (GO:0008285, blue).

Increasing intracellular cAMP concentration is known to increase transcription via the transcription factors CREB, cAMP-responsive modulator (CREM), and activating transcription factor 1 (ATF-1)^3^. Increased cAMP levels have also been shown to have paradoxical effects on transcription downstream of the MAPK cascade, in some cases inhibiting it and in other cases activating it^23^. These divergent features of cAMP-regulated transcription are mediated by influencing the activities of many cellular pathways, most prominently PKA and ERK, which are kinases known to target and modulate the activity of a wide array of transcription factors^6,7^. Despite a qualitative understanding of these transcriptional dynamics, the quantitative features of cAMP transcription across a range of intracellular concentrations remain ill-defined. We hypothesized that because of the complex interacting nature of the protein networks mediating cAMP-dependent transcription, quantitatively defining the nature of cAMP dose-dependent transcription would further elucidate the topology of the signaling networks downstream of cAMP. To test this hypothesis, we carried out mRNA sequencing of MDCKI and MDCKI+bPAC cells under no, low, or high cAMP inputs (0%, 1.7% and 33% blue light duty cycle) for 2 hours.

To control for the effect of bPAC expression on cellular function in the absence of light, we identified genes that were differentially expressed in MDCKI + bPAC versus MDCKI parental cells when both were not exposed to light (duty cycle = 0%). Additionally, we measured the effects of blue light exposure on the MDCKI transcriptome by identifying genes that were differentially expressed in MDCKI parental cells exposed to low or high light inputs (1.7% or 33%) versus no light input (0%). Using a threshold for fold change in expression larger than 1.5 and an adjusted p-value smaller than 0.05, we identified 371 genes that were differentially expressed due to bPAC expression and in the absence of light (Supplementary Figure 1). These genes were enriched in Gene ontology^24^ (GO) terms related to cellular adhesion and mitosis. Reassuringly, these genes were not correlated with cAMP induced expression changes (see below, Supplementary Figure 1). We also identified 434 genes that were differentially regulated due to light exposure even in parental cell lines. These genes were removed from subsequent analyses.

Our aim was to capture all genes that were differentially expressed in response to light-induced cAMP inputs, and then classify them according to their patterns of cAMP dependence. Using the same thresholds as above (|FC|>1.5, p<0.05), we identified 3471 genes that are differentially expressed in response to one or both cAMP inputs, as well as genes that are differentially expressed between cAMP doses (Figure 1C, Supplementary Table 1). We used hierarchical clustering to group them according to their patterns. We identified 8 clusters that were further combined into *4* qualitatively distinct classes. Multiple clusters showed biphasic behavior as a function of cAMP, where gene expression levels showed opposite behavior at low and high cAMP doses. Genes in clusters 1 and 2 were strongly repressed in low cAMP and weakly repressed or activated in high cAMP (Decrease → Increase, Class 1). On the other hand, genes in clusters 3 and 4 were strongly activated in low cAMP and weakly activated or repressed in high cAMP dose (Increase → Decrease, Class 2). The biphasic pattern of Class 1 and Class 2 genes could be contrasted to genes in clusters 5 and 6 that were monotonically activated by cAMP (Class 3) and genes in clusters 7 and 8 that were monotonically repressed by cAMP (Class 4) (Figure 1C and D). We confirmed the RNA sequencing results for four cAMP regulated genes with qPCR experiments on independent RNA samples (Supplementary Figure 2).

To distill the various transcriptional programs corresponding to the different classes, we performed Enrichr analysis^25^ on genes from each class, focusing on transcription factors and kinases that regulate these genes and GO enrichment analysis to identify biological processes in which they are involved (Supplemental Figures 3 and 4). Genes across all clusters showed enrichment for binding of CREM, suggesting that all identified genes share a common dependency on cAMP for their expression. However, given the novelty of the biphasic responses we observed in Class 1 and 2 genes, we sought to focus on investigating their regulation.

Class 1 (Decrease → Increase) was specifically enriched for Vitamin D Receptor (VDR) and GA binding protein (GABP) binding (adjusted p-values 1.6×10^-14^ and 1.0×10^-1¤^, Supplementary Figure 3), which have been shown to have activity that is modulated by ERK^26^·^27^. This class was also enriched for genes that are upregulated upon depletion of the PKA-dependent CREB1^28^ (adjusted p-value 2.2×10^-13^, Supplementary Figure 3), and for genes that are upregulated in a protein kinase B (AKT1) knockout and downregulated in the presence of an AKT1 active mutant (adjusted p-values 2.1×10^-5^ and 1.8×10^-9^, Supplementary Figure 3). These data suggest that ERK and AKT1 might have their activities biphasically regulated by cAMP dose.

By contrast, Class 2 (Increase → Decrease) was enriched for genes that are known targets of CREB1 and the histone demethylase KDM5B (adjusted p-values 1.0×10-14 and 1.3×10-13, Supplementary Figure 3), an effect that is possibly mediated by PKA-driven regulation of chromatin remodelers and its activation of CREB1 transcription factor. In addition, we found a clear enrichment for genes that are known to be upregulated following knockout of the transcription factor jun-B (JUNB, adjusted p-value 1.2×10-13, Supplementary Figure 3), which is known to be activated by ERK as an immediate-early gene^29^.

Finally, we performed GO enrichment analysis for these two gene classes (Supplementary Figure 4). Class 1 (Decrease → Increase) was enriched in GO terms that encompassed desmosome organization, stress activated MAPK cascade and positive regulation of respiratory burst (Figure 1E, Supplementary Figure 4). By contrast, genes in class 2 (Increase → Decrease) had strong enrichment for terms related to chromatin remodeling, kidney morphogenesis and negative regulation of proliferation (Figure 1F, Supplementary Figure 4).

Based on these analyses, we hypothesized that ERK and other MAPKs are major drivers of the Decrease → Increase class, and that PKA is a driver of the Increase → Decrease class. We next sought to characterize if these putative signaling pathways had similar biphasic behavior across cAMP dose regimes.

### PKA and ERK show a biphasic response to bPAC-generated cAMP inputs in MDCK cells, while p38 shows a monotonic increase in activation

For the next exploration, we investigated the activity of PKA and ERK, given their prevalence in the regulation of the transcriptional response described above. We added to this panel the stress responsive MAPK p38 due to its overlapping regulation of some ERK-responsive genes in our dataset and previously demonstrated activation by cAMP in both PKA-dependent and independent manner^30^. Additionally, p38 has been shown to inhibit the PKA-dependent increase in TEER of epithelial cells^12^.

We first investigated the temporal dynamics of the cellular response to increasing cAMP dose, by exposing MDCKI+bPAC cells to low (duty cycle = 1.7%) and high (duty cycle = 33%) bPAC inputs lasting 90 minutes and measuring PKA, ERK, and phosphorylated CREB at different time points during light treatment by immunoblotting with antibodies that recognize the phosphorylated PKA substrates (pPKA Sub. (RRXS*/T*)), phosphorylated cAMP Regulatory Element Binding Protein (pCREB), and phosphorylated ERK1/2 (pERK) (Supplementary Figure 5). Parental MDCKI cells did not show any changes in PKA activity for this regimen of light exposure (Supplementary Figure 5A). By contrast, PKA in MDCKI + bPAC cells was responsive to light activation. Specifically, PKA activity increased in cells exposed to low cAMP, reaching its maximum between 10 and 20 minutes (Supplementary Figure 5A) and then declining to a new steady-state after that. PKA activity decreased when cells were exposed to high cAMP concentrations, with minimum activity reached at 5 minutes and maintained over the time course (Supplementary Figure 5A). This behavior was recapitulated for the PKA-dependent phosphorylation of CREB (Supplementary Figure 5B). Conversely, ERK was suppressed throughout the time course of low cAMP stimulation, but was activated with high cAMP dose (Supplementary Figure 5C).

To determine the full dose response of PKA, ERK, and p38 as a function of cAMP, we subjected MDCKI and MDCKI + bPAC cells to light inputs of different duty cycles, ranging from 0 to continuous exposure and quantified kinase activity by measuring the amount of phosphorylated PKA substrates, phosphorylated ERK, or phosphorylated p38 at 20 minutes, a time that was chosen because of the timescales of PKA activation and inactivation observed above. We identified that PKA activity increases with cAMP concentration and reaches its peak activity at ~1nM/mg protein (1.7% duty cycle). For cAMP concentrations above this value, PKA activity decreases, reaching values that are lower than its basal value for cAMP concentrations above 300nM/ng protein (Figure 2A, Supplementary Figure 6A). PKA activity, therefore, exhibits a markedly biphasic response to cAMP. ERK activity was the inverse of PKA’s and showed a similar biphasic nature, reaching its nadir at ~0.5nM cAMP/mg protein, and peaking above 100nM/ng protein (Figure 2B, Supplementary Figure 6B). In contrast to the biphasic responses of PKA and ERK, p38 showed a strictly monotonic increase in activation with increasing doses of cAMP (Figure 2C, Supplementary Figure 6C).

**Figure 2.**
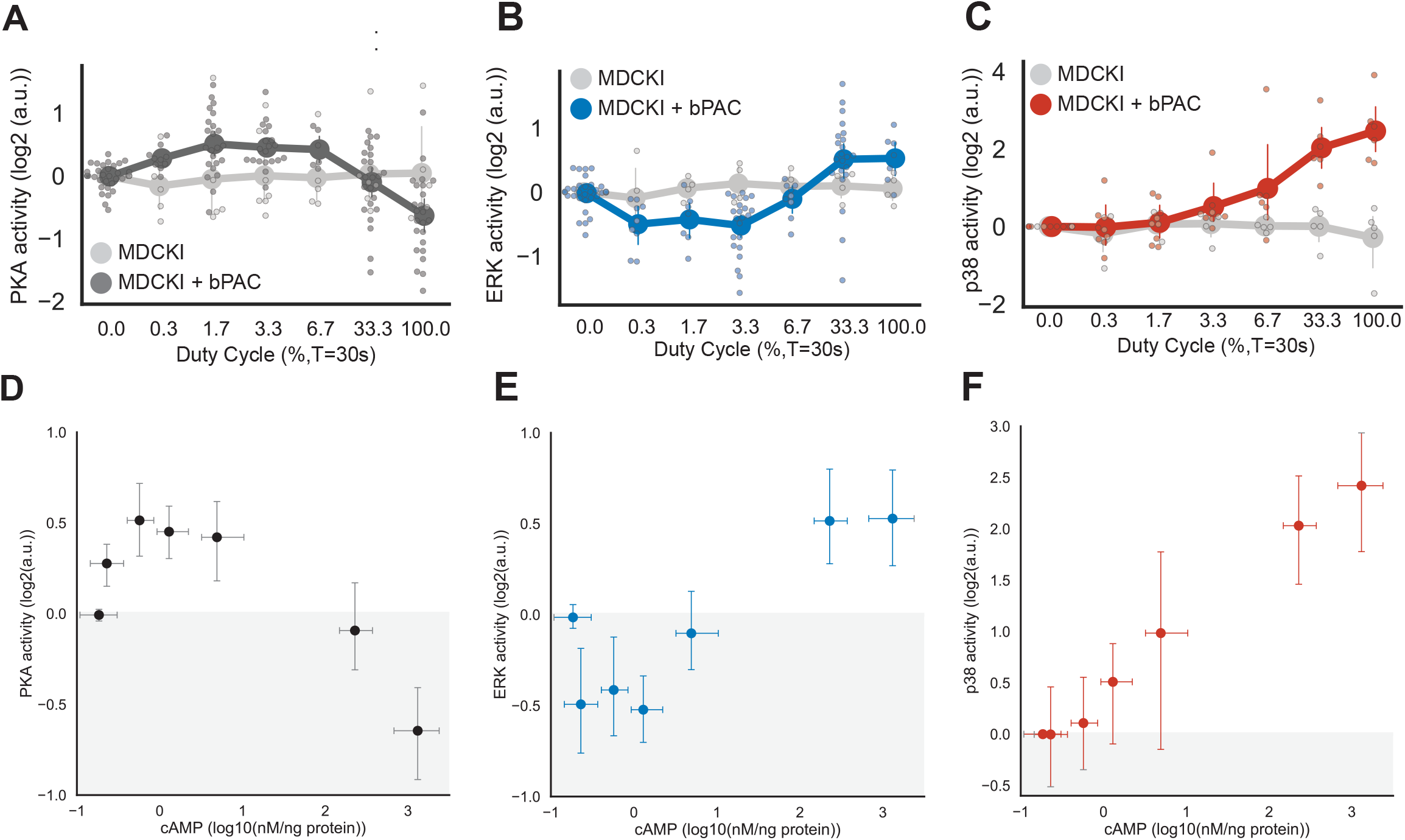
PKA and ERK have a biphasic response to cAMP, while p38 has a monotonic response. Quantification of immunoblots of **(A)** PKA activity (measured with Phospho-PKA substrate antibody), (B) of ERK activity (measured with Phospho-ERK antibody) and (C) of p38 activity (measured with Phospho-p38 antibody) of MDCKI and MDCKI+bPAC cells in response to 20min of blue light with different duty cycles. **(D)** Quantification of PKA activity, **(E)** ERK activity, and **(F)** p38 activity as a function cAMP concentration in MDCKI + bPAC cells. Error bars represent 95% confidence intervals of mean for at least 3 replicates.

Taken together, these data suggest that the biphasic genes identified from RNAseq in Class 1 (Decrease → Increase) followed the pattern of cAMP-dependent ERK activity, and the genes in Class 2 (Increase → Decrease) followed that of PKA activity.

### Inhibition of PKA by high cAMP dose is independent of PDE4 activity and of the dissociation of PKA holoenzyme

The direct molecular mechanisms of PKA’s cAMP-dependent activation are well documented, with cAMP binding to the PKA holoenzyme and causing the release of the catalytic subunits^31^. However, it is not understood how cAMP might inhibit PKA as we observe at high cAMP concentrations. We next sought to understand the mechanism for this dose-dependent inhibition. Previous investigations have described inactivation of PKA by two mechanisms involving feedback regulation that act primarily on cAMP. In the first mechanism, PKA activates PDEs that degrade cAMP thereby reducing PKA activity^32^. In the second mechanism, PKA promotes the desensitization of AC-activating G protein-coupled receptors (GPCRs)^33,34^. Since we were generating cAMP signals using an exogenous adenylyl cyclase, the second mechanism was less relevant. Because most cAMP degradation in MDCK cells is performed by the PDE4 family^35^, we set out to test if PDE4 activity is required for PKA inhibition with high cAMP doses. We grew MDCKI + bPAC cells in the presence of DMSO or Rolipram to inhibit PDE activity for 1h, exposed them to varying doses of light for 20min, and then collected extracts from these cells and probed their PKA activity using immunoblotting (Supplementary Figure 7). As expected, inhibition of PDE4s with Rolipram changed the input-output relationship from cAMP to PKA. PKA activity increased in the absence of light, and with cAMP inputs generated by duty cycles 0.3% and 1.7%, in cells exposed to Rolipram when compared to DMSO (Figure 3A, Supplemental Figure 7A). Notably, however, for duty cycles larger than 1.7%, Rolipram treatment lead to the sharp inhibition of PKA activity, suggesting that cells reached the threshold of cAMP concentration required for PKA inhibition with a smaller input of light due to the lack of cAMP degradation (Figure 3A, Supplemental Figure 7A). We therefore conclude that modulation of cAMP by PDE4 is not likely to be a major contributor to the biphasic relationship between PKA activity and cAMP.

**Figure 3.**
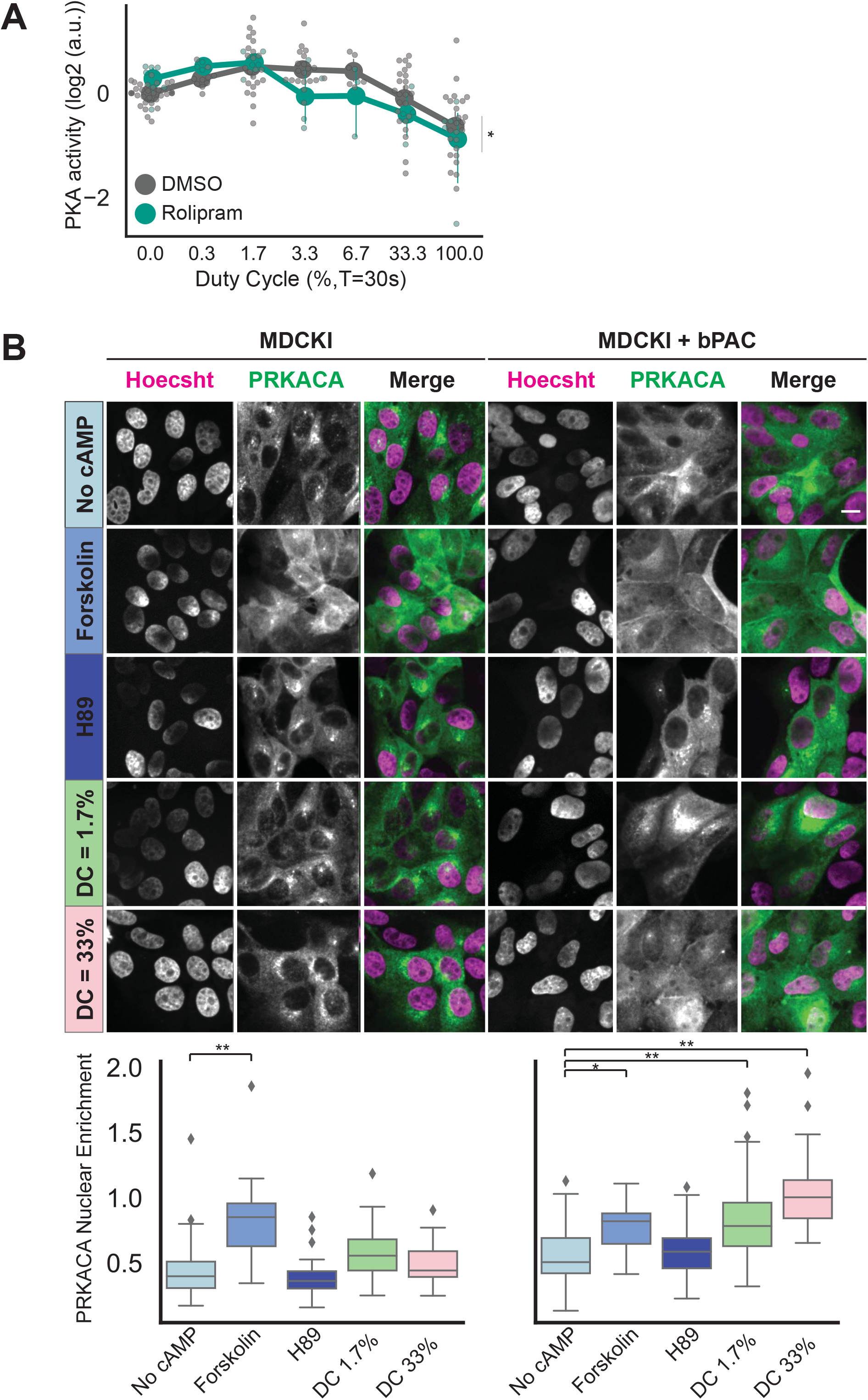
PKA inhibition is independent of PDE4 activity and association of PKA subunits. **(A)** Quantification of PKA activity (measured with Phospho-PKA substrate antibody) of MDCKI + bPAC cells treated with DMSO or PDE4 inhibitor (Rolipram (10μM)) and subsequently exposed to 20min of blue light with different duty cycles. * identifies p-value <0.05 in a two-way ANOVA test for the interaction between treatment and duty cycle. Error bars show 95% confidence intervals of mean for at least 3 replicates. **(B)** Images of fixed MDCKI and MDCKI + bPAC cells, expressing PRKACA fused to mRuby2 and stained with Hoechst 3342. Scale bar represents 10μm. Parental MDCKI and MDCKI + bPAC were treated with adenylyl cyclase agonist Forskolin (50μM) or with PKA inhibitor H89(10μM) or subjected to 2h of low (duty cycle = 1.7%) and high (duty cycle = 33.3%) cAMP doses. Quantification of PRKACA enrichment in nuclei (Nuclear PRKACA *I* Cytoplasmic PRKACA) is shown below. Box plot height indicates inter-quartile deviation. Median, minimum and maximum values of distribution are shown. * and ** identify p-values <0.05 and <0.01 in a Tukey-HSD multiple comparison after ANOVA. Quantification of each condition and genotype was performed in at least 20 cells.

Next, we tested whether this biphasic response is generated by molecular interactions of the PKA complex itself. Even though the holoenzyme itself can show activity^36^, one of the hallmarks of PKA activation by cAMP is the dissociation of its catalytic subunits from the regulatory subunits that are bound to A-kinase anchoring proteins (AKAPs)^5^. PKA inhibition is also usually associated with the reassociation of the holoenzyme. We therefore explored the state of the PKA holoenzyme under different regimes of cAMP input to determine whether the association of the PKA subunits is preserved at high cAMP concentrations. In order to visualize PKA localization in MDCKI cells, we expressed the catalytic subunit of human PKA (PRKACA) fused to mRuby2. We quantified nuclear enrichment of PRKACA-mRuby2 as a proxy for its dissociation from PKA regulatory units and AKAPs at the perinuclear region. As controls, we first quantified PRKACA-mRuby2 nuclear enrichment in cells exposed to Forskolin (adenylyl cyclase activator) or H89 (PKA inhibitor). As expected, PKA activation by Forskolin led to dissociation of PRKACA and increased its nuclear localization in both MDCKI and MDCKI + bPAC cells (Figure 3B). Due to the low basal activity of PKA, H89 treatment did not lead to a significant change in PRKACA localization, which remained mostly in the perinuclear region. Next, we investigated how bPAC-generated cAMP inputs changed the localization of PRKACA. To do so, we exposed MDCKI and MDCKI + bPAC cells to 20min of light with duty cycle 1.7% (low cAMP) or 33% (high cAMP), which increased or decreased PKA activity, respectively (Figure 2A). Light-induced low cAMP doses increased the nuclear fraction of PRKACA in bPAC-expressing cells to levels similar to those achieved by Forskolin treatment (Figure 3B). By contrast, the localization of PRKACA in the parental cell line did not change in either light conditions (Figure 3B). In MDCKI + bPAC cells exposed to the PKA-inhibiting high dose of cAMP, PRKACA nuclear localization increased to levels above those of low cAMP doses and those achieved following Forskolin treatment (Figure 3B). These data indicate that high levels of cAMP are not likely to inhibit PKA by directly increasing the reassociation of the holoenzyme in the perinuclear region.

We therefore next explored whether the inhibition of PKA at high level of cAMP is a result of network interactions, potentially involving MAPKs, that stem from cAMP and then converge on PKA activity, subjecting it to antagonistic activating and inhibiting actions.

### p38 and ERK activation are required for inhibition of PKA by high cAMP doses

Above, we demonstrated that PKA and ERK responded in an anti-correlated biphasic manner to increasing cAMP dose, whereas p38 showed a monotonically increasing response to cAMP. The anticorrelated biphasic relationship of PKA and ERK posed the natural hypothesis that they inhibited each other.

First, PKA has been implicated as a direct negative regulator of ERK activity and we hypothesized that ERK inhibition at low cAMP levels is the result of PKA activity^23^. In agreement, when we treated cells subjected to no, low or high light inputs with H89 (a PKA inhibitor^37^), ERK activity increased above that of control cells (Supplementary Figure 7B and C). Furthermore, the ERK activity of PKA-inhibited cells increased with duty cycle, suggesting that the ERK is activated by cAMP. Therefore, the biphasic behavior of ERK seems to be at least partially due to the antagonistic effects of activation by cAMP and inhibition by PKA.

Second, to investigate the potential of PKA inhibition by ERK, we treated MDCKI + bPAC cells with U0216 (MEK inhibitor), then exposed cells to the same light doses as above and measured PKA activity by immunoblotting (Figure 4A, Supplementary Figure 8A). Cells with and without MEK inhibitor had the same relative PKA activity at light inputs below 3.3% duty cycle (Figure 4A and B), indicating that ERK activity had little influence on PKA at low levels of cAMP. However, U0126 treated cells had a reduced PKA inhibition at larger light inputs, suggesting that ERK activity was only effective in this regime. Taken together with the data demonstrating that ERK activity decreased for low cAMP and increased again for high cAMP (Figure 2A and B), these data argue that the most parsimonious model explaining the relationship between PKA and ERK is one in which they counteract each other in a double negative interaction (Figure 4C). Intuitively, in this model, ERK activity is a balance of activation by cAMP and inhibition by PKA. Low levels of cAMP cause large PKA activity, whose inhibitory effect dominates causing ERK activity to decrease (Figure 2A and B, Supplementary Figure 6A and B). At a certain threshold of cAMP though, the balance of power is flipped and ERK activity starts increasing, therefore subjecting PKA to inhibition at high cAMP doses (Figure 4A and B).

**Figure 4.**
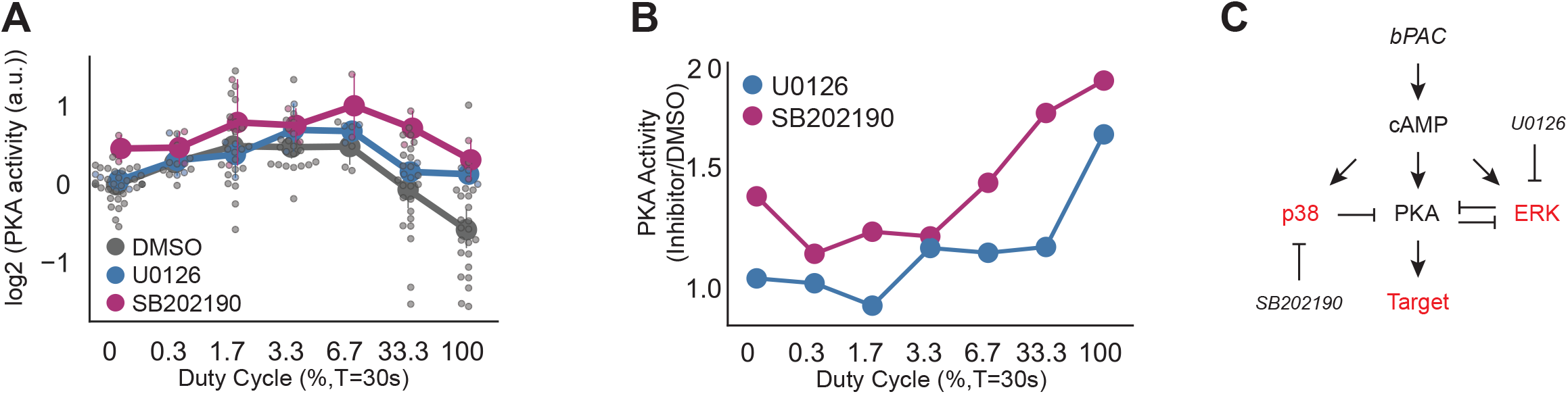
MAPK activation inhibits PKA activity at high cAMP doses. **(A)** Quantification of immunoblots of PKA activity (measured with Phospho-PKA substrate antibody) of MDCKI + bPAC cells treated with DMSO or MEK inhibitor (U0126 (10μM)) or p38 inhibitor (SB202190 (10μM)) and subsequently exposed to 20min of blue light with different duty cycles. Quantification of westerns was performed in at least 3 replicates. Error bars show 95% confidence intervals of mean. **(B)** Ratio of PKA activity in MDCKI+ bPAC cells treated with SB203580 and U0126 to those treated with DMSO when exposed to light inputs with different duty cycles for 20min. **(C)** Schematic of the postulated interactions between cAMP, PKA and MAPK. These interactions are represented in the computational model. bPAC-generated cAMP activates PKA, ERK and p38. Active PKA phosphorylates its targets and inhibits ERK. P38 and ERK inhibit PKA activity. SB202190 inhibits p38 activity and U0126 inhibits ERK activation. Experimentally measured variables are shown in red.

Although we demonstrated that ERK inhibits PKA at high cAMP doses, inhibition of ERK was insufficient to completely disinhibit PKA. Given p38’s monotonic increase in activity as a function of cAMP, we hypothesized that p38 activation might provide the other important inhibition of PKA at high doses. To investigate p38’s role on PKA activity, we treated MDCKI + bPAC cel Is with the p38 inhibitor SB202190, exposed them to varying doses of light for 20min and measured PKA activity by immunoblotting. Inhibition of p38 increased PKA activity in MDCKI + bPAC cells across all light treatments compared to cells treated with vehicle (DMSO) (Figure 4A, Supplementary Figure 8B), indicating that p38 exerts an inhibitory influence on PKA. This inhibition, however, was not uniform across all doses of light (that is, at different cAMP concentrations). Instead, the difference in PKA activity between DMSO and SB202190-treated cells was larger at 0 duty cycle, suggesting that p38 was active and was important for setting the basal activity of PKA. This difference then decreased between duty cycles of 0 and 3.3% and increased again with increasing duty cycles larger than 3.3% (Figure 4B).

Overall, the totality of our data so far suggests a molecular model in which cAMP affects the activity of all three kinases-PKA, p38, ERK-directly. In turn, these kinases seem to interact in elaborate ways to generate the biphasic PKA and ERK responses, as well as the monotonic p38 response to cAMP (Figure 4C). Nonetheless, further support that this qualitative picture is plausible can be provided by a computational model that captures the integrated dynamics of these interactions. Next, we build such a model.

### cAMP-dependent biphasic PKA activation can be explained through an incoherent feedforward loop involving ERK and p38

To capture the interactions in Figure 4C, we built a phenomenological mathematical model whose input is cAMP (different amounts produced by bPAC) and output is PKA activity (as a surrogate for the phosphorylation of its targets). In this model, cAMP activates PKA, p38 and ERK with different kinetics, and ERK and p38 inhibit PKA activity therefore mediating an incoherent feedforward interaction from cAMP onto PKA (Figure 4C, see methods and Supplementary Information). These interactions were modeled phenomenologically using enzymatic Hill functions in order to allow for the possibility that the interactions are not direct, but instead can have multiple intermediate steps. This functional form also allows for cooperativity and saturation in the substrate. In this form, the model was easily able to reproduce all the data from bPAC as well as kinase inhibition experiments. However, several structural features of the model, as well as choice of its parameters were essential for this.

With regard to the topology of the model, the inhibition of ERK by PKA and the dependence of both kinases on cAMP for their activation were vital for the biphasic behavior of ERK (Figure 5B) as a function of cAMP in DMSO-treated cells. While ERK inhibition of PKA was not required by the model to recapitulate the biphasic response of PKA to cAMP (as illustrated by the reduction in PKA activity at high doses of light in the presence of U0126, Figure 5A), this inhibition was essential to induce a reduction in PKA activity in the absence of p38 activity (Figure 5A). In other words, if ERK inhibition of PKA did not exist, PKA activity could never decrease when p38 is inhibited because no negative influences on PKA would exist. Second, the monotonic rise in p38 activity with increasing duty cycles (Figure 5C) indicated that no inhibition of p38 by PKA was needed, an insight further corroborated by the ability of the model to fully reproduce the data without any feedback from p38 onto PKA. However, a basal activity of p38 in the absence of the cAMP input was required in order to reproduce the basal increase in PKA activity upon inhibition of p38 by SB202190 (Figure 5A).

**Figure 5.**
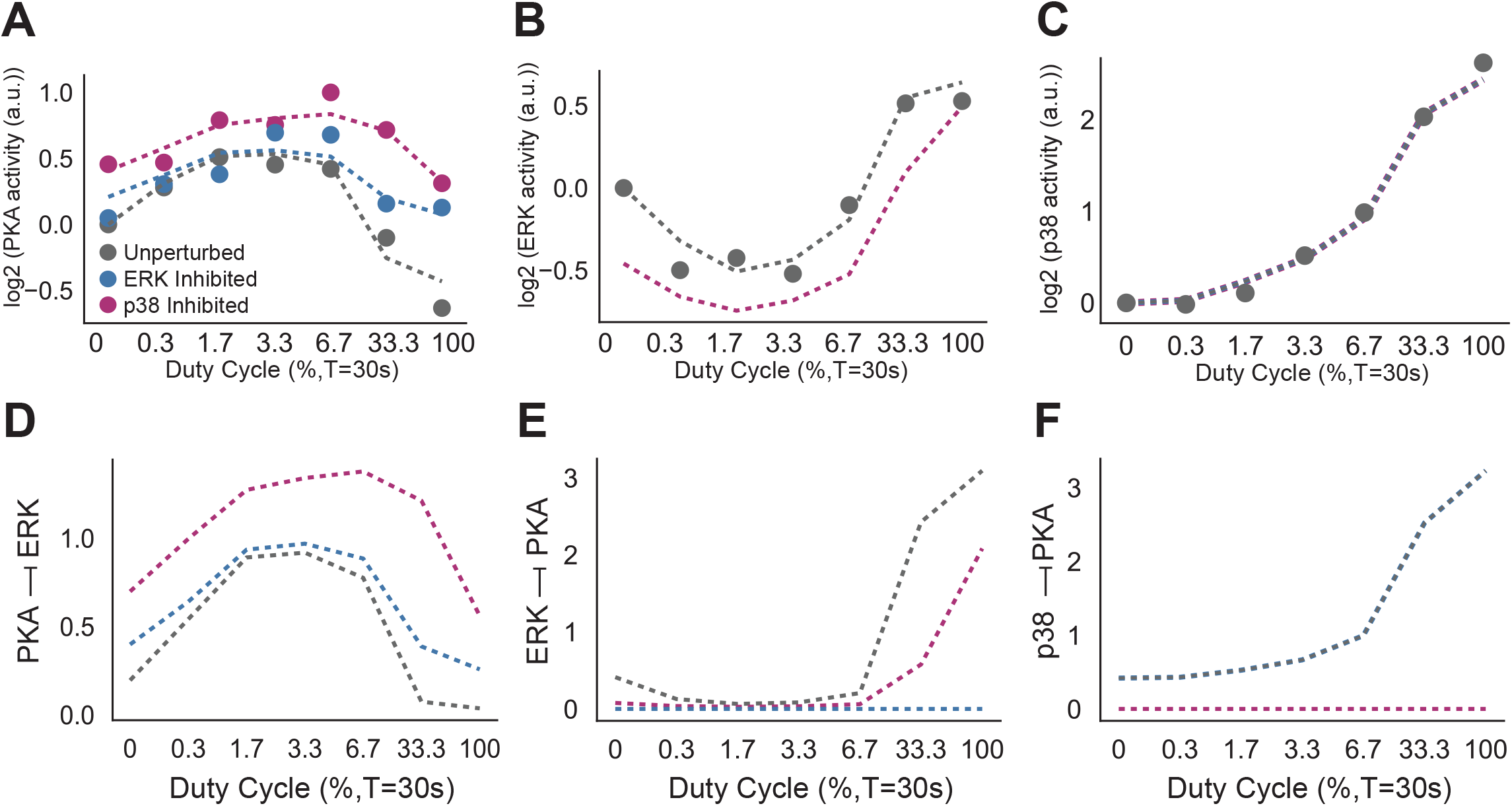
A phenomenological model describes the network conditions required for cAMP and MAPK regulation of PKA. For all graphs, blue denotes DMSO treated cells, orange denotes U0126 treated (ERK inhibited) cells, and pink denotes SB202190 treated (p38 inhibited) cells. Filled circles and solid lines represent experimental data, while dashed lines represent simulations. **(A)** Model simulation results of PKA activity of of MDCKI + bPAC cells treated with DMSO or MEK inhibitor (U0126 (10μM)) or p38 inhibitor (SB202190 (10μM)) and subsequently exposed to 20min of blue light with different duty cycles. **(B)** Model simulation results of ERK activity of MDCKI+bPAC cells treated with DMSO, SB202190 and U0126 and exposed to blue light with different duty cycles. **(C)** Model simulation results of p38 activity of MDCKI+bPAC cells treated with DMSO, SB202190 and U0126 and exposed to blue light with different duty cycles. (D) Magnitude of PKA inhibition of ERK predicted by computational model (PKA -| ERK) as a function of duty cycle for MDCKI+ bPAC cells in the presence of DMSO, U0126 or SB202190. PKA -| ERK was defined as 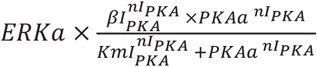 (see Supplementary Information for more detail). **(E)** Magnitude of ERK inhibition of PKA predicted by computational model (ERK -| PKA) as a function of duty cycle for MDCKI+ bPAC cells in the presence of DMSO, U0126 or SB202190. ERK -| PKA was defined as 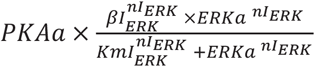 (see Supplementary Information for more detail). **(F)** Magnitude of p38 inhibition of PKA predicted by computation model (p38 -| PKA) as a function of duty cycle for MDCKI+ bPAC cells in the presence of DMSO, U0126 or SB202190. P38 -| PKA was defined as 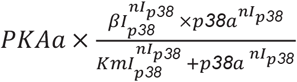 (see Supplementary Information for more detail).

In terms of parameter constraints, to fully represent the PKA phenotype upon p38 inhibition, the model required larger levels of cAMP in order to activate p38 and hence to trigger its negative effect on PKA, giving PKA activity the opportunity to increase at low cAMP levels (Figure 5A and D). This was achieved by a larger K_m_ for the activation of p38 by cAMP (Km_p38_) than that of PKA activation by cAMP (Km_PKA_) (~27x larger in simulations, Supplementary Information, Supplemental Figure 9A, B). When the K_m_’s of PKA and p38 were similar, the model achieved the activation of PKA by having a smaller Hill coefficient for the PKA activation (n_PKA_) than p38 (n_p38_) (Supplementary Figure 9 A, B and C). A different scenario in which Km_p38_ and Km_PK_ Awere similar but the Km of PKA inhibition by p38 (Kml_p38_) is large should be formally able to produce the same effect. However, such a parameter regime produces PKA activity that is the same in the absence of cAMP between SB202190-treated cells and DMSO treated cells, which is contradictory with our data (Figure 5A and D, Supplemental Figure 8).

Additionally, to reproduce the effects of ERK inhibition, the model required a larger K_mERK_ (K_m_ for the activation of ERK by cAMP) than K_mPKA_ (~32x larger, Supplementary Information, Supplemental Figure 9A, B) or a smaller n_PKA_ than n_ERK_ (Supplementary Figure 9 A, B and C), and a high Hill coefficient for the ERK inhibition of PKA (n_ERK_ = 5.37, Supplementary Information, Supplemental Figure 9A and E). These parameters were necessary to reproduce the small effect of ERK inhibition on PKA activity for small cAMP inputs (Figure 4A) and its strong effect at high cAMP inputs. Intuitively, at low cAMP concentrations, PKA inhibition of ERK dominates and hence ERK plays a small role in modulating PKA activity (Figure 5B and E). However, as cAMP concentration increases and ERK passes a predicted sharp threshold where its activity starts increasing again (representing an inflection point in the ERK activity, Figure 2B), it starts exerting a negative effect on PKA (Figures 5A and E). This is the regime where inhibition of ERK has a notable effect as indicated by the data.

Finally, the model predicted that the inhibitory effect of p38 on PKA should increase noticeably only after cAMP inputs larger than Km_p38_ (duty cycle 6.7 and larger) and that it should reach saturation for duty cycle 100% (Figure 5F). The saturation of this inhibitory effect was essential for reproducing the quantitative pattern of PKA inhibition in the presence of U0126, where the rate of inhibition of PKA seemed to decrease at high duty cycles (from 33% to 100%, Figure 5A). Since in this experiment, only p38 is present as an inhibitory influence on PKA, and since its level continues to increase at these high duty cycles, the fact that PKA activity assumes similar values for different duty cycles while cAMP increases indicates that the influence of p38 on PKA has to reach a saturation. This saturation effect should also be present in cells treated with DMSO, which was absent in the experimental data (Figure 5A), suggesting that additional interactions between the MAPKs that become important at high cAMP levels might also be present beyond the parsimonious model we have used.

Taken together, our data and computational modeling suggest that p38 activation modulates the activity of PKA through a cAMP-driven incoherent feedforward loop that plays an important role at all cAMP concentrations and that ERK engages in a double-negative feedback interaction whose importance seems to manifest at higher cAMP concentration. These interactions are integrated to produce a biphasic program of PKA activity for different cAMP concentrations. This program is recapitulated in gene expression phenotypes that follow PKA activity biphasically. Given these genetic phenotypes, as well as the roles of PKA and the MAPKs ERK and p38 as master regulators of physiology, we next turned to investigating whether this biphasic program has physiological I repercussions.

### Cellular proliferation is positively influenced by high cAMP inputs

Many genes in the PKA correlated clusters from the RNAseq (Class 2, Increase → Decrease) were associated with negative control of cellular proliferation, and these were repressed in the presence of high cAMP levels. This suggested that at a high cAMP dose, repression of these genes positions MDCKI cells to a state similar to cells transiting to a mesenchymal state^38^ or in elongating regions of epithelial tubes, which have weaker cell-cell interactions and higher rates of proliferation^39^’^40^. To test this possibility, we exposed parental MDCKI and MDCKI + bPAC cells to no, low or high cAMP doses (duty cycle of 0,1.7 and 33%) for 2 hours and measured their mitotic fraction by histone H3S10 phosphorylation after 14h. As expected, there were no significant changes in the mitotic fraction of MDCKI parental cells exposed to light. Yet, MDCKI + bPAC cells stimulated with a high cAMP dose had a marked ~5-fold increase in their proliferation rate when compared to unstimulated cells (Figure 6A). Cells exposed to the low cAMP input did not have a statistically significant change in the mitotic rate (~1.5-fold reduction in comparison to cells exposed to 0% duty cycle) (Figure 6A), indicating that only high cAMP treatment can significantly change the proliferation rate of MDCKI cells. These data support the notion that different doses of the same secondary messenger (cAMP), which result in different signaling and transcriptional patterns, can be decoded by cells to regulate proliferation in a differential way.

**Figure 6.**
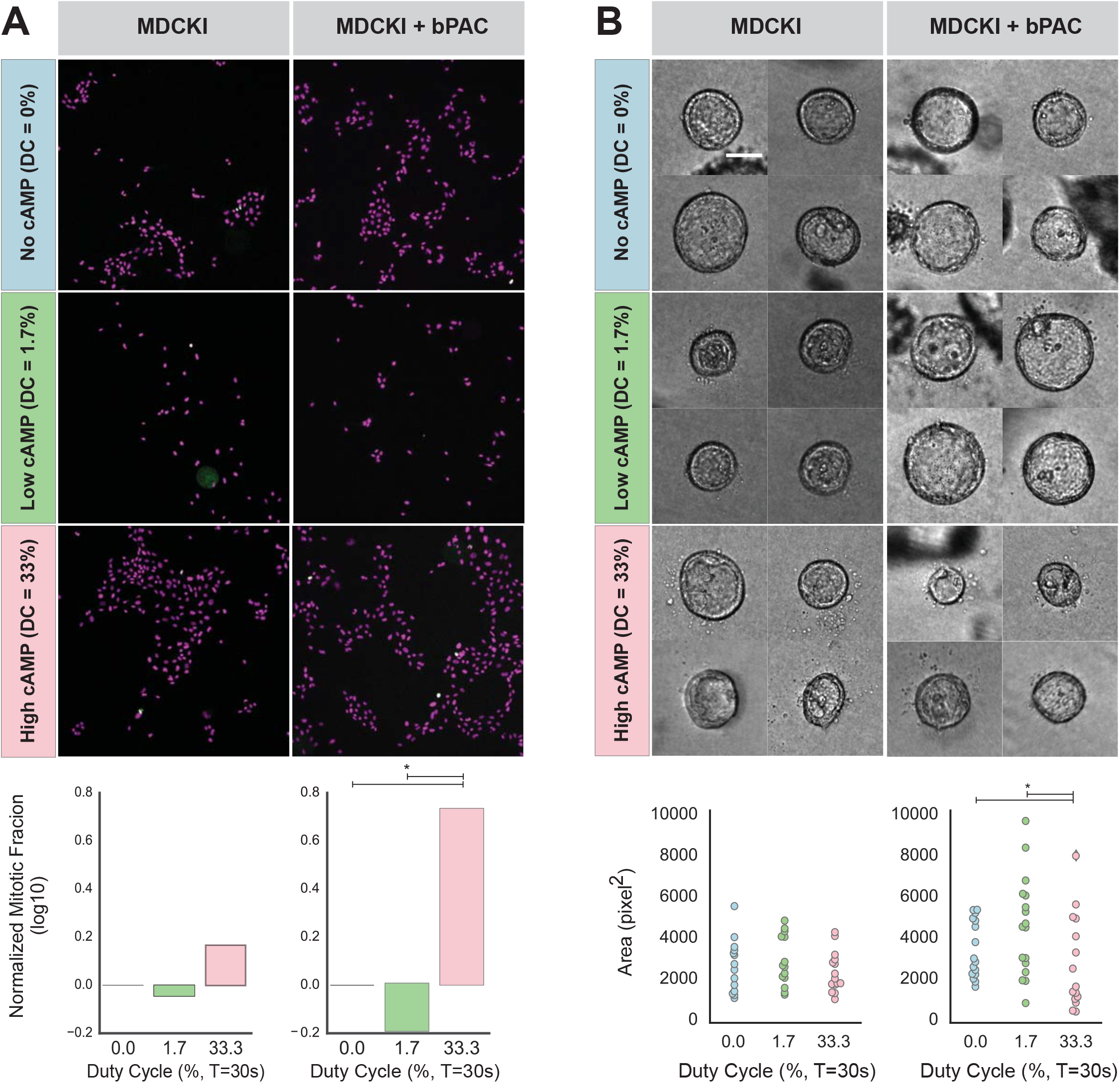
cAMP dose differentially regulates proliferation and acini growth. **(A)** Proliferation rate of cells through measurement of phosphorylation of histone H3S10. MDCKI and MDCKI + bPAC cells were subjected to 2h of low (duty cycle = 1.7%, green) and high (duty cycle = 33.3%, pink) cAMP doses. After 14h, phosphorylation of histone H3S10 was quantified by immunofluorescence. Images show cell nuclei stained with Hoechst 3342 (magenta) and histone H3S10 phosphorylation (green). Mitotic fractions, normalized to cells not subjected to cAMP are show at the bottom. * identifies a p-value <0.01 in Fisher’s exact test. **(B)** Images of acini structures. MDCKI and MDCKI + bPAC cells, grown in collagen I gel, were subjected to 2h of low (duty cycle = 1.7%, green) and high (duty cycle = 33.3%, pink) cAMP doses every 48h for 11 days. Acini were imaged at their center and the area quantified. Data points used to generate distributions are plotted below. * identifies a p-value <0.05 in Kolmogorov-Smirnov test.

### Formation of MDCKI acini is differentially controlled by cAMP dose

*We* noted that kidney developmental genes and cell adhesion genes belonged to the class of genes that correlate with PKA activity (Class 2, Figure 1F). In addition, both PKA and ERK signaling pathways have been shown to regulate different stages of kidney development^13–41^. Therefore, we predicted that different cAMP doses, through their biphasic influence on PKA and ERK, might influence the formation and maintenance of kidney structures differently. To examine this effect, we focused on acinar development of MDCKI cells. MDCK acinus formation is a well-studied model of kidney epithelial structure and function^42,43^. Correct formation and maintenance of MDCKI acini relies on the precise coordination of events that incorporate cell interactions with basement membrane, cell polarization, formation of tight junctions and dependence on an abundance of extracellular signals^19^.

It is known that PKA activity and ERK inhibition is required for the rapid formation of acini^44^. Therefore, we hypothesized that low cAMP doses and accompanying high PKA activity leads to the development of acini of normal size, while high doses of cAMP and associated ERK activation lead to increased cellular mobility and concomitant incorrect acinar development. To test this hypothesis, we quantified acinar size in MDCKI and MDCKI+bPAC cells grown in collagen gels exposed to no, low or high cAMP inputs for 10 days. MDCKI cells exposed to all light conditions showed no significant change in the area of their acini (Figure 6B), demonstrating that the effects of light are negligible on acini formation and maintenance. By contrast, cAMP dose had a striking effect on the area of bPAC-expressing acini (Figure 6B). Specifically, we identified that low and high cAMP inputs had a significant effect on the distribution of acini area, in which the low dose of cAMP increased the area of acini (median increase of 1.6-fold when compared to no cAMP), and acini exposed to high doses of cAMP showed a reduction in their area (median reduction of 1.7-fold when compared to no cAMP) (Figure 6B). These data illustrate that tight control of intracellular cAMP concentration and signaling activity is vital for the correct formation of multicellular structures, and that MDCKI cells can use cAMP concentration and its paradoxical effect on downstream pathways to generate an array of multicellular epithelial phenotypes.

## Discussion

In this work, we used a combination of precise optogenetic stimulation to control cAMP levels in living cells, with various cell biological readouts of its impact, to investigate the relationship between cAMP dose and the PKA signaling pathway. We uncovered that a variety of gene expression programs possess a biphasic response to intracellular cAMP dose, and that PKA and ERK activity correlated with these biphasic programs with PKA activity increasing and then decreasing as a function of cAMP and ERK activity showing the opposite behavior. While the dynamic range of PKA and ERK activity in their biphasic response to cAMP was 2-fold, the repercussions of this activity were substantial, both at the transcriptional and physiological level. Our mRNA sequencing results under two cAMP regimes (low and high) demonstrated the presence of two different molecular programs and physiological outcomes. Molecularly, we explored the intricate roles of p38 and the ERK pathways in producing the biphasic PKA phenotype and presented a data-supported model in which a poorly studied interaction between PKA and MAPKs inhibits PKA signaling pathway in a cAMP-dependent way. As a result, PKA is under positive and negative influences from cAMP, with the negative influence mediated by MAPKs. The interaction of these two modes of regulation, and their relative strength, induces a non-monotonic dependence of PKA on cAMP in the MDCKI cells we study here.

The biphasic response of PKA to cAMP has been previously observed in malpighi tubes of flies^10^, hypothalamus of mice^9^, and human mammary cells^12^ stimulated with small molecules. However, the non-monotonic behaviors reported in most of these earlier works resulted from chemical inputs that perturbed many cellular nodes other than solely cAMP. Our use of bPAC allowed us to circumvent this pleiotropy and uncouple feedback regulation on cAMP production from PKA activity. As a result, we could unambiguously link PKA activity solely to the amount of cAMP produced by bPAC and hence causally link cAMP concentration to the biphasic response of the PKA pathway. Additionally, this precise cAMP input afforded by bPAC enabled us to causally establish additional downstream targets of cAMP, including ERK, AKT and p38.

The biphasic nature of PKA and ERK responses to cAMP is implemented both by the incoherent feedforward loop (IFFL) motif from cAMP to PKA through the activities of p38 and ERK, and by the mutual inhibition of PKA and ERK. It is unknown how p38 and ERK produce their negative effect on PKA signaling. Our transcriptional dataset indicated that DUSPs are upregulated upon high cAMP doses, suggesting that they might mediate some of this interaction. However, further studies will be necessary to discover if their phosphatase activity is altered, if PKA and its targets are also targeted by these DUSPs, or if additional mechanisms are involved^45^. Furthermore, it is unclear how ERK and p38 are activated by cAMP. While previous studies have shown activation of MAPKs by cAMP through the activity of PKA^46^, our results suggest that, in MDCKI cells, additional cAMP-dependent mechanisms are involved. Previous results suggest that Epac, through activation of Rap1, can induce cAMP-dependent ERK activation^47^. Additionally, our data indicate that the nuclear fraction of PKAc increases with cAMP concentration (Figure 3B). If cytoplasmic PKAc is essential for the inhibition of ERK, then as cAMP and nuclear PKAc increase, more ERK can become active and reduce the activity of PKA. It is therefore possible that one or more of these mechanisms are responsible for the regulation of MAPKs at high cAMP doses.

Irrespective of the precise molecular mechanisms underlying cAMP effect on MAPKs and the effect of MAPKs on PKA, the presence of at least two IFFL structures in the system is at the heart of its biphasic behavior. An IFFL is a circuit motif that is usually assumed to respond to a step input with a transient response that then adapts close to its unstimulated state^48^. However, a less studied operational regime of IFFL is one in which they can give rise to dose-dependent biphasic responses^49–50^, such as the ones shown here. We explored these regimes in the context of the cAMP-PKA computational model and identified its specific parameter requirements. It is interesting to think that in other cell types, or even in disease states, changes in parameter values or network topology can originate radically different signaling responses out of the same molecular players. For example, if PKA is unable to inhibit ERK, ERK activity would increase monotonically with cAMP concentration. As a result, the signaling state of cells would be that of active PKA and ERK, instead of active PKA and inhibited ERK, at low cAMP concentration. It is far from clear what new functionalities or pathologies might ensue from these different signaling states. This is particularly important in light of the fact that the capacity of cAMP to coordinate the activity of important signaling pathways in MDCKI cells led to important changes in the expression of many genes. Remarkably, several of those genes that were cAMP responsive followed either a PKA or an ERK pattern, and correspondingly implemented two different programs. In the high cAMP program (repressed PKA and activated ERK) the proliferation rate of cells was increased and their ability to form acini reduced. By contrast, at a lower cAMP dose, proliferation rate was smaller than at a high cAMP dose and the size of acini was larger, in accordance with the previously described effects of PKA on acinar morphogenesis^44^. Interestingly, in Polycystic Kidney disease (PKD) cells, PKA is thought to activate MAPK and that activity of both pathways is required for the measured increase in proliferation rate and acinus size^20^. It is unclear if cells are producing abnormal cAMP doses or if the decoding of cAMP dose is deficient. While therapies for PKD have focused on reducing cAMP levels(53), if the network topology present in MDCKI cells is conserved in PKD cells, an alternative approach would be to increase cAMP, or directly activate p38, to inhibit PKA activity and reduce cystogenesis.

Finally, while the particular interactions in the cAMP-PKA-MAPK network we uncover here might be specific to epithelial cells, it is fascinating to think about how these links might be modulated, to shift the quantitative relationship between cAMP and PKA in different cell types. For example, a quantitative change in the inhibition of PKA by p38 or ERK might shift where PKA activity reaches its peak as a function of cAMP. A different change, balancing the timescale of the inhibiting and activating links on PKA, might abolish the biphasic relationship altogether. One can also imagine that some cell types might modulate these links dynamically, using inputs other than cAMP to impinge on the relationship of p38 and ERK on PKA, weakening or strengthening it based on the particular circumstances encoded in these inputs.

## Methods and Materials

### Cell culture

MDCKI cell line was a kind gift from Pavel Nedvestky. No mycoplasma contamination was found. MDCKI + bPAC cells were generated by infecting MDCKI cells with pLenti-PGK-bPAC::NES (Addgene #130267). PKAc expressing cells were obtained by transfecting MDCKI and MDCKI + bPAC cells with pPB-CAG-PRKACA::mRuby2 (Addgene #130268). Cells were maintained in MEM (Gibco #11095072) supplemented with 10% Fetal CalfSerum (VWR #89510-184) and 1X Anti-Anti (Gibco #15240062), and kept at 37 °C in a humidified incubator with 5% CO_2_.

### Antibodies and reagents

To probe signaling activity we used the following antibodies: anti-Phospho-PKA Substrate (RRXS*/T*) (100G7E) Rabbit mAb (Cell signaling technology #9624S, 1:2000), anti-Phospho-CREB (Ser133) (87G3) Rabbit mAb (Cell signaling technology # 9198S, 1:2000), anti-p44/42 MAPK (Erk1/2) (L34F12) Mouse mAb (Cell signaling technology ##4696, 1:2000), anti-Phospho-p38 MAPK (Thr180/Tyr182) (D3F9) XP^®^ Rabbit mAb (Cell signaling technology #4511, 1:2000), anti-αTubulin Antibody (DM1A) (Santa Cruz biotechnology #sc-32293, 1:2000), anti-Phospho-Akt (Thr308) (D25E6) XP Rabbit mAb (Cell signaling technology #13038S, 1:2000), anti-Phospho-Histone H3 (Ser10) Antibody (Cell signaling technology #9701, 1:500), IRDye 800CW Donkey anti-Rabbit (LI-COR #925-32213, 1:10000), IRDye 680RD Donkey antiMouse (LI-COR #925-68072, 1:10000) and Anti-rabbit IgG (H+L), F(ab’)2 Fragment (Alexa Fluor^®^ 488 Conjugate) (Cell Signaling Technology #4412, 1:1000).

The following small molecules were used: SB202190 (Selleck Chemicals, #S1077, 10μM), U0126 (MedChem Express #HY-12031, 10μM), rolipram ((R)-(-)-Rolipram, Tocris#l349, 100μM), H89 (H 89 dihydrochloride, Tocris#2910, 10μM) and Forskolin (Tocris# 1099/10, 50μM).

### Quantification of cAMP

1×10^6 cells were seeded in wells of 6-well plates (FALCON # 353046) in MEM without phenol red (Gibco # 51200038) supplemented with 0.5% Fetal Calf Serum (VWR #89510-184). After 16h, cells were exposed to no light or blue light (2.2 mW.cm^-2^) with different duty cycles for 20min. After light exposure cells were transferred to ice and cAMP concentration was determined using the cAMP ELISA Kit (Cell biolabs #STA-501), following the manufacturer’s instructions. To determine protein concentration, ELISA samples were assayed using the Pierce BCA Protein Assay Kit (Thermo Scientific Pierce #23227), following the manufacturer’s instructions. Measurements for cAMP ELISA and BCA Protein Assay kits were performed in a FlexStation 3 Multi-Mode Microplate Reader (Molecular Devices). The cAMP dose response to light was fit to a hyperbolic equation and confidence intervals were extracted by bootstrapping in a custom Python Script available on GitHub (https://qithub.com/jpfon/cAMP).

### Quantitative western blot

1×10^6 cells were seeded in wells of 6-well plates (FALCON # 353046) in MEM without phenol red (Gibco # 51200038) supplemented with 0.5% Fetal Calf Serum (VWR #89510-184). After 16h, cells were treated with inhibitors or vehicle (DMSO, SIGMA life science #D2650) for 1h. Cells were then exposed to no light or blue light (2.2 mW.cm^-2^) with different duty cycles for 20min. Cells were lysed in cold M-PER (Thermo Scientific Pierce #PI78501) supplemented with protease (Thermo Scientific Pierce #PIA32953) and phosphatase (Thermo Scientific Pierce #PI-88667) inhibitors for 20min. Proteins were separated in 7.5% Mini-PROTEAN^®^ TGX™ Precast Protein Gels (Biorad), probed with primary antibodies diluted in blocking buffer (Rockland Immunochemical #MB-070) for 16h and with secondary antibodies diluted in blocking buffer for 2h. Proteins were detected by fluorescence in a Odyssey CLx (LI-COR). Phospho-PKA Substrate quantification was performed for proteins above 75KDa. Tubulin was used as a loading control. Quantification was done using ImageJ^51^ and a custom Python Script available on GitHub (https://qithub.com/jpfon/cAMP).

### RNA library preparation and sequencing

1×10^6 cells were seeded in wells of 6-well plates (FALCON # 353046) in MEM without phenol red (Gibco # 51200038) supplemented with 0.5% Fetal Calf Serum (VWR #89510-184). After 16h, cells were exposed to no light or blue light (2.2 mW.cm^-2^) with different duty cycles for 2h. After light exposure cells were transferred to ice and total RNA was isolated using SPLIT RNA Extraction Kit (Lexogen # 008.48), following the manufacturer’s instructions. RNA quality was assessed using RNA 6000 Nano chips (Agilent # 5067-1512) in a Bioanalyzer 2100 (Agilent). RNA libraries for Illumina sequencing were prepared from 2ug of total RNA, using the QuantSeq 3’ mRNA-Seq Library Prep Kit FWD for Illumina (Lexogen # 015.96). Library quality was assessed using High Sensitivity DNA kit (Agilent # 5067-4626) in a Bioanalyzer 2100 (Agilent). Sequencing was performed at the Center for Advanced Technology (UCSF), in a HiSeq 4000 (Illumina). One lane was used to generate 50bp single reads. Sequencing data is available GEO (GSE134650)

### RNA sequencing analysis

Low quality reads and reads containing adapters were removed from raw data. Gene counts were generated using the standard workflow of Bluebee NGS Genomics Analysis Software (QuantSeq-FWD) for the dog genome (https://platform.bluebee.com). Differentially expressed (DE) genes were identified using the DESeq2 R package^52^, with |logFC| >1.5 adjp >0.05, in order to reduce the false discovery rate and genes whose expression change was small. Clustering of DE genes was performed using a custom R script available on GitHub (https://github.com/jpfon/cAMP). Gene Ontology term enrichment for genes in each cluster was performed using DAVID^24^ and displayed in a 2D scatterplot based on the semantic similarity (Revigo^53^). Transcription factor and kinase regulation enrichment analysis was performed using Enrichr^25^.

### Quantitative RT-PCR

Total RNA was prepared as described in the RNA library preparation section. First strand synthesis was performed on 5ug of RNA using Superscript II (Invitrogen #18064-014) and following the manufacturer’s instructions. qPCR was performed in a CFX connect (Bio Rad), using the PerfeCTa SYBR^®^ Green FastMix (Quantabio #95072) and following the manufacturer’s instructions for Fast 2-Step Cycling. GUSB expression was used as normalization factor. Primers used are shown in Supplementary Table 2.

### Quantification of mitotic fraction

4×10^4 cells were seeded in wells of 8-well slides (Ibidi # 80826) in MEM without phenol red (Gibco # 51200038) supplemented with 2% Fetal Calf Serum (VWR #89510-184). After 16h, cells were exposed to no light or blue light (2.2 mW/cm^2) with different duty cycles for 2h. After 14h, cells were fixed in 4% formaldehyde (Thermo Scientific Pierce #PI-28906) for 20min on ice, permeabilized with PBS-T (PBS (UCSF Cell Culture Facility) + 0.3% Tween-20 (Sigma-Aldrich #PI379)), blocked with normal goat serum (Abeam, #ab748l) for 1h, incubated with anti-H3S10p for 16h at 4oC and with secondary antibody for 3h at room temperature, and stained with Hoechst 33342 (Molecular Probes #H3570, 5μg/ml) for 10min. 3 PBS-T washes were performed between each step of the protocol, with a final PBS wash. Cells were imaged on a Nikon Ti inverted scope with arc-lamp illumination. Hoechst 33342 was detected using a 350/50nm excitation filter and a 460/50nm emission filter and H3S10p detected using a 470/40nm excitation filter and 525/50nm emission filter (Semrock, Rochester, NY). Cell nuclei were segmented and mitotic fraction was quantified using a custom python script available on github (https://qithub.com/jpfon/cAMP)

### Quantification of acinar area

MDCKI acini were prepared as described previously^54^. Over 11 days, acini were exposed to no, low (Duty Cycle = 1.7%) or high (Duty Cycle = 33%) blue light (2mW/cm^2) doses, for 1h of every 48h. Media was replenished at the same rate. Acini were imaged on a Nikon Ti inverted scope using brightfield. Acinar area was quantified using ImageJ (NIH).

### Imaging of PKAc and quantification of nuclear enrichment

1×10^5 cells were seeded in wells of 8-well slides (Ibidi # 80826) in MEM without phenol red (Gibco# 51200038) supplemented with 0.5% Fetal Calf Serum (VWR #89510-184). After 16h, cells were treated with small molecules or vehicle (DMSO, SIGMA life science #D265O) for 1h. Cells were then exposed to no light or blue light (2mW/cm^2) with different duty cycles for 20min to. Cells were immediately fixed in 4% formaldehyde (Thermo Scientific Pierce #PI-28906) and stained with Hoechst 33342 (5μg/ml). Cells were imaged on a Zeiss microscope equipped with a Yokagawa CSUX1-A1N-E confocal spinning disk. Images were collected with a 40 x 1.1 NA water immersion objective and Photometries Evolve 512 EMCCD camera. Hoechst 33342 and PRKACA::mRuby2 were detected using the following excitation lines and emission filters: excitation at 405nm, collection between 425 and 475nm for Hoechst 3342; excitation at 561nm, collection between 590 and 650nm for PRKACA::mRuby2. Segmentation of nuclei and quantification of PRKACA Nuclear to Cytoplasmic ratio were performed using a custom python script available on Github (https://qithub.com/jpfon/cAMP).

### Data availability

Raw data in this study is available at the UCSF repository (https://doi.org/10.7272/Q6ST7N00).

### Statistical analysis

Differences in mitotic fraction were calculated by Fischer’s exact test, in acini area by the Kolmogorov-Smirnov test, in PRKACA nuclear enrichment by one-way ANOVA with Tukey-HSD post-tests, and in the effects of Rolipram on PKA activity with a two-way ANOVA. All tests were implemented with Python scripts available on Github (https://qithub.com/jpfon/cAMP)

### Computational model

The computational model representing the cAMP-PKA signaling network is described in the Supplementary Information. Model simulations and optimization were performed with Python scripts available on Github (https://qithub.com/jpfon/cAMP)

## Acknowledgements

We thank Leonardo Morsut for the blue light LEDs and controllers; the El-Samad lab members, Marisa Rosa, Jeremy Chang, Roshanak Irannejad, Mark van Zastrow and Jeremy Reiter for critical feedback on the manuscript. This work was supported by the National Science Foundation through grant NSF-MCB 1715108 to HES

---

**Supplementary Figure 1.**
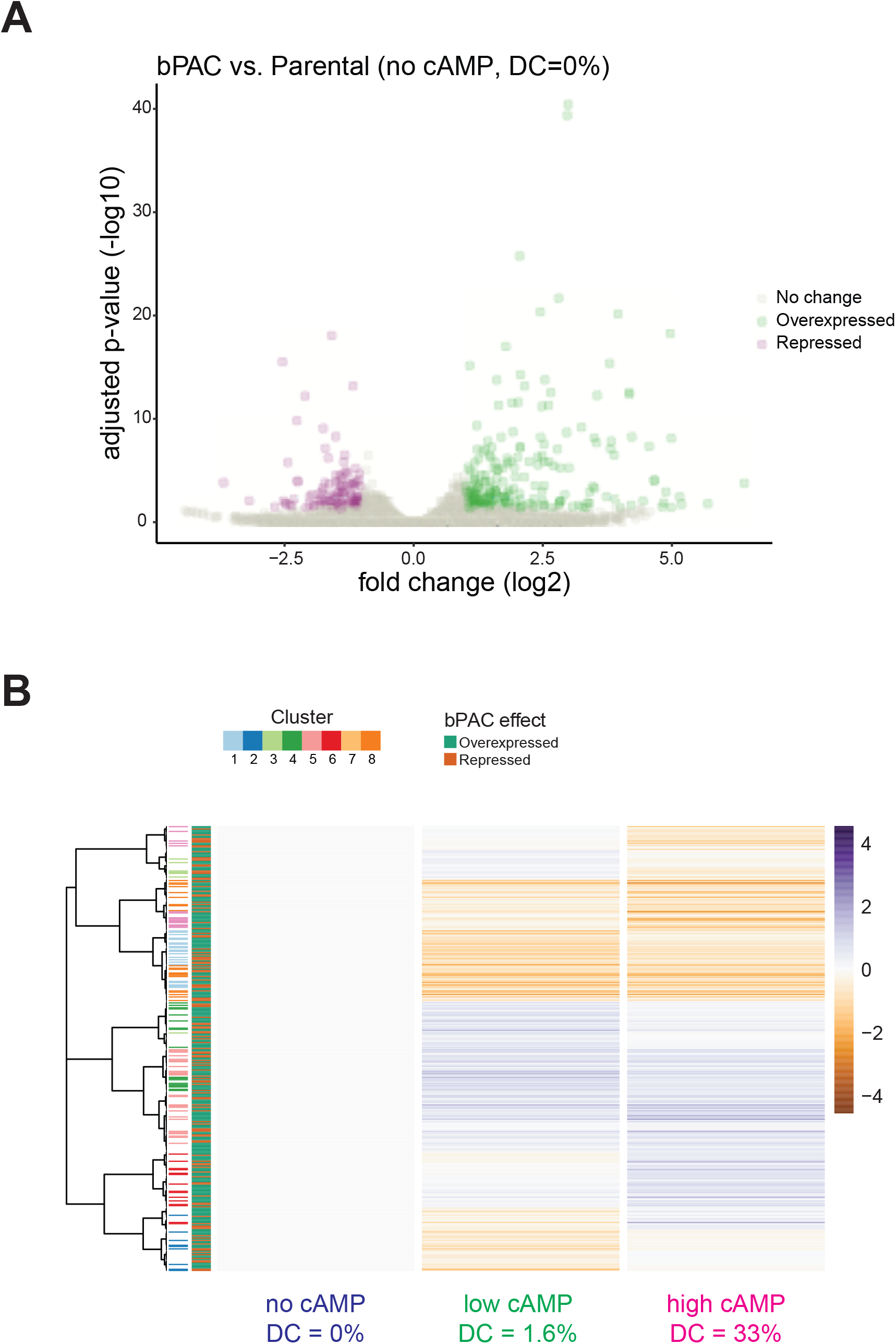
Effect of bPAC expression in MDCKI transcriptome. **(A)** Volcano plot identifying genes that are repressed (magenta) or upregulated (green) upon expression of bPAC in MDCKI cells. **(B)** Normalized expression level of genes regulated by bPAC presence (n = 371) in MDCKI + bPAC cells after 2h of low and high cAMP doses (Duty Cycles of 1.7% and 33%, respectively). Membership in clusters identified in Figure 1 is shown on the left (92 genes) as well as positive or negative effect of bPAC expression.

**Supplementary Figure 2.**
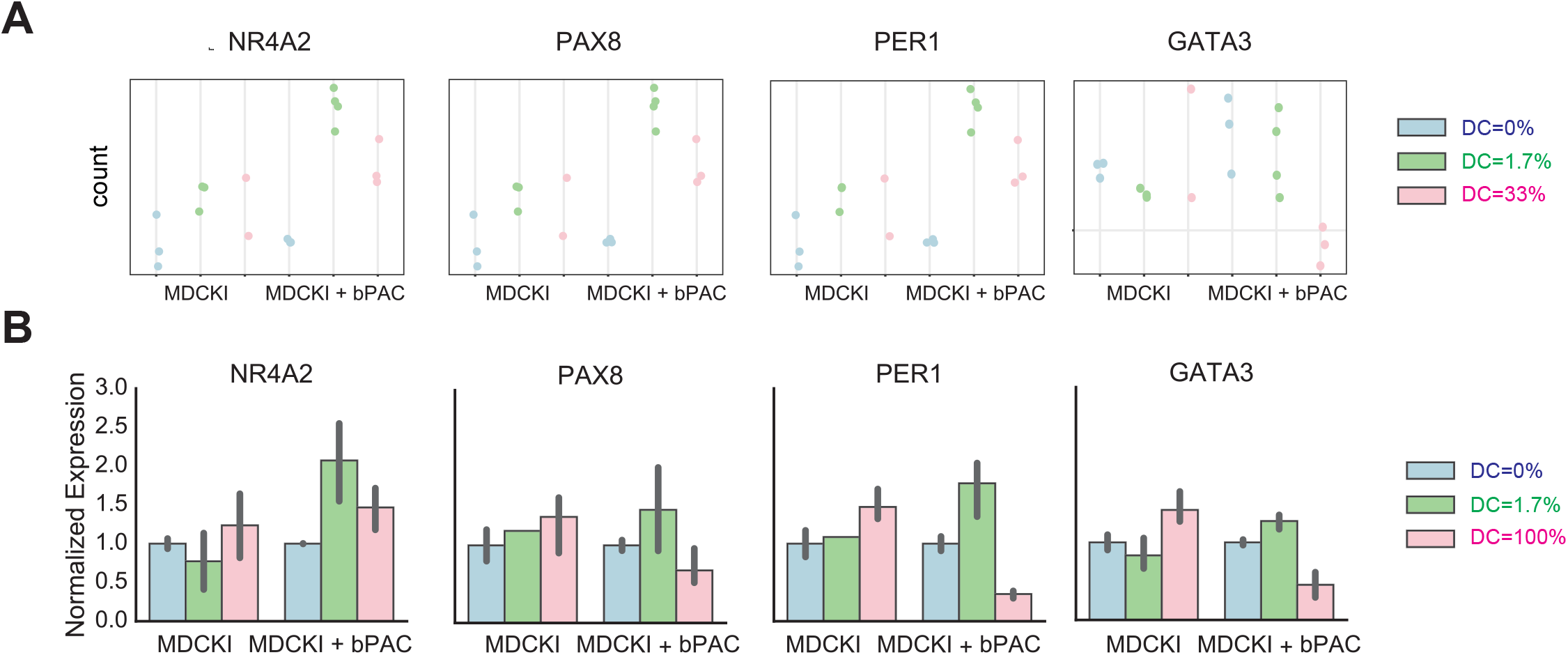
Validation of RNAseq results with qPCR. **(A)** RNA sequencing counts and **(B)** qPCR quantification for NR4A2, PAX8, PER1 and GATA3 genes in MDCKI and MDCKI + bPAC cells exposed to blue light for 2h with different duty cycles for period (T) equal to 30s (0%, 1.7% and 33 or 100%). qPCR was performed from at least 3 independent RNA isolations.

**Supplementary Figure 3.**
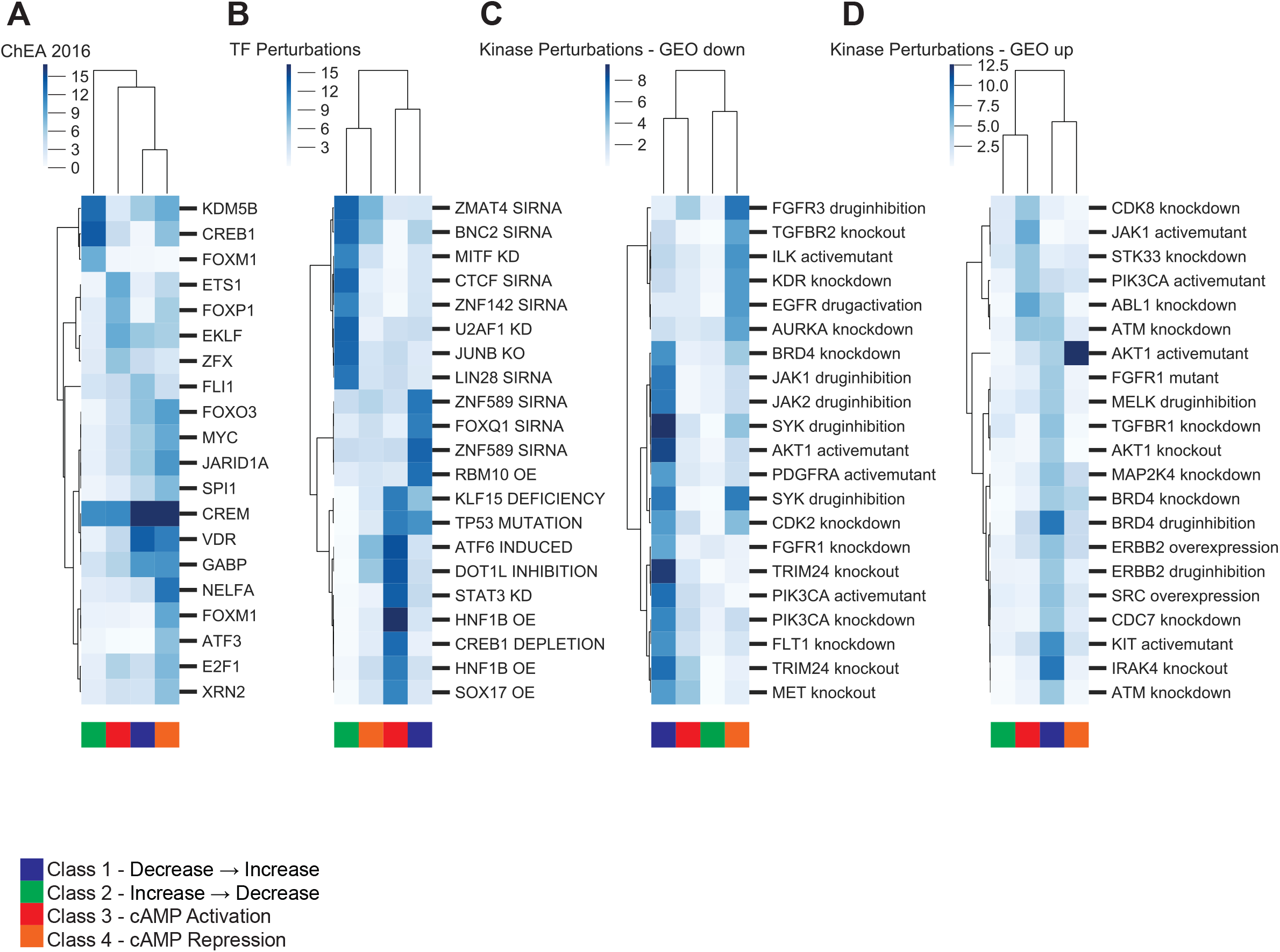
Transcription factor and Kinase Perturbation associated with differentially expressed genes. **(A)** Top twenty transcription factors from ChEA 2016 database in Enrichr whose binding is enriched in the promoters of the genes included in the clusters identified in Figure 1. (B) Top twenty transcription factor perturbations from the Transcription Factor Perturbation database in Enrichr whose transcriptome correlates to the genes included in the clusters identified in Figure 1. **(C)** Top twenty kinase perturbations from Kinase Perturbation – GEO down database in Enrichr tool whose downregulated genes correlate to the genes included in the clusters identified in Figure 4. **(D)** Top twenty kinase perturbations from Kinase Perturbation – GEO up database in Enrichr tool whose upregulated genes correlate to the genes included in the clusters identified in Figure 1.

**Supplementary Figure 4.**
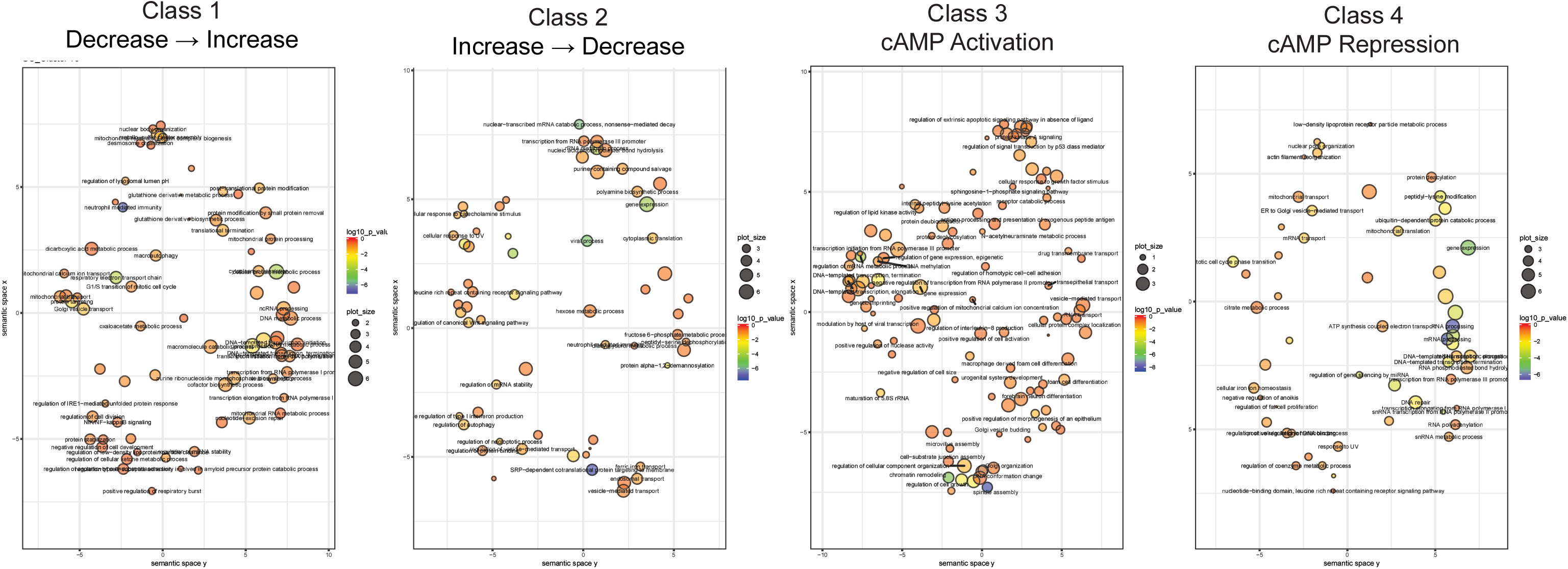
GO terms enriched in clusters of differentially expressed genes. **(A)** REVIGO plot of GO terms enriched in clusters 1 and 2, **(B)** clusters 3 and 4, **(C)** clusters 4 and 5 and **(D)** clusters 7 and 8. Clusters were identified in Figure 1.

**Supplementary Figure 5.**
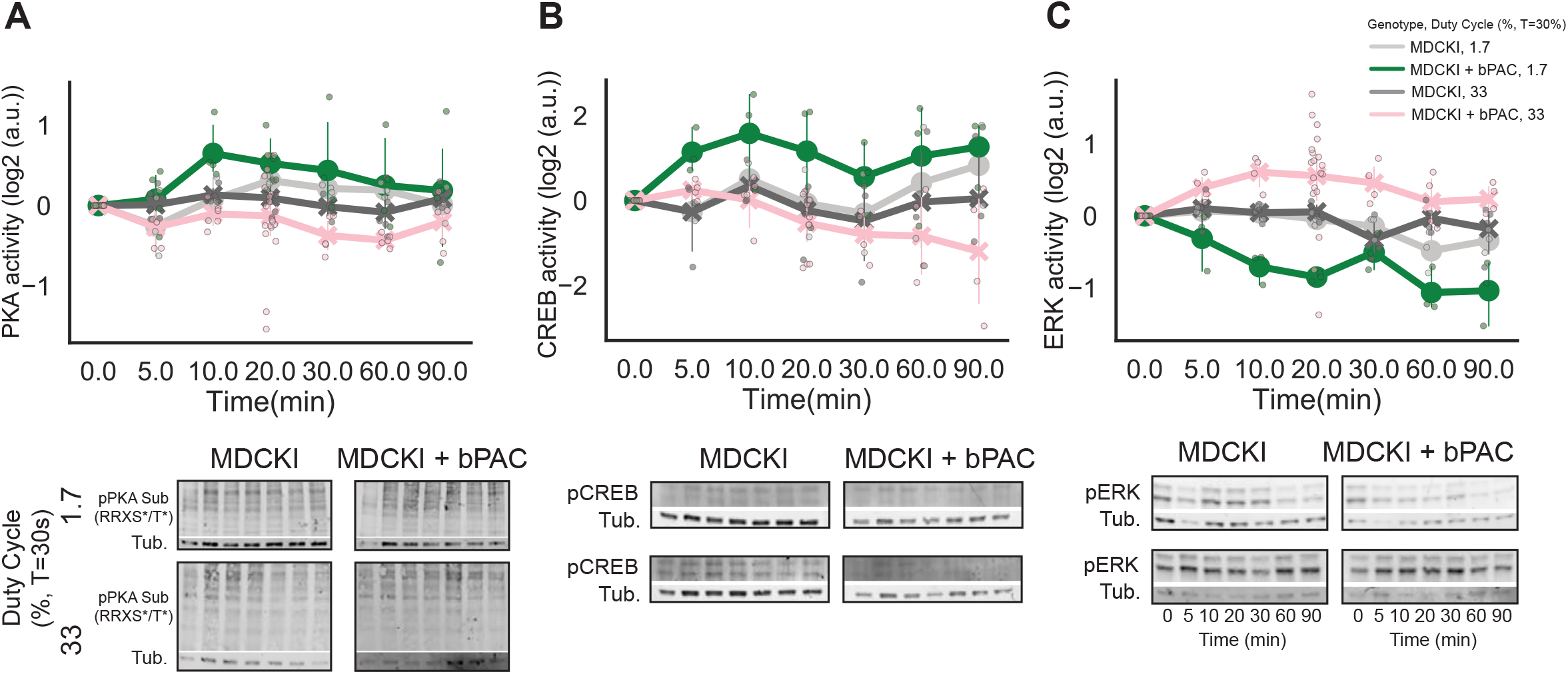
Time course of CREB, PKA and ERK1/2 activity of MDCKI and MDCKI + bPAC cells treated with two cAMP doses. Immunoblots of **(A)** PKA, **(B)** CREB and **(C)** ERK activity (measured with Phospho-PKA substrate (pPKASub. (RRXS*/T*)), Phospho-CREB (pCREB) and Phospho-ERK1/2 (pCREB) antibodies) of MDCKI and MDCKI + bPAC cells treated for 0, 5, 10, 20, 30, 60 and 90 minutes with low and high cAMP doses (Duty Cycle = 1.7% and 33%, respectively). Quantification of immunoblots below. For quantification, data points from replicates are plotted, together with mean and its 95% confidence intervals.

**Supplementary Figure 6.**
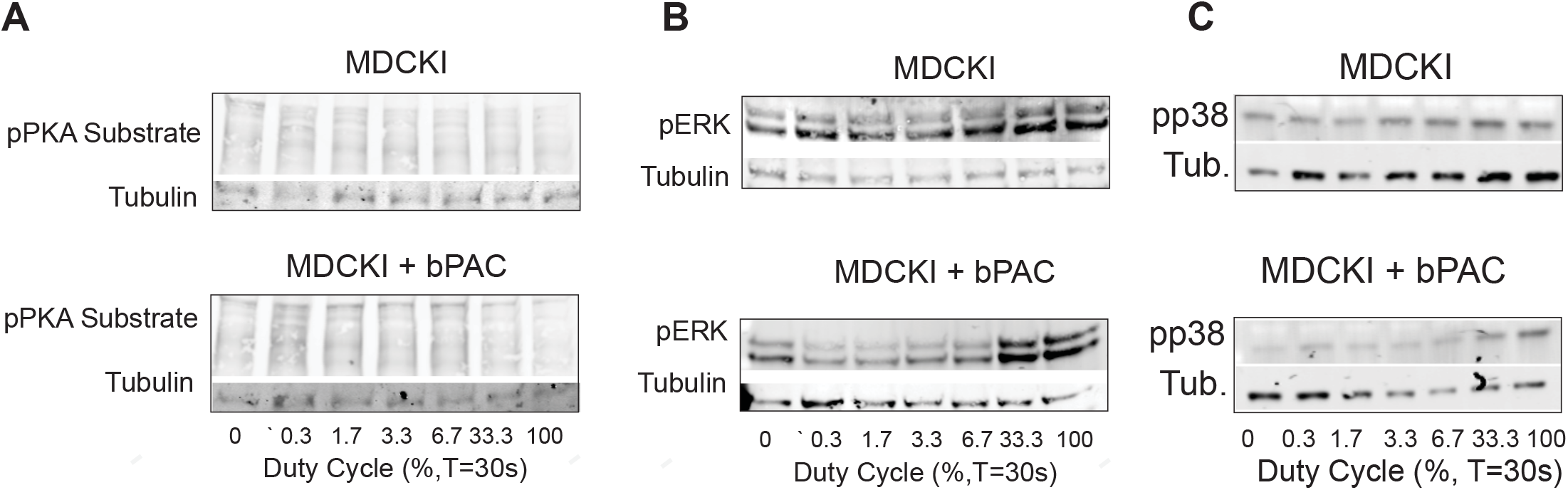
PKA, ERK, and p38 relationship to cAMP dose. Representative immunoblots of (A) phosphorylated PKA substrates above 100KDa (pPKA Sub. (RRXS*/T*)), **(B)** phosphorylated ERK1/2 (pERK), and **(C)** phosphorylated p38 of parental MDCKI and MDCKI + bPAC cells after 20min of light treatment with different duty cycles. Alpha-Tubulin (Tub.) was used for normalization.

**Supplementary Figure 7.**
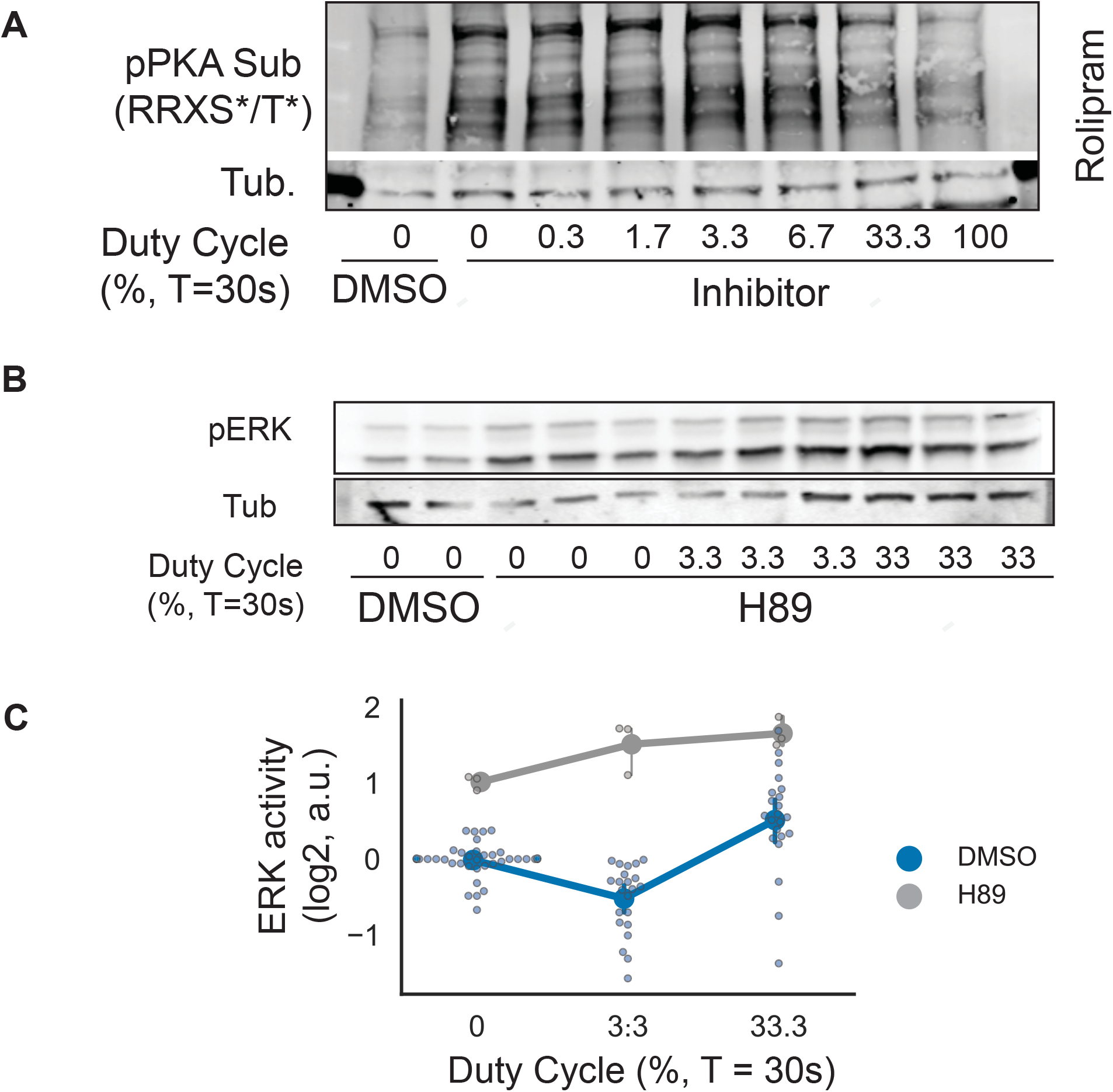
PKA regulation by PDE4 and ERK regulation by PKA. **(A)** Representative immunoblots of PKA activity (measured with Phospho-PKA substrate antibody) of MDCKI + bPAC cells treated with DMSO or PDE4 inhibitor (Rolipram (10μM)) and subsequently exposed to 20min of blue light with different duty cycles. **(B)** Immunoblots of ERK activity (measured with Phospho-ERK antibody) of MDCKI+ cells treated with DMSO or PKA inhibitor (H89 (10μM)) and subsequently exposed to 20min of blue light with different duty cycles. **(C)** Quantification of immunoblot shown in (B). Data points from replicates are plotted, together with mean and its 95% confidence intervals.

**Supplementary Figure 8.**
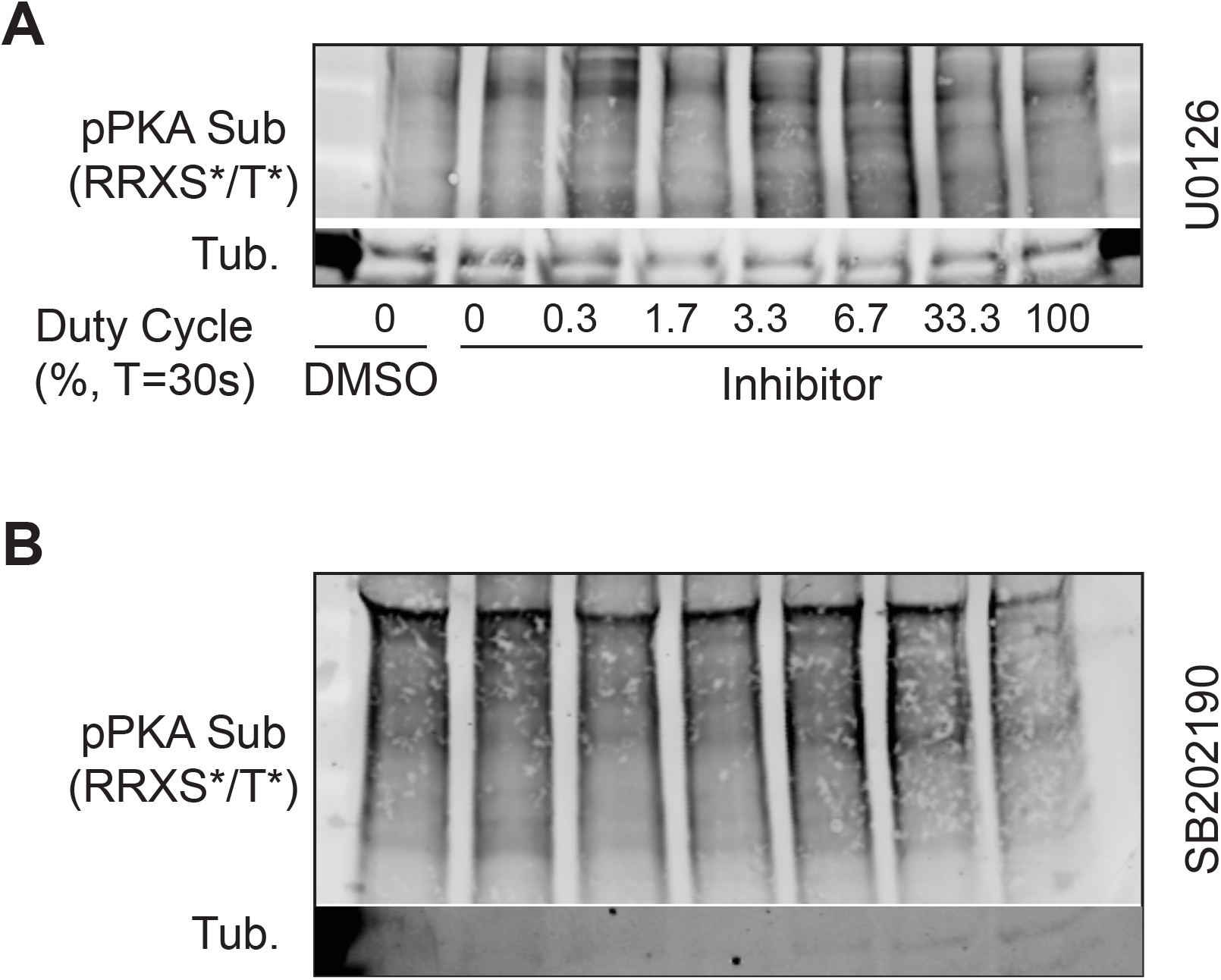
PKA activity of MDCKI + bPAC cells treated with light and different MAPKs inhibitors. Representative immunoblots of PKA activity (measured with Phospho-PKA substrate antibody) of MDCKI + bPAC cells treated with DMSO, **(A)** MEK inhibitor (U0126 (10 μM)) or **(B)** p38 inhibitor (SB203580 (10 μM)) and subsequently exposed to 20min of blue light with different duty cycles.

**Supplementary Figure 9.**
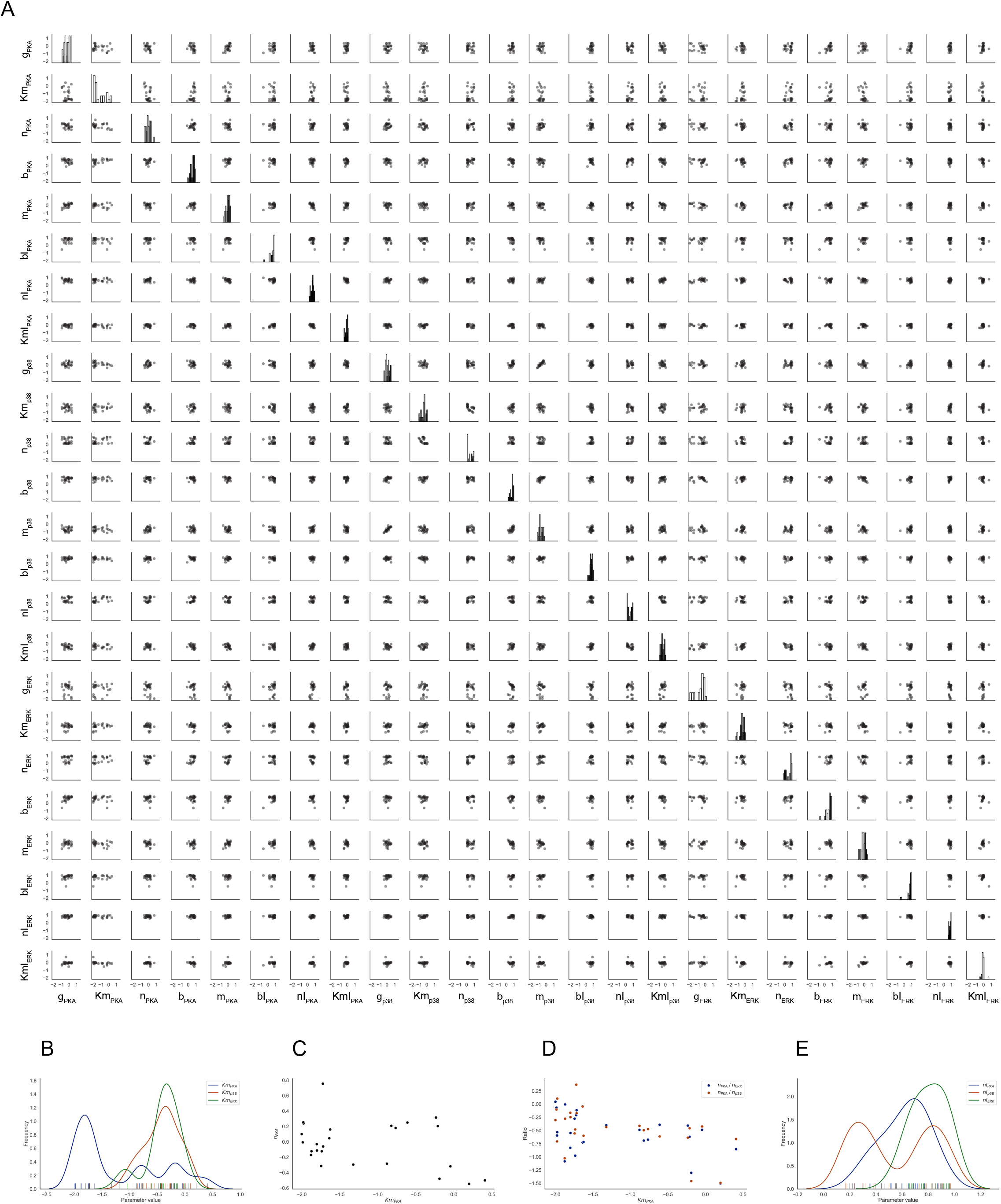
Distributions of cAMP-PKA model parameters. **(A)** Distributions (diagonal) and pair-wise joint distributions of parameter values in the cAMP-PKA computational model that were able to describe PKA, ERK and p38 activity in the presence of DMSO, and PKA in the presence of U0126 and SB202190. (B) Distributions of Km_PKA_ (blue), Km_ERK_ (green) and Km_P38_ (red). Most fitted Km_PKA_ parameters are smaller than Km_ERK_ and Km_p38_. **(C)** Relationship between Km_PKA_ and n_PKA_. For large Km_PKA_, n_PKA_ is always small. **(D)** Relationship between Km_PKA_, and the ratio of n_PKA_ to n_ERK_ (blue) or to n_p38_ (red). n_PKA_ is much smaller than n_ERK_ and n_p38_ for large Km_PKA_. (E) Distributions of nl_PKA_ (blue), nl_ERK_ (green) and nl_p38_ (red). Fitted nl_ERK_ parameters are always large. Detailed description of model and simulations in Supplementary Information.

Supplementary Table 1.

**List of differential expressed genes and their adjusted p-value, log fold change and cluster identity.** Related to Figure 1.

Supplementary Table 2.

**List of primers used for qPCR.** Related to Supplementary Figure 2.

Supplementary Information.

**Description of computational model and parameters used in simulations.** Related to Figure 5

