## Supplementary Information and Tables for "Biphasic Response of Protein Kinase A to Cyclic Adenosine Monophosphate Triggers Distinct Epithelial Phenotypes"

### Supplementary Information for *Biphasic response of Protein Kinase A to cyclic adenosine monophosphate triggers distinct epithelial phenotypes*

#### 1 Computational model description

An ordinary differential equation (ODE) model of the cAMP-PKA pathway was built to capture the interactions between cAMP, PKA, p38 and ERK. We described the activation of PKA, p38 and ERK by cAMP and the inhibition reactions of p38 and ERK on PKA, and of PKA on ERK using enzymatic Michaelis-Menten functional forms with saturation of the substrate, but these relationships were allowed to assume higher Hill coefficients in order to allow for the possibility that these interactions were not direct. In total, the model has 6 variables and 24 parameters described in table 1. In addition, cAMP ranged from 0 to 1 (in order to mimic the duty cycles used in other experiments). For the simulation of ERK inhibition with U0126 experiments,basal activation of ERK ($\mu_{ERK}$) and the maxium velocity of ERK activation by cAMP ($\beta_{ERK}$) were set to 0. For the simulation of p38 inhibition with SB202190 experiments, the maxium velocity of PKA inactivation by p38 ($_{p38}$) was set to 0.
In order to find the model parameters that were able to reproduce the data, Latin hypercube sampling of 10000 parameter sets was performed, and simulations with 7 cAMP inputs (0, 0.003, 0.017, 0.033, 0.067, 0.33, 1) were run in DMSO, U0126 and SB202190 conditions. 25 parameter sets that showed (i) biphasic behavior of PKA and ERK, (ii) monotonic activation p38, (iii) alleviation of PKA repression by SB202190 with all cAMP inputs and (iv) greater alleviation of PKA repression by U0126 at high cAMP inputs than at low cAMP inputs were further optimized using a non-linear least squares optimization scheme. One of this optimized parameter set is shown in table 3

#### 2 Computational model variables

In total,there are 6 state variables for PKA, p38 and ERK in active ($_a$) and inactive form ($_i$). Additionally, there are 24 parameters, shown in table 1

Table 1: Description of parameters for used in model

| Parameter | Description |
|---|---|
| $\gamma_{PKA}$ | Basal inactivation of PKAa |
| $\mathrm{Km}_{PKA}$ | Michaelis constant of PKA activation by cAMP |
| $\mathrm{n}_{PKA}$ | Hill coefficient of PKA activation by cAMP |
| $\beta_{PKA}$ | Maximum velocity of PKA activation by cAMP |
| $\mu_{PKA}$ | Basal activation of PKAa |
| $\beta I_{PKA}$ | Maximum velocity of ERK inactivation by PKA |
| $\mathrm{nI}_{PKA}$ | Hill coefficient of ERK inactivation by PKA |
| $\mathrm{KmI}_{PKA}$ | Michaelis constant of ERK inactivation by PKA |
| $\gamma_{p38}$ | Basal inactivation of p38 |
| $\mathrm{Km}_{p38}$ | Michaelis constant of p38 activation by cAMP |
| $\mathrm{n}_{p38}$ | Hill coefficient of p38 activation by cAMP |
| $\beta_{p38}$ | Maximum velocity of p38 activation by cAMP |
| $\mu_{p38}$ | Basal activation of p38 |
| $\beta I_{p38}$ | Maximum velocity of PKA inactivation by p38 |
| $\mathrm{nI}_{p38}$ | Hill coefficient of PKA inactivation by p38 |
| $\mathrm{KmI}_{p38}$ | Michaelis constant of PKA inactivation by p38 |
| $\gamma_{ERK}$ | Basal inactivation of ERK |
| $\mathrm{Km}_{ERK}$ | Michaelis constant of ERK activation by cAMP |
| $\mathrm{n}_{ERK}$ | Hill coefficient of ERK activation by cAMP |
| $\beta_{ERK}$ | Maximum velocity of ERK activation by cAMP |
| $\mu_{ERK}$ | Basal activation of ERK |
| $\beta I_{ERK}$ | Maximum velocity of PKA inactivation by ERK |
| $\mathrm{nI}_{ERK}$ | Hill coefficient of PKA inactivation by ERK |
| $\mathrm{KmI}_{ERK}$ | Michaelis constant of PKA inactivation by ERK |

### 3 Computational model ODEs

The reactions described in the model are shown below and summarized in table 2

Table 2: Model ODEs

| Description | Equation |
|---|---|
| Change in active PKA over time | (1) |
| Change in inactive PKA over time | (2) |
| Change in active p38 over time | (3) |
| Change in inactive p38 over time | (4) |
| Change in active ERK over time | (5) |
| Change in inactive ERK over time | (6) |

$$\frac{dPKAa}{dt} = PKAi * (\mu_{PKA} + \beta_{PKA} * \frac{cAMP^{n_{PKA}}}{cAMP^{n_{PKA}} + Km_{PKA}^{n_{PKA}}}) - PKAa * (\gamma_{PKA} + \beta I_{p38} * \frac{p38a^{nI_{p38}}}{p38a^{nI_{p38}} + KmI_{p38}^{nI_{p38}}} + \beta I_{ERK} * \frac{ERKa^{nI_{ERK}}}{ERKa^{nI_{ERK}} + KmI_{ERK}^{nI_{ERK}}}) \tag{1}$$

$$\frac{dPKAi}{dt} = PKAa * (\gamma_{PKA} + \beta I_{p38} * \frac{p38a^{nI_{p38}}}{p38a^{nI_{p38}} + KmI_{p38}^{nI_{p38}}} + \beta I_{ERK} * \frac{ERKa^{nI_{ERK}}}{ERKa^{nI_{ERK}} + KmI_{ERK}^{nI_{ERK}}}) - PKAi * (\mu_{PKA} + \beta_{PKA} * \frac{cAMP^{n_{PKA}}}{cAMP^{n_{PKA}} + Km_{PKA}^{n_{PKA}}}) \tag{2}$$

$$\frac{dp38a}{dt} = p38i * (\mu_{p38} + \beta_{p38} * \frac{cAMP^{n_{p38}}}{cAMP^{n_{p38}} + Km_{p38}^{n_{p38}}}) - p38a * (\gamma_{p38}) \tag{3}$$

$$\frac{dp38i}{dt} = p38a * (\gamma_{p38} - p38i * (\mu_{p38} + \beta_{p38} * \frac{cAMP^{n_{p38}}}{cAMP^{n_{p38}} + Km_{p38}^{n_{p38}}}) \tag{4}$$

$$\frac{dERKa}{dt} = ERKi * (\mu_{ERK} + \beta_{ERK} * \frac{cAMP^{n_{ERK}}}{cAMP^{n_{ERK}} + Km_{ERK}^{n_{ERK}}}) - ERKa * (\gamma_{ERK} + \beta I_{PKA} * \frac{PKAa^{nI_{PKA}}}{PKAa^{nI_{PKA}} + KmI_{PKA}^{nI_{PKA}}}) \tag{5}$$

$$\frac{dERKi}{dt} = ERKa * (\gamma_{ERK} + \beta I_{PKA} * \frac{PKAa^{nI_{PKA}}}{PKAa^{nI_{PKA}} + KmI_{PKA}^{nI_{PKA}}}) - ERKi * (\mu_{EKR} + \beta_{ERK} * \frac{cAMP^{n_{ERK}}}{cAMP^{n_{ERK}} + Km_{ERK}^{n_{EKR}}}) \tag{6}$$

#### 4 Parameters for simulations shown in Figure 3

The parameters used to produce Figure 3 are summarized in table 3

Table 3: Parameters for model simulation. Related to Figure 3

| Parameter | Value |
|---|---|
| $\gamma_{PKA}$ | 1 |
| $\mathrm{Km}_{PKA}$ | $2 \cdot 10^{-2}$ |
| $\mathrm{n}_{PKA}$ | 0.87 |
| $\beta_{PKA}$ | 1.94 |
| $\mu_{PKA}$ | 1.23 |
| $\beta I_{PKA}$ | 1.73 |
| $\mathrm{nI}_{PKA}$ | 5.91 |
| $\mathrm{KmI}_{PKA}$ | 0.57 |
| $\gamma_{p38}$ | 3.4 |
| $\mathrm{Km}_{p38}$ | 0.54 |
| $\mathrm{n}_{p38}$ | 1.3 |
| $\beta_{p38}$ | 8.75 |
| $\mu_{p38}$ | 0.47 |
| $\beta I_{p38}$ | 8.18 |
| $\mathrm{nI}_{p38}$ | 1.48 |
| $\mathrm{KmI}_{p38}$ | 0.87 |
| $\gamma_{ERK}$ | 0.36 |
| $\mathrm{Km}_{ERK}$ | 0.65 |
| $\mathrm{n}_{ERK}$ | 1.14 |
| $\beta_{ERK}$ | 5.88 |
| $\mu_{ERK}$ | 0.79 |
| $\beta I_{ERK}$ | 9.02 |
| $\mathrm{nI}_{ERK}$ | 5.37 |
| $\mathrm{KmI}_{ERK}$ | 1.04 |
| initial PKAa | 0 |
| initial p38a | 0 |
| initial ERKa | 0 |
| initial PKAi | 1 |
| initial p38i | 1 |
| initial ERKi | 1 |
| Time | 50 |

| Name | LFC_NOvLOW | padj_NOvLOW | LFC_NOvHIGH | padj_NOvHIGH | LFC_HIGHvLOW | padj_HIGHvLOW | Cluster |
|---|---|---|---|---|---|---|---|
| ZADH2 | -1.9E+00 | 1.4E-05 | -2.9E+00 | 8.6E-10 | -1.0E+00 | 1.2E-01 | 8 |
| PIGN | -6.7E-01 | 8.7E-02 | -9.5E-01 | 2.5E-02 | -2.9E-01 | 8.6E-01 | 8 |
| POLI | -2.6E+00 | 1.5E-03 | -4.3E+00 | 9.1E-04 | -1.7E+00 | 4.9E-01 | 8 |
| HSD17B4 | -1.1E+00 | 3.4E-03 | -1.5E+00 | 3.6E-04 | -4.1E-01 | 7.0E-01 | 8 |
| ELAC1 | -1.3E+00 | 4.0E-02 | -1.5E+00 | 4.1E-02 | -2.3E-01 | 9.9E-01 | 8 |
| SHMT2 | -2.6E+00 | 3.3E-02 | -4.2E+00 | 7.8E-03 | -1.6E+00 | 6.5E-01 | 8 |
| RMND5B | -3.5E+00 | 1.4E-02 | -4.8E+00 | 1.4E-02 | -1.3E+00 | 9.2E-01 | 8 |
| IFNGR1 | -2.5E+00 | 4.5E-04 | -3.0E+00 | 2.0E-04 | -4.6E-01 | 9.6E-01 | 8 |
| NHSL1 | -1.2E+00 | 6.7E-03 | -1.4E+00 | 5.2E-03 | -1.8E-01 | 9.9E-01 | 8 |
| SAS | -1.4E+00 | 2.3E-04 | -1.3E+00 | 1.9E-03 | 4.1E-02 | 9.9E-01 | 8 |
| CDK4 | -2.8E+00 | 4.3E-05 | -2.4E+00 | 1.8E-03 | 3.6E-01 | 9.9E-01 | 8 |
| FUCA2 | -2.2E+00 | 4.7E-02 | -4.2E+00 | 6.2E-03 | -2.0E+00 | 4.6E-01 | 8 |
| HNRNPH1 | -7.9E-01 | 1.4E-02 | -1.2E+00 | 4.0E-04 | -4.1E-01 | 5.0E-01 | 8 |
| TBK1 | -1.6E+00 | 3.1E-04 | -2.1E+00 | 3.6E-05 | -4.3E-01 | 7.8E-01 | 8 |
| SASH1 | -3.0E-01 | 3.4E-01 | -7.2E-01 | 8.8E-03 | -4.2E-01 | 2.6E-01 | 8 |
| TAB2 | -2.1E+00 | 2.6E-04 | -2.4E+00 | 1.7E-04 | -3.3E-01 | 9.9E-01 | 8 |
| CAND1 | -8.2E-01 | 4.6E-02 | -1.0E+00 | 2.4E-02 | -2.3E-01 | 9.8E-01 | 8 |
| NUP107 | -1.4E+00 | 3.0E-04 | -1.4E+00 | 3.0E-03 | 6.6E-02 | 9.9E-01 | 8 |
| CNOT6 | -8.7E-01 | 1.6E-03 | -1.3E+00 | 1.4E-05 | -4.1E-01 | 4.2E-01 | 8 |
| FBXO5 | -2.3E+00 | 2.5E-02 | -3.0E+00 | 2.8E-02 | -6.3E-01 | 9.9E-01 | 8 |
| NA | -2.5E+00 | 1.4E-02 | -3.3E+00 | 2.0E-02 | -7.8E-01 | 9.8E-01 | 8 |
| TMEM242 | -2.5E+00 | 1.6E-05 | -2.2E+00 | 5.6E-04 | 2.4E-01 | 9.9E-01 | 8 |
| TNXB | -1.1E+00 | 4.2E-03 | -1.5E+00 | 1.3E-03 | -3.4E-01 | 8.7E-01 | 8 |
| MAP3K4 | -6.2E-01 | 1.3E-02 | -7.5E-01 | 8.0E-03 | -1.2E-01 | 9.9E-01 | 8 |
| EXT1 | -5.0E-01 | 2.5E-01 | -1.2E+00 | 4.7E-03 | -6.6E-01 | 2.4E-01 | 8 |
| TAP1 | -3.2E+00 | 9.0E-03 | -5.2E+00 | 4.6E-03 | -2.0E+00 | 6.4E-01 | 8 |
| DLA-DMA | -1.4E+00 | 4.2E-02 | -1.3E+00 | 1.2E-01 | 7.1E-02 | 9.9E-01 | 8 |
| ZUFSP | -8.8E-01 | 3.1E-03 | -1.4E+00 | 9.0E-06 | -5.6E-01 | 2.4E-01 | 8 |
| TSPO | -5.6E-01 | 9.8E-02 | -8.6E-01 | 1.3E-02 | -3.0E-01 | 7.1E-01 | 8 |
| ARFGAP3 | -1.7E+00 | 3.0E-05 | -1.7E+00 | 1.3E-04 | -1.4E-02 | 9.9E-01 | 8 |
| MTBP | -1.8E+00 | 2.3E-03 | -2.9E+00 | 2.2E-05 | -1.2E+00 | 2.7E-01 | 8 |
| TBC1D31 | -1.5E+00 | 1.7E-02 | -2.9E+00 | 8.6E-05 | -1.4E+00 | 1.8E-01 | 8 |
| PHF1 | -3.0E+00 | 5.5E-02 | -5.8E+00 | 4.8E-03 | -2.8E+00 | 4.4E-01 | 8 |
| UBE2B | -2.1E+00 | 5.7E-05 | -2.4E+00 | 4.4E-05 | -3.5E-01 | 9.7E-01 | 8 |
| ACO2 | -2.7E+00 | 4.5E-04 | -3.2E+00 | 3.0E-04 | -5.6E-01 | 9.5E-01 | 8 |
| MYC | -2.4E+00 | 9.4E-04 | -2.4E+00 | 3.8E-03 | -8.5E-03 | 1.0E+00 | 8 |
| MKL1 | -1.8E+00 | 1.5E-01 | -3.1E+00 | 3.7E-02 | -1.4E+00 | 6.9E-01 | 8 |
| BRD8 | -8.9E-01 | 3.9E-02 | -1.7E+00 | 4.9E-04 | -8.1E-01 | 2.6E-01 | 8 |
| HMGA1 | -1.3E+00 | 1.4E-04 | -1.2E+00 | 1.7E-03 | 9.6E-02 | 9.9E-01 | 8 |
| ERCC6L2 | -1.3E+00 | 1.7E-02 | -1.1E+00 | 9.4E-02 | 1.8E-01 | 9.9E-01 | 8 |
| ERCC6L2 | -1.3E+00 | 1.7E-02 | -1.1E+00 | 9.4E-02 | 1.8E-01 | 9.9E-01 | 8 |
| EGR1 | -3.8E+00 | 6.0E-06 | -4.2E+00 | 6.3E-06 | -3.9E-01 | 9.9E-01 | 8 |
| PDGFB | -1.8E+00 | 6.1E-03 | -2.4E+00 | 8.3E-04 | -6.0E-01 | 7.8E-01 | 8 |
| SUN2 | -3.5E+00 | 1.4E-02 | -5.3E+00 | 6.9E-04 | -1.8E+00 | 5.4E-01 | 8 |
| RMI1 | -1.4E+00 | 7.0E-03 | -1.5E+00 | 1.5E-02 | -1.0E-01 | 9.9E-01 | 8 |
| UBE2H | -5.8E-01 | 6.1E-02 | -1.3E+00 | 2.1E-05 | -7.6E-01 | 4.3E-02 | 8 |
| CYC1 | -8.1E-01 | 1.3E-03 | -1.1E+00 | 5.1E-05 | -2.7E-01 | 6.6E-01 | 8 |
| NA | -7.2E-01 | 8.0E-03 | -6.7E-01 | 3.8E-02 | 5.3E-02 | 9.9E-01 | 8 |
| TOMM6 | -2.6E+00 | 2.5E-02 | -2.3E+00 | 9.5E-02 | 2.6E-01 | 9.9E-01 | 8 |
| BYSL | -1.9E+00 | 1.3E-02 | -2.2E+00 | 2.2E-02 | -2.9E-01 | 9.9E-01 | 8 |
| TAF8 | -1.3E+00 | 6.9E-03 | -1.4E+00 | 1.2E-02 | -9.8E-02 | 9.9E-01 | 8 |
| C9orf72 | -1.6E+00 | 2.2E-02 | -1.4E+00 | 1.0E-01 | 1.8E-01 | 9.9E-01 | 8 |
| TBCC | -1.6E+00 | 2.7E-04 | -2.0E+00 | 5.5E-05 | -4.1E-01 | 8.2E-01 | 8 |
| POT1 | -6.7E-01 | 3.7E-02 | -8.5E-01 | 1.8E-02 | -1.8E-01 | 9.8E-01 | 8 |
| MEA1 | -1.7E+00 | 2.5E-03 | -1.6E+00 | 2.2E-02 | 1.8E-01 | 9.9E-01 | 8 |
| ZNF300 | -1.6E+00 | 2.8E-02 | -1.4E+00 | 1.3E-01 | 2.3E-01 | 9.9E-01 | 8 |
| NA | -1.9E+00 | 3.8E-02 | -3.4E+00 | 5.6E-03 | -1.6E+00 | 4.9E-01 | 8 |
| CKAP4 | -6.6E-01 | 6.9E-02 | -9.5E-01 | 1.2E-02 | -2.9E-01 | 8.0E-01 | 8 |
| VEGFA | -7.9E-01 | 2.0E-02 | -7.9E-01 | 4.5E-02 | 4.0E-03 | 1.0E+00 | 8 |
| KRIT1 | -7.1E-01 | 1.4E-01 | -1.1E+00 | 2.3E-02 | -4.3E-01 | 7.2E-01 | 8 |
| FRYL | -3.8E-01 | 8.2E-02 | -7.4E-01 | 7.1E-04 | -3.6E-01 | 2.3E-01 | 8 |
| FANCG | -1.5E+00 | 2.0E-02 | -1.5E+00 | 5.6E-02 | 2.9E-02 | 9.9E-01 | 8 |
| CHIC2 | -1.5E+00 | 9.7E-02 | -2.5E+00 | 1.5E-02 | -9.8E-01 | 6.6E-01 | 8 |
| COL1A2 | -2.8E-01 | 4.3E-01 | -6.1E-01 | 3.6E-02 | -3.3E-01 | 4.7E-01 | 8 |
| PPAPDC2 | -7.0E-01 | 8.2E-02 | -1.1E+00 | 9.8E-03 | -3.9E-01 | 6.8E-01 | 8 |
| NA | -2.6E+00 | 1.2E-02 | -5.9E+00 | 6.2E-04 | -3.3E+00 | 1.7E-01 | 8 |
| FHL2 | -2.6E+00 | 3.5E-05 | -2.4E+00 | 4.6E-04 | 1.4E-01 | 9.9E-01 | 8 |
| PON2 | -1.8E+00 | 1.5E-03 | -2.8E+00 | 8.4E-05 | -9.9E-01 | 4.6E-01 | 8 |
| CKS2 | -3.9E+00 | 8.5E-03 | -3.3E+00 | 5.4E-02 | 5.5E-01 | 9.9E-01 | 8 |
| CREB3 | -1.2E+00 | 1.8E-02 | -1.2E+00 | 5.1E-02 | 1.8E-02 | 9.9E-01 | 8 |
| SPIN1 | -2.4E+00 | 7.1E-06 | -2.2E+00 | 2.4E-04 | 2.4E-01 | 9.9E-01 | 8 |
| TMEM14A | -2.0E+00 | 2.5E-02 | -2.3E+00 | 3.5E-02 | -2.7E-01 | 9.9E-01 | 8 |
| PDCL3 | -1.4E+00 | 4.8E-03 | -2.1E+00 | 6.0E-05 | -7.9E-01 | 3.7E-01 | 8 |
| PAICS | -2.7E+00 | 1.0E-05 | -3.2E+00 | 2.2E-05 | -4.4E-01 | 9.8E-01 | 8 |
| REV1 | -8.7E-01 | 4.0E-02 | -1.3E+00 | 4.3E-03 | -4.7E-01 | 6.5E-01 | 8 |
| PHF14 | -1.0E+00 | 1.7E-03 | -1.2E+00 | 4.5E-04 | -2.2E-01 | 9.3E-01 | 8 |
| NA | -2.2E+00 | 2.8E-03 | -2.5E+00 | 2.9E-03 | -2.7E-01 | 9.9E-01 | 8 |
| CLTA | -9.2E-01 | 1.5E-03 | -1.0E+00 | 1.3E-03 | -1.1E-01 | 9.9E-01 | 8 |
| POLR2B | -1.0E+00 | 3.7E-05 | -1.4E+00 | 6.9E-08 | -4.0E-01 | 3.5E-01 | 8 |
| LIPT1 | -2.1E+00 | 3.1E-02 | -3.1E+00 | 1.5E-02 | -1.0E+00 | 8.2E-01 | 8 |
| IARS | -5.9E-01 | 2.3E-01 | -1.1E+00 | 2.2E-02 | -5.0E-01 | 5.6E-01 | 8 |
| INPP4A | -2.2E+00 | 1.2E-04 | -2.1E+00 | 1.1E-03 | 9.5E-02 | 9.9E-01 | 8 |
| UBE2M | -2.8E+00 | 7.3E-04 | -3.2E+00 | 7.1E-04 | -4.1E-01 | 9.9E-01 | 8 |
| ADH5 | -3.7E+00 | 1.3E-02 | -6.5E+00 | 1.3E-03 | -2.8E+00 | 4.7E-01 | 8 |
| TRIM28 | -7.0E-01 | 9.0E-02 | -1.2E+00 | 5.3E-03 | -5.2E-01 | 5.0E-01 | 8 |
| EXOSC3 | -1.3E+00 | 1.4E-01 | -2.1E+00 | 4.2E-02 | -7.4E-01 | 8.2E-01 | 8 |
| MACC1 | -1.4E+00 | 4.1E-04 | -1.6E+00 | 1.5E-04 | -2.3E-01 | 9.8E-01 | 8 |
| PIGF | -2.2E+00 | 4.2E-03 | -2.0E+00 | 2.3E-02 | 1.7E-01 | 9.9E-01 | 8 |
| RDH13 | -1.3E+00 | 1.0E-01 | -2.0E+00 | 3.3E-02 | -7.0E-01 | 8.2E-01 | 8 |
| TSEN34 | -2.0E+00 | 1.3E-02 | -2.3E+00 | 1.8E-02 | -2.7E-01 | 9.9E-01 | 8 |
| TMC4 | -1.5E+00 | 1.8E-02 | -2.0E+00 | 6.8E-03 | -4.9E-01 | 8.9E-01 | 8 |
| ASB3 | -1.2E+00 | 1.0E-01 | -1.7E+00 | 4.2E-02 | -5.1E-01 | 9.0E-01 | 8 |
| ASB3 | -1.2E+00 | 1.0E-01 | -1.7E+00 | 4.2E-02 | -5.1E-01 | 9.0E-01 | 8 |

| | | | | | | | |
|---|---|---|---|---|---|---|---|
| RTN4 | -1.8E+00 | 3.5E-05 | -2.8E+00 | 1.1E-07 | -9.4E-01 | 2.5E-01 | 8 |
| PPP2R1A | -1.2E+00 | 2.1E-02 | -1.3E+00 | 1.8E-02 | -1.7E-01 | 9.9E-01 | 8 |
| TTK | -1.6E+00 | 4.9E-02 | -1.8E+00 | 6.2E-02 | -2.2E-01 | 9.9E-01 | 8 |
| NA | -2.4E+00 | 5.4E-02 | -3.4E+00 | 2.3E-02 | -1.0E+00 | 8.9E-01 | 8 |
| PNPT1 | -7.8E-01 | 3.3E-01 | -1.9E+00 | 1.2E-02 | -1.1E+00 | 3.1E-01 | 8 |
| DOPEY1 | -1.0E+00 | 1.0E-01 | -1.9E+00 | 1.1E-02 | -8.3E-01 | 5.5E-01 | 8 |
| RUFY3 | -5.1E-01 | 1.0E-01 | -1.0E+00 | 8.0E-04 | -5.0E-01 | 2.2E-01 | 8 |
| ANKRD17 | -8.0E-01 | 9.4E-05 | -7.4E-01 | 1.3E-03 | 6.0E-02 | 9.9E-01 | 8 |
| UGCG | -6.0E-01 | 1.0E-02 | -1.3E+00 | 8.7E-08 | -6.5E-01 | 1.6E-02 | 8 |
| MFSD2A | -2.7E+00 | 3.1E-03 | -4.7E+00 | 1.8E-03 | -2.1E+00 | 4.7E-01 | 8 |
| SYNCRIP | -5.8E-01 | 3.4E-02 | -1.2E+00 | 1.5E-05 | -6.1E-01 | 7.3E-02 | 8 |
| PPIE | -9.0E-01 | 1.6E-01 | -1.4E+00 | 3.2E-02 | -5.5E-01 | 7.5E-01 | 8 |
| NA | -6.4E-01 | 1.1E-01 | -9.9E-01 | 1.5E-02 | -3.5E-01 | 7.0E-01 | 8 |
| NA | -6.4E-01 | 1.1E-01 | -9.9E-01 | 1.5E-02 | -3.5E-01 | 7.0E-01 | 8 |
| NA | -6.4E-01 | 1.1E-01 | -9.9E-01 | 1.5E-02 | -3.5E-01 | 7.0E-01 | 8 |
| NA | -6.4E-01 | 1.1E-01 | -9.9E-01 | 1.5E-02 | -3.5E-01 | 7.0E-01 | 8 |
| NUP205 | -1.6E+00 | 9.2E-03 | -2.0E+00 | 3.4E-03 | -3.8E-01 | 9.5E-01 | 8 |
| PNO1 | -2.4E+00 | 1.1E-02 | -3.6E+00 | 2.2E-03 | -1.2E+00 | 6.8E-01 | 8 |
| IFRD1 | -2.9E+00 | 9.4E-03 | -3.1E+00 | 1.2E-02 | -2.4E-01 | 9.9E-01 | 8 |
| NR1H2 | -2.9E+00 | 1.4E-02 | -3.6E+00 | 2.0E-02 | -6.8E-01 | 9.9E-01 | 8 |
| GRHL1 | -1.3E+00 | 5.1E-04 | -1.4E+00 | 1.0E-03 | -7.5E-02 | 9.9E-01 | 8 |
| CAV1 | -1.2E+00 | 2.7E-02 | -1.5E+00 | 1.3E-02 | -2.8E-01 | 9.9E-01 | 8 |
| FBXL4 | -2.0E+00 | 3.3E-04 | -1.9E+00 | 4.2E-03 | 1.8E-01 | 9.9E-01 | 8 |
| ING3 | -5.1E-01 | 1.9E-01 | -8.6E-01 | 2.6E-02 | -3.5E-01 | 6.7E-01 | 8 |
| PTOV1 | -4.5E-01 | 2.3E-01 | -8.4E-01 | 1.8E-02 | -4.0E-01 | 5.1E-01 | 8 |
| HACE1 | -1.1E+00 | 2.4E-02 | -1.2E+00 | 3.1E-02 | -1.0E-01 | 9.9E-01 | 8 |
| ZNF330 | -1.1E+00 | 2.2E-03 | -1.1E+00 | 7.2E-03 | 2.3E-03 | 1.0E+00 | 8 |
| DDX1 | -7.6E-01 | 2.7E-03 | -9.5E-01 | 5.3E-04 | -1.9E-01 | 8.8E-01 | 8 |
| FAF1 | -1.0E+00 | 9.1E-02 | -2.0E+00 | 4.8E-03 | -1.0E+00 | 4.0E-01 | 8 |
| ADCK2 | -4.2E+00 | 2.7E-02 | -3.7E+00 | 1.2E-01 | 5.0E-01 | 9.9E-01 | 8 |
| NA | -5.4E+00 | 1.0E-03 | -5.8E+00 | 2.2E-03 | -4.3E-01 | 9.9E-01 | 8 |
| NA | -1.4E+00 | 2.7E-03 | -1.4E+00 | 6.7E-03 | -3.7E-02 | 9.9E-01 | 8 |
| TUBE1 | -9.9E-01 | 1.3E-02 | -1.1E+00 | 2.2E-02 | -7.3E-02 | 9.9E-01 | 8 |
| TPRA1 | -2.9E+00 | 6.1E-03 | -3.4E+00 | 1.3E-02 | -5.4E-01 | 9.9E-01 | 8 |
| CCNA2 | -2.5E+00 | 5.2E-02 | -3.2E+00 | 4.4E-02 | -7.1E-01 | 9.9E-01 | 8 |
| GATA2 | -1.9E+00 | 1.7E-03 | -2.1E+00 | 2.2E-03 | -1.6E-01 | 9.9E-01 | 8 |
| SLC1A5 | -3.0E+00 | 2.5E-02 | -5.1E+00 | 4.9E-03 | -2.1E+00 | 6.3E-01 | 8 |
| COPG1 | -2.5E+00 | 5.4E-04 | -3.3E+00 | 4.5E-05 | -7.6E-01 | 7.2E-01 | 8 |
| LRRC41 | -2.4E+00 | 1.9E-06 | -2.7E+00 | 2.9E-06 | -2.7E-01 | 9.9E-01 | 8 |
| HADHA | -6.6E-01 | 1.0E-01 | -9.2E-01 | 3.3E-02 | -2.6E-01 | 9.0E-01 | 8 |
| TMEM60 | -2.1E+00 | 8.1E-05 | -2.5E+00 | 4.9E-05 | -3.6E-01 | 9.7E-01 | 8 |
| ACBD5 | -1.1E+00 | 9.8E-03 | -1.0E+00 | 4.1E-02 | 7.3E-02 | 9.9E-01 | 8 |
| PRSS23 | -2.8E+00 | 8.0E-03 | -2.6E+00 | 4.3E-02 | 2.3E-01 | 9.9E-01 | 8 |
| ZNF282 | -1.5E+00 | 1.7E-02 | -2.0E+00 | 7.1E-03 | -4.7E-01 | 9.2E-01 | 8 |
| NDUFB6 | -2.8E+00 | 9.1E-03 | -3.3E+00 | 4.3E-03 | -5.6E-01 | 9.9E-01 | 8 |
| EED | -1.5E+00 | 1.9E-05 | -1.8E+00 | 1.5E-06 | -3.2E-01 | 8.0E-01 | 8 |
| SLC6A6 | -1.3E+00 | 1.8E-03 | -1.8E+00 | 1.4E-05 | -5.8E-01 | 4.2E-01 | 8 |
| ERCC3 | -1.7E+00 | 6.3E-03 | -3.1E+00 | 4.7E-05 | -1.4E+00 | 2.3E-01 | 8 |
| PRDX1 | -1.8E+00 | 6.7E-05 | -2.0E+00 | 2.5E-05 | -2.5E-01 | 9.8E-01 | 8 |
| NA | -2.6E+00 | 1.4E-03 | -2.6E+00 | 5.0E-03 | 2.7E-02 | 9.9E-01 | 8 |
| NA | -2.6E+00 | 1.4E-03 | -2.6E+00 | 5.0E-03 | 2.7E-02 | 9.9E-01 | 8 |
| NA | -2.6E+00 | 1.4E-03 | -2.6E+00 | 5.0E-03 | 2.7E-02 | 9.9E-01 | 8 |
| UBP1 | -5.9E-01 | 2.8E-03 | -7.1E-01 | 9.4E-04 | -1.2E-01 | 9.5E-01 | 8 |
| CBLC | -1.4E+00 | 5.8E-02 | -1.8E+00 | 4.2E-02 | -3.9E-01 | 9.9E-01 | 8 |
| HSPA14 | -1.2E+00 | 1.2E-03 | -1.2E+00 | 6.2E-03 | 3.7E-02 | 9.9E-01 | 8 |
| PLK3 | -1.8E+00 | 4.2E-02 | -2.8E+00 | 7.3E-03 | -9.7E-01 | 7.0E-01 | 8 |
| SUMO2 | -1.8E+00 | 1.5E-05 | -2.5E+00 | 2.1E-07 | -6.9E-01 | 4.4E-01 | 8 |
| MRPS7 | -1.5E+00 | 2.9E-03 | -2.3E+00 | 8.5E-05 | -8.1E-01 | 4.5E-01 | 8 |
| TSEN54 | -2.2E+00 | 6.2E-02 | -3.6E+00 | 3.3E-02 | -1.4E+00 | 7.7E-01 | 8 |
| ITGA9 | -1.5E+00 | 5.2E-03 | -1.8E+00 | 5.0E-03 | -2.9E-01 | 9.9E-01 | 8 |
| cfa-mir-671 | -8.4E-01 | 1.7E-02 | -1.2E+00 | 1.5E-03 | -3.7E-01 | 6.8E-01 | 8 |
| ATP6V0B | -9.1E-01 | 6.7E-03 | -1.1E+00 | 4.2E-03 | -1.5E-01 | 9.9E-01 | 8 |
| SNX17 | -2.2E+00 | 1.7E-02 | -3.4E+00 | 3.2E-03 | -1.2E+00 | 6.8E-01 | 8 |
| OXSR1 | -7.8E-01 | 2.9E-02 | -1.0E+00 | 8.0E-03 | -2.5E-01 | 8.9E-01 | 8 |
| PPARG | -2.4E+00 | 1.6E-02 | -2.8E+00 | 9.8E-03 | -4.4E-01 | 9.9E-01 | 8 |
| SEC13 | -3.2E+00 | 3.8E-05 | -3.3E+00 | 1.6E-04 | -1.1E-01 | 9.9E-01 | 8 |
| SFMBT2 | -1.1E+00 | 7.5E-03 | -2.5E+00 | 1.0E-08 | -1.3E+00 | 6.9E-03 | 8 |
| VHL | -1.7E+00 | 5.3E-02 | -2.0E+00 | 4.7E-02 | -3.3E-01 | 9.9E-01 | 8 |
| GZF1 | -8.7E-01 | 9.7E-03 | -7.9E-01 | 5.0E-02 | 8.0E-02 | 9.9E-01 | 8 |
| NA | -3.8E+00 | 9.9E-04 | -4.9E+00 | 1.2E-03 | -1.0E+00 | 9.2E-01 | 8 |
| NA | -3.8E+00 | 9.9E-04 | -4.9E+00 | 1.2E-03 | -1.0E+00 | 9.2E-01 | 8 |
| BRK1 | -2.2E+00 | 2.4E-02 | -2.6E+00 | 4.3E-02 | -3.2E-01 | 9.9E-01 | 8 |
| EMC3 | -1.4E+00 | 5.1E-03 | -1.5E+00 | 1.1E-02 | -8.5E-02 | 9.9E-01 | 8 |
| RBM17 | -2.1E+00 | 2.5E-05 | -2.2E+00 | 4.0E-05 | -1.2E-01 | 9.9E-01 | 8 |
| NCAPG2 | -1.3E+00 | 2.1E-02 | -1.4E+00 | 3.2E-02 | -8.9E-02 | 9.9E-01 | 8 |
| NA | -1.8E+00 | 8.3E-02 | -2.6E+00 | 2.9E-02 | -8.8E-01 | 8.4E-01 | 8 |
| SEC22C | -3.6E+00 | 1.6E-03 | -3.0E+00 | 2.0E-02 | 5.4E-01 | 9.9E-01 | 8 |
| SERTAD1 | -3.6E+00 | 2.9E-05 | -6.0E+00 | 1.4E-04 | -2.4E+00 | 4.1E-01 | 8 |
| KLF6 | -4.1E+00 | 3.1E-05 | -3.7E+00 | 8.8E-04 | 4.1E-01 | 9.9E-01 | 8 |
| HGSNAT | -9.5E-01 | 3.7E-02 | -9.2E-01 | 9.6E-02 | 3.5E-02 | 9.9E-01 | 8 |
| HGSNAT | -9.5E-01 | 3.7E-02 | -9.2E-01 | 9.6E-02 | 3.5E-02 | 9.9E-01 | 8 |
| TGFBR2 | -1.6E+00 | 3.0E-05 | -2.0E+00 | 5.7E-07 | -4.4E-01 | 6.3E-01 | 8 |
| SUPT5H | -4.9E-01 | 9.6E-03 | -7.4E-01 | 1.9E-04 | -2.5E-01 | 4.7E-01 | 8 |
| ZFP36 | -1.4E+00 | 2.9E-03 | -2.3E+00 | 6.8E-06 | -9.0E-01 | 2.3E-01 | 8 |
| DPY30 | -2.0E+00 | 2.0E-04 | -2.5E+00 | 1.3E-04 | -4.9E-01 | 8.8E-01 | 8 |
| CXXC5 | -1.8E+00 | 1.4E-02 | -2.4E+00 | 4.3E-03 | -5.4E-01 | 9.0E-01 | 8 |
| PPP1R12A | -1.1E+00 | 6.1E-06 | -1.4E+00 | 4.9E-08 | -3.0E-01 | 5.6E-01 | 8 |
| SRA1 | -1.0E+00 | 1.6E-02 | -1.0E+00 | 4.0E-02 | -3.9E-04 | 1.0E+00 | 8 |
| ASPSCR1 | -9.6E-01 | 1.6E-03 | -1.2E+00 | 2.7E-04 | -2.6E-01 | 8.3E-01 | 8 |
| PCCA | -2.2E+00 | 6.8E-02 | -4.2E+00 | 2.3E-02 | -2.0E+00 | 6.2E-01 | 8 |
| ADAM9 | -1.6E+00 | 2.7E-03 | -2.1E+00 | 1.8E-04 | -5.3E-01 | 7.1E-01 | 8 |
| HARS | -1.9E+00 | 5.3E-02 | -5.5E+00 | 1.0E-03 | -3.6E+00 | 9.2E-02 | 8 |
| HARS | -1.9E+00 | 5.3E-02 | -5.5E+00 | 1.0E-03 | -3.6E+00 | 9.2E-02 | 8 |
| BTD | -5.9E-01 | 4.0E-02 | -1.2E+00 | 3.7E-05 | -6.1E-01 | 1.0E-01 | 8 |

| | | | | | | | |
|---|---|---|---|---|---|---|---|
| DUS1L | -1.1E+00 | 5.4E-04 | -1.2E+00 | 4.6E-04 | -1.3E-01 | 9.9E-01 | 8 |
| HEATR5B | -8.4E-01 | 1.2E-02 | -9.0E-01 | 2.0E-02 | -5.7E-02 | 9.9E-01 | 8 |
| PROSER1 | -2.0E+00 | 5.6E-03 | -1.8E+00 | 2.6E-02 | 1.4E-01 | 9.9E-01 | 8 |
| NA | -2.4E+00 | 5.1E-04 | -3.4E+00 | 5.3E-06 | -1.1E+00 | 4.3E-01 | 8 |
| ASTE1 | -1.0E+00 | 7.4E-03 | -1.2E+00 | 5.5E-03 | -1.9E-01 | 9.9E-01 | 8 |
| LSM1 | -2.3E+00 | 2.7E-03 | -2.2E+00 | 1.6E-02 | 1.2E-01 | 9.9E-01 | 8 |
| ARHGEF7 | -4.2E-01 | 3.5E-01 | -8.7E-01 | 3.4E-02 | -4.5E-01 | 5.0E-01 | 8 |
| ATL2 | -1.8E+00 | 1.5E-03 | -1.7E+00 | 8.5E-03 | 1.0E-01 | 9.9E-01 | 8 |
| STARD13 | -1.1E+00 | 1.2E-05 | -1.7E+00 | 5.6E-10 | -5.4E-01 | 1.5E-01 | 8 |
| TMCO3 | -1.5E+00 | 1.4E-02 | -1.5E+00 | 4.4E-02 | 6.6E-02 | 9.9E-01 | 8 |
| ULK1 | -6.1E-01 | 7.0E-02 | -9.9E-01 | 3.8E-03 | -3.9E-01 | 5.3E-01 | 8 |
| SIRT3 | -2.3E+00 | 8.9E-03 | -2.4E+00 | 2.0E-02 | -1.3E-01 | 9.9E-01 | 8 |
| RBM27 | -3.7E-01 | 1.3E-01 | -6.3E-01 | 1.1E-02 | -2.6E-01 | 5.7E-01 | 8 |
| FRMD4B | -1.1E+00 | 1.7E-05 | -2.7E+00 | 8.3E-18 | -1.6E+00 | 4.9E-06 | 8 |
| NA | -9.5E-01 | 2.1E-02 | -9.7E-01 | 3.8E-02 | -2.8E-02 | 9.9E-01 | 8 |
| APAF1 | -1.1E+00 | 2.3E-02 | -1.5E+00 | 3.8E-03 | -4.2E-01 | 7.8E-01 | 8 |
| SFSWAP | -3.4E-01 | 1.8E-01 | -6.0E-01 | 1.6E-02 | -2.7E-01 | 5.6E-01 | 8 |
| SNAI2 | -2.4E+00 | 1.1E-16 | -2.3E+00 | 3.4E-12 | 1.6E-01 | 9.9E-01 | 8 |
| RYK | -1.5E+00 | 8.6E-03 | -2.8E+00 | 5.5E-05 | -1.2E+00 | 2.2E-01 | 8 |
| ACTR6 | -6.8E-01 | 2.3E-02 | -8.1E-01 | 1.7E-02 | -1.3E-01 | 9.9E-01 | 8 |
| NA | -3.3E+00 | 2.2E-02 | -4.8E+00 | 3.5E-03 | -1.5E+00 | 7.2E-01 | 8 |
| PARG | -5.3E-01 | 5.8E-02 | -1.0E+00 | 4.8E-04 | -5.0E-01 | 2.3E-01 | 8 |
| FZD10 | -1.9E+00 | 1.6E-02 | -2.3E+00 | 1.7E-02 | -4.0E-01 | 9.9E-01 | 8 |
| TRAF6 | -4.4E-01 | 1.6E-01 | -6.7E-01 | 4.2E-02 | -2.3E-01 | 8.2E-01 | 8 |
| CDK8 | -1.6E+00 | 3.5E-03 | -1.9E+00 | 2.1E-03 | -2.6E-01 | 9.9E-01 | 8 |
| RNF6 | -8.6E-01 | 8.0E-02 | -1.3E+00 | 1.8E-02 | -3.9E-01 | 8.2E-01 | 8 |
| BRI3BP | -1.5E+00 | 1.1E-02 | -2.0E+00 | 7.7E-04 | -5.8E-01 | 6.9E-01 | 8 |
| TMEM9B | -2.2E+00 | 5.3E-02 | -2.7E+00 | 4.3E-02 | -5.4E-01 | 9.9E-01 | 8 |
| MRPL15 | -2.5E+00 | 2.6E-02 | -5.6E+00 | 1.7E-03 | -3.1E+00 | 2.4E-01 | 8 |
| ARL1 | -3.0E+00 | 5.6E-04 | -2.8E+00 | 5.8E-03 | 2.3E-01 | 9.9E-01 | 8 |
| MFSD5 | -1.5E+00 | 4.3E-04 | -1.7E+00 | 3.0E-04 | -2.0E-01 | 9.9E-01 | 8 |
| NA | -2.2E+00 | 2.6E-02 | -4.9E+00 | 8.8E-07 | -2.7E+00 | 2.0E-02 | 8 |
| UBXN2B | -5.3E-01 | 3.2E-02 | -6.4E-01 | 2.1E-02 | -1.0E-01 | 9.9E-01 | 8 |
| MICU2 | -1.7E+00 | 1.7E-03 | -1.7E+00 | 5.4E-03 | 1.1E-03 | 1.0E+00 | 8 |
| LATS2 | -1.1E+00 | 2.1E-03 | -1.4E+00 | 2.6E-04 | -3.0E-01 | 8.2E-01 | 8 |
| IPO11 | -9.6E-01 | 1.6E-02 | -9.8E-01 | 3.9E-02 | -1.7E-02 | 9.9E-01 | 8 |
| ANKRD1 | -3.1E+00 | 2.3E-02 | -3.2E+00 | 3.9E-02 | -1.0E-01 | 9.9E-01 | 8 |
| NUP37 | -1.0E+00 | 3.5E-02 | -1.1E+00 | 6.6E-02 | -1.7E-02 | 9.9E-01 | 8 |
| TNKS2 | -1.1E+00 | 7.2E-03 | -1.7E+00 | 4.3E-05 | -6.2E-01 | 3.3E-01 | 8 |
| DEPDC7 | -1.9E+00 | 4.6E-02 | -2.6E+00 | 8.4E-03 | -7.3E-01 | 8.2E-01 | 8 |
| REEP5 | -2.6E+00 | 2.3E-02 | -2.4E+00 | 7.9E-02 | 2.0E-01 | 9.9E-01 | 8 |
| NA | -6.2E-01 | 1.4E-01 | -1.3E+00 | 3.4E-03 | -6.9E-01 | 2.9E-01 | 8 |
| LSM14A | -8.5E-01 | 2.5E-03 | -1.0E+00 | 7.8E-04 | -1.8E-01 | 9.4E-01 | 8 |
| STAG1 | -7.8E-01 | 2.1E-03 | -8.0E-01 | 5.8E-03 | -1.8E-02 | 9.9E-01 | 8 |
| WDR36 | -1.6E+00 | 2.4E-05 | -1.4E+00 | 5.9E-04 | 1.4E-01 | 9.9E-01 | 8 |
| PPWD1 | -2.2E+00 | 1.3E-05 | -3.2E+00 | 1.3E-07 | -9.5E-01 | 3.6E-01 | 8 |
| SLC35G2 | -2.5E+00 | 3.9E-03 | -2.9E+00 | 3.8E-03 | -4.3E-01 | 9.9E-01 | 8 |
| TMED2 | -2.0E+00 | 6.4E-05 | -2.0E+00 | 2.7E-04 | -2.5E-03 | 1.0E+00 | 8 |
| GIN1 | -1.4E+00 | 2.6E-02 | -1.3E+00 | 9.0E-02 | 1.2E-01 | 9.9E-01 | 8 |
| ANKRD37 | -8.3E-01 | 1.6E-02 | -1.5E+00 | 5.8E-05 | -6.3E-01 | 2.3E-01 | 8 |
| PTCD3 | -1.4E+00 | 1.0E-02 | -2.2E+00 | 1.1E-03 | -7.6E-01 | 6.1E-01 | 8 |
| SLC25A4 | -1.5E+00 | 5.2E-04 | -1.4E+00 | 2.4E-03 | 3.0E-02 | 9.9E-01 | 8 |
| GSS | -1.9E+00 | 5.3E-03 | -2.1E+00 | 1.0E-02 | -1.8E-01 | 9.9E-01 | 8 |
| C4ORF27 | -1.4E+00 | 2.4E-03 | -1.3E+00 | 1.8E-02 | 1.2E-01 | 9.9E-01 | 8 |
| SLMAP | -8.1E-01 | 5.7E-03 | -8.0E-01 | 1.8E-02 | 1.3E-02 | 9.9E-01 | 8 |
| LSM6 | -3.6E+00 | 2.7E-04 | -4.5E+00 | 5.3E-05 | -8.4E-01 | 8.5E-01 | 8 |
| MYOF | -3.3E-01 | 3.3E-01 | -7.1E-01 | 1.7E-02 | -3.8E-01 | 3.9E-01 | 8 |
| MAT2A | -1.1E+00 | 3.7E-02 | -1.8E+00 | 5.4E-03 | -6.5E-01 | 6.6E-01 | 8 |
| GALNT6 | -5.5E-01 | 2.7E-01 | -1.0E+00 | 3.6E-02 | -4.7E-01 | 6.2E-01 | 8 |
| ZDHHC5 | -1.8E+00 | 6.9E-06 | -1.9E+00 | 8.1E-06 | -1.4E-01 | 9.9E-01 | 8 |
| PRMT10 | -1.7E+00 | 2.6E-03 | -1.9E+00 | 2.9E-03 | -2.3E-01 | 9.9E-01 | 8 |
| CLP1 | -2.5E+00 | 7.0E-05 | -3.1E+00 | 7.3E-06 | -5.6E-01 | 8.2E-01 | 8 |
| TIMM10 | -1.3E+00 | 1.3E-03 | -1.2E+00 | 9.3E-03 | 8.2E-02 | 9.9E-01 | 8 |
| HMGB2 | -1.2E+00 | 3.9E-02 | -2.1E+00 | 5.5E-04 | -8.8E-01 | 3.4E-01 | 8 |
| SAP30 | -2.0E+00 | 1.7E-04 | -2.2E+00 | 2.4E-04 | -1.8E-01 | 9.9E-01 | 8 |
| ZCCHC8 | -6.2E-01 | 1.5E-01 | -1.2E+00 | 5.3E-03 | -5.7E-01 | 3.8E-01 | 8 |
| ATR | -8.3E-01 | 2.2E-01 | -1.7E+00 | 1.9E-02 | -8.9E-01 | 4.6E-01 | 8 |
| DIABLO | -2.3E+00 | 2.7E-03 | -2.1E+00 | 2.3E-02 | 2.2E-01 | 9.9E-01 | 8 |
| RPS3A | -2.6E+00 | 3.4E-03 | -2.3E+00 | 2.5E-02 | 2.8E-01 | 9.9E-01 | 8 |
| RPS3A | -2.6E+00 | 3.4E-03 | -2.3E+00 | 2.5E-02 | 2.8E-01 | 9.9E-01 | 8 |
| RDH10 | -1.1E+00 | 1.6E-01 | -1.9E+00 | 1.7E-02 | -8.1E-01 | 5.9E-01 | 8 |
| NA | -2.5E+00 | 4.0E-03 | -2.3E+00 | 2.3E-02 | 1.4E-01 | 9.9E-01 | 8 |
| TCEB1 | -2.3E+00 | 9.9E-03 | -2.0E+00 | 6.2E-02 | 3.1E-01 | 9.9E-01 | 8 |
| RBM12 | -6.8E-01 | 3.4E-02 | -8.8E-01 | 1.2E-02 | -2.0E-01 | 9.4E-01 | 8 |
| INTS9 | -1.4E+00 | 2.5E-02 | -1.9E+00 | 4.8E-03 | -5.0E-01 | 8.4E-01 | 8 |
| HK2 | -1.4E+00 | 4.4E-03 | -1.2E+00 | 3.4E-02 | 1.7E-01 | 9.9E-01 | 8 |
| RACGAP1 | -1.3E+00 | 1.3E-03 | -1.4E+00 | 3.6E-03 | -3.6E-02 | 9.9E-01 | 8 |
| HTRA2 | -1.8E+00 | 2.0E-02 | -2.0E+00 | 2.8E-02 | -1.4E-01 | 9.9E-01 | 8 |
| DPH6 | -6.5E-01 | 1.6E-01 | -1.2E+00 | 1.3E-02 | -5.4E-01 | 5.2E-01 | 8 |
| CNOT1 | -1.4E+00 | 1.5E-05 | -2.1E+00 | 5.5E-09 | -7.5E-01 | 1.6E-01 | 8 |
| MCRS1 | -2.9E+00 | 1.9E-03 | -3.0E+00 | 6.7E-03 | -7.5E-02 | 9.9E-01 | 8 |
| SCARB2 | -1.5E+00 | 2.3E-02 | -1.3E+00 | 1.0E-01 | 1.6E-01 | 9.9E-01 | 8 |
| SHROOM3 | -5.3E-01 | 1.8E-01 | -1.1E+00 | 4.4E-03 | -5.4E-01 | 3.2E-01 | 8 |
| SHROOM3 | -5.3E-01 | 1.8E-01 | -1.1E+00 | 4.4E-03 | -5.4E-01 | 3.2E-01 | 8 |
| PPID | -8.4E-01 | 1.3E-02 | -8.3E-01 | 4.1E-02 | 1.6E-02 | 9.9E-01 | 8 |
| FNIP2 | -7.3E-01 | 8.3E-03 | -9.6E-01 | 1.4E-03 | -2.3E-01 | 8.3E-01 | 8 |
| BACH1 | -4.9E-01 | 7.1E-02 | -6.3E-01 | 3.3E-02 | -1.4E-01 | 9.8E-01 | 8 |
| BACH1 | -4.9E-01 | 7.1E-02 | -6.3E-01 | 3.3E-02 | -1.4E-01 | 9.8E-01 | 8 |
| EIF2AK4 | -9.1E-01 | 1.0E-04 | -1.1E+00 | 3.2E-05 | -1.7E-01 | 9.2E-01 | 8 |
| WWP1 | -2.7E-01 | 4.7E-01 | -6.2E-01 | 4.9E-02 | -3.5E-01 | 4.8E-01 | 8 |
| NAA25 | -4.8E-01 | 3.1E-01 | -1.1E+00 | 1.3E-02 | -6.1E-01 | 3.4E-01 | 8 |
| FKBP11 | -1.9E+00 | 1.1E-02 | -1.9E+00 | 2.4E-02 | -1.9E-02 | 9.9E-01 | 8 |
| DUSP11 | -3.1E+00 | 9.5E-03 | -3.2E+00 | 1.8E-02 | -1.2E-01 | 9.9E-01 | 8 |
| SERINC5 | -1.3E+00 | 6.5E-03 | -1.2E+00 | 3.1E-02 | 1.1E-01 | 9.9E-01 | 8 |

| | | | | | | | |
|---|---|---|---|---|---|---|---|
| CCNT1 | -1.2E+00 | 4.3E-03 | -1.4E+00 | 2.6E-03 | -2.1E-01 | 9.9E-01 | 8 |
| SPCS1 | -2.4E+00 | 1.7E-04 | -3.8E+00 | 2.0E-07 | -1.3E+00 | 2.2E-01 | 8 |
| GLT8D1 | -1.5E+00 | 1.3E-02 | -2.2E+00 | 3.0E-03 | -6.6E-01 | 7.6E-01 | 8 |
| KCTD9 | -1.7E+00 | 1.0E-05 | -1.8E+00 | 2.8E-05 | -7.6E-02 | 9.9E-01 | 8 |
| ANXA10 | -2.9E+00 | 1.6E-02 | -2.4E+00 | 9.0E-02 | 4.1E-01 | 9.9E-01 | 8 |
| TMEM64 | -1.9E+00 | 4.8E-02 | -1.6E+00 | 1.9E-01 | 2.9E-01 | 9.9E-01 | 8 |
| NUP93 | -8.6E-01 | 3.6E-02 | -7.6E-01 | 1.3E-01 | 9.7E-02 | 9.9E-01 | 8 |
| MINA | -1.3E+00 | 1.0E-02 | -1.5E+00 | 6.7E-03 | -2.7E-01 | 9.9E-01 | 8 |
| GTF2H1 | -2.4E+00 | 4.1E-06 | -2.8E+00 | 6.9E-07 | -3.9E-01 | 9.0E-01 | 8 |
| CLDND1 | -1.7E+00 | 4.1E-02 | -1.5E+00 | 1.5E-01 | 2.1E-01 | 9.9E-01 | 8 |
| RPAP3 | -1.3E+00 | 7.1E-04 | -1.2E+00 | 7.2E-03 | 1.2E-01 | 9.9E-01 | 8 |
| WRNIP1 | -7.5E-01 | 9.2E-03 | -7.0E-01 | 3.9E-02 | 5.3E-02 | 9.9E-01 | 8 |
| TSG101 | -2.7E+00 | 1.2E-02 | -4.0E+00 | 5.2E-04 | -1.3E+00 | 5.7E-01 | 8 |
| TOMM70A | -9.1E-01 | 5.4E-02 | -1.7E+00 | 8.9E-04 | -7.9E-01 | 3.1E-01 | 8 |
| MYBL2 | -1.4E+00 | 1.0E-01 | -3.0E+00 | 7.2E-03 | -1.6E+00 | 3.9E-01 | 8 |
| SLC40A1 | -1.1E+00 | 3.7E-02 | -1.8E+00 | 2.5E-03 | -6.7E-01 | 5.6E-01 | 8 |
| POLK | -1.8E+00 | 3.0E-06 | -1.8E+00 | 2.1E-05 | 2.5E-04 | 1.0E+00 | 8 |
| HIBCH | -2.0E+00 | 5.9E-03 | -2.3E+00 | 7.1E-03 | -2.9E-01 | 9.9E-01 | 8 |
| COL4A3BP | -9.8E-01 | 1.2E-02 | -1.1E+00 | 1.2E-02 | -1.2E-01 | 9.9E-01 | 8 |
| NDUFAF1 | -1.5E+00 | 5.7E-04 | -1.9E+00 | 4.7E-05 | -4.5E-01 | 7.3E-01 | 8 |
| ZC3H11A | -1.1E+00 | 2.0E-03 | -1.7E+00 | 1.6E-05 | -5.4E-01 | 4.0E-01 | 8 |
| ZC3H11A | -1.1E+00 | 2.0E-03 | -1.7E+00 | 1.6E-05 | -5.4E-01 | 4.0E-01 | 8 |
| SETD4 | -1.3E+00 | 6.0E-03 | -1.3E+00 | 2.0E-02 | -1.7E-02 | 9.9E-01 | 8 |
| CSDE1 | -6.0E-01 | 7.2E-02 | -7.7E-01 | 3.8E-02 | -1.7E-01 | 9.9E-01 | 8 |
| ZNF248 | -1.6E+00 | 8.4E-06 | -2.2E+00 | 3.4E-08 | -5.9E-01 | 4.0E-01 | 8 |
| DBNDD2 | -2.9E+00 | 9.4E-04 | -4.0E+00 | 2.3E-04 | -1.0E+00 | 7.3E-01 | 8 |
| PIGT | -2.5E+00 | 3.3E-02 | -3.0E+00 | 4.9E-02 | -4.6E-01 | 9.9E-01 | 8 |
| PUS7L | -8.2E-01 | 2.0E-01 | -2.1E+00 | 2.1E-03 | -1.3E+00 | 1.6E-01 | 8 |
| ACAD8 | -9.6E-01 | 2.2E-01 | -1.9E+00 | 2.9E-02 | -9.0E-01 | 5.7E-01 | 8 |
| NEDD9 | -8.0E-01 | 1.5E-03 | -7.8E-01 | 7.0E-03 | 2.7E-02 | 9.9E-01 | 8 |
| CD58 | -2.8E+00 | 1.2E-03 | -3.6E+00 | 1.5E-03 | -8.2E-01 | 9.0E-01 | 8 |
| SIRT5 | -1.7E+00 | 5.5E-06 | -2.1E+00 | 1.0E-06 | -3.7E-01 | 8.2E-01 | 8 |
| NA | -2.3E+00 | 1.9E-05 | -2.4E+00 | 1.2E-04 | -7.8E-02 | 9.9E-01 | 8 |
| GMPR | -3.1E+00 | 7.2E-02 | -5.4E+00 | 1.3E-02 | -2.3E+00 | 6.5E-01 | 8 |
| PPRC1 | -1.5E+00 | 1.0E-02 | -2.0E+00 | 4.0E-03 | -4.5E-01 | 9.1E-01 | 8 |
| FAM173B | -1.6E+00 | 2.2E-02 | -3.1E+00 | 8.2E-04 | -1.5E+00 | 3.1E-01 | 8 |
| CD81 | -1.6E+00 | 4.7E-05 | -1.5E+00 | 3.6E-04 | 4.3E-02 | 9.9E-01 | 8 |
| FASTKD3 | -8.4E-01 | 4.4E-02 | -1.9E+00 | 5.7E-05 | -1.0E+00 | 8.2E-02 | 8 |
| CUL3 | -8.7E-01 | 3.3E-02 | -1.6E+00 | 2.5E-04 | -7.5E-01 | 2.4E-01 | 8 |
| RBM15B | -3.3E+00 | 3.0E-02 | -3.7E+00 | 5.2E-02 | -4.5E-01 | 9.9E-01 | 8 |
| MANF | -2.5E+00 | 3.6E-02 | -4.3E+00 | 5.2E-04 | -1.7E+00 | 3.6E-01 | 8 |
| ELMO2 | -4.2E-01 | 1.6E-01 | -7.6E-01 | 1.0E-02 | -3.3E-01 | 5.1E-01 | 8 |
| DYNLL1 | -1.9E+00 | 2.2E-04 | -1.9E+00 | 7.4E-04 | -2.3E-02 | 9.9E-01 | 8 |
| NA | -8.7E-01 | 4.6E-03 | -1.4E+00 | 2.8E-05 | -5.0E-01 | 3.3E-01 | 8 |
| RNF19B | -1.6E+00 | 5.0E-02 | -2.0E+00 | 3.5E-02 | -4.3E-01 | 9.9E-01 | 8 |
| CCDC80 | -1.0E+00 | 2.4E-04 | -1.1E+00 | 3.7E-04 | -6.3E-02 | 9.9E-01 | 8 |
| LRRC16A | -8.9E-01 | 4.8E-02 | -1.0E+00 | 4.2E-02 | -1.4E-01 | 9.9E-01 | 8 |
| NA | -2.0E+00 | 5.9E-04 | -2.0E+00 | 2.9E-03 | 1.0E-02 | 9.9E-01 | 8 |
| RGS1 | -6.7E-01 | 2.8E-01 | -1.5E+00 | 7.7E-03 | -8.3E-01 | 2.9E-01 | 8 |
| NA | -3.7E+00 | 1.3E-02 | -7.1E+00 | 5.4E-04 | -3.4E+00 | 2.7E-01 | 8 |
| HYAL2 | -2.6E+00 | 1.5E-01 | -5.2E+00 | 2.3E-02 | -2.6E+00 | 5.2E-01 | 8 |
| LINS | -1.3E+00 | 7.0E-02 | -1.6E+00 | 4.7E-02 | -3.3E-01 | 9.9E-01 | 8 |
| CCND1 | -9.4E-01 | 8.8E-02 | -1.3E+00 | 3.0E-02 | -3.2E-01 | 9.4E-01 | 8 |
| KCTD3 | -5.7E-01 | 6.7E-02 | -1.1E+00 | 5.5E-04 | -5.1E-01 | 2.5E-01 | 8 |
| ATP6V1A | -1.1E+00 | 1.5E-03 | -1.1E+00 | 3.5E-03 | -4.6E-02 | 9.9E-01 | 8 |
| CAPRIN2 | -5.8E-01 | 7.1E-02 | -9.5E-01 | 4.5E-03 | -3.7E-01 | 5.5E-01 | 8 |
| NA | -2.9E+00 | 5.3E-04 | -2.6E+00 | 7.0E-03 | 2.7E-01 | 9.9E-01 | 8 |
| SBDS | -1.3E+00 | 1.3E-02 | -1.8E+00 | 1.0E-03 | -5.4E-01 | 6.8E-01 | 8 |
| ZBTB20 | -1.7E+00 | 7.2E-04 | -2.3E+00 | 4.5E-05 | -6.7E-01 | 6.0E-01 | 8 |
| ERGIC2 | -1.2E+00 | 2.6E-02 | -1.2E+00 | 4.2E-02 | -5.5E-02 | 9.9E-01 | 8 |
| NA | -9.7E-01 | 7.1E-05 | -9.9E-01 | 2.5E-04 | -2.0E-02 | 9.9E-01 | 8 |
| NA | -9.7E-01 | 7.1E-05 | -9.9E-01 | 2.5E-04 | -2.0E-02 | 9.9E-01 | 8 |
| KCTD10 | -7.0E-01 | 1.4E-03 | -1.6E+00 | 1.1E-12 | -9.0E-01 | 1.7E-04 | 8 |
| RBM6 | -6.5E-01 | 4.5E-02 | -7.6E-01 | 3.8E-02 | -1.0E-01 | 9.9E-01 | 8 |
| PTHLH | -1.4E+00 | 1.9E-01 | -2.4E+00 | 1.7E-02 | -1.0E+00 | 5.8E-01 | 8 |
| PEF1 | -2.7E+00 | 4.4E-02 | -3.8E+00 | 3.5E-02 | -1.1E+00 | 9.2E-01 | 8 |
| SLC25A6 | -2.4E+00 | 3.8E-02 | -2.7E+00 | 4.5E-02 | -2.3E-01 | 9.9E-01 | 8 |
| CHD2 | -6.2E-01 | 1.3E-02 | -1.2E+00 | 8.9E-07 | -6.1E-01 | 4.0E-02 | 8 |
| DAD1 | -2.4E+00 | 3.3E-02 | -3.5E+00 | 6.8E-03 | -1.2E+00 | 7.3E-01 | 8 |
| NA | -3.2E+00 | 6.2E-04 | -3.2E+00 | 2.2E-03 | -1.7E-02 | 9.9E-01 | 8 |
| MRPL52 | -1.7E+00 | 1.1E-02 | -2.4E+00 | 4.2E-03 | -6.8E-01 | 8.2E-01 | 8 |
| PRMT5 | -2.5E+00 | 7.4E-03 | -2.8E+00 | 1.5E-02 | -3.3E-01 | 9.9E-01 | 8 |
| CSE1L | -9.4E-01 | 5.3E-03 | -9.1E-01 | 2.0E-02 | 2.8E-02 | 9.9E-01 | 8 |
| AJUBA | -1.4E+00 | 4.4E-02 | -1.3E+00 | 1.2E-01 | 6.1E-02 | 9.9E-01 | 8 |
| GGPS1 | -5.9E-01 | 4.3E-01 | -1.6E+00 | 1.9E-02 | -1.0E+00 | 2.9E-01 | 8 |
| CDK2AP2 | -2.8E+00 | 1.5E-02 | -3.6E+00 | 1.4E-02 | -8.2E-01 | 9.6E-01 | 8 |
| CCDC109B | -1.9E+00 | 5.0E-02 | -2.0E+00 | 8.4E-02 | -1.2E-01 | 9.9E-01 | 8 |
| SCN9A | -9.3E-01 | 3.6E-03 | -1.1E+00 | 1.5E-03 | -1.8E-01 | 9.8E-01 | 8 |
| PSMD12 | -3.5E-01 | 3.1E-01 | -8.5E-01 | 5.7E-03 | -5.0E-01 | 2.2E-01 | 8 |
| COA6 | -1.8E+00 | 6.3E-02 | -3.0E+00 | 2.0E-02 | -1.2E+00 | 6.9E-01 | 8 |
| PPP1CA | -1.3E+00 | 4.1E-03 | -1.6E+00 | 1.1E-03 | -3.1E-01 | 9.2E-01 | 8 |
| NA | -1.1E+00 | 1.4E-01 | -2.0E+00 | 2.3E-02 | -8.7E-01 | 6.4E-01 | 8 |
| KPNA2 | -4.1E+00 | 4.0E-03 | -7.2E+00 | 2.8E-05 | -3.1E+00 | 2.1E-01 | 8 |
| CEP95 | -1.4E+00 | 1.4E-02 | -1.9E+00 | 2.6E-03 | -5.1E-01 | 8.0E-01 | 8 |
| DDX5 | -1.1E+00 | 5.3E-03 | -1.9E+00 | 7.1E-06 | -7.7E-01 | 1.8E-01 | 8 |
| DDX5 | -1.1E+00 | 5.3E-03 | -1.9E+00 | 7.1E-06 | -7.7E-01 | 1.8E-01 | 8 |
| DDX5 | -1.1E+00 | 5.3E-03 | -1.9E+00 | 7.1E-06 | -7.7E-01 | 1.8E-01 | 8 |
| EXOC8 | -1.9E+00 | 1.6E-01 | -4.3E+00 | 2.0E-02 | -2.4E+00 | 4.3E-01 | 8 |
| PRPF3 | -1.4E+00 | 2.0E-04 | -1.8E+00 | 2.8E-05 | -3.9E-01 | 7.7E-01 | 8 |
| IGSF9 | -1.4E+00 | 9.7E-04 | -1.6E+00 | 9.5E-04 | -2.0E-01 | 9.9E-01 | 8 |
| LAMB3 | -2.8E+00 | 1.8E-03 | -3.0E+00 | 2.7E-03 | -1.9E-01 | 9.9E-01 | 8 |
| PARP9 | -1.2E+00 | 4.7E-02 | -2.2E+00 | 6.5E-04 | -9.5E-01 | 3.3E-01 | 8 |
| RFC2 | -6.9E-01 | 2.3E-01 | -1.2E+00 | 3.8E-02 | -5.6E-01 | 6.6E-01 | 8 |
| DIEXF | -7.6E-01 | 2.3E-02 | -1.8E+00 | 2.8E-06 | -1.0E+00 | 2.6E-02 | 8 |

| | | | | | | | |
|---|---|---|---|---|---|---|---|
| IRF9 | -2.3E+00 | 7.1E-02 | -3.9E+00 | 1.9E-02 | -1.5E+00 | 7.0E-01 | 8 |
| MCL1 | -1.7E+00 | 2.0E-08 | -2.0E+00 | 1.1E-10 | -3.5E-01 | 6.3E-01 | 8 |
| PTPLB | -1.7E+00 | 3.3E-02 | -1.6E+00 | 9.5E-02 | 6.9E-02 | 9.9E-01 | 8 |
| NEDD8 | -1.1E+00 | 2.3E-03 | -1.1E+00 | 6.3E-03 | -1.3E-02 | 9.9E-01 | 8 |
| IP6K2 | -1.7E+00 | 1.8E-03 | -3.2E+00 | 8.3E-07 | -1.5E+00 | 8.1E-02 | 8 |
| TIAL1 | -3.4E+00 | 2.7E-03 | -3.6E+00 | 6.0E-03 | -1.1E-01 | 9.9E-01 | 8 |
| CTSZ | -2.9E+00 | 2.6E-05 | -4.1E+00 | 1.5E-06 | -1.1E+00 | 4.9E-01 | 8 |
| ZNRD1 | -3.5E+00 | 1.9E-02 | -3.6E+00 | 5.6E-02 | -1.3E-01 | 9.9E-01 | 8 |
| LDHB | -2.9E+00 | 4.8E-04 | -4.0E+00 | 4.8E-05 | -1.1E+00 | 6.1E-01 | 8 |
| BAG3 | -1.1E+00 | 1.2E-02 | -9.5E-01 | 8.2E-02 | 1.7E-01 | 9.9E-01 | 8 |
| ABCF3 | -2.3E+00 | 1.6E-03 | -2.7E+00 | 1.3E-03 | -4.4E-01 | 9.9E-01 | 8 |
| TATDN3 | -2.8E+00 | 5.8E-03 | -3.1E+00 | 1.7E-02 | -3.5E-01 | 9.9E-01 | 8 |
| BLM | -1.4E+00 | 7.1E-02 | -2.0E+00 | 2.2E-02 | -6.1E-01 | 8.5E-01 | 8 |
| NA | -3.1E+00 | 2.7E-04 | -3.6E+00 | 5.0E-04 | -4.4E-01 | 9.9E-01 | 8 |
| NA | -3.1E+00 | 2.7E-04 | -3.6E+00 | 5.0E-04 | -4.4E-01 | 9.9E-01 | 8 |
| NA | -3.1E+00 | 2.7E-04 | -3.6E+00 | 5.0E-04 | -4.4E-01 | 9.9E-01 | 8 |
| SUMO1 | -2.9E+00 | 1.9E-08 | -2.7E+00 | 2.3E-06 | 2.1E-01 | 9.9E-01 | 8 |
| C4orf3 | -2.5E+00 | 1.2E-04 | -2.7E+00 | 2.0E-04 | -1.8E-01 | 9.9E-01 | 8 |
| ARCN1 | -1.8E+00 | 1.5E-02 | -2.6E+00 | 1.3E-03 | -7.3E-01 | 7.1E-01 | 8 |
| DERA | -2.0E+00 | 1.3E-02 | -1.7E+00 | 6.8E-02 | 2.3E-01 | 9.9E-01 | 8 |
| DPP3 | -1.9E+00 | 2.0E-02 | -2.8E+00 | 2.6E-03 | -9.1E-01 | 6.9E-01 | 8 |
| SCFD1 | -1.4E+00 | 3.4E-04 | -1.5E+00 | 6.4E-04 | -8.0E-02 | 9.9E-01 | 8 |
| RBBP7 | -1.8E+00 | 9.4E-06 | -2.9E+00 | 1.1E-10 | -1.1E+00 | 6.8E-02 | 8 |
| ANK3 | -8.4E-01 | 6.2E-06 | -8.3E-01 | 4.7E-05 | 9.5E-03 | 9.9E-01 | 8 |
| CGN | -5.0E-01 | 1.7E-01 | -1.1E+00 | 1.7E-03 | -5.9E-01 | 2.0E-01 | 8 |
| NIT1 | -1.9E+00 | 8.7E-04 | -1.7E+00 | 1.1E-02 | 2.2E-01 | 9.9E-01 | 8 |
| DIDO1 | -5.3E-01 | 8.0E-02 | -1.2E+00 | 5.8E-05 | -6.8E-01 | 6.3E-02 | 8 |
| PSMC5 | -2.2E+00 | 2.4E-04 | -2.7E+00 | 6.5E-05 | -4.6E-01 | 9.1E-01 | 8 |
| EDRF1 | -8.1E-01 | 2.2E-01 | -1.5E+00 | 2.6E-02 | -6.9E-01 | 5.8E-01 | 8 |
| ADAMTS4 | -1.7E+00 | 7.0E-03 | -1.8E+00 | 1.3E-02 | -7.0E-02 | 9.9E-01 | 8 |
| MORC2 | -1.4E+00 | 4.3E-02 | -2.2E+00 | 8.5E-03 | -7.3E-01 | 7.3E-01 | 8 |
| NA | -2.4E+00 | 2.9E-02 | -3.6E+00 | 2.8E-02 | -1.2E+00 | 8.7E-01 | 8 |
| PDE8A | -1.3E+00 | 1.2E-02 | -1.3E+00 | 2.1E-02 | -6.3E-02 | 9.9E-01 | 8 |
| WDR53 | -1.0E+00 | 3.3E-02 | -1.2E+00 | 3.0E-02 | -1.7E-01 | 9.9E-01 | 8 |
| BANF1 | -2.4E+00 | 9.2E-04 | -3.1E+00 | 2.9E-04 | -6.5E-01 | 8.6E-01 | 8 |
| RNASEL | -1.1E+00 | 5.5E-03 | -1.3E+00 | 4.2E-03 | -1.7E-01 | 9.9E-01 | 8 |
| GALE | -1.9E+00 | 1.9E-02 | -3.2E+00 | 4.3E-04 | -1.3E+00 | 4.0E-01 | 8 |
| NRP2 | -8.4E-01 | 3.1E-02 | -9.2E-01 | 3.5E-02 | -7.8E-02 | 9.9E-01 | 8 |
| DHX9 | -9.1E-01 | 9.6E-05 | -9.9E-01 | 1.0E-04 | -7.9E-02 | 9.9E-01 | 8 |
| DDX47 | -1.1E+00 | 9.3E-03 | -1.7E+00 | 6.9E-04 | -5.2E-01 | 6.3E-01 | 8 |
| PTPN23 | -3.9E-01 | 3.4E-01 | -8.4E-01 | 2.0E-02 | -4.5E-01 | 4.1E-01 | 8 |
| SRP54 | -9.3E-01 | 8.8E-03 | -1.1E+00 | 7.8E-03 | -1.5E-01 | 9.9E-01 | 8 |
| MGST3 | -2.9E+00 | 7.3E-02 | -3.7E+00 | 3.5E-02 | -7.9E-01 | 9.9E-01 | 8 |
| PCNXL3 | -8.5E-01 | 1.2E-02 | -9.8E-01 | 1.2E-02 | -1.2E-01 | 9.9E-01 | 8 |
| PSMA6 | -1.6E+00 | 5.7E-03 | -1.6E+00 | 2.3E-02 | 7.9E-02 | 9.9E-01 | 8 |
| RHOC | -2.5E+00 | 2.1E-03 | -2.8E+00 | 2.2E-03 | -3.1E-01 | 9.9E-01 | 8 |
| MOV10 | -1.3E+00 | 9.4E-03 | -1.3E+00 | 2.3E-02 | -3.9E-02 | 9.9E-01 | 8 |
| ATP13A3 | -1.2E+00 | 4.3E-04 | -1.1E+00 | 3.6E-03 | 7.2E-02 | 9.9E-01 | 8 |
| NA | -3.6E+00 | 7.5E-04 | -4.5E+00 | 1.7E-03 | -9.7E-01 | 9.3E-01 | 8 |
| HNRNPH3 | -2.7E+00 | 2.0E-02 | -3.0E+00 | 2.0E-02 | -3.0E-01 | 9.9E-01 | 8 |
| RBM45 | -2.2E+00 | 5.6E-04 | -2.0E+00 | 7.4E-03 | 2.2E-01 | 9.9E-01 | 8 |
| MBIP | -2.3E+00 | 1.2E-02 | -3.0E+00 | 2.6E-03 | -6.6E-01 | 8.9E-01 | 8 |
| GK | -5.6E-01 | 5.0E-02 | -8.1E-01 | 8.1E-03 | -2.4E-01 | 7.7E-01 | 8 |
| SPATA5L1 | -1.2E+00 | 5.0E-02 | -1.5E+00 | 3.0E-02 | -3.4E-01 | 9.8E-01 | 8 |
| KIAA1279 | -8.5E-01 | 2.6E-03 | -1.6E+00 | 1.9E-07 | -7.1E-01 | 5.8E-02 | 8 |
| NA | -1.5E+00 | 1.6E-02 | -1.5E+00 | 4.4E-02 | 2.9E-02 | 9.9E-01 | 8 |
| AIFM2 | -1.4E+00 | 3.9E-03 | -1.4E+00 | 1.6E-02 | 4.9E-02 | 9.9E-01 | 8 |
| PPA1 | -2.6E+00 | 2.8E-03 | -3.0E+00 | 2.9E-03 | -3.9E-01 | 9.9E-01 | 8 |
| SLC12A9 | -2.2E+00 | 2.5E-03 | -2.0E+00 | 1.6E-02 | 1.0E-01 | 9.9E-01 | 8 |
| ATP6AP2 | -2.8E+00 | 3.2E-05 | -2.4E+00 | 1.2E-03 | 3.4E-01 | 9.9E-01 | 8 |
| EHD1 | -1.4E+00 | 5.6E-02 | -2.4E+00 | 8.1E-03 | -9.9E-01 | 6.0E-01 | 8 |
| NA | -3.7E+00 | 3.1E-03 | -5.9E+00 | 7.6E-04 | -2.2E+00 | 6.0E-01 | 8 |
| MGAT2 | -6.6E-01 | 7.1E-02 | -1.7E+00 | 2.9E-06 | -1.0E+00 | 1.3E-02 | 8 |
| POP7 | -1.1E+00 | 3.5E-02 | -1.4E+00 | 3.1E-02 | -2.4E-01 | 9.9E-01 | 8 |
| L2HGDH | -2.7E+00 | 1.3E-02 | -4.6E+00 | 1.4E-03 | -1.9E+00 | 4.6E-01 | 8 |
| ENY2 | -1.2E+00 | 3.4E-04 | -1.1E+00 | 5.5E-03 | 1.2E-01 | 9.9E-01 | 8 |
| CRAMP1L | -1.3E+00 | 4.0E-02 | -1.5E+00 | 3.2E-02 | -2.3E-01 | 9.9E-01 | 8 |
| GNB2 | -3.3E+00 | 1.1E-01 | -6.5E+00 | 9.2E-03 | -3.2E+00 | 4.4E-01 | 8 |
| HDAC5 | -8.7E-01 | 4.0E-02 | -9.4E-01 | 5.5E-02 | -6.7E-02 | 9.9E-01 | 8 |
| NA | -2.6E+00 | 1.2E-03 | -3.0E+00 | 1.4E-03 | -3.8E-01 | 9.9E-01 | 8 |
| TSC22D4 | -1.6E+00 | 2.2E-02 | -2.2E+00 | 1.1E-02 | -5.4E-01 | 9.2E-01 | 8 |
| COPE | -4.1E-01 | 1.6E-01 | -6.8E-01 | 1.9E-02 | -2.7E-01 | 6.4E-01 | 8 |
| SPSB2 | -1.6E+00 | 4.9E-02 | -1.3E+00 | 2.1E-01 | 2.8E-01 | 9.9E-01 | 8 |
| NA | -1.6E+00 | 4.0E-04 | -1.7E+00 | 6.9E-04 | -9.8E-02 | 9.9E-01 | 8 |
| NA | -2.7E+00 | 2.3E-03 | -3.9E+00 | 8.5E-04 | -1.1E+00 | 7.2E-01 | 8 |
| RPL27 | -2.6E+00 | 3.9E-03 | -2.4E+00 | 2.7E-02 | 2.9E-01 | 9.9E-01 | 8 |
| PNKD | -2.9E+00 | 2.8E-03 | -3.3E+00 | 7.7E-03 | -3.5E-01 | 9.9E-01 | 8 |
| SLC35F2 | -1.7E+00 | 1.6E-04 | -1.5E+00 | 3.7E-03 | 2.1E-01 | 9.9E-01 | 8 |
| AP4M1 | -2.2E+00 | 1.3E-01 | -5.0E+00 | 1.2E-02 | -2.8E+00 | 3.7E-01 | 8 |
| PIGC | -2.1E+00 | 4.7E-02 | -2.9E+00 | 2.2E-02 | -8.2E-01 | 9.0E-01 | 8 |
| NA | -2.1E+00 | 4.1E-02 | -2.1E+00 | 9.5E-02 | -3.1E-02 | 9.9E-01 | 8 |
| ANXA7 | -2.6E+00 | 1.0E-04 | -2.7E+00 | 2.5E-04 | -1.5E-01 | 9.9E-01 | 8 |
| EID1 | -1.0E+00 | 9.4E-02 | -1.4E+00 | 4.2E-02 | -3.5E-01 | 9.6E-01 | 8 |
| SKIL | -1.3E+00 | 1.8E-08 | -1.3E+00 | 3.6E-08 | -6.7E-02 | 9.9E-01 | 8 |
| RNF4 | -9.4E-01 | 3.1E-04 | -1.2E+00 | 2.2E-05 | -2.9E-01 | 7.0E-01 | 8 |
| CHCHD1 | -2.4E+00 | 1.5E-04 | -2.4E+00 | 5.7E-04 | -2.0E-02 | 9.9E-01 | 8 |
| HAUS3 | -1.1E+00 | 6.5E-03 | -1.2E+00 | 1.3E-02 | -6.4E-02 | 9.9E-01 | 8 |
| RPL22L1 | -8.0E-01 | 5.0E-02 | -1.7E+00 | 1.0E-05 | -9.5E-01 | 4.3E-02 | 8 |
| NA | -4.0E+00 | 3.2E-04 | -5.8E+00 | 4.3E-04 | -1.9E+00 | 6.4E-01 | 8 |
| GCH1 | -3.3E+00 | 2.4E-03 | -3.5E+00 | 5.0E-03 | -1.2E-01 | 9.9E-01 | 8 |
| TACC3 | -2.4E+00 | 1.4E-02 | -2.9E+00 | 1.0E-02 | -5.0E-01 | 9.9E-01 | 8 |
| NOP2 | -6.3E-01 | 2.3E-01 | -1.3E+00 | 1.0E-02 | -7.1E-01 | 3.7E-01 | 8 |
| DCUN1D5 | -2.8E+00 | 6.7E-05 | -3.4E+00 | 2.6E-05 | -6.2E-01 | 8.7E-01 | 8 |
| KIFAP3 | -9.8E-01 | 9.9E-03 | -1.3E+00 | 1.8E-03 | -3.1E-01 | 8.3E-01 | 8 |

| | | | | | | | |
|---|---|---|---|---|---|---|---|
| LAP3 | -1.6E+00 | 1.2E-01 | -3.3E+00 | 2.1E-02 | -1.7E+00 | 5.0E-01 | 8 |
| GABPB1 | -8.7E-01 | 4.5E-02 | -1.1E+00 | 2.1E-02 | -2.5E-01 | 9.7E-01 | 8 |
| ZZEF1 | -1.5E+00 | 2.1E-03 | -1.4E+00 | 1.4E-02 | 1.0E-01 | 9.9E-01 | 8 |
| DNAJB2 | -2.6E+00 | 1.4E-02 | -2.4E+00 | 5.8E-02 | 1.8E-01 | 9.9E-01 | 8 |
| SLC19A2 | -2.3E+00 | 1.1E-02 | -2.9E+00 | 1.3E-02 | -6.3E-01 | 9.8E-01 | 8 |
| TRRAP | -1.5E+00 | 1.8E-03 | -1.5E+00 | 5.4E-03 | -4.5E-02 | 9.9E-01 | 8 |
| SFT2D2 | -1.4E+00 | 3.7E-01 | -4.0E+00 | 4.9E-02 | -2.5E+00 | 4.2E-01 | 8 |
| SPPL2A | -7.0E-01 | 6.1E-03 | -7.4E-01 | 1.0E-02 | -4.3E-02 | 9.9E-01 | 8 |
| BRI3 | -2.5E+00 | 1.2E-03 | -2.4E+00 | 8.2E-03 | 1.2E-01 | 9.9E-01 | 8 |
| KIAA0586 | -1.9E+00 | 5.0E-04 | -2.2E+00 | 3.1E-04 | -2.7E-01 | 9.9E-01 | 8 |
| SAMD8 | -1.1E+00 | 3.0E-04 | -1.3E+00 | 2.4E-04 | -1.4E-01 | 9.9E-01 | 8 |
| NA | -8.1E-01 | 1.6E-01 | -1.5E+00 | 8.2E-03 | -6.8E-01 | 4.6E-01 | 8 |
| NA | -8.1E-01 | 1.6E-01 | -1.5E+00 | 8.2E-03 | -6.8E-01 | 4.6E-01 | 8 |
| NA | -8.1E-01 | 1.6E-01 | -1.5E+00 | 8.2E-03 | -6.8E-01 | 4.6E-01 | 8 |
| NA | -8.1E-01 | 1.6E-01 | -1.5E+00 | 8.2E-03 | -6.8E-01 | 4.6E-01 | 8 |
| POLR2G | -2.6E+00 | 4.3E-03 | -3.2E+00 | 3.8E-03 | -5.4E-01 | 9.9E-01 | 8 |
| DAAM1 | -1.3E+00 | 4.6E-06 | -2.1E+00 | 3.4E-13 | -8.2E-01 | 1.6E-02 | 8 |
| JKAMP | -9.0E-01 | 3.6E-03 | -9.2E-01 | 8.8E-03 | -1.8E-02 | 9.9E-01 | 8 |
| NXF1 | -1.3E+00 | 1.1E-01 | -2.6E+00 | 8.3E-03 | -1.3E+00 | 4.5E-01 | 8 |
| RHNO1 | -2.9E+00 | 2.2E-02 | -3.2E+00 | 4.9E-02 | -3.7E-01 | 9.9E-01 | 8 |
| LEO1 | -1.3E+00 | 7.3E-03 | -1.6E+00 | 3.4E-03 | -2.8E-01 | 9.8E-01 | 8 |
| SIX4 | -1.4E+00 | 8.9E-02 | -1.8E+00 | 3.4E-02 | -4.4E-01 | 9.5E-01 | 8 |
| SGMS1 | -4.6E-01 | 8.9E-02 | -6.5E-01 | 2.2E-02 | -1.9E-01 | 8.5E-01 | 8 |
| C19ORF42 | -1.3E+00 | 4.5E-02 | -1.4E+00 | 8.5E-02 | -6.4E-02 | 9.9E-01 | 8 |
| SLC38A6 | -1.8E+00 | 4.9E-03 | -2.1E+00 | 5.2E-03 | -2.5E-01 | 9.9E-01 | 8 |
| TTC9C | -1.9E+00 | 9.4E-04 | -2.0E+00 | 3.1E-03 | -1.0E-01 | 9.9E-01 | 8 |
| TIMM17B | -1.6E+00 | 3.5E-03 | -1.6E+00 | 1.2E-02 | -2.4E-02 | 9.9E-01 | 8 |
| LYAR | -7.4E-01 | 1.2E-02 | -1.5E+00 | 8.0E-07 | -7.8E-01 | 3.5E-02 | 8 |
| ZBTB49 | -2.2E+00 | 4.8E-04 | -1.9E+00 | 1.0E-02 | 3.1E-01 | 9.9E-01 | 8 |
| MSX1 | -8.9E-01 | 4.3E-02 | -1.6E+00 | 3.2E-04 | -7.1E-01 | 2.7E-01 | 8 |
| NA | -2.2E+00 | 9.4E-05 | -2.1E+00 | 9.5E-04 | 6.5E-02 | 9.9E-01 | 8 |
| INTS5 | -6.1E-01 | 5.6E-02 | -1.1E+00 | 9.6E-04 | -4.7E-01 | 3.5E-01 | 8 |
| EPS15L1 | -1.0E+00 | 2.5E-02 | -1.3E+00 | 1.7E-02 | -2.1E-01 | 9.9E-01 | 8 |
| CNP | -1.0E+00 | 1.8E-02 | -1.4E+00 | 2.0E-03 | -4.1E-01 | 7.3E-01 | 8 |
| C7orf26 | -1.9E+00 | 1.3E-03 | -2.7E+00 | 1.7E-04 | -7.4E-01 | 6.8E-01 | 8 |
| WDR45 | -3.4E+00 | 2.3E-03 | -4.5E+00 | 1.2E-03 | -1.2E+00 | 8.2E-01 | 8 |
| NA | -2.5E+00 | 3.2E-02 | -2.6E+00 | 5.5E-02 | -8.2E-02 | 9.9E-01 | 8 |
| ACLY | -5.1E-01 | 1.2E-01 | -9.0E-01 | 5.0E-03 | -3.9E-01 | 4.6E-01 | 8 |
| DESI2 | -2.2E+00 | 5.6E-04 | -2.4E+00 | 6.8E-04 | -2.1E-01 | 9.9E-01 | 8 |
| PFN1 | -1.9E+00 | 1.6E-02 | -3.3E+00 | 1.4E-04 | -1.4E+00 | 2.8E-01 | 8 |
| SLC25A11 | -2.0E+00 | 7.6E-03 | -3.0E+00 | 2.1E-03 | -9.6E-01 | 7.0E-01 | 8 |
| CXCL16 | -1.9E+00 | 1.5E-04 | -1.7E+00 | 1.9E-03 | 1.4E-01 | 9.9E-01 | 8 |
| EPHA2 | -2.7E+00 | 3.9E-02 | -4.2E+00 | 1.5E-03 | -1.5E+00 | 5.2E-01 | 8 |
| CECR5 | -2.9E+00 | 1.1E-01 | -4.9E+00 | 3.7E-02 | -2.0E+00 | 7.5E-01 | 8 |
| GABARAP | -8.3E-01 | 1.1E-01 | -1.5E+00 | 7.1E-03 | -6.5E-01 | 5.0E-01 | 8 |
| CLDN7 | -1.4E+00 | 1.4E-02 | -2.0E+00 | 1.3E-03 | -6.8E-01 | 6.3E-01 | 8 |
| MRPL44 | -7.9E-01 | 2.0E-03 | -8.1E-01 | 5.3E-03 | -1.8E-02 | 9.9E-01 | 8 |
| PEX26 | -1.6E+00 | 3.0E-04 | -1.6E+00 | 1.3E-03 | -1.1E-03 | 1.0E+00 | 8 |
| NA | -8.9E-01 | 1.2E-01 | -1.3E+00 | 3.1E-02 | -3.8E-01 | 8.7E-01 | 8 |
| NA | -8.9E-01 | 1.2E-01 | -1.3E+00 | 3.1E-02 | -3.8E-01 | 8.7E-01 | 8 |
| C17ORF37 | -2.3E+00 | 5.1E-03 | -3.2E+00 | 1.8E-03 | -8.7E-01 | 8.0E-01 | 8 |
| TBC1D19 | -7.9E-01 | 1.4E-01 | -1.2E+00 | 2.9E-02 | -4.6E-01 | 7.6E-01 | 8 |
| PHF8 | -1.9E+00 | 5.2E-04 | -2.5E+00 | 2.9E-05 | -6.0E-01 | 6.8E-01 | 8 |
| ARG2 | -6.0E-01 | 1.5E-02 | -6.2E-01 | 2.7E-02 | -2.1E-02 | 9.9E-01 | 8 |
| MED1 | -8.0E-01 | 2.4E-04 | -8.5E-01 | 4.3E-04 | -5.0E-02 | 9.9E-01 | 8 |
| NAA38 | -2.5E+00 | 1.3E-02 | -2.3E+00 | 5.9E-02 | 2.3E-01 | 9.9E-01 | 8 |
| MFN2 | -2.1E+00 | 3.4E-04 | -2.7E+00 | 4.8E-05 | -5.8E-01 | 7.8E-01 | 8 |
| PLOD1 | -1.8E+00 | 5.5E-02 | -2.6E+00 | 2.4E-02 | -8.0E-01 | 8.7E-01 | 8 |
| CWC25 | -6.6E-01 | 6.2E-02 | -1.5E+00 | 2.7E-05 | -8.5E-01 | 5.0E-02 | 8 |
| PHKG2 | -1.2E+00 | 1.7E-03 | -1.3E+00 | 4.2E-03 | -6.4E-02 | 9.9E-01 | 8 |
| GTF2A2 | -2.0E+00 | 2.3E-02 | -3.5E+00 | 8.8E-04 | -1.5E+00 | 3.9E-01 | 8 |
| MED6 | -2.1E+00 | 3.3E-04 | -3.4E+00 | 2.4E-06 | -1.2E+00 | 2.8E-01 | 8 |
| NARG2 | -1.1E+00 | 1.5E-02 | -1.6E+00 | 1.8E-03 | -4.9E-01 | 6.9E-01 | 8 |
| RNF40 | -6.1E-01 | 2.5E-01 | -1.1E+00 | 4.9E-02 | -4.4E-01 | 7.1E-01 | 8 |
| GAK | -9.8E-01 | 7.6E-02 | -1.6E+00 | 9.8E-03 | -6.0E-01 | 6.6E-01 | 8 |
| MRPL24 | -2.6E+00 | 3.8E-02 | -5.1E+00 | 6.0E-03 | -2.6E+00 | 4.2E-01 | 8 |
| VKORC1 | -2.1E+00 | 7.8E-03 | -2.6E+00 | 3.7E-03 | -4.6E-01 | 9.8E-01 | 8 |
| C1orf85 | -2.3E+00 | 1.9E-02 | -2.5E+00 | 3.9E-02 | -1.7E-01 | 9.9E-01 | 8 |
| ARMC5 | -7.7E-01 | 1.3E-01 | -1.7E+00 | 1.5E-03 | -9.3E-01 | 2.0E-01 | 8 |
| IGF2BP1 | -2.0E+00 | 9.3E-04 | -1.8E+00 | 7.7E-03 | 1.1E-01 | 9.9E-01 | 8 |
| RANBP17 | -1.8E+00 | 7.3E-02 | -2.9E+00 | 2.5E-02 | -1.1E+00 | 7.8E-01 | 8 |
| MED12 | -2.2E+00 | 8.9E-03 | -1.8E+00 | 6.9E-02 | 3.3E-01 | 9.9E-01 | 8 |
| FOS | -2.1E+00 | 3.5E-03 | -1.9E+00 | 2.7E-02 | 2.3E-01 | 9.9E-01 | 8 |
| PPIB | -2.7E+00 | 2.6E-04 | -3.2E+00 | 5.4E-05 | -5.2E-01 | 9.1E-01 | 8 |
| PPP4C | -2.3E+00 | 8.3E-02 | -3.4E+00 | 1.5E-02 | -1.1E+00 | 7.9E-01 | 8 |
| MTFMT | -1.7E+00 | 6.1E-03 | -2.8E+00 | 1.3E-04 | -1.1E+00 | 3.9E-01 | 8 |
| IGDCC4 | -2.0E+00 | 1.7E-04 | -2.0E+00 | 5.4E-04 | -2.9E-02 | 9.9E-01 | 8 |
| PMVK | -1.7E+00 | 4.3E-02 | -1.6E+00 | 1.1E-01 | 4.5E-02 | 9.9E-01 | 8 |
| UPRT | -1.0E+00 | 1.5E-02 | -1.1E+00 | 2.1E-02 | -9.4E-02 | 9.9E-01 | 8 |
| UBAP2L | -9.1E-01 | 2.0E-01 | -1.9E+00 | 1.1E-02 | -1.0E+00 | 3.9E-01 | 8 |
| NA | -3.4E+00 | 1.2E-02 | -5.0E+00 | 3.1E-04 | -1.7E+00 | 5.1E-01 | 8 |
| TIPIN | -9.8E-01 | 4.1E-02 | -1.4E+00 | 5.4E-03 | -4.7E-01 | 7.2E-01 | 8 |
| GTF2A1 | -2.2E+00 | 5.6E-03 | -2.0E+00 | 2.6E-02 | 1.5E-01 | 9.9E-01 | 8 |
| APOOL | -1.3E+00 | 4.2E-02 | -2.0E+00 | 5.9E-03 | -7.2E-01 | 6.8E-01 | 8 |
| COIL | -8.3E-01 | 2.2E-03 | -1.3E+00 | 5.0E-06 | -4.9E-01 | 2.5E-01 | 8 |
| MRPS23 | -1.3E+00 | 1.7E-04 | -1.5E+00 | 2.0E-04 | -1.6E-01 | 9.9E-01 | 8 |
| NA | -1.8E+00 | 5.1E-02 | -2.2E+00 | 2.6E-02 | -4.3E-01 | 9.9E-01 | 8 |
| NA | -1.8E+00 | 5.1E-02 | -2.2E+00 | 2.6E-02 | -4.3E-01 | 9.9E-01 | 8 |
| ERN2 | -9.9E-01 | 1.2E-02 | -9.2E-01 | 5.1E-02 | 7.0E-02 | 9.9E-01 | 8 |
| EPG5 | -1.6E+00 | 1.8E-03 | -1.8E+00 | 1.2E-03 | -2.4E-01 | 9.9E-01 | 8 |
| PTRH2 | -3.7E-01 | 3.2E-01 | -7.2E-01 | 2.9E-02 | -3.6E-01 | 5.2E-01 | 8 |
| INTS2 | -1.3E+00 | 4.8E-04 | -1.7E+00 | 2.6E-05 | -4.1E-01 | 6.9E-01 | 8 |
| P2RY11 | -6.4E-01 | 4.7E-02 | -9.0E-01 | 1.0E-02 | -2.6E-01 | 8.2E-01 | 8 |
| P2RY11 | -6.4E-01 | 4.7E-02 | -9.0E-01 | 1.0E-02 | -2.6E-01 | 8.2E-01 | 8 |

| | | | | | | | |
|---|---|---|---|---|---|---|---|
| P2RY11 | -6.4E-01 | 4.7E-02 | -9.0E-01 | 1.0E-02 | -2.6E-01 | 8.2E-01 | 8 |
| NA | -6.1E-01 | 4.6E-01 | -1.5E+00 | 3.8E-02 | -8.4E-01 | 4.3E-01 | 8 |
| PSMD10 | -1.9E+00 | 1.7E-04 | -2.1E+00 | 9.4E-05 | -2.4E-01 | 9.9E-01 | 8 |
| C16orf62 | -9.8E-01 | 4.8E-02 | -1.2E+00 | 4.2E-02 | -2.0E-01 | 9.9E-01 | 8 |
| TMC7 | -3.1E-01 | 3.9E-01 | -7.6E-01 | 1.1E-02 | -4.5E-01 | 2.6E-01 | 8 |
| COQ7 | -1.5E+00 | 4.3E-02 | -2.0E+00 | 2.1E-02 | -4.8E-01 | 9.5E-01 | 8 |
| CAMSAP3 | -2.1E+00 | 1.4E-02 | -2.5E+00 | 1.9E-02 | -3.9E-01 | 9.9E-01 | 8 |
| NA | -3.5E+00 | 3.2E-03 | -5.0E+00 | 4.4E-03 | -1.5E+00 | 8.2E-01 | 8 |
| PPP1R13B | -1.6E+00 | 3.9E-04 | -1.6E+00 | 1.3E-03 | -2.7E-02 | 9.9E-01 | 8 |
| FBXO38 | -6.1E-01 | 1.2E-01 | -9.1E-01 | 3.5E-02 | -2.9E-01 | 8.5E-01 | 8 |
| PSMD11 | -2.2E+00 | 2.7E-02 | -3.4E+00 | 3.1E-03 | -1.2E+00 | 6.3E-01 | 8 |
| SNX18 | -3.1E+00 | 9.4E-03 | -2.7E+00 | 5.9E-02 | 3.7E-01 | 9.9E-01 | 8 |
| PARP8 | -2.5E+00 | 1.0E-03 | -2.8E+00 | 1.5E-03 | -3.6E-01 | 9.9E-01 | 8 |
| FLCN | -1.7E+00 | 4.1E-02 | -1.6E+00 | 1.1E-01 | 7.9E-02 | 9.9E-01 | 8 |
| TEFM | -1.3E+00 | 3.3E-02 | -2.0E+00 | 4.9E-03 | -6.7E-01 | 6.9E-01 | 8 |
| UTP6 | -7.8E-01 | 9.2E-03 | -1.1E+00 | 4.8E-04 | -3.6E-01 | 5.9E-01 | 8 |
| HNRNPM | -6.6E-01 | 4.4E-02 | -8.2E-01 | 2.1E-02 | -1.6E-01 | 9.9E-01 | 8 |
| MAP2K3 | -8.4E-01 | 6.7E-02 | -1.3E+00 | 4.1E-03 | -4.9E-01 | 5.7E-01 | 8 |
| MAP2K3 | -8.4E-01 | 6.7E-02 | -1.3E+00 | 4.1E-03 | -4.9E-01 | 5.7E-01 | 8 |
| DLGAP1 | -1.1E+00 | 4.8E-02 | -1.3E+00 | 4.8E-02 | -1.8E-01 | 9.9E-01 | 8 |
| FBXO4 | -1.1E+00 | 2.6E-02 | -1.7E+00 | 9.7E-04 | -6.4E-01 | 5.0E-01 | 8 |
| TRIP10 | -2.4E+00 | 4.5E-04 | -2.1E+00 | 7.5E-03 | 2.7E-01 | 9.9E-01 | 8 |
| IMPA2 | -2.6E+00 | 1.2E-03 | -3.7E+00 | 4.4E-04 | -1.1E+00 | 7.0E-01 | 8 |
| TUBB6 | -3.1E+00 | 1.0E-04 | -2.8E+00 | 1.5E-03 | 2.3E-01 | 9.9E-01 | 8 |
| TM2D1 | -4.1E+00 | 6.8E-03 | -5.9E+00 | 2.1E-03 | -1.8E+00 | 8.0E-01 | 8 |
| CEP76 | -2.6E+00 | 3.8E-02 | -3.1E+00 | 5.2E-02 | -5.5E-01 | 9.9E-01 | 8 |
| NA | -1.0E+00 | 9.3E-03 | -1.5E+00 | 5.0E-04 | -4.9E-01 | 5.7E-01 | 8 |
| LIPG | -9.9E-01 | 6.2E-02 | -1.4E+00 | 1.3E-02 | -4.2E-01 | 8.2E-01 | 8 |
| NA | -2.6E+00 | 9.5E-04 | -3.0E+00 | 3.7E-04 | -4.4E-01 | 9.8E-01 | 8 |
| NA | -2.6E+00 | 9.5E-04 | -3.0E+00 | 3.7E-04 | -4.4E-01 | 9.8E-01 | 8 |
| NA | -2.6E+00 | 9.5E-04 | -3.0E+00 | 3.7E-04 | -4.4E-01 | 9.8E-01 | 8 |
| NA | -2.6E+00 | 9.5E-04 | -3.0E+00 | 3.7E-04 | -4.4E-01 | 9.8E-01 | 8 |
| DYM | -1.4E+00 | 5.2E-05 | -1.2E+00 | 1.9E-03 | 1.6E-01 | 9.9E-01 | 8 |
| NA | -2.1E+00 | 5.4E-02 | -4.1E+00 | 1.2E-02 | -2.0E+00 | 5.1E-01 | 8 |
| HMOX2 | -6.2E-01 | 2.5E-01 | -1.4E+00 | 5.5E-03 | -8.3E-01 | 2.5E-01 | 8 |
| LSM5 | -3.0E+00 | 1.8E-03 | -6.2E+00 | 1.7E-04 | -3.2E+00 | 2.1E-01 | 8 |
| ATP6V0C | -1.6E+00 | 4.7E-02 | -3.4E+00 | 1.2E-03 | -1.9E+00 | 2.2E-01 | 8 |
| C16orf59 | -2.8E+00 | 6.2E-02 | -4.9E+00 | 1.2E-02 | -2.1E+00 | 6.6E-01 | 8 |
| GSG2 | -1.3E+00 | 1.4E-01 | -4.9E+00 | 2.5E-03 | -3.6E+00 | 7.9E-02 | 8 |
| DPH7 | -8.6E-01 | 2.5E-02 | -1.3E+00 | 2.5E-03 | -4.0E-01 | 6.8E-01 | 8 |
| MRPL41 | -2.4E+00 | 1.7E-02 | -2.7E+00 | 3.0E-02 | -3.1E-01 | 9.9E-01 | 8 |
| LSM7 | -7.0E-01 | 1.2E-01 | -9.4E-01 | 5.0E-02 | -2.4E-01 | 9.6E-01 | 8 |
| SSNA1 | -2.0E+00 | 5.5E-02 | -3.3E+00 | 1.6E-02 | -1.3E+00 | 6.9E-01 | 8 |
| C1orf174 | -1.8E+00 | 1.7E-02 | -2.2E+00 | 9.5E-03 | -3.9E-01 | 9.9E-01 | 8 |
| FLNA | -1.3E+00 | 1.8E-04 | -1.5E+00 | 8.1E-05 | -1.9E-01 | 9.9E-01 | 8 |
| POLR2E | -2.0E+00 | 5.6E-03 | -4.0E+00 | 5.6E-05 | -2.0E+00 | 1.4E-01 | 8 |
| AGPAT2 | -1.5E+00 | 7.8E-02 | -3.1E+00 | 6.0E-03 | -1.6E+00 | 3.9E-01 | 8 |
| SLC10A3 | -1.9E+00 | 2.0E-02 | -2.0E+00 | 3.7E-02 | -1.1E-01 | 9.9E-01 | 8 |
| PMPCA | -6.9E-01 | 3.9E-02 | -6.3E-01 | 1.3E-01 | 6.1E-02 | 9.9E-01 | 8 |
| NA | -2.4E+00 | 1.4E-01 | -5.0E+00 | 1.5E-02 | -2.6E+00 | 4.9E-01 | 8 |
| MRPL28 | -6.9E-01 | 3.1E-02 | -1.3E+00 | 7.7E-05 | -6.2E-01 | 1.6E-01 | 8 |
| SNAPC4 | -8.7E-01 | 3.6E-01 | -2.1E+00 | 3.3E-02 | -1.2E+00 | 4.2E-01 | 8 |
| NA | -2.2E+00 | 1.1E-04 | -2.3E+00 | 2.9E-04 | -9.3E-02 | 9.9E-01 | 8 |
| QSOX2 | -1.2E+00 | 7.5E-03 | -1.5E+00 | 2.1E-03 | -3.3E-01 | 9.0E-01 | 8 |
| POLR3K | -3.3E+00 | 3.2E-02 | -4.7E+00 | 1.8E-02 | -1.4E+00 | 8.7E-01 | 8 |
| RXRALPHA | -4.3E-01 | 9.6E-02 | -6.4E-01 | 1.5E-02 | -2.1E-01 | 7.5E-01 | 8 |
| PIEZO1 | -2.5E+00 | 4.5E-03 | -2.5E+00 | 1.5E-02 | 5.6E-02 | 9.9E-01 | 8 |
| MAP1LC3B2 | -2.4E+00 | 3.6E-02 | -4.0E+00 | 7.0E-04 | -1.6E+00 | 4.0E-01 | 8 |
| EMC8 | -2.3E+00 | 1.5E-05 | -2.1E+00 | 2.8E-04 | 1.5E-01 | 9.9E-01 | 8 |
| C16orf74 | -2.5E+00 | 4.9E-02 | -3.0E+00 | 7.9E-02 | -4.5E-01 | 9.9E-01 | 8 |
| LAMTOR5 | -2.5E+00 | 3.1E-03 | -2.6E+00 | 6.1E-03 | -1.5E-01 | 9.9E-01 | 8 |
| RNPC3 | -1.0E+00 | 7.5E-03 | -9.7E-01 | 3.6E-02 | 7.7E-02 | 9.9E-01 | 8 |
| TOR1A | -2.4E+00 | 2.0E-02 | -4.4E+00 | 7.1E-03 | -1.9E+00 | 5.5E-01 | 8 |
| CCBL1 | -2.4E+00 | 3.4E-02 | -5.0E+00 | 5.3E-03 | -2.6E+00 | 3.8E-01 | 8 |
| ZNRF1 | -2.7E+00 | 7.3E-04 | -3.7E+00 | 3.7E-05 | -1.0E+00 | 6.0E-01 | 8 |
| FAM129B | -7.9E-01 | 6.8E-02 | -1.3E+00 | 5.5E-03 | -5.1E-01 | 5.7E-01 | 8 |
| AARS | -2.0E+00 | 4.5E-04 | -2.1E+00 | 1.7E-03 | -3.0E-02 | 9.9E-01 | 8 |
| SLC2A8 | -1.5E+00 | 5.0E-02 | -1.6E+00 | 7.0E-02 | -1.5E-01 | 9.9E-01 | 8 |
| CDC7 | -1.7E+00 | 2.0E-04 | -1.9E+00 | 2.4E-04 | -1.6E-01 | 9.9E-01 | 8 |
| GTF2B | -1.8E+00 | 2.2E-05 | -2.2E+00 | 2.5E-06 | -4.2E-01 | 7.8E-01 | 8 |
| ZNF821 | -2.2E+00 | 1.4E-04 | -2.1E+00 | 2.0E-03 | 1.3E-01 | 9.9E-01 | 8 |
| IST1 | -1.1E+00 | 1.2E-03 | -2.5E+00 | 2.2E-10 | -1.3E+00 | 2.5E-03 | 8 |
| NA | -2.4E+00 | 1.1E-04 | -2.4E+00 | 7.5E-04 | 2.9E-02 | 9.9E-01 | 8 |
| CYR61 | -1.8E+00 | 7.1E-05 | -3.1E+00 | 1.7E-11 | -1.3E+00 | 1.8E-02 | 8 |
| CDH1 | -7.8E-01 | 7.0E-02 | -1.2E+00 | 1.4E-02 | -3.9E-01 | 7.7E-01 | 8 |
| SLC12A4 | -1.8E+00 | 4.8E-02 | -2.6E+00 | 1.6E-02 | -8.6E-01 | 8.1E-01 | 8 |
| DNAJB4 | -7.4E-01 | 4.8E-02 | -6.9E-01 | 1.4E-01 | 5.5E-02 | 9.9E-01 | 8 |
| ELMO3 | -1.7E+00 | 1.1E-05 | -1.6E+00 | 2.4E-04 | 1.3E-01 | 9.9E-01 | 8 |
| C16orf70 | -9.6E-01 | 5.1E-02 | -1.2E+00 | 3.9E-02 | -2.1E-01 | 9.9E-01 | 8 |
| CTH | -1.8E+00 | 5.4E-04 | -1.8E+00 | 2.4E-03 | 2.5E-02 | 9.9E-01 | 8 |
| RHOA | -2.3E+00 | 3.6E-03 | -3.8E+00 | 2.9E-05 | -1.4E+00 | 3.1E-01 | 8 |
| RBM8A | -7.9E-01 | 2.0E-02 | -7.2E-01 | 7.6E-02 | 6.9E-02 | 9.9E-01 | 8 |
| PRIM2 | -1.0E+00 | 8.9E-02 | -2.2E+00 | 1.3E-03 | -1.2E+00 | 2.3E-01 | 8 |
| PRIM2 | -1.0E+00 | 8.9E-02 | -2.2E+00 | 1.3E-03 | -1.2E+00 | 2.3E-01 | 8 |
| GLRX5 | -2.0E+00 | 1.2E-02 | -2.0E+00 | 3.7E-02 | 1.5E-02 | 9.9E-01 | 8 |
| ZBTB43 | -8.8E-01 | 9.0E-02 | -1.2E+00 | 4.2E-02 | -2.9E-01 | 9.7E-01 | 8 |
| SETDB1 | -1.3E+00 | 4.6E-03 | -1.1E+00 | 4.3E-02 | 1.8E-01 | 9.9E-01 | 8 |
| C9orf3 | -3.2E+00 | 3.6E-02 | -3.9E+00 | 2.1E-02 | -7.5E-01 | 9.9E-01 | 8 |
| GMPPB | -1.1E+00 | 3.1E-02 | -1.0E+00 | 1.0E-01 | 8.8E-02 | 9.9E-01 | 8 |
| GMPPB | -1.1E+00 | 3.1E-02 | -1.0E+00 | 1.0E-01 | 8.8E-02 | 9.9E-01 | 8 |
| GNAS | -2.4E+00 | 2.3E-05 | -2.8E+00 | 5.5E-06 | -4.2E-01 | 9.1E-01 | 8 |
| HERC3 | -9.6E-01 | 2.2E-02 | -1.0E+00 | 3.3E-02 | -7.6E-02 | 9.9E-01 | 8 |
| HERC3 | -9.6E-01 | 2.2E-02 | -1.0E+00 | 3.3E-02 | -7.6E-02 | 9.9E-01 | 8 |
| GSTA4 | -2.6E+00 | 9.4E-04 | -2.7E+00 | 2.9E-03 | -1.0E-01 | 9.9E-01 | 8 |

| | | | | | | | |
|---|---|---|---|---|---|---|---|
| MINOS1 | -2.7E+00 | 1.2E-03 | -3.1E+00 | 2.2E-03 | -3.8E-01 | 9.9E-01 | 8 |
| CFL1 | -2.8E+00 | 4.9E-03 | -5.9E+00 | 2.3E-05 | -3.1E+00 | 8.8E-02 | 8 |
| NA | -1.8E+00 | 2.5E-01 | -4.4E+00 | 3.2E-02 | -2.6E+00 | 4.4E-01 | 8 |
| NA | -1.8E+00 | 1.0E-02 | -1.8E+00 | 4.4E-02 | 6.0E-02 | 9.9E-01 | 8 |
| NA | -3.2E+00 | 1.2E-02 | -3.0E+00 | 4.6E-02 | 2.1E-01 | 9.9E-01 | 8 |
| IFNLR1 | -1.4E+00 | 1.1E-02 | -1.5E+00 | 2.1E-02 | -8.1E-02 | 9.9E-01 | 8 |
| HYLS1 | -7.7E-01 | 2.2E-01 | -1.9E+00 | 3.1E-03 | -1.2E+00 | 1.8E-01 | 8 |
| NA | -3.6E+00 | 3.8E-04 | -5.5E+00 | 8.4E-05 | -1.9E+00 | 4.9E-01 | 8 |
| C1orf122 | -2.5E+00 | 3.1E-03 | -2.8E+00 | 8.2E-03 | -3.3E-01 | 9.9E-01 | 8 |
| MRPS6 | -9.7E-01 | 1.1E-01 | -1.5E+00 | 2.5E-02 | -5.7E-01 | 7.5E-01 | 8 |
| ZBED5 | -6.7E-01 | 1.6E-03 | -8.8E-01 | 1.1E-04 | -2.2E-01 | 7.0E-01 | 8 |
| ZBED5 | -6.7E-01 | 1.6E-03 | -8.8E-01 | 1.1E-04 | -2.2E-01 | 7.0E-01 | 8 |
| TRIB2 | -8.2E-01 | 1.6E-04 | -1.2E+00 | 3.8E-07 | -3.5E-01 | 3.5E-01 | 8 |
| NA | -9.5E-01 | 1.5E-01 | -1.8E+00 | 1.4E-02 | -8.2E-01 | 5.3E-01 | 8 |
| NA | -2.0E+00 | 7.6E-05 | -2.0E+00 | 4.9E-04 | -2.7E-02 | 9.9E-01 | 8 |
| NA | -2.0E+00 | 7.6E-05 | -2.0E+00 | 4.9E-04 | -2.7E-02 | 9.9E-01 | 8 |
| NA | -3.6E+00 | 2.8E-06 | -4.2E+00 | 2.9E-06 | -5.6E-01 | 9.6E-01 | 8 |
| RBAK | -9.4E-01 | 8.6E-03 | -9.3E-01 | 2.6E-02 | 1.1E-02 | 9.9E-01 | 8 |
| CTGF | -4.3E+00 | 7.4E-05 | -4.9E+00 | 3.1E-05 | -6.2E-01 | 9.9E-01 | 8 |
| OSGIN2 | -1.8E+00 | 4.7E-07 | -1.8E+00 | 6.8E-06 | 4.1E-02 | 9.9E-01 | 8 |
| ZNF425 | -1.4E+00 | 8.7E-02 | -2.4E+00 | 1.5E-02 | -9.8E-01 | 6.5E-01 | 8 |
| TM2D2 | -2.3E+00 | 1.5E-04 | -2.5E+00 | 4.1E-04 | -1.6E-01 | 9.9E-01 | 8 |
| SRSF7 | -9.5E-01 | 1.9E-04 | -1.1E+00 | 1.5E-04 | -1.0E-01 | 9.9E-01 | 8 |
| SIX1 | -3.6E-01 | 4.6E-01 | -8.3E-01 | 3.4E-02 | -4.7E-01 | 4.2E-01 | 8 |
| LAMTOR3 | -1.5E+00 | 3.5E-03 | -2.2E+00 | 4.4E-05 | -7.4E-01 | 4.2E-01 | 8 |
| ZNF572 | -9.8E-01 | 3.9E-01 | -2.7E+00 | 2.3E-02 | -1.7E+00 | 3.1E-01 | 8 |
| TXN | -2.9E+00 | 2.7E-02 | -4.1E+00 | 3.0E-03 | -1.2E+00 | 7.3E-01 | 8 |
| SDC4 | -9.5E-01 | 6.9E-02 | -1.4E+00 | 7.0E-03 | -5.0E-01 | 6.8E-01 | 8 |
| PEX11A | -1.3E+00 | 1.9E-05 | -1.6E+00 | 7.5E-07 | -3.4E-01 | 6.7E-01 | 8 |
| NA | -1.8E+00 | 3.3E-05 | -3.8E+00 | 1.0E-10 | -2.0E+00 | 4.6E-03 | 8 |
| FTH1 | -5.2E+00 | 4.0E-03 | -6.3E+00 | 2.5E-03 | -1.1E+00 | 9.9E-01 | 8 |
| PDCL | -1.1E+00 | 3.8E-03 | -1.2E+00 | 4.9E-03 | -1.2E-01 | 9.9E-01 | 8 |
| ZNF782 | -8.5E-01 | 2.9E-01 | -1.8E+00 | 3.3E-02 | -9.2E-01 | 5.2E-01 | 8 |
| KIAA0040 | -2.7E+00 | 1.2E-01 | -5.2E+00 | 1.7E-02 | -2.5E+00 | 5.6E-01 | 8 |
| IDI1 | -1.5E+00 | 1.5E-05 | -1.5E+00 | 1.6E-04 | 6.0E-02 | 9.9E-01 | 8 |
| PNRC1 | -1.5E+00 | 3.2E-03 | -2.1E+00 | 1.3E-04 | -6.1E-01 | 5.9E-01 | 8 |
| C7orf60 | -1.3E+00 | 9.5E-03 | -1.9E+00 | 3.1E-04 | -6.1E-01 | 5.3E-01 | 8 |
| E2F3 | -6.8E-01 | 1.8E-02 | -7.9E-01 | 1.4E-02 | -1.1E-01 | 9.9E-01 | 8 |
| NA | -1.0E+00 | 5.3E-03 | -1.2E+00 | 2.9E-03 | -1.8E-01 | 9.9E-01 | 8 |
| NA | -1.0E+00 | 5.3E-03 | -1.2E+00 | 2.9E-03 | -1.8E-01 | 9.9E-01 | 8 |
| NA | -1.0E+00 | 5.3E-03 | -1.2E+00 | 2.9E-03 | -1.8E-01 | 9.9E-01 | 8 |
| NA | -1.0E+00 | 5.3E-03 | -1.2E+00 | 2.9E-03 | -1.8E-01 | 9.9E-01 | 8 |
| NA | -1.0E+00 | 5.3E-03 | -1.2E+00 | 2.9E-03 | -1.8E-01 | 9.9E-01 | 8 |
| NA | -2.5E+00 | 3.7E-02 | -1.8E+00 | 2.6E-01 | 6.9E-01 | 9.7E-01 | 8 |
| NA | -2.5E+00 | 3.7E-02 | -1.8E+00 | 2.6E-01 | 6.9E-01 | 9.7E-01 | 8 |
| GLIS3 | -6.9E-01 | 7.1E-02 | -9.0E-01 | 2.7E-02 | -2.1E-01 | 9.5E-01 | 8 |
| MAP2K7 | -5.4E-01 | 3.0E-01 | -1.2E+00 | 1.9E-02 | -6.4E-01 | 4.1E-01 | 8 |
| PRELID2 | -8.8E-01 | 1.1E-02 | -8.0E-01 | 5.6E-02 | 7.8E-02 | 9.9E-01 | 8 |
| ITPA | -1.6E+00 | 1.1E-03 | -1.9E+00 | 5.2E-04 | -3.6E-01 | 9.3E-01 | 8 |
| NUP35 | -7.0E-01 | 1.1E-01 | -1.4E+00 | 3.8E-03 | -6.6E-01 | 3.7E-01 | 8 |
| ZNF235 | -1.7E+00 | 6.0E-03 | -2.5E+00 | 4.5E-04 | -8.7E-01 | 5.5E-01 | 8 |
| RNF219 | -7.5E-01 | 2.5E-02 | -1.8E+00 | 2.4E-06 | -1.0E+00 | 2.1E-02 | 8 |
| GEM | -1.9E+00 | 1.7E-02 | -2.4E+00 | 6.3E-03 | -5.5E-01 | 9.1E-01 | 8 |
| SAR1B | -5.5E-01 | 7.4E-02 | -7.4E-01 | 2.5E-02 | -1.9E-01 | 9.2E-01 | 8 |
| CCDC58 | -6.3E-01 | 5.4E-02 | -8.7E-01 | 1.5E-02 | -2.4E-01 | 8.6E-01 | 8 |
| PDCD7 | -1.4E+00 | 5.0E-03 | -1.9E+00 | 3.4E-04 | -5.6E-01 | 6.5E-01 | 8 |
| C14orf142 | -1.8E+00 | 9.3E-03 | -1.7E+00 | 3.3E-02 | 7.2E-02 | 9.9E-01 | 8 |
| MYL12B | -2.3E+00 | 2.1E-06 | -2.2E+00 | 4.1E-05 | 1.3E-01 | 9.9E-01 | 8 |
| NA | -1.7E+00 | 3.7E-02 | -1.5E+00 | 1.6E-01 | 2.7E-01 | 9.9E-01 | 8 |
| DDIT4 | -1.3E+00 | 3.0E-01 | -2.4E+00 | 3.1E-02 | -1.2E+00 | 5.6E-01 | 8 |
| ISCA2 | -1.1E+00 | 2.9E-04 | -1.4E+00 | 2.1E-05 | -3.5E-01 | 6.8E-01 | 8 |
| CRELD1 | -3.3E+00 | 4.8E-02 | -6.1E+00 | 4.4E-03 | -2.8E+00 | 4.7E-01 | 8 |
| LYRM4 | -2.1E+00 | 2.2E-01 | -4.8E+00 | 2.1E-02 | -2.7E+00 | 4.4E-01 | 8 |
| CFL2 | -5.4E-01 | 1.2E-01 | -8.2E-01 | 2.2E-02 | -2.8E-01 | 7.6E-01 | 8 |
| NA | -1.5E+00 | 3.0E-07 | -1.7E+00 | 1.2E-07 | -1.8E-01 | 9.8E-01 | 8 |
| NA | -9.3E-01 | 1.5E-02 | -9.0E-01 | 4.4E-02 | 3.3E-02 | 9.9E-01 | 8 |
| NA | -1.6E+00 | 8.2E-02 | -2.7E+00 | 1.7E-02 | -1.1E+00 | 6.6E-01 | 8 |
| NA | -4.1E+00 | 1.3E-02 | -4.2E+00 | 3.3E-02 | -1.8E-01 | 9.9E-01 | 8 |
| NA | -3.7E+00 | 2.7E-02 | -3.7E+00 | 5.0E-02 | -9.0E-02 | 9.9E-01 | 8 |
| CYB5A | -2.1E+00 | 2.2E-03 | -1.6E+00 | 5.3E-02 | 4.9E-01 | 9.1E-01 | 1 |
| LMAN1 | -1.2E+00 | 2.7E-05 | -1.1E+00 | 1.5E-03 | 1.8E-01 | 9.7E-01 | 1 |
| NA | -1.0E+00 | 1.5E-06 | -3.4E-01 | 2.9E-01 | 6.6E-01 | 6.9E-03 | 1 |
| ATP8B1 | -6.8E-01 | 2.1E-02 | -3.0E-01 | 5.9E-01 | 3.8E-01 | 4.7E-01 | 1 |
| TMEM194A | -2.2E+00 | 1.7E-03 | -1.6E+00 | 5.6E-02 | 5.5E-01 | 8.5E-01 | 1 |
| MEX3C | -1.3E+00 | 5.9E-03 | -8.8E-01 | 1.4E-01 | 4.0E-01 | 8.0E-01 | 1 |
| TRIM36 | -9.1E-01 | 4.1E-02 | -4.0E-01 | 6.6E-01 | 5.1E-01 | 5.5E-01 | 1 |
| MTFR2 | -4.1E+00 | 1.6E-03 | -2.3E+00 | 7.1E-02 | 1.8E+00 | 5.4E-01 | 1 |
| PEX7 | -1.1E+00 | 7.4E-03 | -4.4E-01 | 5.8E-01 | 6.9E-01 | 3.0E-01 | 1 |
| METTL21B | -1.6E+00 | 2.4E-03 | -5.4E-01 | 6.1E-01 | 1.0E+00 | 1.5E-01 | 1 |
| LTV1 | -1.4E+00 | 8.7E-05 | -9.7E-01 | 2.7E-02 | 4.6E-01 | 5.7E-01 | 1 |
| XPOT | -1.3E+00 | 1.0E-02 | -5.4E-01 | 5.7E-01 | 7.5E-01 | 3.6E-01 | 1 |
| KATNA1 | -1.4E+00 | 1.5E-05 | -5.5E-01 | 2.7E-01 | 9.0E-01 | 3.5E-02 | 1 |
| NUP43 | -7.5E-01 | 2.8E-02 | -3.1E-01 | 6.7E-01 | 4.4E-01 | 4.6E-01 | 1 |
| MDM1 | -9.6E-01 | 1.9E-02 | -8.4E-01 | 9.7E-02 | 1.2E-01 | 9.9E-01 | 1 |
| C6orf136 | -8.4E-01 | 4.1E-02 | -5.4E-01 | 3.6E-01 | 2.9E-01 | 8.7E-01 | 1 |
| AKAP12 | -7.7E-01 | 1.0E-02 | -3.8E-01 | 4.4E-01 | 3.9E-01 | 4.8E-01 | 1 |
| RMND1 | -1.1E+00 | 9.7E-03 | -4.7E-01 | 5.6E-01 | 6.5E-01 | 3.6E-01 | 1 |
| CCT2 | -8.0E-01 | 3.9E-02 | -6.7E-01 | 1.8E-01 | 1.3E-01 | 9.9E-01 | 1 |
| TSPAN8 | -9.9E-01 | 5.7E-03 | -3.8E-01 | 5.8E-01 | 6.1E-01 | 2.5E-01 | 1 |
| RNF19A | -9.1E-01 | 3.5E-02 | -4.2E-01 | 6.1E-01 | 4.9E-01 | 5.6E-01 | 1 |
| ZNF706 | -2.7E+00 | 7.4E-04 | -1.1E+00 | 3.4E-01 | 1.6E+00 | 1.7E-01 | 1 |
| 3-Mar | -1.2E+00 | 1.9E-06 | -9.5E-01 | 4.8E-04 | 2.1E-01 | 8.2E-01 | 1 |
| CSNK2B | -1.0E+00 | 1.8E-03 | -3.4E-01 | 5.9E-01 | 6.6E-01 | 1.3E-01 | 1 |
| EIF3E | -1.0E+00 | 7.3E-03 | -4.1E-01 | 5.8E-01 | 6.2E-01 | 3.0E-01 | 1 |

| | | | | | | | |
|---|---|---|---|---|---|---|---|
| TMEM74 | -1.8E+00 | 1.8E-02 | -8.5E-01 | 5.0E-01 | 1.0E+00 | 5.2E-01 | 1 |
| GTSE1 | -2.1E+00 | 4.6E-02 | -1.1E+00 | 5.4E-01 | 1.0E+00 | 7.0E-01 | 1 |
| MRPL13 | -1.4E+00 | 1.8E-03 | -8.9E-01 | 1.4E-01 | 5.4E-01 | 5.9E-01 | 1 |
| SKP1 | -1.3E+00 | 6.4E-03 | -8.6E-01 | 1.5E-01 | 3.9E-01 | 8.4E-01 | 1 |
| TATDN1 | -1.2E+00 | 6.1E-03 | -6.2E-01 | 3.3E-01 | 5.5E-01 | 5.0E-01 | 1 |
| POLR3H | -1.3E+00 | 8.2E-04 | -1.1E+00 | 1.2E-02 | 1.7E-01 | 9.9E-01 | 1 |
| KIAA0196 | -1.1E+00 | 8.2E-03 | -6.6E-01 | 2.5E-01 | 4.3E-01 | 6.9E-01 | 1 |
| NA | -1.3E+00 | 3.4E-06 | -8.3E-01 | 9.7E-03 | 4.4E-01 | 3.5E-01 | 1 |
| KIF20A | -2.5E+00 | 4.2E-02 | -1.9E+00 | 2.3E-01 | 5.6E-01 | 9.9E-01 | 1 |
| AKR1B1 | -4.4E+00 | 2.0E-02 | -2.4E+00 | 3.2E-01 | 2.0E+00 | 7.2E-01 | 1 |
| HSPA9 | -1.1E+00 | 5.9E-03 | -4.0E-01 | 6.0E-01 | 6.6E-01 | 2.5E-01 | 1 |
| SYNGR1 | -1.6E+00 | 1.0E-03 | -1.4E+00 | 1.6E-02 | 2.4E-01 | 9.9E-01 | 1 |
| RPL3 | -2.0E+00 | 4.7E-08 | -1.3E+00 | 9.4E-04 | 6.1E-01 | 3.2E-01 | 1 |
| KDELR3 | -1.4E+00 | 5.4E-03 | -7.6E-01 | 2.9E-01 | 6.2E-01 | 5.5E-01 | 1 |
| MKLN1 | -9.7E-01 | 1.4E-02 | -7.0E-01 | 1.8E-01 | 2.7E-01 | 9.1E-01 | 1 |
| MTCH1 | -1.9E+00 | 1.7E-03 | -1.6E+00 | 2.4E-02 | 3.0E-01 | 9.9E-01 | 1 |
| PIM1 | -1.3E+00 | 4.0E-02 | -9.3E-01 | 3.0E-01 | 3.9E-01 | 9.3E-01 | 1 |
| TMEM209 | -9.4E-01 | 4.6E-02 | -5.8E-01 | 4.2E-01 | 3.6E-01 | 8.4E-01 | 1 |
| ATP6V1F | -9.6E-01 | 2.2E-03 | -7.4E-01 | 5.3E-02 | 2.2E-01 | 9.1E-01 | 1 |
| ATP6V1F | -9.6E-01 | 2.2E-03 | -7.4E-01 | 5.3E-02 | 2.2E-01 | 9.1E-01 | 1 |
| CALU | -6.2E-01 | 1.6E-02 | -2.4E-01 | 6.4E-01 | 3.7E-01 | 3.7E-01 | 1 |
| PLAA | -1.1E+00 | 2.2E-02 | -3.9E-01 | 7.4E-01 | 7.2E-01 | 3.6E-01 | 1 |
| TMEM33 | -5.1E+00 | 6.7E-03 | -1.4E+00 | 6.7E-01 | 3.6E+00 | 1.9E-01 | 1 |
| CROT | -9.2E-01 | 3.4E-03 | -7.7E-01 | 3.9E-02 | 1.4E-01 | 9.9E-01 | 1 |
| COMMD8 | -2.1E+00 | 8.3E-06 | -1.6E+00 | 3.1E-03 | 5.1E-01 | 6.9E-01 | 1 |
| MTERF | -1.8E+00 | 2.5E-04 | -1.5E+00 | 7.1E-03 | 2.7E-01 | 9.9E-01 | 1 |
| SLC41A2 | -1.5E+00 | 1.2E-02 | -8.1E-01 | 3.7E-01 | 7.1E-01 | 5.9E-01 | 1 |
| PEX1 | -1.5E+00 | 1.7E-03 | -1.0E+00 | 9.0E-02 | 4.8E-01 | 7.4E-01 | 1 |
| TFPI2 | -1.2E+00 | 9.1E-04 | -6.5E-01 | 1.7E-01 | 5.0E-01 | 4.1E-01 | 1 |
| TMEM165 | -1.7E+00 | 8.7E-05 | -1.3E+00 | 1.3E-02 | 4.5E-01 | 7.1E-01 | 1 |
| NCK2 | -1.4E+00 | 2.0E-03 | -1.0E+00 | 8.9E-02 | 4.4E-01 | 7.5E-01 | 1 |
| ERMP1 | -1.2E+00 | 1.4E-02 | -6.7E-01 | 3.8E-01 | 5.6E-01 | 6.1E-01 | 1 |
| NFIL3 | -1.8E+00 | 1.6E-03 | -1.2E+00 | 8.7E-02 | 5.7E-01 | 7.2E-01 | 1 |
| PHF2 | -1.5E+00 | 1.9E-04 | -7.9E-01 | 1.5E-01 | 7.2E-01 | 2.5E-01 | 1 |
| IGFBP7 | -9.3E-01 | 3.0E-04 | -3.1E-01 | 5.1E-01 | 6.1E-01 | 6.3E-02 | 1 |
| POLR1E | -1.2E+00 | 1.3E-02 | -6.7E-01 | 3.3E-01 | 4.9E-01 | 6.8E-01 | 1 |
| ACTR1B | -2.8E+00 | 2.3E-03 | -2.1E+00 | 6.5E-02 | 7.7E-01 | 8.7E-01 | 1 |
| AHR | -1.5E+00 | 8.2E-03 | -8.3E-01 | 3.3E-01 | 6.8E-01 | 5.5E-01 | 1 |
| LRPPRC | -7.6E-01 | 1.2E-03 | -4.4E-01 | 1.6E-01 | 3.2E-01 | 4.7E-01 | 1 |
| TGFBR1 | -9.1E-01 | 1.5E-02 | -4.5E-01 | 4.8E-01 | 4.6E-01 | 5.2E-01 | 1 |
| SEC61B | -1.2E+00 | 1.4E-03 | -4.6E-01 | 5.1E-01 | 7.7E-01 | 1.5E-01 | 1 |
| FOXJ3 | -7.7E-01 | 2.1E-02 | -2.5E-01 | 7.8E-01 | 5.2E-01 | 3.1E-01 | 1 |
| CRIPT | -9.9E-01 | 5.5E-04 | -8.0E-01 | 1.8E-02 | 1.9E-01 | 9.5E-01 | 1 |
| EPCAM | -1.0E+00 | 2.7E-05 | -4.0E-01 | 2.6E-01 | 6.0E-01 | 4.9E-02 | 1 |
| CD109 | -8.6E-01 | 1.5E-02 | -6.3E-01 | 1.7E-01 | 2.2E-01 | 9.3E-01 | 1 |
| NA | -2.3E+00 | 3.2E-04 | -1.8E+00 | 1.7E-02 | 5.1E-01 | 8.7E-01 | 1 |
| NA | -2.1E+00 | 6.4E-03 | -1.5E+00 | 1.4E-01 | 6.3E-01 | 8.2E-01 | 1 |
| MYEOV2 | -1.2E+00 | 2.4E-02 | -1.1E+00 | 8.5E-02 | 1.1E-01 | 9.9E-01 | 1 |
| FSD1L | -1.6E+00 | 1.0E-02 | -1.2E+00 | 1.4E-01 | 4.2E-01 | 9.3E-01 | 1 |
| NA | -2.0E+00 | 6.9E-04 | -1.4E+00 | 6.9E-02 | 6.7E-01 | 6.4E-01 | 1 |
| CCDC104 | -1.8E+00 | 2.5E-02 | -7.4E-01 | 6.4E-01 | 1.1E+00 | 4.6E-01 | 1 |
| SKAP2 | -6.0E-01 | 7.2E-03 | -3.8E-01 | 2.2E-01 | 2.3E-01 | 6.9E-01 | 1 |
| SUSD1 | -1.5E+00 | 2.2E-02 | -1.2E+00 | 1.5E-01 | 2.9E-01 | 9.9E-01 | 1 |
| NA | -2.3E+00 | 3.0E-02 | -1.3E+00 | 3.9E-01 | 1.0E+00 | 7.5E-01 | 1 |
| MDH1 | -1.1E+00 | 3.6E-02 | -4.2E-01 | 7.5E-01 | 7.2E-01 | 4.3E-01 | 1 |
| UGP2 | -6.0E-01 | 2.2E-02 | -4.0E-01 | 2.8E-01 | 2.0E-01 | 8.4E-01 | 1 |
| NA | -1.8E+00 | 7.2E-04 | -7.5E-01 | 3.4E-01 | 1.1E+00 | 1.9E-01 | 1 |
| RRAGC | -9.0E-01 | 1.7E-02 | -6.2E-01 | 2.2E-01 | 2.8E-01 | 8.7E-01 | 1 |
| UTP11L | -1.3E+00 | 1.9E-03 | -1.1E+00 | 2.9E-02 | 2.3E-01 | 9.9E-01 | 1 |
| ELMO1 | -2.1E+00 | 2.7E-02 | -1.5E+00 | 2.2E-01 | 5.7E-01 | 9.6E-01 | 1 |
| ITGB1BP1 | -1.1E+00 | 2.4E-02 | -4.6E-01 | 6.2E-01 | 6.2E-01 | 4.6E-01 | 1 |
| ST7 | -6.2E-01 | 3.6E-02 | -4.8E-01 | 2.2E-01 | 1.4E-01 | 9.9E-01 | 1 |
| ST7 | -6.2E-01 | 3.6E-02 | -4.8E-01 | 2.2E-01 | 1.4E-01 | 9.9E-01 | 1 |
| ST7 | -6.2E-01 | 3.6E-02 | -4.8E-01 | 2.2E-01 | 1.4E-01 | 9.9E-01 | 1 |
| EZH2 | -6.1E-01 | 4.5E-02 | -5.0E-01 | 2.1E-01 | 1.2E-01 | 9.9E-01 | 1 |
| GAB1 | -1.2E+00 | 1.9E-03 | -6.7E-01 | 1.9E-01 | 5.0E-01 | 4.9E-01 | 1 |
| ATG5 | -1.0E+00 | 5.5E-04 | -6.5E-01 | 9.4E-02 | 3.9E-01 | 5.2E-01 | 1 |
| MGST2 | -8.0E-01 | 3.8E-03 | -3.2E-01 | 5.4E-01 | 4.9E-01 | 2.4E-01 | 1 |
| PPIL6 | -9.6E-01 | 3.7E-03 | -4.6E-01 | 3.7E-01 | 5.0E-01 | 3.8E-01 | 1 |
| PLK4 | -1.8E+00 | 2.6E-03 | -1.4E+00 | 4.6E-02 | 3.5E-01 | 9.8E-01 | 1 |
| MPP7 | -1.4E+00 | 8.1E-04 | -7.4E-01 | 2.0E-01 | 6.6E-01 | 3.5E-01 | 1 |
| SLC37A3 | -1.3E+00 | 1.1E-03 | -1.1E+00 | 2.4E-02 | 2.5E-01 | 9.6E-01 | 1 |
| ZEB1 | -1.9E+00 | 1.8E-02 | -1.4E+00 | 2.0E-01 | 5.4E-01 | 9.3E-01 | 1 |
| ZEB1 | -1.9E+00 | 1.8E-02 | -1.4E+00 | 2.0E-01 | 5.4E-01 | 9.3E-01 | 1 |
| EFCAB10 | -1.1E+00 | 3.5E-02 | -4.0E-01 | 7.7E-01 | 7.3E-01 | 4.0E-01 | 1 |
| PHF10 | -1.1E+00 | 2.7E-04 | -6.1E-01 | 1.5E-01 | 5.3E-01 | 2.9E-01 | 1 |
| TTC26 | -1.9E+00 | 3.1E-04 | -1.4E+00 | 2.7E-02 | 5.1E-01 | 7.7E-01 | 1 |
| MLLT10 | -9.5E-01 | 1.5E-03 | -8.0E-01 | 2.4E-02 | 1.5E-01 | 9.9E-01 | 1 |
| ORC5 | -9.8E-01 | 2.0E-02 | -4.8E-01 | 5.1E-01 | 5.0E-01 | 5.5E-01 | 1 |
| RPN1 | -1.5E+00 | 4.7E-02 | -6.6E-01 | 6.8E-01 | 8.7E-01 | 5.5E-01 | 1 |
| HMCES | -8.6E-01 | 1.4E-04 | -5.6E-01 | 4.3E-02 | 3.0E-01 | 5.1E-01 | 1 |
| PMPCB | -6.9E-01 | 2.0E-02 | -5.4E-01 | 1.6E-01 | 1.5E-01 | 9.9E-01 | 1 |
| ATP6V0A4 | -2.4E+00 | 4.6E-05 | -9.3E-01 | 2.7E-01 | 1.4E+00 | 6.3E-02 | 1 |
| UQCRH | -1.3E+00 | 3.0E-06 | -7.0E-01 | 4.1E-02 | 5.9E-01 | 1.3E-01 | 1 |
| UQCRH | -1.3E+00 | 3.0E-06 | -7.0E-01 | 4.1E-02 | 5.9E-01 | 1.3E-01 | 1 |
| TRIM13 | -6.9E-01 | 1.9E-02 | -5.7E-01 | 1.2E-01 | 1.2E-01 | 9.9E-01 | 1 |
| SVOPL | -1.3E+00 | 3.7E-02 | -8.1E-01 | 3.6E-01 | 4.6E-01 | 8.5E-01 | 1 |
| EBPL | -2.2E+00 | 2.7E-03 | -1.5E+00 | 9.6E-02 | 6.7E-01 | 8.0E-01 | 1 |
| NA | -3.1E+00 | 1.4E-04 | -2.2E+00 | 2.1E-02 | 9.0E-01 | 6.8E-01 | 1 |
| YME1L1 | -9.3E-01 | 4.2E-02 | -4.3E-01 | 6.2E-01 | 5.0E-01 | 5.9E-01 | 1 |
| NA | -6.6E-01 | 2.8E-02 | -4.6E-01 | 2.6E-01 | 2.0E-01 | 9.1E-01 | 1 |
| ITM2B | -2.0E+00 | 1.5E-04 | -1.4E+00 | 2.8E-02 | 6.0E-01 | 6.3E-01 | 1 |
| ABHD5 | -9.1E-01 | 1.3E-02 | -5.5E-01 | 3.0E-01 | 3.6E-01 | 7.0E-01 | 1 |
| CYP27C1 | -1.3E+00 | 1.9E-02 | -1.1E+00 | 1.2E-01 | 2.2E-01 | 9.9E-01 | 1 |

| | | | | | | | |
|---|---|---|---|---|---|---|---|
| GEMIN7 | -4.1E+00 | 4.3E-03 | -3.2E+00 | 4.6E-02 | 9.3E-01 | 9.8E-01 | 1 |
| RSU1 | -1.5E+00 | 1.2E-05 | -9.1E-01 | 3.0E-02 | 6.0E-01 | 2.8E-01 | 1 |
| CCDC90B | -1.8E+00 | 1.9E-05 | -1.5E+00 | 1.4E-03 | 2.9E-01 | 9.4E-01 | 1 |
| HPDL | -2.6E+00 | 5.2E-03 | -2.2E+00 | 4.6E-02 | 3.8E-01 | 9.9E-01 | 1 |
| DNAJC15 | -1.6E+00 | 4.2E-02 | -9.5E-01 | 4.1E-01 | 6.1E-01 | 8.3E-01 | 1 |
| PDCD6IP | -9.3E-01 | 2.9E-03 | -5.2E-01 | 2.4E-01 | 4.1E-01 | 4.9E-01 | 1 |
| CAMK1D | -1.5E+00 | 3.2E-02 | -1.2E+00 | 1.7E-01 | 2.7E-01 | 9.9E-01 | 1 |
| ATRAID | -1.4E+00 | 1.1E-04 | -1.0E+00 | 2.1E-02 | 4.1E-01 | 6.7E-01 | 1 |
| TMEM185B | -1.9E+00 | 3.5E-04 | -1.4E+00 | 2.5E-02 | 4.8E-01 | 8.0E-01 | 1 |
| RHEB | -8.7E-01 | 5.3E-03 | -4.1E-01 | 4.1E-01 | 4.5E-01 | 3.9E-01 | 1 |
| PLCD1 | -8.4E-01 | 3.6E-02 | -4.9E-01 | 4.4E-01 | 3.5E-01 | 7.4E-01 | 1 |
| MYD88 | -1.0E+00 | 4.7E-02 | -8.8E-01 | 1.8E-01 | 1.5E-01 | 9.9E-01 | 1 |
| SLC35F5 | -1.1E+00 | 4.8E-02 | -8.1E-01 | 3.2E-01 | 3.4E-01 | 9.5E-01 | 1 |
| NA | -2.5E+00 | 3.0E-02 | -1.4E+00 | 4.2E-01 | 1.1E+00 | 7.3E-01 | 1 |
| TBC1D4 | -1.0E+00 | 6.1E-04 | -3.5E-01 | 5.1E-01 | 6.9E-01 | 9.9E-02 | 1 |
| NA | -7.0E-01 | 3.1E-02 | -4.3E-01 | 3.9E-01 | 2.8E-01 | 7.9E-01 | 1 |
| JMJD6 | -2.5E+00 | 6.4E-04 | -1.7E+00 | 4.5E-02 | 8.1E-01 | 7.3E-01 | 1 |
| FAM208B | -1.1E+00 | 1.4E-02 | -8.3E-01 | 1.3E-01 | 2.3E-01 | 9.8E-01 | 1 |
| NDFIP2 | -1.3E+00 | 2.4E-03 | -8.0E-01 | 1.4E-01 | 4.6E-01 | 6.3E-01 | 1 |
| C11orf30 | -1.3E+00 | 1.0E-02 | -1.1E+00 | 6.3E-02 | 1.6E-01 | 9.9E-01 | 1 |
| INTS10 | -1.4E+00 | 4.8E-04 | -6.2E-01 | 3.1E-01 | 7.9E-01 | 1.8E-01 | 1 |
| ABCC4 | -5.9E-01 | 1.4E-02 | -4.2E-01 | 1.9E-01 | 1.7E-01 | 9.0E-01 | 1 |
| DNAJC3 | -9.6E-01 | 1.4E-02 | -5.3E-01 | 3.7E-01 | 4.3E-01 | 6.1E-01 | 1 |
| ORC4 | -1.6E+00 | 2.6E-02 | -6.7E-01 | 6.4E-01 | 9.5E-01 | 4.7E-01 | 1 |
| MBNL2 | -6.0E-01 | 4.4E-02 | -2.3E-01 | 7.5E-01 | 3.7E-01 | 4.8E-01 | 1 |
| EPC2 | -1.1E+00 | 2.7E-04 | -6.5E-01 | 9.0E-02 | 4.3E-01 | 4.2E-01 | 1 |
| PAAF1 | -2.3E+00 | 3.9E-02 | -1.2E+00 | 4.6E-01 | 1.0E+00 | 7.4E-01 | 1 |
| DOCK9 | -1.4E+00 | 5.5E-03 | -9.8E-01 | 1.3E-01 | 4.1E-01 | 8.2E-01 | 1 |
| ACTG1 | -1.0E+00 | 6.0E-03 | -6.4E-01 | 2.1E-01 | 3.9E-01 | 6.8E-01 | 1 |
| ACTG1 | -1.0E+00 | 6.0E-03 | -6.4E-01 | 2.1E-01 | 3.9E-01 | 6.8E-01 | 1 |
| NR1D2 | -1.1E+00 | 1.3E-02 | -6.9E-01 | 2.9E-01 | 4.4E-01 | 7.1E-01 | 1 |
| YIPF4 | -2.4E+00 | 9.1E-04 | -2.1E+00 | 1.2E-02 | 2.9E-01 | 9.9E-01 | 1 |
| RNF121 | -2.0E+00 | 1.0E-02 | -1.7E+00 | 6.3E-02 | 2.3E-01 | 9.9E-01 | 1 |
| P4HB | -9.2E-01 | 4.7E-04 | -8.0E-01 | 8.2E-03 | 1.2E-01 | 9.9E-01 | 1 |
| NDUFA2 | -1.8E+00 | 1.1E-04 | -1.4E+00 | 1.1E-02 | 4.2E-01 | 7.9E-01 | 1 |
| TMTC2 | -1.7E+00 | 1.3E-02 | -7.6E-01 | 5.4E-01 | 9.6E-01 | 4.3E-01 | 1 |
| CRLS1 | -2.2E+00 | 4.7E-03 | -1.9E+00 | 4.2E-02 | 3.0E-01 | 9.9E-01 | 1 |
| SPINT2 | -1.0E+00 | 2.6E-02 | -6.2E-01 | 3.5E-01 | 3.8E-01 | 7.7E-01 | 1 |
| MRPS31 | -1.8E+00 | 7.9E-04 | -1.4E+00 | 2.1E-02 | 3.5E-01 | 9.4E-01 | 1 |
| CSNK1D | -3.9E+00 | 4.8E-04 | -2.9E+00 | 1.9E-02 | 9.4E-01 | 8.7E-01 | 1 |
| PGAM5 | -6.4E-01 | 6.5E-03 | -2.6E-01 | 5.5E-01 | 3.8E-01 | 3.0E-01 | 1 |
| EIF4EBP1 | -3.3E+00 | 4.5E-05 | -2.6E+00 | 5.2E-03 | 7.6E-01 | 8.3E-01 | 1 |
| RMDN2 | -1.6E+00 | 4.7E-02 | -1.1E+00 | 3.4E-01 | 4.9E-01 | 9.3E-01 | 1 |
| ATP11A | -1.6E+00 | 7.2E-06 | -1.3E+00 | 6.9E-04 | 2.4E-01 | 9.2E-01 | 1 |
| NDUFA12 | -7.4E-01 | 2.0E-02 | -4.4E-01 | 3.6E-01 | 3.0E-01 | 7.1E-01 | 1 |
| DNAJC13 | -8.2E-01 | 1.7E-02 | -6.8E-01 | 1.1E-01 | 1.4E-01 | 9.9E-01 | 1 |
| PCID2 | -1.1E+00 | 8.4E-04 | -4.6E-01 | 3.9E-01 | 6.4E-01 | 1.7E-01 | 1 |
| TTI2 | -1.1E+00 | 2.8E-02 | -7.0E-01 | 3.4E-01 | 4.1E-01 | 8.3E-01 | 1 |
| NA | -9.0E-01 | 3.0E-04 | -5.4E-01 | 8.8E-02 | 3.5E-01 | 4.6E-01 | 1 |
| NA | -9.0E-01 | 3.0E-04 | -5.4E-01 | 8.8E-02 | 3.5E-01 | 4.6E-01 | 1 |
| MRPS26 | -2.1E+00 | 4.0E-04 | -1.5E+00 | 3.0E-02 | 5.5E-01 | 7.8E-01 | 1 |
| TOPBP1 | -6.1E-01 | 5.5E-03 | -2.3E-01 | 5.8E-01 | 3.8E-01 | 2.6E-01 | 1 |
| ELK3 | -7.5E-01 | 3.0E-04 | -5.4E-01 | 3.1E-02 | 2.1E-01 | 7.1E-01 | 1 |
| PTPRA | -2.3E+00 | 3.2E-03 | -1.6E+00 | 9.3E-02 | 6.5E-01 | 8.2E-01 | 1 |
| DCTN6 | -2.0E+00 | 2.5E-02 | -1.4E+00 | 2.5E-01 | 6.1E-01 | 9.1E-01 | 1 |
| HSPH1 | -1.5E+00 | 1.4E-04 | -1.2E+00 | 6.0E-03 | 2.5E-01 | 9.5E-01 | 1 |
| MCM4 | -1.5E+00 | 2.1E-03 | -7.1E-01 | 3.7E-01 | 8.3E-01 | 2.9E-01 | 1 |
| NA | -3.5E+00 | 3.9E-02 | -1.9E+00 | 3.4E-01 | 1.5E+00 | 8.0E-01 | 1 |
| PCMTD1 | -1.0E+00 | 5.6E-03 | -7.8E-01 | 9.5E-02 | 2.5E-01 | 9.1E-01 | 1 |
| POLR1D | -1.4E+00 | 1.3E-02 | -5.9E-01 | 5.9E-01 | 8.3E-01 | 3.8E-01 | 1 |
| NSFL1C | -2.1E+00 | 5.8E-05 | -1.8E+00 | 2.6E-03 | 3.0E-01 | 9.9E-01 | 1 |
| NA | -8.7E-01 | 3.0E-02 | -6.4E-01 | 2.3E-01 | 2.2E-01 | 9.7E-01 | 1 |
| NR1H4 | -2.7E+00 | 3.4E-02 | -1.5E+00 | 3.9E-01 | 1.2E+00 | 7.5E-01 | 1 |
| PPAP2A | -2.0E+00 | 4.5E-07 | -1.7E+00 | 8.8E-05 | 2.9E-01 | 9.2E-01 | 1 |
| PSMD6 | -1.9E+00 | 5.8E-06 | -1.1E+00 | 4.4E-02 | 8.7E-01 | 1.6E-01 | 1 |
| PDHX | -2.4E+00 | 3.3E-06 | -1.4E+00 | 1.5E-02 | 9.8E-01 | 2.7E-01 | 1 |
| APIP | -1.3E+00 | 7.9E-04 | -5.7E-01 | 3.2E-01 | 7.0E-01 | 2.2E-01 | 1 |
| MAP3K1 | -1.4E+00 | 5.2E-03 | -1.0E+00 | 1.2E-01 | 4.0E-01 | 8.7E-01 | 1 |
| CAT | -1.8E+00 | 4.5E-05 | -1.3E+00 | 1.1E-02 | 4.7E-01 | 6.6E-01 | 1 |
| RGS20 | -1.2E+00 | 1.3E-02 | -7.0E-01 | 3.2E-01 | 5.1E-01 | 6.8E-01 | 1 |
| CSGALNACT2 | -9.9E-01 | 4.4E-02 | -3.1E-01 | 8.5E-01 | 6.8E-01 | 3.9E-01 | 1 |
| IMPAD1 | -2.9E+00 | 1.5E-03 | -2.3E+00 | 3.3E-02 | 6.3E-01 | 9.6E-01 | 1 |
| BMS1 | -6.8E-01 | 2.3E-03 | -5.5E-01 | 3.9E-02 | 1.3E-01 | 9.8E-01 | 1 |
| BMS1 | -6.8E-01 | 2.3E-03 | -5.5E-01 | 3.9E-02 | 1.3E-01 | 9.8E-01 | 1 |
| MIPEP | -1.6E+00 | 8.0E-03 | -7.4E-01 | 4.5E-01 | 8.5E-01 | 4.2E-01 | 1 |
| ELOVL7 | -6.8E-01 | 2.6E-02 | -2.8E-01 | 6.6E-01 | 4.1E-01 | 4.4E-01 | 1 |
| GNPTAB | -8.5E-01 | 2.9E-02 | -7.1E-01 | 1.5E-01 | 1.4E-01 | 9.9E-01 | 1 |
| KRT5 | -1.2E+00 | 1.4E-02 | -8.2E-01 | 2.1E-01 | 3.6E-01 | 8.8E-01 | 1 |
| IPO7 | -1.3E+00 | 1.9E-06 | -4.9E-01 | 2.2E-01 | 8.2E-01 | 1.4E-02 | 1 |
| GPI | -1.6E+00 | 3.9E-04 | -1.4E+00 | 9.4E-03 | 2.3E-01 | 9.9E-01 | 1 |
| COPS5 | -9.0E-01 | 1.6E-02 | -3.5E-01 | 6.4E-01 | 5.5E-01 | 3.7E-01 | 1 |
| TRAPPC13 | -2.1E+00 | 2.5E-03 | -1.6E+00 | 5.6E-02 | 4.8E-01 | 9.1E-01 | 1 |
| PAM | -6.0E-01 | 4.9E-02 | -2.6E-01 | 7.0E-01 | 3.5E-01 | 5.5E-01 | 1 |
| VTI1B | -2.4E+00 | 7.4E-03 | -1.7E+00 | 1.5E-01 | 7.7E-01 | 8.3E-01 | 1 |
| VAMP8 | -1.3E+00 | 1.3E-03 | -1.1E+00 | 2.1E-02 | 2.0E-01 | 9.9E-01 | 1 |
| CDK7 | -2.1E+00 | 1.2E-04 | -1.8E+00 | 4.2E-03 | 3.1E-01 | 9.9E-01 | 1 |
| NA | -1.4E+00 | 3.7E-04 | -8.6E-01 | 9.6E-02 | 5.7E-01 | 4.5E-01 | 1 |
| NA | -1.4E+00 | 3.7E-04 | -8.6E-01 | 9.6E-02 | 5.7E-01 | 4.5E-01 | 1 |
| NA | -1.4E+00 | 3.7E-04 | -8.6E-01 | 9.6E-02 | 5.7E-01 | 4.5E-01 | 1 |
| NA | -1.4E+00 | 3.7E-04 | -8.6E-01 | 9.6E-02 | 5.7E-01 | 4.5E-01 | 1 |
| SMN1 | -1.9E+00 | 7.1E-04 | -8.0E-01 | 3.7E-01 | 1.1E+00 | 1.7E-01 | 1 |
| SMN1 | -1.9E+00 | 7.1E-04 | -8.0E-01 | 3.7E-01 | 1.1E+00 | 1.7E-01 | 1 |
| SMN1 | -1.9E+00 | 7.1E-04 | -8.0E-01 | 3.7E-01 | 1.1E+00 | 1.7E-01 | 1 |
| SMN1 | -1.9E+00 | 7.1E-04 | -8.0E-01 | 3.7E-01 | 1.1E+00 | 1.7E-01 | 1 |

| | | | | | | | |
|---|---|---|---|---|---|---|---|
| TRAPPC11 | -1.1E+00 | 2.6E-05 | -4.3E-01 | 2.9E-01 | 6.9E-01 | 4.1E-02 | 1 |
| TFCP2 | -5.9E-01 | 3.7E-02 | -2.2E-01 | 7.5E-01 | 3.7E-01 | 4.3E-01 | 1 |
| TRAM1 | -9.3E-01 | 2.3E-02 | -4.5E-01 | 5.4E-01 | 4.9E-01 | 5.3E-01 | 1 |
| TMSB10 | -1.2E+00 | 5.3E-03 | -6.7E-01 | 2.6E-01 | 5.0E-01 | 5.6E-01 | 1 |
| NA | -6.9E-01 | 1.3E-02 | -4.9E-01 | 1.9E-01 | 2.0E-01 | 8.9E-01 | 1 |
| EMC4 | -2.4E+00 | 2.3E-06 | -1.8E+00 | 1.6E-03 | 6.0E-01 | 6.5E-01 | 1 |
| EIF6 | -1.5E+00 | 9.1E-05 | -1.1E+00 | 1.3E-02 | 3.9E-01 | 7.3E-01 | 1 |
| NA | -1.5E+00 | 5.5E-04 | -7.8E-01 | 2.1E-01 | 7.5E-01 | 2.9E-01 | 1 |
| TMEM70 | -8.2E-01 | 7.2E-03 | -3.0E-01 | 6.4E-01 | 5.2E-01 | 2.5E-01 | 1 |
| MRPL19 | -1.3E+00 | 2.7E-04 | -1.1E+00 | 1.3E-02 | 2.6E-01 | 9.1E-01 | 1 |
| ARFIP1 | -6.9E-01 | 8.6E-03 | -5.7E-01 | 7.5E-02 | 1.2E-01 | 9.9E-01 | 1 |
| ANKRA2 | -1.5E+00 | 5.0E-03 | -7.4E-01 | 3.7E-01 | 7.6E-01 | 4.2E-01 | 1 |
| LIMA1 | -1.7E+00 | 5.5E-05 | -9.8E-01 | 6.3E-02 | 7.0E-01 | 3.0E-01 | 1 |
| ACTR8 | -1.9E+00 | 2.6E-03 | -9.7E-01 | 3.0E-01 | 9.2E-01 | 3.9E-01 | 1 |
| HEXB | -1.0E+00 | 8.2E-03 | -3.5E-01 | 6.8E-01 | 6.8E-01 | 2.3E-01 | 1 |
| NA | -2.3E+00 | 4.3E-05 | -1.5E+00 | 3.3E-02 | 8.7E-01 | 4.1E-01 | 1 |
| PFN2 | -8.5E-01 | 3.1E-03 | -3.8E-01 | 4.2E-01 | 4.7E-01 | 3.0E-01 | 1 |
| SETD6 | -1.2E+00 | 1.9E-04 | -7.3E-01 | 7.1E-02 | 4.7E-01 | 4.5E-01 | 1 |
| ZFAND1 | -1.9E+00 | 4.2E-02 | -9.7E-01 | 5.3E-01 | 9.1E-01 | 6.9E-01 | 1 |
| C16orf80 | -1.4E+00 | 3.2E-02 | -8.4E-01 | 4.2E-01 | 6.0E-01 | 7.4E-01 | 1 |
| BRAP | -1.1E+00 | 1.2E-02 | -6.9E-01 | 2.3E-01 | 3.7E-01 | 7.9E-01 | 1 |
| E2F5 | -1.1E+00 | 3.2E-02 | -4.0E-01 | 7.4E-01 | 6.8E-01 | 4.2E-01 | 1 |
| ETFDH | -1.5E+00 | 1.8E-03 | -7.4E-01 | 3.0E-01 | 7.5E-01 | 3.4E-01 | 1 |
| TM9SF3 | -1.0E+00 | 2.0E-04 | -7.5E-01 | 2.6E-02 | 3.0E-01 | 6.9E-01 | 1 |
| ANXA3 | -1.5E+00 | 1.8E-05 | -1.0E+00 | 1.7E-02 | 5.2E-01 | 4.4E-01 | 1 |
| URB1 | -1.6E+00 | 2.8E-02 | -1.3E+00 | 1.6E-01 | 2.9E-01 | 9.9E-01 | 1 |
| NA | -1.6E+00 | 1.6E-08 | -5.2E-01 | 1.7E-01 | 1.0E+00 | 1.2E-03 | 1 |
| PAPD4 | -1.1E+00 | 1.6E-03 | -4.5E-01 | 4.2E-01 | 6.1E-01 | 2.2E-01 | 1 |
| ZNF638 | -1.1E+00 | 4.6E-03 | -7.8E-01 | 1.2E-01 | 3.2E-01 | 8.2E-01 | 1 |
| ZNF638 | -1.1E+00 | 4.6E-03 | -7.8E-01 | 1.2E-01 | 3.2E-01 | 8.2E-01 | 1 |
| MMS19 | -2.7E+00 | 2.2E-04 | -1.9E+00 | 2.3E-02 | 8.0E-01 | 7.3E-01 | 1 |
| RPUSD2 | -1.4E+00 | 5.6E-03 | -7.3E-01 | 3.1E-01 | 6.5E-01 | 5.2E-01 | 1 |
| CMSS1 | -2.0E+00 | 1.8E-06 | -1.5E+00 | 8.8E-04 | 4.3E-01 | 7.4E-01 | 1 |
| F2RL1 | -1.4E+00 | 7.8E-03 | -1.2E+00 | 5.4E-02 | 1.8E-01 | 9.9E-01 | 1 |
| NIT2 | -9.2E-01 | 9.4E-03 | -3.6E-01 | 6.1E-01 | 5.7E-01 | 3.1E-01 | 1 |
| WDR75 | -9.7E-01 | 3.6E-02 | -4.8E-01 | 5.6E-01 | 4.9E-01 | 6.2E-01 | 1 |
| ASNSD1 | -9.5E-01 | 6.5E-04 | -5.8E-01 | 1.1E-01 | 3.7E-01 | 4.9E-01 | 1 |
| ANO6 | -1.4E+00 | 1.0E-03 | -1.0E+00 | 3.9E-02 | 3.4E-01 | 8.4E-01 | 1 |
| WDSUB1 | -8.9E-01 | 1.3E-02 | -3.5E-01 | 6.2E-01 | 5.4E-01 | 3.5E-01 | 1 |
| SLC39A14 | -1.2E+00 | 1.7E-02 | -1.0E+00 | 1.1E-01 | 2.1E-01 | 9.9E-01 | 1 |
| GLS | -6.6E-01 | 1.7E-02 | -4.3E-01 | 2.6E-01 | 2.3E-01 | 8.1E-01 | 1 |
| COX15 | -7.8E-01 | 1.5E-02 | -4.0E-01 | 4.4E-01 | 3.8E-01 | 5.7E-01 | 1 |
| NUDT9 | -2.2E+00 | 1.9E-03 | -1.3E+00 | 2.0E-01 | 9.7E-01 | 4.8E-01 | 1 |
| SPP1 | -2.3E+00 | 6.1E-03 | -1.6E+00 | 1.3E-01 | 6.5E-01 | 8.4E-01 | 1 |
| PPP1R15B | -1.5E+00 | 2.0E-02 | -8.1E-01 | 4.1E-01 | 6.4E-01 | 6.6E-01 | 1 |
| NA | -2.2E+00 | 9.4E-06 | -1.8E+00 | 1.5E-03 | 4.2E-01 | 8.7E-01 | 1 |
| NA | -2.2E+00 | 9.4E-06 | -1.8E+00 | 1.5E-03 | 4.2E-01 | 8.7E-01 | 1 |
| PRMT3 | -6.2E-01 | 6.0E-03 | -4.6E-01 | 1.1E-01 | 1.6E-01 | 9.0E-01 | 1 |
| TANK | -8.4E-01 | 4.5E-02 | -5.9E-01 | 3.2E-01 | 2.5E-01 | 9.5E-01 | 1 |
| NA | -1.2E+00 | 3.6E-03 | -1.0E+00 | 4.6E-02 | 2.1E-01 | 9.9E-01 | 1 |
| MFSD4 | -2.3E+00 | 6.1E-03 | -1.4E+00 | 1.9E-01 | 8.7E-01 | 7.2E-01 | 1 |
| BACE2 | -1.6E+00 | 9.4E-05 | -1.2E+00 | 1.4E-02 | 4.2E-01 | 7.1E-01 | 1 |
| CHCHD2 | -1.5E+00 | 6.9E-05 | -6.3E-01 | 2.6E-01 | 8.7E-01 | 8.2E-02 | 1 |
| VKORC1L1 | -1.8E+00 | 2.8E-02 | -1.6E+00 | 1.2E-01 | 2.4E-01 | 9.9E-01 | 1 |
| ARL3 | -1.7E+00 | 5.8E-04 | -1.5E+00 | 7.2E-03 | 1.7E-01 | 9.9E-01 | 1 |
| CISH | -1.0E+00 | 1.3E-02 | -6.9E-01 | 2.2E-01 | 3.4E-01 | 8.3E-01 | 1 |
| ETS1 | -8.8E-01 | 1.7E-03 | -4.0E-01 | 3.7E-01 | 4.9E-01 | 2.6E-01 | 1 |
| SRSF9 | -2.2E+00 | 4.4E-04 | -1.8E+00 | 1.7E-02 | 4.4E-01 | 9.3E-01 | 1 |
| SRSF9 | -2.2E+00 | 4.4E-04 | -1.8E+00 | 1.7E-02 | 4.4E-01 | 9.3E-01 | 1 |
| GMNN | -1.6E+00 | 1.3E-02 | -7.2E-01 | 5.2E-01 | 8.6E-01 | 4.5E-01 | 1 |
| NA | -1.0E+00 | 6.8E-05 | -3.7E-01 | 3.8E-01 | 6.5E-01 | 4.4E-02 | 1 |
| GLRX2 | -1.3E+00 | 1.6E-03 | -7.5E-01 | 1.6E-01 | 5.1E-01 | 5.2E-01 | 1 |
| PPP3CA | -7.2E-01 | 4.2E-03 | -3.6E-01 | 3.5E-01 | 3.6E-01 | 4.1E-01 | 1 |
| STK17B | -1.8E+00 | 5.0E-03 | -1.3E+00 | 1.1E-01 | 4.9E-01 | 8.9E-01 | 1 |
| NFKB1 | -1.9E+00 | 3.4E-04 | -1.5E+00 | 1.6E-02 | 4.0E-01 | 8.9E-01 | 1 |
| CAB39 | -1.0E+00 | 1.5E-03 | -4.5E-01 | 3.7E-01 | 5.5E-01 | 2.5E-01 | 1 |
| GRAMD1C | -1.3E+00 | 1.6E-02 | -1.2E+00 | 6.1E-02 | 8.4E-02 | 9.9E-01 | 1 |
| TC2N | -1.5E+00 | 7.0E-03 | -7.1E-01 | 4.6E-01 | 8.2E-01 | 3.9E-01 | 1 |
| LYPLAL1 | -1.8E+00 | 4.0E-02 | -1.5E+00 | 1.8E-01 | 3.2E-01 | 9.9E-01 | 1 |
| PPA2 | -1.7E+00 | 4.4E-03 | -1.5E+00 | 3.6E-02 | 2.0E-01 | 9.9E-01 | 1 |
| NA | -2.2E+00 | 3.1E-03 | -1.8E+00 | 3.7E-02 | 3.4E-01 | 9.9E-01 | 1 |
| NA | -2.2E+00 | 3.1E-03 | -1.8E+00 | 3.7E-02 | 3.4E-01 | 9.9E-01 | 1 |
| NA | -2.2E+00 | 3.1E-03 | -1.8E+00 | 3.7E-02 | 3.4E-01 | 9.9E-01 | 1 |
| NA | -2.2E+00 | 3.1E-03 | -1.8E+00 | 3.7E-02 | 3.4E-01 | 9.9E-01 | 1 |
| TMEM39A | -2.1E+00 | 5.1E-04 | -1.5E+00 | 3.3E-02 | 5.5E-01 | 8.0E-01 | 1 |
| BPNT1 | -1.4E+00 | 8.2E-03 | -4.6E-01 | 7.1E-01 | 9.3E-01 | 2.2E-01 | 1 |
| POGLUT1 | -7.3E-01 | 1.2E-02 | -4.8E-01 | 2.2E-01 | 2.5E-01 | 7.9E-01 | 1 |
| SDHA | -2.0E+00 | 2.6E-06 | -1.1E+00 | 4.5E-02 | 9.7E-01 | 1.1E-01 | 1 |
| ZDHHC6 | -9.0E-01 | 2.0E-02 | -7.2E-01 | 1.4E-01 | 1.8E-01 | 9.9E-01 | 1 |
| PPFIBP1 | -7.5E-01 | 1.0E-03 | -6.2E-01 | 2.1E-02 | 1.3E-01 | 9.8E-01 | 1 |
| LGMN | -2.3E+00 | 6.3E-03 | -1.4E+00 | 2.0E-01 | 8.6E-01 | 7.0E-01 | 1 |
| TYW5 | -1.8E+00 | 9.5E-03 | -1.4E+00 | 9.5E-02 | 3.6E-01 | 9.9E-01 | 1 |
| NA | -1.9E+00 | 7.9E-07 | -1.1E+00 | 2.0E-02 | 8.6E-01 | 1.4E-01 | 1 |
| NA | -1.0E+00 | 2.7E-03 | -4.4E-01 | 4.6E-01 | 6.0E-01 | 2.6E-01 | 1 |
| PDCD6 | -4.8E+00 | 2.0E-03 | -2.3E+00 | 1.3E-01 | 2.5E+00 | 3.9E-01 | 1 |
| CCDC127 | -8.9E-01 | 1.9E-03 | -5.0E-01 | 2.1E-01 | 3.9E-01 | 4.7E-01 | 1 |
| PDE6D | -9.3E-01 | 1.5E-02 | -3.6E-01 | 6.6E-01 | 5.7E-01 | 3.5E-01 | 1 |
| FABP3 | -1.2E+00 | 4.5E-02 | -1.0E+00 | 1.5E-01 | 1.2E-01 | 9.9E-01 | 1 |
| LRP10 | -1.2E+00 | 9.1E-03 | -1.1E+00 | 5.7E-02 | 1.3E-01 | 9.9E-01 | 1 |
| TBRG1 | -7.5E-01 | 4.6E-03 | -4.2E-01 | 2.7E-01 | 3.3E-01 | 5.4E-01 | 1 |
| B3GALNT2 | -3.5E+00 | 1.7E-02 | -2.1E+00 | 2.2E-01 | 1.4E+00 | 7.8E-01 | 1 |
| CORO1C | -1.4E+00 | 2.6E-06 | -1.1E+00 | 4.9E-04 | 2.3E-01 | 8.7E-01 | 1 |
| NA | -2.7E+00 | 4.4E-02 | -2.4E+00 | 1.6E-01 | 3.0E-01 | 9.9E-01 | 1 |
| RPL4 | -3.3E+00 | 1.6E-02 | -2.1E+00 | 1.9E-01 | 1.2E+00 | 8.3E-01 | 1 |

| | | | | | | | |
|---|---|---|---|---|---|---|---|
| RPL4 | -3.3E+00 | 1.6E-02 | -2.1E+00 | 1.9E-01 | 1.2E+00 | 8.3E-01 | 1 |
| RPL4 | -3.3E+00 | 1.6E-02 | -2.1E+00 | 1.9E-01 | 1.2E+00 | 8.3E-01 | 1 |
| RPL4 | -3.3E+00 | 1.6E-02 | -2.1E+00 | 1.9E-01 | 1.2E+00 | 8.3E-01 | 1 |
| RPL4 | -3.3E+00 | 1.6E-02 | -2.1E+00 | 1.9E-01 | 1.2E+00 | 8.3E-01 | 1 |
| UBR1 | -1.4E+00 | 9.0E-08 | -1.2E+00 | 2.8E-05 | 1.9E-01 | 9.2E-01 | 1 |
| PITPNM1 | -8.9E-01 | 4.1E-02 | -2.7E-01 | 8.5E-01 | 6.2E-01 | 3.7E-01 | 1 |
| GAR1 | -2.3E+00 | 4.4E-06 | -2.0E+00 | 3.3E-04 | 3.0E-01 | 9.7E-01 | 1 |
| TOMM20 | -2.3E+00 | 5.6E-03 | -1.6E+00 | 1.1E-01 | 6.4E-01 | 8.7E-01 | 1 |
| TARBP1 | -2.4E+00 | 1.0E-03 | -2.0E+00 | 2.3E-02 | 4.4E-01 | 9.7E-01 | 1 |
| TTC21B | -1.8E+00 | 1.3E-02 | -7.4E-01 | 5.8E-01 | 1.0E+00 | 4.0E-01 | 1 |
| TTC21B | -1.8E+00 | 1.3E-02 | -7.4E-01 | 5.8E-01 | 1.0E+00 | 4.0E-01 | 1 |
| ATP11B | -6.0E-01 | 4.4E-02 | -3.1E-01 | 5.6E-01 | 2.9E-01 | 6.8E-01 | 1 |
| SCAND3 | -8.4E-01 | 3.2E-02 | -5.8E-01 | 2.9E-01 | 2.6E-01 | 9.1E-01 | 1 |
| POLG2 | -1.6E+00 | 9.7E-04 | -1.2E+00 | 3.5E-02 | 3.8E-01 | 8.9E-01 | 1 |
| SORL1 | -1.7E+00 | 1.4E-04 | -1.1E+00 | 6.2E-02 | 6.9E-01 | 4.3E-01 | 1 |
| TRAPPC2 | -7.6E-01 | 3.3E-02 | -3.8E-01 | 5.4E-01 | 3.8E-01 | 6.3E-01 | 1 |
| TRAPPC2 | -7.6E-01 | 3.3E-02 | -3.8E-01 | 5.4E-01 | 3.8E-01 | 6.3E-01 | 1 |
| SPRTN | -2.1E+00 | 7.6E-03 | -1.6E+00 | 1.0E-01 | 4.9E-01 | 9.7E-01 | 1 |
| C1orf131 | -5.9E-01 | 2.9E-03 | -3.6E-01 | 1.8E-01 | 2.3E-01 | 5.9E-01 | 1 |
| MOSPD2 | -1.4E+00 | 1.2E-02 | -1.1E+00 | 1.1E-01 | 2.8E-01 | 9.9E-01 | 1 |
| IMPDH2 | -1.1E+00 | 1.3E-03 | -7.9E-01 | 4.9E-02 | 2.7E-01 | 8.3E-01 | 1 |
| TARS2 | -1.3E+00 | 2.3E-02 | -1.1E+00 | 1.4E-01 | 2.4E-01 | 9.9E-01 | 1 |
| TARS2 | -1.3E+00 | 2.3E-02 | -1.1E+00 | 1.4E-01 | 2.4E-01 | 9.9E-01 | 1 |
| C1orf198 | -1.1E+00 | 2.1E-03 | -3.9E-01 | 5.6E-01 | 6.9E-01 | 1.6E-01 | 1 |
| C3H15orf38 | -9.5E-01 | 5.0E-03 | -5.1E-01 | 3.1E-01 | 4.5E-01 | 5.1E-01 | 1 |
| C3H15orf38 | -9.5E-01 | 5.0E-03 | -5.1E-01 | 3.1E-01 | 4.5E-01 | 5.1E-01 | 1 |
| IPO4 | -9.5E-01 | 1.9E-02 | -7.4E-01 | 1.5E-01 | 2.1E-01 | 9.9E-01 | 1 |
| IDH2 | -1.4E+00 | 1.1E-02 | -1.1E+00 | 1.0E-01 | 2.7E-01 | 9.9E-01 | 1 |
| ZNF311 | -1.1E+00 | 1.9E-02 | -1.0E+00 | 8.8E-02 | 1.2E-01 | 9.9E-01 | 1 |
| CIB1 | -1.1E+00 | 1.3E-02 | -7.7E-01 | 1.8E-01 | 3.2E-01 | 8.8E-01 | 1 |
| NGRN | -2.9E+00 | 2.7E-03 | -2.2E+00 | 5.3E-02 | 6.9E-01 | 9.4E-01 | 1 |
| NDUFB3 | -1.9E+00 | 4.3E-02 | -9.6E-01 | 5.6E-01 | 9.3E-01 | 6.6E-01 | 1 |
| URB2 | -2.3E+00 | 3.9E-03 | -1.6E+00 | 1.0E-01 | 6.4E-01 | 8.2E-01 | 1 |
| NUP133 | -1.9E+00 | 7.9E-04 | -1.5E+00 | 3.2E-02 | 4.6E-01 | 8.8E-01 | 1 |
| CCDC14 | -1.6E+00 | 3.6E-03 | -1.1E+00 | 1.1E-01 | 4.7E-01 | 8.3E-01 | 1 |
| CISD1 | -1.1E+00 | 1.1E-03 | -8.5E-01 | 3.5E-02 | 2.4E-01 | 9.1E-01 | 1 |
| TRAPPC4 | -2.0E+00 | 8.1E-05 | -1.6E+00 | 7.9E-03 | 4.3E-01 | 8.4E-01 | 1 |
| TRAPPC4 | -2.0E+00 | 8.1E-05 | -1.6E+00 | 7.9E-03 | 4.3E-01 | 8.4E-01 | 1 |
| NA | -1.8E+00 | 7.7E-06 | -1.2E+00 | 9.2E-03 | 5.7E-01 | 4.5E-01 | 1 |
| ATE1 | -1.1E+00 | 2.4E-04 | -5.9E-01 | 1.3E-01 | 4.9E-01 | 3.1E-01 | 1 |
| TAF4 | -1.7E+00 | 1.7E-05 | -1.4E+00 | 1.4E-03 | 2.7E-01 | 9.3E-01 | 1 |
| NA | -2.5E+00 | 3.4E-03 | -1.0E+00 | 4.4E-01 | 1.4E+00 | 2.9E-01 | 1 |
| SNX4 | -9.7E-01 | 7.6E-04 | -7.4E-01 | 3.2E-02 | 2.3E-01 | 8.4E-01 | 1 |
| OSBPL11 | -1.0E+00 | 9.2E-04 | -4.3E-01 | 3.6E-01 | 5.6E-01 | 2.1E-01 | 1 |
| LMLN | -1.7E+00 | 2.7E-03 | -1.2E+00 | 8.1E-02 | 4.7E-01 | 8.5E-01 | 1 |
| BCCIP | -8.8E-01 | 8.3E-03 | -8.0E-01 | 4.2E-02 | 7.7E-02 | 9.9E-01 | 1 |
| TMEM50A | -1.3E+00 | 1.0E-03 | -5.7E-01 | 3.3E-01 | 7.2E-01 | 2.4E-01 | 1 |
| NA | -9.5E-01 | 4.3E-05 | -6.4E-01 | 2.1E-02 | 3.1E-01 | 5.1E-01 | 1 |
| NA | -2.2E+00 | 1.2E-03 | -1.8E+00 | 2.6E-02 | 4.1E-01 | 9.7E-01 | 1 |
| TBL2 | -1.9E+00 | 4.6E-02 | -1.2E+00 | 3.6E-01 | 6.8E-01 | 9.0E-01 | 1 |
| NUBPL | -1.7E+00 | 8.2E-03 | -8.8E-01 | 3.7E-01 | 8.5E-01 | 5.2E-01 | 1 |
| NA | -2.6E+00 | 4.4E-02 | -1.5E+00 | 4.0E-01 | 1.1E+00 | 8.2E-01 | 1 |
| MBTPS2 | -1.3E+00 | 1.2E-02 | -8.1E-01 | 2.6E-01 | 4.8E-01 | 7.6E-01 | 1 |
| DLG1 | -1.1E+00 | 2.6E-05 | -6.0E-01 | 6.6E-02 | 4.8E-01 | 2.2E-01 | 1 |
| NA | -2.1E+00 | 3.4E-03 | -1.4E+00 | 1.3E-01 | 7.2E-01 | 7.5E-01 | 1 |
| NDUFS1 | -7.8E-01 | 1.1E-02 | -5.2E-01 | 2.1E-01 | 2.5E-01 | 8.2E-01 | 1 |
| TRA2B | -1.6E+00 | 2.0E-02 | -8.1E-01 | 4.5E-01 | 7.5E-01 | 6.1E-01 | 1 |
| BUD13 | -1.4E+00 | 1.3E-05 | -1.0E+00 | 4.6E-03 | 3.4E-01 | 6.8E-01 | 1 |
| MTX2 | -1.6E+00 | 1.9E-04 | -1.2E+00 | 2.1E-02 | 4.2E-01 | 7.4E-01 | 1 |
| NA | -1.6E+00 | 2.9E-04 | -1.1E+00 | 3.5E-02 | 4.8E-01 | 6.8E-01 | 1 |
| ZNF511 | -8.1E-01 | 4.5E-02 | -6.5E-01 | 2.2E-01 | 1.6E-01 | 9.9E-01 | 1 |
| XXYLT1 | -1.6E+00 | 1.8E-02 | -9.8E-01 | 3.3E-01 | 6.5E-01 | 7.4E-01 | 1 |
| REXO2 | -1.5E+00 | 2.2E-03 | -1.2E+00 | 4.2E-02 | 3.1E-01 | 9.7E-01 | 1 |
| NA | -3.9E+00 | 1.8E-02 | -1.6E+00 | 4.4E-01 | 2.3E+00 | 4.6E-01 | 1 |
| FRMD8 | -1.3E+00 | 1.1E-02 | -6.9E-01 | 3.8E-01 | 6.0E-01 | 5.6E-01 | 1 |
| B2M | -2.5E+00 | 2.8E-04 | -1.9E+00 | 1.8E-02 | 5.7E-01 | 8.2E-01 | 1 |
| DDX20 | -9.0E-01 | 6.2E-04 | -5.7E-01 | 8.9E-02 | 3.3E-01 | 5.6E-01 | 1 |
| RFC4 | -9.8E-01 | 3.8E-03 | -4.1E-01 | 4.9E-01 | 5.7E-01 | 2.8E-01 | 1 |
| LRWD1 | -8.2E-01 | 4.1E-02 | -6.9E-01 | 1.8E-01 | 1.4E-01 | 9.9E-01 | 1 |
| LRWD1 | -8.2E-01 | 4.1E-02 | -6.9E-01 | 1.8E-01 | 1.4E-01 | 9.9E-01 | 1 |
| ORAI2 | -8.8E-01 | 9.0E-03 | -3.0E-01 | 6.8E-01 | 5.8E-01 | 2.6E-01 | 1 |
| MIPOL1 | -1.4E+00 | 9.4E-04 | -1.1E+00 | 3.2E-02 | 3.2E-01 | 8.9E-01 | 1 |
| DDX50 | -1.6E+00 | 1.7E-05 | -1.3E+00 | 1.5E-03 | 2.7E-01 | 9.1E-01 | 1 |
| PLA2G4A | -7.3E-01 | 1.0E-02 | -6.3E-01 | 7.1E-02 | 1.0E-01 | 9.9E-01 | 1 |
| SEC23A | -1.7E+00 | 2.1E-02 | -8.1E-01 | 5.4E-01 | 8.9E-01 | 5.2E-01 | 1 |
| SUPV3L1 | -9.6E-01 | 2.9E-02 | -6.5E-01 | 3.0E-01 | 3.1E-01 | 8.9E-01 | 1 |
| NA | -1.3E+00 | 1.9E-02 | -4.3E-01 | 7.3E-01 | 8.2E-01 | 3.3E-01 | 1 |
| RTP4 | -1.7E+00 | 2.0E-02 | -1.4E+00 | 1.1E-01 | 2.5E-01 | 9.9E-01 | 1 |
| KIF18B | -2.7E+00 | 1.4E-02 | -2.1E+00 | 1.3E-01 | 6.1E-01 | 9.9E-01 | 1 |
| CCDC50 | -1.0E+00 | 2.2E-02 | -7.2E-01 | 2.4E-01 | 3.1E-01 | 9.1E-01 | 1 |
| GRN | -1.5E+00 | 1.5E-05 | -9.9E-01 | 1.9E-02 | 5.3E-01 | 4.0E-01 | 1 |
| IQCJ-SCHIP1 | -1.5E+00 | 1.2E-02 | -5.1E-01 | 7.2E-01 | 1.0E+00 | 2.5E-01 | 1 |
| IQCJ-SCHIP1 | -1.5E+00 | 1.2E-02 | -5.1E-01 | 7.2E-01 | 1.0E+00 | 2.5E-01 | 1 |
| MOSPD3 | -1.4E+00 | 3.1E-03 | -5.9E-01 | 4.6E-01 | 8.2E-01 | 2.7E-01 | 1 |
| DNAAF2 | -1.3E+00 | 4.4E-02 | -1.1E+00 | 2.2E-01 | 2.6E-01 | 9.9E-01 | 1 |
| MAP4K2 | -4.2E+00 | 2.5E-02 | -2.0E+00 | 4.4E-01 | 2.2E+00 | 6.1E-01 | 1 |
| DUT | -1.3E+00 | 9.5E-04 | -1.0E+00 | 3.2E-02 | 3.0E-01 | 8.9E-01 | 1 |
| ATP5S | -1.2E+00 | 5.0E-03 | -6.3E-01 | 3.4E-01 | 6.0E-01 | 4.5E-01 | 1 |
| PHB2 | -2.1E+00 | 2.6E-04 | -1.4E+00 | 4.8E-02 | 7.2E-01 | 5.7E-01 | 1 |
| PRDX6 | -9.7E-01 | 8.6E-05 | -6.7E-01 | 2.4E-02 | 3.0E-01 | 5.7E-01 | 1 |
| CUL5 | -7.5E-01 | 1.6E-03 | -6.6E-01 | 1.7E-02 | 8.6E-02 | 9.9E-01 | 1 |
| ARPC2 | -1.4E+00 | 1.8E-02 | -1.1E+00 | 1.5E-01 | 3.0E-01 | 9.9E-01 | 1 |
| FRMD6 | -7.6E-01 | 1.7E-02 | -4.7E-01 | 3.1E-01 | 3.0E-01 | 7.3E-01 | 1 |
| SLC11A1 | -2.0E+00 | 8.6E-03 | -1.4E+00 | 1.6E-01 | 6.7E-01 | 8.4E-01 | 1 |

| | | | | | | | |
|---|---|---|---|---|---|---|---|
| NA | -1.9E+00 | 3.3E-02 | -7.9E-01 | 6.2E-01 | 1.1E+00 | 5.3E-01 | 1 |
| CRLF1 | -1.6E+00 | 3.5E-03 | -1.4E+00 | 3.2E-02 | 2.1E-01 | 9.9E-01 | 1 |
| AASDHPPT | -1.0E+00 | 3.4E-03 | -5.2E-01 | 3.2E-01 | 5.1E-01 | 4.1E-01 | 1 |
| NA | -1.4E+00 | 3.4E-05 | -9.5E-01 | 2.1E-02 | 4.8E-01 | 4.8E-01 | 1 |
| NA | -1.4E+00 | 3.4E-05 | -9.5E-01 | 2.1E-02 | 4.8E-01 | 4.8E-01 | 1 |
| NA | -1.4E+00 | 3.4E-05 | -9.5E-01 | 2.1E-02 | 4.8E-01 | 4.8E-01 | 1 |
| NA | -1.4E+00 | 3.4E-05 | -9.5E-01 | 2.1E-02 | 4.8E-01 | 4.8E-01 | 1 |
| NAGLU | -1.2E+00 | 3.0E-03 | -8.8E-01 | 6.9E-02 | 2.8E-01 | 9.2E-01 | 1 |
| LGALS3 | -1.6E+00 | 1.1E-04 | -1.1E+00 | 3.7E-02 | 5.7E-01 | 4.9E-01 | 1 |
| NA | -2.0E+00 | 9.4E-03 | -1.6E+00 | 8.5E-02 | 3.7E-01 | 9.9E-01 | 1 |
| MED28 | -1.7E+00 | 2.8E-03 | -1.1E+00 | 1.1E-01 | 5.3E-01 | 7.4E-01 | 1 |
| ARPC1A | -1.3E+00 | 1.0E-03 | -1.0E+00 | 3.0E-02 | 2.7E-01 | 9.0E-01 | 1 |
| CD9 | -2.4E+00 | 2.8E-02 | -2.0E+00 | 1.4E-01 | 3.7E-01 | 9.9E-01 | 1 |
| CRKL | -1.1E+00 | 2.3E-03 | -7.9E-01 | 8.0E-02 | 3.1E-01 | 8.1E-01 | 1 |
| PLAU | -3.5E+00 | 1.4E-04 | -2.1E+00 | 7.3E-02 | 1.4E+00 | 3.6E-01 | 1 |
| PROM1 | -1.9E+00 | 9.2E-04 | -1.6E+00 | 2.0E-02 | 3.4E-01 | 9.8E-01 | 1 |
| ACTR10 | -1.1E+00 | 4.6E-04 | -6.6E-01 | 1.0E-01 | 4.3E-01 | 4.6E-01 | 1 |
| PSMA3 | -1.0E+00 | 2.5E-06 | -4.0E-01 | 2.1E-01 | 6.4E-01 | 1.8E-02 | 1 |
| AP4E1 | -6.4E-01 | 2.6E-02 | -3.1E-01 | 5.5E-01 | 3.3E-01 | 5.6E-01 | 1 |
| DCAF6 | -1.7E+00 | 1.7E-05 | -1.0E+00 | 3.1E-02 | 6.4E-01 | 3.1E-01 | 1 |
| MPC2 | -2.9E+00 | 3.7E-06 | -2.4E+00 | 5.1E-04 | 4.7E-01 | 9.0E-01 | 1 |
| VDAC2 | -3.0E+00 | 2.3E-05 | -2.5E+00 | 1.8E-03 | 5.1E-01 | 9.3E-01 | 1 |
| CREG1 | -7.2E-01 | 4.4E-02 | -2.8E-01 | 7.4E-01 | 4.4E-01 | 4.8E-01 | 1 |
| RPL14 | -2.5E+00 | 6.3E-03 | -1.8E+00 | 1.2E-01 | 7.3E-01 | 8.9E-01 | 1 |
| STAT5B | -1.7E+00 | 1.5E-04 | -1.3E+00 | 1.2E-02 | 3.7E-01 | 8.3E-01 | 1 |
| HAUS8 | -9.8E-01 | 1.1E-02 | -8.4E-01 | 7.7E-02 | 1.4E-01 | 9.9E-01 | 1 |
| DHRS7 | -2.4E+00 | 1.6E-03 | -2.1E+00 | 1.8E-02 | 3.2E-01 | 9.9E-01 | 1 |
| POGK | -1.1E+00 | 5.1E-04 | -7.1E-01 | 8.0E-02 | 4.0E-01 | 5.5E-01 | 1 |
| HDAC6 | -2.5E+00 | 6.3E-03 | -1.3E+00 | 2.7E-01 | 1.2E+00 | 5.6E-01 | 1 |
| TMEM179B | -1.1E+00 | 1.1E-03 | -6.7E-01 | 1.5E-01 | 4.7E-01 | 4.8E-01 | 1 |
| TMEM179B | -1.1E+00 | 1.1E-03 | -6.7E-01 | 1.5E-01 | 4.7E-01 | 4.8E-01 | 1 |
| KDELR2 | -1.2E+00 | 2.3E-04 | -6.4E-01 | 1.7E-01 | 6.1E-01 | 2.4E-01 | 1 |
| ADSS | -7.7E-01 | 4.2E-03 | -5.2E-01 | 1.4E-01 | 2.5E-01 | 7.4E-01 | 1 |
| SDHB | -2.2E+00 | 5.8E-04 | -1.3E+00 | 9.9E-02 | 8.3E-01 | 5.0E-01 | 1 |
| TSPAN14 | -1.2E+00 | 3.6E-02 | -7.2E-01 | 3.9E-01 | 4.6E-01 | 8.0E-01 | 1 |
| RNASEK | -1.0E+00 | 1.6E-04 | -8.5E-01 | 6.5E-03 | 1.7E-01 | 9.7E-01 | 1 |
| FADS3 | -1.5E+00 | 4.9E-03 | -1.3E+00 | 3.8E-02 | 1.7E-01 | 9.9E-01 | 1 |
| RPL9 | -2.1E+00 | 1.7E-06 | -1.1E+00 | 3.3E-02 | 9.4E-01 | 1.2E-01 | 1 |
| NA | -2.4E+00 | 4.1E-05 | -1.9E+00 | 4.5E-03 | 5.0E-01 | 8.4E-01 | 1 |
| NA | -2.4E+00 | 4.1E-05 | -1.9E+00 | 4.5E-03 | 5.0E-01 | 8.4E-01 | 1 |
| WDR26 | -1.0E+00 | 1.8E-04 | -8.0E-01 | 1.0E-02 | 2.0E-01 | 8.9E-01 | 1 |
| RAB24 | -2.1E+00 | 3.4E-04 | -1.5E+00 | 2.6E-02 | 5.4E-01 | 7.9E-01 | 1 |
| PSMG3 | -1.6E+00 | 1.3E-02 | -8.9E-01 | 3.4E-01 | 7.2E-01 | 6.4E-01 | 1 |
| RPL19 | -1.0E+00 | 5.1E-03 | -7.4E-01 | 1.2E-01 | 2.9E-01 | 8.3E-01 | 1 |
| DCAF5 | -8.7E-01 | 1.3E-02 | -6.8E-01 | 1.3E-01 | 1.9E-01 | 9.8E-01 | 1 |
| NA | -1.1E+00 | 2.5E-03 | -5.9E-01 | 2.6E-01 | 5.2E-01 | 4.6E-01 | 1 |
| IQGAP3 | -3.1E+00 | 1.2E-02 | -1.8E+00 | 2.4E-01 | 1.3E+00 | 7.1E-01 | 1 |
| PCNX | -9.9E-01 | 1.9E-02 | -7.0E-01 | 2.3E-01 | 3.0E-01 | 9.0E-01 | 1 |
| ABCD4 | -1.1E+00 | 4.0E-02 | -6.9E-01 | 3.9E-01 | 4.0E-01 | 8.4E-01 | 1 |
| DAP3 | -7.2E-01 | 3.9E-03 | -3.2E-01 | 4.4E-01 | 4.0E-01 | 3.2E-01 | 1 |
| NA | -1.7E+00 | 1.8E-02 | -1.0E+00 | 2.9E-01 | 6.3E-01 | 8.0E-01 | 1 |
| C1orf43 | -3.0E+00 | 1.1E-02 | -2.1E+00 | 1.4E-01 | 9.6E-01 | 8.8E-01 | 1 |
| FLRT2 | -2.0E+00 | 1.6E-04 | -1.6E+00 | 1.2E-02 | 4.5E-01 | 8.7E-01 | 1 |
| FLRT2 | -2.0E+00 | 1.6E-04 | -1.6E+00 | 1.2E-02 | 4.5E-01 | 8.7E-01 | 1 |
| USP43 | -1.2E+00 | 9.8E-03 | -7.1E-01 | 2.6E-01 | 4.5E-01 | 7.0E-01 | 1 |
| SPESP1 | -2.5E+00 | 8.3E-03 | -1.3E+00 | 2.9E-01 | 1.1E+00 | 6.0E-01 | 1 |
| SUPT4H1 | -1.9E+00 | 2.9E-04 | -1.5E+00 | 1.2E-02 | 3.5E-01 | 9.4E-01 | 1 |
| MED7 | -1.1E+00 | 2.2E-02 | -9.6E-01 | 1.2E-01 | 1.6E-01 | 9.9E-01 | 1 |
| C18orf25 | -1.1E+00 | 4.3E-03 | -7.8E-01 | 1.2E-01 | 3.3E-01 | 8.0E-01 | 1 |
| NA | -1.9E+00 | 1.4E-03 | -1.0E+00 | 2.3E-01 | 9.0E-01 | 3.7E-01 | 1 |
| COG7 | -2.5E+00 | 7.1E-05 | -2.2E+00 | 1.9E-03 | 2.8E-01 | 9.9E-01 | 1 |
| PIK3C3 | -8.3E-01 | 1.4E-02 | -2.9E-01 | 7.1E-01 | 5.4E-01 | 3.0E-01 | 1 |
| POLR3E | -2.4E+00 | 1.9E-03 | -9.6E-01 | 3.9E-01 | 1.4E+00 | 2.4E-01 | 1 |
| POLR3E | -2.4E+00 | 1.9E-03 | -9.6E-01 | 3.9E-01 | 1.4E+00 | 2.4E-01 | 1 |
| AP1M2 | -3.2E+00 | 3.2E-03 | -2.5E+00 | 4.5E-02 | 6.9E-01 | 9.8E-01 | 1 |
| ATG4D | -2.5E+00 | 1.8E-03 | -2.0E+00 | 3.8E-02 | 5.4E-01 | 9.7E-01 | 1 |
| BEX4 | -2.8E+00 | 7.4E-05 | -2.2E+00 | 6.8E-03 | 6.0E-01 | 8.4E-01 | 1 |
| NA | -2.6E+00 | 8.3E-03 | -2.0E+00 | 1.0E-01 | 5.9E-01 | 9.7E-01 | 1 |
| NA | -2.6E+00 | 8.3E-03 | -2.0E+00 | 1.0E-01 | 5.9E-01 | 9.7E-01 | 1 |
| ZNHIT3 | -1.9E+00 | 5.6E-05 | -1.4E+00 | 1.5E-02 | 5.7E-01 | 6.6E-01 | 1 |
| MYO19 | -2.4E+00 | 4.4E-03 | -1.3E+00 | 2.9E-01 | 1.2E+00 | 4.8E-01 | 1 |
| NA | -3.5E+00 | 5.5E-04 | -1.8E+00 | 1.3E-01 | 1.7E+00 | 3.5E-01 | 1 |
| NA | -1.4E+00 | 4.0E-03 | -6.7E-01 | 3.4E-01 | 6.9E-01 | 4.2E-01 | 1 |
| DSG2 | -9.4E-01 | 4.5E-04 | -4.4E-01 | 2.6E-01 | 5.0E-01 | 2.1E-01 | 1 |
| ETFA | -1.9E+00 | 9.7E-04 | -1.5E+00 | 2.0E-02 | 3.2E-01 | 9.8E-01 | 1 |
| SS18 | -1.4E+00 | 1.4E-06 | -5.5E-01 | 1.6E-01 | 8.3E-01 | 1.7E-02 | 1 |
| INSR | -1.6E+00 | 1.7E-02 | -5.9E-01 | 6.6E-01 | 9.9E-01 | 3.6E-01 | 1 |
| NA | -1.8E+00 | 5.3E-06 | -1.5E+00 | 9.4E-04 | 3.3E-01 | 8.4E-01 | 1 |
| NA | -1.7E+00 | 2.0E-04 | -1.2E+00 | 2.7E-02 | 4.9E-01 | 7.0E-01 | 1 |
| NA | -1.7E+00 | 2.0E-04 | -1.2E+00 | 2.7E-02 | 4.9E-01 | 7.0E-01 | 1 |
| NA | -1.7E+00 | 2.0E-04 | -1.2E+00 | 2.7E-02 | 4.9E-01 | 7.0E-01 | 1 |
| NA | -1.7E+00 | 2.0E-04 | -1.2E+00 | 2.7E-02 | 4.9E-01 | 7.0E-01 | 1 |
| NA | -1.7E+00 | 2.0E-04 | -1.2E+00 | 2.7E-02 | 4.9E-01 | 7.0E-01 | 1 |
| NA | -1.7E+00 | 2.0E-04 | -1.2E+00 | 2.7E-02 | 4.9E-01 | 7.0E-01 | 1 |
| PGRMC1 | -1.0E+00 | 5.1E-06 | -3.7E-01 | 2.9E-01 | 6.7E-01 | 1.6E-02 | 1 |
| SLC25A5 | -1.3E+00 | 1.9E-02 | -1.0E+00 | 1.3E-01 | 2.4E-01 | 9.9E-01 | 1 |
| UBE2A | -1.8E+00 | 9.1E-04 | -1.1E+00 | 1.1E-01 | 7.1E-01 | 5.5E-01 | 1 |
| NKRF | -8.1E-01 | 2.5E-02 | -4.5E-01 | 4.3E-01 | 3.6E-01 | 6.9E-01 | 1 |
| SREBF1 | -1.2E+00 | 2.6E-02 | -6.8E-01 | 4.2E-01 | 5.2E-01 | 6.9E-01 | 1 |
| CRLF3 | -8.5E-01 | 8.6E-03 | -7.2E-01 | 6.9E-02 | 1.3E-01 | 9.9E-01 | 1 |
| BIRC4 | -1.4E+00 | 1.4E-04 | -9.0E-01 | 3.7E-02 | 4.5E-01 | 5.5E-01 | 1 |
| BIRC4 | -1.4E+00 | 1.4E-04 | -9.0E-01 | 3.7E-02 | 4.5E-01 | 5.5E-01 | 1 |
| NA | -3.3E+00 | 2.2E-02 | -3.0E+00 | 9.4E-02 | 3.2E-01 | 9.9E-01 | 1 |
| UTP14A | -9.5E-01 | 1.4E-02 | -6.6E-01 | 2.1E-01 | 3.0E-01 | 8.6E-01 | 1 |

| | | | | | | | |
|---|---|---|---|---|---|---|---|
| UTP14A | -9.5E-01 | 1.4E-02 | -6.6E-01 | 2.1E-01 | 3.0E-01 | 8.6E-01 | 1 |
| NTAN1 | -7.4E-01 | 3.2E-03 | -2.7E-01 | 5.9E-01 | 4.8E-01 | 1.9E-01 | 1 |
| PSMG2 | -1.7E+00 | 1.9E-04 | -1.3E+00 | 1.3E-02 | 3.7E-01 | 8.4E-01 | 1 |
| TARS | -1.2E+00 | 1.5E-02 | -9.1E-01 | 1.3E-01 | 2.6E-01 | 9.8E-01 | 1 |
| HPRT | -1.0E+00 | 1.1E-02 | -7.1E-01 | 1.9E-01 | 3.3E-01 | 8.3E-01 | 1 |
| OMA1 | -1.9E+00 | 1.4E-02 | -9.3E-01 | 4.6E-01 | 9.7E-01 | 5.3E-01 | 1 |
| MYO5B | -6.6E-01 | 2.5E-02 | -3.3E-01 | 5.1E-01 | 3.2E-01 | 6.0E-01 | 1 |
| HSPB11 | -1.4E+00 | 1.3E-02 | -4.7E-01 | 7.3E-01 | 9.6E-01 | 2.7E-01 | 1 |
| GLOD4 | -1.6E+00 | 9.9E-05 | -1.0E+00 | 4.6E-02 | 6.0E-01 | 4.4E-01 | 1 |
| PLXNB3 | -2.8E+00 | 9.8E-04 | -2.3E+00 | 2.0E-02 | 4.9E-01 | 9.9E-01 | 1 |
| METTL16 | -1.4E+00 | 5.7E-03 | -6.7E-01 | 4.0E-01 | 7.1E-01 | 4.2E-01 | 1 |
| METTL16 | -1.4E+00 | 5.7E-03 | -6.7E-01 | 4.0E-01 | 7.1E-01 | 4.2E-01 | 1 |
| GADD45B | -1.5E+00 | 1.1E-02 | -8.6E-01 | 3.2E-01 | 6.3E-01 | 6.3E-01 | 1 |
| EHMT1 | -1.0E+00 | 1.9E-03 | -8.7E-01 | 2.1E-02 | 1.3E-01 | 9.9E-01 | 1 |
| NDUFB10 | -9.7E-01 | 8.1E-04 | -7.6E-01 | 2.7E-02 | 2.1E-01 | 9.0E-01 | 1 |
| TELO2 | -9.3E-01 | 1.6E-02 | -6.0E-01 | 2.7E-01 | 3.3E-01 | 8.1E-01 | 1 |
| GNPTG | -1.3E+00 | 9.9E-05 | -4.7E-01 | 3.9E-01 | 8.4E-01 | 5.1E-02 | 1 |
| NOTCH1 | -7.8E-01 | 2.0E-02 | -6.0E-01 | 1.6E-01 | 1.7E-01 | 9.9E-01 | 1 |
| VAMP7 | -1.5E+00 | 9.4E-03 | -1.4E+00 | 4.2E-02 | 1.0E-01 | 9.9E-01 | 1 |
| RBM15 | -3.3E+00 | 1.1E-03 | -2.9E+00 | 1.4E-02 | 3.9E-01 | 9.9E-01 | 1 |
| NA | -1.3E+00 | 3.7E-02 | -8.8E-01 | 2.9E-01 | 3.7E-01 | 9.4E-01 | 1 |
| NA | -1.3E+00 | 3.7E-02 | -8.8E-01 | 2.9E-01 | 3.7E-01 | 9.4E-01 | 1 |
| NA | -1.3E+00 | 3.7E-02 | -8.8E-01 | 2.9E-01 | 3.7E-01 | 9.4E-01 | 1 |
| NA | -1.3E+00 | 3.7E-02 | -8.8E-01 | 2.9E-01 | 3.7E-01 | 9.4E-01 | 1 |
| PSMA5 | -9.4E-01 | 5.7E-03 | -5.1E-01 | 3.1E-01 | 4.3E-01 | 5.2E-01 | 1 |
| PPP1R26 | -2.1E+00 | 2.7E-04 | -1.6E+00 | 1.5E-02 | 4.5E-01 | 8.8E-01 | 1 |
| ATMIN | -8.6E-01 | 5.3E-03 | -3.4E-01 | 5.4E-01 | 5.1E-01 | 2.9E-01 | 1 |
| SLC35A3 | -1.7E+00 | 4.7E-04 | -9.3E-01 | 1.5E-01 | 7.8E-01 | 3.6E-01 | 1 |
| KARS | -9.6E-01 | 1.6E-06 | -3.2E-01 | 3.0E-01 | 6.4E-01 | 7.0E-03 | 1 |
| BCAR1 | -2.6E+00 | 2.2E-03 | -1.2E+00 | 3.5E-01 | 1.3E+00 | 3.1E-01 | 1 |
| PTRH1 | -1.4E+00 | 3.7E-03 | -1.2E+00 | 3.8E-02 | 2.0E-01 | 9.9E-01 | 1 |
| GCLM | -2.9E+00 | 2.0E-02 | -2.2E+00 | 1.5E-01 | 7.1E-01 | 9.9E-01 | 1 |
| BCAR3 | -2.2E+00 | 9.4E-03 | -1.4E+00 | 2.3E-01 | 8.2E-01 | 7.2E-01 | 1 |
| RPL12 | -1.5E+00 | 3.4E-07 | -6.3E-01 | 9.8E-02 | 8.4E-01 | 1.8E-02 | 1 |
| MAPKAP1 | -1.1E+00 | 1.2E-02 | -6.5E-01 | 3.0E-01 | 4.4E-01 | 6.8E-01 | 1 |
| LRRC8C | -7.1E-01 | 4.3E-02 | -2.9E-01 | 7.1E-01 | 4.2E-01 | 5.1E-01 | 1 |
| LRRC8C | -7.1E-01 | 4.3E-02 | -2.9E-01 | 7.1E-01 | 4.2E-01 | 5.1E-01 | 1 |
| NA | -3.1E+00 | 4.1E-03 | -2.1E+00 | 1.0E-01 | 1.0E+00 | 8.0E-01 | 1 |
| PRKACB | -6.3E-01 | 2.4E-02 | -5.3E-01 | 1.3E-01 | 1.0E-01 | 9.9E-01 | 1 |
| FAM65A | -1.4E+00 | 3.9E-03 | -9.2E-01 | 1.3E-01 | 4.4E-01 | 7.8E-01 | 1 |
| USP33 | -7.8E-01 | 5.2E-03 | -4.7E-01 | 2.2E-01 | 3.1E-01 | 6.3E-01 | 1 |
| RABGGTB | -2.8E+00 | 7.1E-03 | -1.7E+00 | 2.2E-01 | 1.0E+00 | 6.9E-01 | 1 |
| RABGGTB | -2.8E+00 | 7.1E-03 | -1.7E+00 | 2.2E-01 | 1.0E+00 | 6.9E-01 | 1 |
| RABGGTB | -2.8E+00 | 7.1E-03 | -1.7E+00 | 2.2E-01 | 1.0E+00 | 6.9E-01 | 1 |
| RABGGTB | -2.8E+00 | 7.1E-03 | -1.7E+00 | 2.2E-01 | 1.0E+00 | 6.9E-01 | 1 |
| FAM96B | -2.0E+00 | 2.6E-02 | -1.1E+00 | 3.7E-01 | 8.1E-01 | 7.6E-01 | 1 |
| RANBP2 | -9.4E-01 | 3.1E-04 | -7.4E-01 | 1.7E-02 | 2.0E-01 | 8.7E-01 | 1 |
| METTL18 | -6.7E-01 | 2.8E-02 | -4.5E-01 | 3.0E-01 | 2.2E-01 | 8.7E-01 | 1 |
| NA | -3.0E+00 | 4.0E-02 | -2.9E+00 | 1.1E-01 | 1.1E-01 | 9.9E-01 | 1 |
| SERPINB7 | -1.5E+00 | 3.4E-02 | -8.5E-01 | 4.5E-01 | 6.8E-01 | 7.3E-01 | 1 |
| ZNF622 | -8.8E-01 | 1.0E-03 | -7.7E-01 | 1.5E-02 | 1.1E-01 | 9.9E-01 | 1 |
| SARNP | -2.0E+00 | 2.0E-04 | -1.5E+00 | 1.8E-02 | 4.8E-01 | 7.9E-01 | 1 |
| FASTKD5 | -1.6E+00 | 1.2E-05 | -9.0E-01 | 3.9E-02 | 7.0E-01 | 2.4E-01 | 1 |
| HNRNPD | -7.3E-01 | 3.7E-03 | -6.4E-01 | 2.9E-02 | 8.4E-02 | 9.9E-01 | 1 |
| LAMB1 | -9.4E-01 | 1.2E-03 | -5.0E-01 | 2.2E-01 | 4.4E-01 | 3.7E-01 | 1 |
| SPCS3 | -1.2E+00 | 3.9E-05 | -1.0E+00 | 2.2E-03 | 1.9E-01 | 9.5E-01 | 1 |
| HUS1 | -7.1E-01 | 9.2E-03 | -3.2E-01 | 4.9E-01 | 3.8E-01 | 4.2E-01 | 1 |
| LOX | -1.1E+00 | 1.7E-02 | -9.2E-01 | 1.2E-01 | 2.2E-01 | 9.9E-01 | 1 |
| GNG12 | -1.0E+00 | 5.0E-03 | -5.3E-01 | 3.3E-01 | 4.9E-01 | 4.7E-01 | 1 |
| C18orf32 | -1.1E+00 | 2.7E-02 | -4.9E-01 | 6.0E-01 | 6.0E-01 | 5.1E-01 | 1 |
| ZNF548 | -1.4E+00 | 2.0E-02 | -1.0E+00 | 2.0E-01 | 3.6E-01 | 9.6E-01 | 1 |
| IMPACT | -7.5E-01 | 2.0E-02 | -5.7E-01 | 1.7E-01 | 1.8E-01 | 9.8E-01 | 1 |
| NA | -3.4E+00 | 1.8E-03 | -2.4E+00 | 4.9E-02 | 9.9E-01 | 8.4E-01 | 1 |
| ACSL3 | -6.8E-01 | 1.5E-02 | -5.5E-01 | 1.1E-01 | 1.2E-01 | 9.9E-01 | 1 |
| ARF4 | -1.1E+00 | 2.9E-03 | -6.8E-01 | 1.6E-01 | 4.1E-01 | 6.2E-01 | 1 |
| NA | -1.6E+00 | 3.1E-02 | -1.1E+00 | 3.1E-01 | 5.4E-01 | 8.9E-01 | 1 |
| NFYB | -9.8E-01 | 1.6E-02 | -4.4E-01 | 5.5E-01 | 5.4E-01 | 4.5E-01 | 1 |
| TXNDC12 | -1.5E+00 | 3.5E-03 | -1.1E+00 | 9.6E-02 | 4.2E-01 | 8.3E-01 | 1 |
| ABT1 | -6.2E-01 | 3.7E-02 | -5.1E-01 | 1.8E-01 | 1.1E-01 | 9.9E-01 | 1 |
| RND3 | -1.6E+00 | 3.4E-02 | -6.4E-01 | 7.0E-01 | 9.6E-01 | 4.6E-01 | 1 |
| ZNF582 | -1.2E+00 | 2.2E-02 | -8.1E-01 | 2.9E-01 | 4.4E-01 | 8.3E-01 | 1 |
| FOPNL | -7.2E-01 | 2.0E-02 | -5.3E-01 | 1.9E-01 | 1.9E-01 | 9.5E-01 | 1 |
| NA | -1.6E+00 | 4.0E-04 | -1.0E+00 | 7.0E-02 | 5.8E-01 | 5.4E-01 | 1 |
| TMA7 | -2.2E+00 | 9.4E-03 | -1.8E+00 | 7.4E-02 | 3.7E-01 | 9.9E-01 | 1 |
| LAMTOR2 | -1.6E+00 | 3.3E-04 | -1.2E+00 | 2.7E-02 | 4.3E-01 | 7.7E-01 | 1 |
| PIGK | -9.9E-01 | 1.4E-03 | -5.1E-01 | 2.5E-01 | 4.8E-01 | 3.6E-01 | 1 |
| NA | -4.3E+00 | 4.4E-02 | -2.3E+00 | 5.1E-01 | 2.0E+00 | 7.8E-01 | 1 |
| C5orf15 | -1.7E+00 | 3.1E-03 | -9.9E-01 | 2.3E-01 | 7.5E-01 | 5.1E-01 | 1 |
| MMD | -1.1E+00 | 1.3E-05 | -9.4E-01 | 1.4E-03 | 2.0E-01 | 8.9E-01 | 1 |
| LYRM5 | -3.9E+00 | 6.0E-04 | -3.3E+00 | 1.1E-02 | 5.7E-01 | 9.9E-01 | 1 |
| COMMD6 | -1.7E+00 | 1.6E-03 | -1.2E+00 | 6.5E-02 | 4.9E-01 | 8.0E-01 | 1 |
| SRP14 | -8.7E-01 | 3.1E-03 | -7.9E-01 | 2.3E-02 | 8.3E-02 | 9.9E-01 | 1 |
| C11orf73 | -1.0E+00 | 9.6E-03 | -5.6E-01 | 3.7E-01 | 4.9E-01 | 5.4E-01 | 1 |
| SENP7 | -1.7E+00 | 1.4E-04 | -1.2E+00 | 3.0E-02 | 5.3E-01 | 5.9E-01 | 1 |
| IMPA1 | -8.1E-01 | 4.3E-02 | -3.1E-01 | 7.5E-01 | 5.0E-01 | 4.7E-01 | 1 |
| NA | -2.5E+00 | 2.0E-03 | -1.6E+00 | 1.4E-01 | 9.1E-01 | 6.1E-01 | 1 |
| LONRF1 | -7.4E-01 | 6.4E-03 | -3.5E-01 | 4.4E-01 | 4.0E-01 | 3.9E-01 | 1 |
| NA | -1.4E+00 | 3.2E-03 | -1.1E+00 | 5.8E-02 | 3.1E-01 | 9.5E-01 | 1 |
| CPPED1 | -1.2E+00 | 2.7E-04 | -6.0E-01 | 1.9E-01 | 5.9E-01 | 2.4E-01 | 1 |
| CISD3 | -7.2E-01 | 8.4E-03 | -3.1E-01 | 5.4E-01 | 4.1E-01 | 3.6E-01 | 1 |
| C3orf52 | -1.3E+00 | 2.7E-05 | -9.1E-01 | 1.6E-02 | 4.2E-01 | 5.2E-01 | 1 |
| BCL2L11 | -1.0E+00 | 2.4E-03 | -7.2E-01 | 9.1E-02 | 3.0E-01 | 7.8E-01 | 1 |
| NA | -1.9E+00 | 7.5E-05 | -9.6E-01 | 1.4E-01 | 9.5E-01 | 1.8E-01 | 1 |
| LLPH | -9.7E-01 | 2.3E-03 | -4.1E-01 | 4.3E-01 | 5.5E-01 | 2.5E-01 | 1 |

| | | | | | | | |
|---|---|---|---|---|---|---|---|
| SMIM11 | -2.6E+00 | 2.8E-06 | -2.0E+00 | 1.7E-03 | 6.8E-01 | 6.8E-01 | 1 |
| LCLAT1 | -9.6E-01 | 2.4E-03 | -4.3E-01 | 3.9E-01 | 5.3E-01 | 3.0E-01 | 1 |
| PERP | -6.3E-01 | 3.2E-03 | -2.2E-01 | 6.2E-01 | 4.1E-01 | 1.7E-01 | 1 |
| NA | -3.1E+00 | 2.3E-02 | -2.5E+00 | 1.4E-01 | 5.9E-01 | 9.9E-01 | 1 |
| NA | -1.5E+00 | 3.8E-04 | -9.5E-01 | 8.3E-02 | 5.8E-01 | 4.9E-01 | 1 |
| TIMM8B | -1.8E+00 | 4.2E-06 | -1.5E+00 | 3.6E-04 | 2.4E-01 | 9.7E-01 | 1 |
| NA | -8.8E-01 | 1.6E-03 | -4.8E-01 | 2.2E-01 | 4.0E-01 | 4.3E-01 | 1 |
| PDF | -4.7E+00 | 1.6E-02 | -2.6E+00 | 3.0E-01 | 2.1E+00 | 7.0E-01 | 1 |
| MANSC1 | -8.6E-01 | 5.0E-03 | -6.8E-01 | 6.9E-02 | 1.8E-01 | 9.7E-01 | 1 |
| NA | -7.5E-01 | 1.6E-02 | -2.9E-01 | 6.5E-01 | 4.6E-01 | 3.6E-01 | 1 |
| MED21 | -1.1E+00 | 2.5E-02 | -4.0E-01 | 7.1E-01 | 6.7E-01 | 4.0E-01 | 1 |
| MED21 | -1.1E+00 | 2.5E-02 | -4.0E-01 | 7.1E-01 | 6.7E-01 | 4.0E-01 | 1 |
| MPZL1 | -9.5E-01 | 2.4E-03 | -5.6E-01 | 1.9E-01 | 3.9E-01 | 5.4E-01 | 1 |
| RAP2A | -2.1E+00 | 1.4E-02 | -1.7E+00 | 1.0E-01 | 3.9E-01 | 9.9E-01 | 1 |
| EREG | -1.4E+00 | 4.5E-06 | -8.3E-01 | 3.2E-02 | 6.2E-01 | 1.8E-01 | 1 |
| NA | -2.0E+00 | 1.5E-02 | -1.5E+00 | 1.4E-01 | 4.5E-01 | 9.9E-01 | 1 |
| NA | -2.0E+00 | 1.5E-02 | -1.5E+00 | 1.4E-01 | 4.5E-01 | 9.9E-01 | 1 |
| NA | -2.0E+00 | 1.5E-02 | -1.5E+00 | 1.4E-01 | 4.5E-01 | 9.9E-01 | 1 |
| NA | -2.0E+00 | 1.5E-02 | -1.5E+00 | 1.4E-01 | 4.5E-01 | 9.9E-01 | 1 |
| NA | -2.0E+00 | 1.5E-02 | -1.5E+00 | 1.4E-01 | 4.5E-01 | 9.9E-01 | 1 |
| TIMM22 | -1.1E+00 | 1.7E-02 | -6.3E-01 | 3.5E-01 | 4.5E-01 | 6.9E-01 | 1 |
| GRPEL2 | -9.9E-01 | 3.3E-04 | -6.8E-01 | 4.3E-02 | 3.1E-01 | 6.5E-01 | 1 |
| CRADD | -1.4E+00 | 2.5E-07 | -8.7E-01 | 4.9E-03 | 5.0E-01 | 2.4E-01 | 1 |
| CGRRF1 | -8.2E-01 | 2.9E-02 | -6.2E-01 | 2.1E-01 | 2.0E-01 | 9.8E-01 | 1 |
| COX7A2L | -1.2E+00 | 8.2E-03 | -8.2E-01 | 1.5E-01 | 3.4E-01 | 8.4E-01 | 1 |
| PPIF | -1.2E+00 | 4.5E-03 | -7.0E-01 | 2.5E-01 | 5.1E-01 | 5.6E-01 | 1 |
| UBE2D1 | -9.6E-01 | 3.7E-02 | -3.8E-01 | 7.1E-01 | 5.8E-01 | 4.7E-01 | 1 |
| SF3B5 | -1.9E+00 | 1.7E-03 | -1.8E+00 | 1.3E-02 | 1.5E-01 | 9.9E-01 | 1 |
| VGLL3 | -1.4E+00 | 1.1E-02 | -8.8E-01 | 2.5E-01 | 5.4E-01 | 7.5E-01 | 1 |
| EMD | -1.2E+00 | 3.3E-03 | -9.7E-01 | 4.2E-02 | 2.0E-01 | 9.9E-01 | 1 |
| SNRPG | -2.5E+00 | 8.6E-03 | -1.6E+00 | 2.4E-01 | 9.6E-01 | 6.9E-01 | 1 |
| WNT7A | -8.1E-01 | 3.9E-02 | -4.5E-01 | 4.9E-01 | 3.7E-01 | 7.1E-01 | 1 |
| TFB2M | -6.6E-01 | 3.7E-02 | -4.1E-01 | 3.8E-01 | 2.4E-01 | 8.3E-01 | 1 |
| GTF2H5 | -1.6E+00 | 5.4E-04 | -1.1E+00 | 4.2E-02 | 4.6E-01 | 7.3E-01 | 1 |
| NA | -2.3E+00 | 4.9E-03 | -1.7E+00 | 1.2E-01 | 6.9E-01 | 8.4E-01 | 1 |
| NA | -2.3E+00 | 1.2E-03 | -1.6E+00 | 6.3E-02 | 6.8E-01 | 7.7E-01 | 1 |
| NA | -4.9E+00 | 4.9E-03 | -1.3E+00 | 6.2E-01 | 3.6E+00 | 1.3E-01 | 1 |
| NA | -7.9E-01 | 1.3E-02 | -3.6E-01 | 5.3E-01 | 4.4E-01 | 4.4E-01 | 1 |
| NA | -1.8E+00 | 1.7E-02 | -1.0E+00 | 3.3E-01 | 7.5E-01 | 7.2E-01 | 1 |
| CNOT2 | -3.3E-02 | 9.8E-01 | -5.9E-01 | 4.4E-03 | -5.6E-01 | 9.0E-03 | 7 |
| UTP23 | -1.7E-01 | 9.8E-01 | -1.9E+00 | 4.6E-03 | -1.8E+00 | 1.5E-02 | 7 |
| FAM53C | -9.1E-01 | 3.3E-02 | -2.9E+00 | 8.3E-12 | -2.0E+00 | 5.3E-06 | 7 |
| PTPLAD2 | -4.6E-01 | 6.0E-01 | -1.5E+00 | 3.3E-02 | -1.0E+00 | 2.6E-01 | 7 |
| HMGXB4 | -2.3E-01 | 8.1E-01 | -1.9E+00 | 1.9E-04 | -1.6E+00 | 1.9E-03 | 7 |
| IREB2 | -6.6E-01 | 7.1E-02 | -2.3E+00 | 1.4E-11 | -1.6E+00 | 3.2E-06 | 7 |
| ZFAND5 | -2.0E-01 | 5.8E-01 | -7.9E-01 | 1.2E-03 | -5.8E-01 | 3.1E-02 | 7 |
| TMEM2 | -1.9E-01 | 6.7E-01 | -1.8E+00 | 3.3E-14 | -1.6E+00 | 5.4E-13 | 7 |
| SAMD9L | -8.3E-01 | 5.6E-02 | -2.6E+00 | 1.9E-08 | -1.8E+00 | 3.9E-04 | 7 |
| FAM214B | 2.6E-01 | 7.6E-01 | -1.2E+00 | 1.2E-02 | -1.4E+00 | 1.2E-03 | 7 |
| MAP4K4 | -3.0E-01 | 2.2E-01 | -8.9E-01 | 4.1E-05 | -5.9E-01 | 1.6E-02 | 7 |
| STX17 | -1.6E-01 | 9.7E-01 | -2.8E+00 | 5.5E-05 | -2.6E+00 | 2.3E-04 | 7 |
| XPO1 | -2.7E-01 | 5.7E-01 | -9.5E-01 | 4.4E-03 | -6.8E-01 | 7.9E-02 | 7 |
| HDHD3 | -2.6E-01 | 8.7E-01 | -1.5E+00 | 2.1E-02 | -1.3E+00 | 9.8E-02 | 7 |
| AMD1 | -1.5E-01 | 9.4E-01 | -1.2E+00 | 9.3E-03 | -1.0E+00 | 3.6E-02 | 7 |
| GTF3C6 | -2.1E-01 | 7.7E-01 | -1.3E+00 | 2.3E-04 | -1.1E+00 | 3.1E-03 | 7 |
| C2orf44 | -1.6E-02 | 9.8E-01 | -1.1E+00 | 2.1E-02 | -1.1E+00 | 2.8E-02 | 7 |
| SVIL | -2.1E-01 | 9.2E-01 | -1.3E+00 | 2.1E-02 | -1.1E+00 | 8.1E-02 | 7 |
| WDR33 | -4.2E-01 | 4.8E-01 | -2.2E+00 | 4.1E-06 | -1.8E+00 | 3.4E-04 | 7 |
| WDR33 | -4.2E-01 | 4.8E-01 | -2.2E+00 | 4.1E-06 | -1.8E+00 | 3.4E-04 | 7 |
| MASTL | -3.0E-01 | 8.5E-01 | -1.8E+00 | 8.8E-03 | -1.5E+00 | 5.1E-02 | 7 |
| VASP | 1.2E-02 | 9.8E-01 | -9.6E-01 | 6.5E-05 | -9.8E-01 | 3.0E-05 | 7 |
| FOSB | 7.9E-02 | 9.8E-01 | -9.3E-01 | 3.9E-02 | -1.0E+00 | 2.4E-02 | 7 |
| TMCC1 | -2.9E-01 | 3.2E-01 | -7.5E-01 | 2.5E-03 | -4.7E-01 | 1.3E-01 | 7 |
| CLASP1 | -3.3E-01 | 6.9E-01 | -1.2E+00 | 2.1E-02 | -8.4E-01 | 1.8E-01 | 7 |
| GATA3 | -1.3E-01 | 8.6E-01 | -7.4E-01 | 5.2E-03 | -6.1E-01 | 3.3E-02 | 7 |
| INSIG1 | -8.2E-02 | 9.8E-01 | -9.5E-01 | 1.1E-02 | -8.7E-01 | 2.8E-02 | 7 |
| RELT | -5.9E-01 | 1.2E-01 | -2.1E+00 | 7.3E-08 | -1.5E+00 | 2.6E-04 | 7 |
| TOP2B | -1.2E-02 | 9.8E-01 | -8.7E-01 | 1.0E-02 | -8.6E-01 | 1.4E-02 | 7 |
| HBEGF | -5.3E-01 | 1.4E-01 | -2.0E+00 | 1.2E-08 | -1.4E+00 | 6.4E-05 | 7 |
| ARHGDIA | -1.7E-01 | 9.8E-01 | -1.6E+00 | 2.1E-02 | -1.4E+00 | 5.9E-02 | 7 |
| NA | -3.7E-01 | 8.8E-01 | -2.4E+00 | 3.5E-02 | -2.1E+00 | 1.3E-01 | 7 |
| FGFR1 | 2.0E-01 | 8.7E-01 | -1.0E+00 | 3.3E-02 | -1.2E+00 | 8.3E-03 | 7 |
| TRIM21 | 2.0E-01 | 7.5E-01 | -1.1E+00 | 8.4E-04 | -1.3E+00 | 2.6E-05 | 7 |
| NOP56 | -2.0E-01 | 5.7E-01 | -6.3E-01 | 9.2E-03 | -4.3E-01 | 1.4E-01 | 7 |
| NOP56 | -2.0E-01 | 5.7E-01 | -6.3E-01 | 9.2E-03 | -4.3E-01 | 1.4E-01 | 7 |
| NOP56 | -2.0E-01 | 5.7E-01 | -6.3E-01 | 9.2E-03 | -4.3E-01 | 1.4E-01 | 7 |
| NOP56 | -2.0E-01 | 5.7E-01 | -6.3E-01 | 9.2E-03 | -4.3E-01 | 1.4E-01 | 7 |
| NOP56 | -2.0E-01 | 5.7E-01 | -6.3E-01 | 9.2E-03 | -4.3E-01 | 1.4E-01 | 7 |
| STX2 | -2.3E-02 | 9.8E-01 | -1.1E+00 | 1.7E-02 | -1.0E+00 | 2.3E-02 | 7 |
| AMOTL2 | -4.2E-01 | 6.5E-01 | -2.0E+00 | 2.1E-04 | -1.6E+00 | 6.2E-03 | 7 |
| UTP20 | -2.6E-01 | 5.1E-01 | -7.7E-01 | 1.1E-02 | -5.1E-01 | 1.7E-01 | 7 |
| PLK2 | -4.8E-01 | 5.3E-01 | -1.3E+00 | 2.7E-02 | -8.7E-01 | 2.9E-01 | 7 |
| CENPJ | -2.0E-01 | 9.3E-01 | -2.0E+00 | 1.9E-04 | -1.8E+00 | 1.1E-03 | 7 |
| ABHD6 | -2.0E-01 | 6.1E-01 | -8.0E-01 | 2.1E-03 | -6.0E-01 | 3.8E-02 | 7 |
| GPATCH1 | -2.2E-01 | 7.8E-01 | -1.1E+00 | 8.5E-03 | -8.7E-01 | 6.3E-02 | 7 |
| IMMT | 8.7E-02 | 9.8E-01 | -7.3E-01 | 7.1E-02 | -8.2E-01 | 4.1E-02 | 7 |
| COPB2 | 1.1E-01 | 8.7E-01 | -8.3E-01 | 4.5E-04 | -9.4E-01 | 2.4E-05 | 7 |
| NA | 1.8E-01 | 8.0E-01 | -9.0E-01 | 9.9E-03 | -1.1E+00 | 9.7E-04 | 7 |
| KIAA0922 | 1.7E-01 | 9.5E-01 | -1.2E+00 | 8.4E-03 | -1.4E+00 | 2.0E-03 | 7 |
| CCNJ | 9.7E-02 | 9.8E-01 | -9.8E-01 | 1.3E-02 | -1.1E+00 | 5.4E-03 | 7 |
| OAS3 | -7.9E-01 | 1.0E-01 | -3.6E+00 | 3.4E-08 | -2.8E+00 | 5.1E-05 | 7 |
| HPS5 | 1.9E-01 | 9.0E-01 | -9.0E-01 | 9.0E-02 | -1.1E+00 | 3.0E-02 | 7 |
| DCBLD2 | -3.6E-01 | 3.1E-01 | -1.4E+00 | 5.7E-06 | -1.0E+00 | 1.6E-03 | 7 |
| CHD6 | 1.6E-01 | 8.4E-01 | -9.0E-01 | 5.2E-03 | -1.1E+00 | 5.4E-04 | 7 |

| | | | | | | | |
|---|---|---|---|---|---|---|---|
| SLC38A2 | 3.1E-03 | 9.9E-01 | -1.2E+00 | 1.5E-06 | -1.2E+00 | 5.6E-07 | 7 |
| SERPINB1 | -2.4E-01 | 4.5E-01 | -8.6E-01 | 3.4E-04 | -6.2E-01 | 1.9E-02 | 7 |
| HERC6 | -5.1E-01 | 5.0E-01 | -1.3E+00 | 2.5E-02 | -8.3E-01 | 3.1E-01 | 7 |
| MYLIP | -2.3E-01 | 6.5E-01 | -1.1E+00 | 3.1E-04 | -9.1E-01 | 7.6E-03 | 7 |
| RBMS1 | -3.8E-01 | 8.9E-02 | -1.3E+00 | 2.6E-10 | -8.9E-01 | 1.5E-05 | 7 |
| IKBKE | -6.3E-01 | 4.2E-01 | -2.6E+00 | 7.7E-04 | -2.0E+00 | 2.6E-02 | 7 |
| CHSY1 | -5.1E-01 | 5.1E-01 | -1.5E+00 | 2.4E-02 | -9.4E-01 | 2.8E-01 | 7 |
| TIMM17A | -2.1E-01 | 5.7E-01 | -5.9E-01 | 2.3E-02 | -3.8E-01 | 2.6E-01 | 7 |
| SP140 | 2.2E-01 | 8.6E-01 | -9.5E-01 | 9.2E-02 | -1.2E+00 | 2.6E-02 | 7 |
| SP140 | 2.2E-01 | 8.6E-01 | -9.5E-01 | 9.2E-02 | -1.2E+00 | 2.6E-02 | 7 |
| TXLNA | -2.8E-02 | 9.8E-01 | -6.4E-01 | 1.0E-02 | -6.2E-01 | 1.7E-02 | 7 |
| ZMYND8 | 1.9E-01 | 6.8E-01 | -7.3E-01 | 6.7E-03 | -9.1E-01 | 2.6E-04 | 7 |
| HEATR1 | -3.9E-01 | 5.3E-01 | -1.1E+00 | 2.5E-02 | -7.4E-01 | 2.7E-01 | 7 |
| TCF7L2 | -1.6E-01 | 7.5E-01 | -8.8E-01 | 4.4E-04 | -7.1E-01 | 6.9E-03 | 7 |
| FSTL1 | -1.3E-01 | 9.8E-01 | -1.3E+00 | 2.4E-02 | -1.2E+00 | 6.0E-02 | 7 |
| PRKX | -3.7E-01 | 1.9E-01 | -1.3E+00 | 1.1E-07 | -9.3E-01 | 2.3E-04 | 7 |
| ZNFX1 | 1.8E-01 | 8.7E-01 | -7.5E-01 | 6.2E-02 | -9.3E-01 | 1.6E-02 | 7 |
| ACIN1 | -5.3E-01 | 3.2E-01 | -1.3E+00 | 4.6E-03 | -7.6E-01 | 2.0E-01 | 7 |
| SOX2 | -2.8E-01 | 8.5E-01 | -1.6E+00 | 9.4E-03 | -1.3E+00 | 5.4E-02 | 7 |
| ALPK1 | -5.9E-01 | 7.3E-01 | -2.3E+00 | 4.5E-02 | -1.7E+00 | 2.4E-01 | 7 |
| C4orf21 | -1.7E-01 | 9.8E-01 | -2.7E+00 | 5.5E-03 | -2.5E+00 | 1.7E-02 | 7 |
| CD86 | 1.6E-01 | 9.8E-01 | -1.2E+00 | 4.1E-02 | -1.3E+00 | 1.9E-02 | 7 |
| PER2 | -8.6E-02 | 9.8E-01 | -1.1E+00 | 3.6E-02 | -1.0E+00 | 7.4E-02 | 7 |
| SS18L1 | 1.2E-01 | 9.0E-01 | -6.7E-01 | 3.5E-02 | -7.9E-01 | 9.2E-03 | 7 |
| FAM175B | 4.1E-02 | 9.8E-01 | -9.9E-01 | 2.4E-04 | -1.0E+00 | 7.1E-05 | 7 |
| UBE4A | -1.7E-01 | 7.8E-01 | -7.1E-01 | 2.6E-02 | -5.3E-01 | 1.6E-01 | 7 |
| CDK1 | -1.4E+00 | 2.8E-01 | -4.9E+00 | 6.7E-03 | -3.5E+00 | 1.3E-01 | 7 |
| STPG1 | -3.1E-01 | 3.6E-01 | -8.4E-01 | 3.9E-03 | -5.3E-01 | 1.5E-01 | 7 |
| LIMK2 | -2.8E-01 | 7.0E-01 | -1.3E+00 | 2.1E-03 | -9.9E-01 | 2.9E-02 | 7 |
| WHAMM | -5.4E-01 | 3.7E-01 | -1.5E+00 | 8.3E-03 | -9.2E-01 | 2.1E-01 | 7 |
| TAGLN | -3.0E-01 | 7.4E-01 | -1.2E+00 | 1.9E-02 | -9.1E-01 | 1.4E-01 | 7 |
| PRR14L | -3.0E-01 | 6.5E-01 | -8.8E-01 | 4.7E-02 | -5.8E-01 | 3.3E-01 | 7 |
| HTR1D | -3.6E-01 | 7.5E-01 | -1.3E+00 | 3.9E-02 | -9.6E-01 | 2.3E-01 | 7 |
| FASTKD2 | -3.7E-01 | 4.0E-01 | -1.1E+00 | 2.2E-03 | -7.4E-01 | 8.8E-02 | 7 |
| KLHL28 | -2.0E-01 | 4.6E-01 | -1.0E+00 | 1.9E-07 | -8.1E-01 | 3.2E-05 | 7 |
| ALG9 | -3.3E-01 | 4.0E-01 | -8.6E-01 | 1.0E-02 | -5.3E-01 | 2.3E-01 | 7 |
| CWC22 | 7.7E-02 | 9.8E-01 | -1.3E+00 | 1.3E-03 | -1.4E+00 | 5.1E-04 | 7 |
| DHX8 | 3.3E-02 | 9.8E-01 | -7.5E-01 | 4.0E-02 | -7.9E-01 | 3.4E-02 | 7 |
| GOLIM4 | -3.2E-01 | 5.8E-01 | -1.1E+00 | 1.0E-02 | -7.5E-01 | 1.4E-01 | 7 |
| IFI35 | -3.6E-01 | 5.5E-01 | -1.1E+00 | 1.2E-02 | -7.5E-01 | 1.7E-01 | 7 |
| ZNF394 | -2.9E-01 | 6.7E-01 | -9.2E-01 | 4.4E-02 | -6.2E-01 | 3.0E-01 | 7 |
| CAPZB | -5.3E-01 | 6.0E-01 | -1.7E+00 | 2.9E-02 | -1.2E+00 | 2.5E-01 | 7 |
| SUV39H1 | 2.4E-01 | 9.7E-01 | -1.7E+00 | 4.9E-02 | -2.0E+00 | 2.3E-02 | 7 |
| TRMT5 | -3.2E-01 | 6.5E-01 | -1.3E+00 | 7.0E-03 | -9.4E-01 | 8.2E-02 | 7 |
| SNAPC1 | -1.8E-01 | 8.5E-01 | -8.5E-01 | 2.3E-02 | -6.8E-01 | 1.2E-01 | 7 |
| LIN9 | -4.7E-01 | 4.0E-01 | -1.2E+00 | 1.3E-02 | -7.4E-01 | 2.6E-01 | 7 |
| H3F3C | -4.3E-01 | 3.3E-01 | -1.2E+00 | 3.2E-03 | -7.2E-01 | 1.4E-01 | 7 |
| FNTB | -1.9E-01 | 7.4E-01 | -7.7E-01 | 9.8E-03 | -5.8E-01 | 8.7E-02 | 7 |
| FNTB | -1.9E-01 | 7.4E-01 | -7.7E-01 | 9.8E-03 | -5.8E-01 | 8.7E-02 | 7 |
| LFNG | -2.9E-01 | 8.2E-01 | -1.7E+00 | 2.1E-03 | -1.4E+00 | 1.8E-02 | 7 |
| ZFYVE26 | 7.0E-02 | 9.8E-01 | -1.7E+00 | 9.6E-11 | -1.7E+00 | 5.4E-13 | 7 |
| EFNB1 | -2.9E-01 | 6.9E-01 | -1.3E+00 | 1.9E-03 | -9.9E-01 | 2.8E-02 | 7 |
| PFAS | -1.3E-01 | 9.8E-01 | -2.4E+00 | 8.9E-03 | -2.3E+00 | 1.9E-02 | 7 |
| RARS | -3.5E-02 | 9.8E-01 | -6.1E-01 | 2.1E-02 | -5.7E-01 | 3.8E-02 | 7 |
| EPN3 | -3.8E-01 | 4.3E-01 | -8.7E-01 | 3.1E-02 | -4.8E-01 | 4.2E-01 | 7 |
| SLU7 | -3.9E-01 | 7.8E-02 | -9.7E-01 | 3.0E-06 | -5.8E-01 | 1.3E-02 | 7 |
| TPM3 | -1.0E+00 | 6.9E-01 | -5.5E+00 | 7.3E-03 | -4.5E+00 | 6.0E-02 | 7 |
| PIAS1 | -1.5E-03 | 1.0E+00 | -8.2E-01 | 1.9E-04 | -8.2E-01 | 1.4E-04 | 7 |
| ARHGAP17 | 1.1E-02 | 9.8E-01 | -6.1E-01 | 7.4E-03 | -6.3E-01 | 6.6E-03 | 7 |
| TNRC6A | -3.7E-01 | 2.5E-01 | -1.3E+00 | 2.6E-06 | -9.0E-01 | 1.6E-03 | 7 |
| RBBP6 | -3.9E-01 | 2.1E-01 | -1.2E+00 | 1.8E-05 | -7.8E-01 | 1.0E-02 | 7 |
| NA | -3.8E-01 | 8.9E-02 | -1.2E+00 | 2.2E-08 | -7.8E-01 | 3.2E-04 | 7 |
| DICER1 | 6.5E-03 | 9.9E-01 | -1.0E+00 | 1.2E-04 | -1.0E+00 | 6.9E-05 | 7 |
| MAP2K4 | -7.5E-01 | 2.0E-01 | -2.1E+00 | 5.9E-04 | -1.4E+00 | 7.0E-02 | 7 |
| SEMA7A | -4.3E-01 | 5.4E-01 | -2.0E+00 | 2.3E-05 | -1.5E+00 | 1.4E-03 | 7 |
| ZNF839 | 1.5E-01 | 9.5E-01 | -9.7E-01 | 5.1E-02 | -1.1E+00 | 2.0E-02 | 7 |
| RBBP8 | -4.2E-01 | 1.5E-01 | -1.2E+00 | 5.8E-06 | -7.7E-01 | 8.4E-03 | 7 |
| KIAA0430 | -9.1E-02 | 9.5E-01 | -7.1E-01 | 6.4E-03 | -6.2E-01 | 2.5E-02 | 7 |
| NSRP1 | 1.7E-02 | 9.8E-01 | -1.4E+00 | 2.9E-05 | -1.4E+00 | 1.3E-05 | 7 |
| TMEM210 | -2.2E-01 | 6.7E-01 | -7.4E-01 | 2.3E-02 | -5.2E-01 | 1.9E-01 | 7 |
| IKBKG | -1.1E-01 | 9.7E-01 | -1.2E+00 | 2.8E-04 | -1.1E+00 | 1.1E-03 | 7 |
| SPTAN1 | -1.3E-01 | 8.7E-01 | -7.9E-01 | 8.9E-03 | -6.6E-01 | 4.6E-02 | 7 |
| GAPVD1 | 1.5E-01 | 8.5E-01 | -8.5E-01 | 6.1E-03 | -1.0E+00 | 6.7E-04 | 7 |
| MT-ND1 | 3.3E-02 | 9.8E-01 | -1.1E+00 | 2.8E-05 | -1.2E+00 | 8.2E-06 | 7 |
| MT-ND6 | -3.2E-01 | 8.2E-01 | -2.2E+00 | 1.1E-04 | -1.9E+00 | 1.3E-03 | 7 |
| MT-CYB | 6.0E-02 | 9.7E-01 | -7.0E-01 | 8.4E-05 | -7.6E-01 | 8.2E-06 | 7 |
| NA | -5.3E-01 | 5.8E-01 | -1.8E+00 | 1.5E-02 | -1.3E+00 | 1.6E-01 | 7 |
| EXO5 | 5.7E-01 | 7.2E-01 | -2.9E+00 | 4.6E-02 | -3.5E+00 | 1.7E-02 | 7 |
| NA | 4.1E-01 | 9.1E-01 | -3.7E+00 | 2.8E-02 | -4.1E+00 | 1.9E-02 | 7 |
| ANKRD49 | -1.7E-01 | 8.2E-01 | -8.6E-01 | 7.5E-03 | -7.0E-01 | 5.1E-02 | 7 |
| SLC35D1 | 1.6E-01 | 9.8E-01 | -1.4E+00 | 1.1E-02 | -1.6E+00 | 4.0E-03 | 7 |
| HAS2 | -9.3E-01 | 1.3E-01 | -2.7E+00 | 1.7E-06 | -1.7E+00 | 4.1E-03 | 7 |
| NA | -7.0E-01 | 7.0E-01 | -5.4E+00 | 2.0E-03 | -4.7E+00 | 1.7E-02 | 7 |
| NA | -1.5E-01 | 9.8E-01 | -1.9E+00 | 9.7E-03 | -1.8E+00 | 2.6E-02 | 7 |
| PLAGL2 | -5.8E-01 | 2.0E-01 | -2.3E+00 | 5.4E-09 | -1.8E+00 | 1.5E-05 | 7 |
| FAM111A | -2.7E-01 | 6.6E-01 | -1.1E+00 | 6.5E-03 | -8.1E-01 | 7.6E-02 | 7 |
| NA | -4.3E-01 | 2.8E-01 | -1.6E+00 | 6.1E-06 | -1.1E+00 | 2.3E-03 | 7 |
| RAP2B | 1.1E-01 | 9.8E-01 | -2.1E+00 | 1.6E-02 | -2.2E+00 | 1.3E-02 | 7 |
| CBX8 | -4.8E-01 | 3.2E-01 | -1.5E+00 | 4.9E-04 | -9.8E-01 | 4.4E-02 | 7 |
| MSANTD4 | 2.2E-01 | 7.8E-01 | -1.2E+00 | 2.5E-03 | -1.4E+00 | 1.6E-04 | 7 |
| CBX2 | 1.8E-01 | 8.8E-01 | -7.7E-01 | 7.9E-02 | -9.5E-01 | 2.3E-02 | 7 |
| NMNAT1 | -3.6E-01 | 4.8E-01 | -1.6E+00 | 8.8E-05 | -1.2E+00 | 5.8E-03 | 7 |
| NA | 2.6E-01 | 9.0E-01 | -1.7E+00 | 9.4E-03 | -1.9E+00 | 1.9E-03 | 7 |
| BAZ2A | 8.8E-01 | 7.6E-04 | -2.7E-01 | 6.4E-01 | -1.2E+00 | 2.2E-05 | 3 |

| | | | | | | | |
|---|---|---|---|---|---|---|---|
| BCLAF1 | 2.6E-01 | 5.4E-01 | -4.2E-01 | 3.1E-01 | -6.8E-01 | 4.2E-02 | 3 |
| SRGAP1 | 5.3E-01 | 2.2E-01 | -3.9E-01 | 6.0E-01 | -9.2E-01 | 3.7E-02 | 3 |
| PPP1R10 | 8.6E-01 | 1.6E-03 | -1.8E+00 | 2.3E-10 | -2.7E+00 | 2.2E-26 | 3 |
| SRFBP1 | 8.4E-01 | 6.4E-02 | -3.2E-01 | 8.3E-01 | -1.2E+00 | 2.1E-02 | 3 |
| LMNB1 | 8.4E-01 | 1.1E-02 | -2.4E-01 | 8.2E-01 | -1.1E+00 | 2.5E-03 | 3 |
| SLC12A2 | 8.1E-01 | 3.1E-02 | -1.3E-01 | 1.0E+00 | -9.4E-01 | 2.6E-02 | 3 |
| DCBLD1 | 1.0E+00 | 6.5E-02 | -3.6E-01 | 8.6E-01 | -1.4E+00 | 2.6E-02 | 3 |
| TOB2 | 1.1E+00 | 9.9E-02 | -4.7E-01 | 8.1E-01 | -1.5E+00 | 3.6E-02 | 3 |
| ZCCHC6 | 3.7E-01 | 3.4E-01 | -3.5E-01 | 5.2E-01 | -7.2E-01 | 4.9E-02 | 3 |
| PLEC | 1.3E+00 | 8.2E-03 | -4.0E-01 | 7.7E-01 | -1.7E+00 | 1.4E-03 | 3 |
| UBAP2 | 4.0E-01 | 3.7E-01 | -4.9E-01 | 3.2E-01 | -8.9E-01 | 2.2E-02 | 3 |
| VCP | 4.1E-01 | 4.0E-01 | -6.3E-01 | 1.9E-01 | -1.0E+00 | 9.8E-03 | 3 |
| MSH6 | 2.6E-01 | 6.6E-01 | -8.2E-01 | 4.5E-02 | -1.1E+00 | 4.2E-03 | 3 |
| FBXO11 | 8.9E-01 | 1.5E-02 | -1.1E-01 | 1.0E+00 | -1.0E+00 | 1.6E-02 | 3 |
| NOD1 | 1.4E+00 | 1.1E-02 | -3.1E-02 | 1.0E+00 | -1.4E+00 | 2.5E-02 | 3 |
| ETAA1 | 1.2E+00 | 9.3E-03 | -1.9E-01 | 1.0E+00 | -1.4E+00 | 7.7E-03 | 3 |
| AKT1S1 | 5.2E-01 | 2.6E-02 | -2.2E-01 | 6.5E-01 | -7.4E-01 | 2.7E-03 | 3 |
| FAM120B | 7.0E-01 | 8.2E-02 | -3.7E-01 | 6.7E-01 | -1.1E+00 | 1.5E-02 | 3 |
| SOX9 | 7.3E-01 | 3.8E-02 | -3.6E-02 | 1.0E+00 | -7.6E-01 | 6.7E-02 | 3 |
| CCDC163P | 1.6E+00 | 6.7E-02 | -5.7E-01 | 8.7E-01 | -2.1E+00 | 3.3E-02 | 3 |
| PCF11 | 8.3E-01 | 5.9E-02 | -1.2E+00 | 2.5E-02 | -2.0E+00 | 8.2E-06 | 3 |
| CASKIN2 | 3.4E-01 | 4.3E-01 | -4.7E-01 | 3.3E-01 | -8.1E-01 | 3.0E-02 | 3 |
| PPM1G | 5.5E-01 | 1.5E-01 | -2.9E-01 | 7.4E-01 | -8.4E-01 | 4.0E-02 | 3 |
| WDR37 | 5.5E-01 | 3.5E-02 | -2.7E-01 | 5.7E-01 | -8.3E-01 | 2.5E-03 | 3 |
| CHD8 | 1.3E+00 | 4.5E-05 | -1.7E-01 | 9.7E-01 | -1.5E+00 | 1.2E-05 | 3 |
| KAT6A | 3.7E-01 | 2.3E-01 | -5.9E-01 | 6.5E-02 | -9.7E-01 | 4.2E-04 | 3 |
| NAV3 | 5.3E-01 | 6.4E-02 | -9.6E-01 | 9.4E-04 | -1.5E+00 | 7.2E-09 | 3 |
| FAM98A | 4.1E-01 | 1.7E-01 | -4.7E-01 | 1.7E-01 | -8.9E-01 | 1.4E-03 | 3 |
| ZMAT2 | 1.4E+00 | 2.9E-03 | -1.1E-01 | 1.0E+00 | -1.5E+00 | 4.4E-03 | 3 |
| CUL4A | 1.1E+00 | 4.1E-02 | -1.1E-01 | 1.0E+00 | -1.2E+00 | 5.6E-02 | 3 |
| PDS5B | 3.8E-01 | 6.1E-01 | -9.6E-01 | 1.1E-01 | -1.3E+00 | 1.2E-02 | 3 |
| USP44 | 9.7E-01 | 6.2E-01 | -3.4E+00 | 1.1E-01 | -4.4E+00 | 2.8E-02 | 3 |
| CD44 | 6.9E-01 | 7.3E-02 | -2.2E-01 | 8.8E-01 | -9.1E-01 | 3.1E-02 | 3 |
| CHD7 | 7.4E-01 | 2.5E-03 | -2.7E-01 | 5.8E-01 | -1.0E+00 | 7.1E-05 | 3 |
| ASXL1 | 5.5E-01 | 1.2E-02 | -4.2E-01 | 1.4E-01 | -9.6E-01 | 8.2E-06 | 3 |
| CKAP2L | 6.9E-01 | 7.3E-01 | -2.6E+00 | 9.5E-02 | -3.3E+00 | 2.6E-02 | 3 |
| MAN2A1 | 4.8E-01 | 8.8E-02 | -1.9E-01 | 7.9E-01 | -6.7E-01 | 2.4E-02 | 3 |
| PIK3R1 | 4.3E-01 | 1.4E-01 | -2.8E-01 | 6.1E-01 | -7.1E-01 | 2.1E-02 | 3 |
| ZNF827 | 7.4E-01 | 5.8E-02 | -3.0E-01 | 7.7E-01 | -1.0E+00 | 1.5E-02 | 3 |
| ZNF827 | 7.4E-01 | 5.8E-02 | -3.0E-01 | 7.7E-01 | -1.0E+00 | 1.5E-02 | 3 |
| RNF34 | 1.4E+00 | 7.0E-02 | -5.0E-01 | 8.8E-01 | -1.9E+00 | 3.5E-02 | 3 |
| AAR2 | 4.0E-01 | 9.3E-02 | -2.1E-01 | 6.5E-01 | -6.1E-01 | 1.4E-02 | 3 |
| ATXN2 | 4.8E-01 | 3.5E-01 | -5.0E-01 | 4.4E-01 | -9.8E-01 | 3.5E-02 | 3 |
| TTI1 | 6.2E-01 | 2.5E-01 | -6.0E-01 | 4.3E-01 | -1.2E+00 | 2.1E-02 | 3 |
| RPRD1B | 1.1E+00 | 2.3E-02 | -3.1E-01 | 8.8E-01 | -1.4E+00 | 1.0E-02 | 3 |
| SRSF6 | 9.9E-01 | 7.0E-02 | -3.0E-01 | 9.2E-01 | -1.3E+00 | 3.6E-02 | 3 |
| PHF21A | 8.6E-01 | 9.9E-02 | -3.9E-01 | 7.8E-01 | -1.3E+00 | 3.0E-02 | 3 |
| VPS18 | 1.3E+00 | 2.1E-02 | -1.9E-01 | 1.0E+00 | -1.5E+00 | 2.2E-02 | 3 |
| RREB1 | 3.1E-01 | 1.8E-01 | -5.9E-01 | 6.7E-03 | -8.9E-01 | 4.5E-06 | 3 |
| ATP2B4 | 1.4E+00 | 5.4E-04 | -3.3E-02 | 1.0E+00 | -1.4E+00 | 1.3E-03 | 3 |
| CHD9 | 4.8E-01 | 1.3E-01 | -2.3E-01 | 7.5E-01 | -7.2E-01 | 3.4E-02 | 3 |
| SNX21 | 1.6E-01 | 8.4E-01 | -5.5E-01 | 1.4E-01 | -7.2E-01 | 3.6E-02 | 3 |
| NAV2 | 4.1E-01 | 1.5E-01 | -7.9E-01 | 4.0E-03 | -1.2E+00 | 9.3E-07 | 3 |
| MGEA5 | 3.9E-01 | 4.4E-01 | -5.6E-01 | 3.2E-01 | -9.5E-01 | 3.0E-02 | 3 |
| N4BP1 | 3.8E-01 | 2.1E-01 | -7.2E-01 | 1.5E-02 | -1.1E+00 | 2.1E-05 | 3 |
| BRWD1 | 7.9E-01 | 8.8E-02 | -3.6E-01 | 7.5E-01 | -1.2E+00 | 2.2E-02 | 3 |
| MX2 | 1.0E+00 | 1.7E-01 | -1.2E+00 | 1.4E-01 | -2.2E+00 | 9.0E-04 | 3 |
| SLC41A1 | 3.3E-01 | 5.2E-01 | -6.9E-01 | 1.0E-01 | -1.0E+00 | 6.9E-03 | 3 |
| PRDM15 | 4.0E-01 | 4.4E-01 | -1.1E+00 | 2.1E-02 | -1.5E+00 | 4.3E-04 | 3 |
| MLEC | 8.8E-01 | 8.6E-03 | -1.1E-01 | 1.0E+00 | -9.9E-01 | 9.0E-03 | 3 |
| S100PBP | 1.2E+00 | 5.7E-04 | -1.5E-01 | 1.0E+00 | -1.3E+00 | 3.2E-04 | 3 |
| KIAA1551 | 3.3E-01 | 6.7E-01 | -9.0E-01 | 7.7E-02 | -1.2E+00 | 8.7E-03 | 3 |
| PSMD1 | 3.8E-01 | 2.3E-01 | -2.7E-01 | 5.9E-01 | -6.6E-01 | 3.6E-02 | 3 |
| B3GNT7 | 1.1E+00 | 9.0E-02 | -5.2E-01 | 7.3E-01 | -1.6E+00 | 2.2E-02 | 3 |
| AKAP13 | 6.7E-01 | 4.6E-03 | -1.8E-01 | 8.2E-01 | -8.4E-01 | 8.5E-04 | 3 |
| GIGYF2 | 7.7E-01 | 2.6E-03 | -6.7E-01 | 3.4E-02 | -1.4E+00 | 1.3E-08 | 3 |
| PTPN1 | 9.7E-01 | 4.9E-03 | -2.3E-01 | 8.7E-01 | -1.2E+00 | 1.4E-03 | 3 |
| MID1 | 8.4E-01 | 3.8E-02 | -1.0E+00 | 4.5E-02 | -1.9E+00 | 9.0E-06 | 3 |
| OTUD7B | 1.1E+00 | 4.4E-02 | -5.4E-02 | 1.0E+00 | -1.1E+00 | 7.7E-02 | 3 |
| ZNRF3 | 1.6E+00 | 1.5E-01 | -1.1E+00 | 6.7E-01 | -2.6E+00 | 4.7E-02 | 3 |
| TFAM | 1.2E+00 | 2.6E-01 | -1.2E+00 | 5.0E-01 | -2.5E+00 | 4.7E-02 | 3 |
| AIM1L | 5.9E-01 | 2.6E-01 | -9.3E-01 | 7.4E-02 | -1.5E+00 | 6.6E-04 | 3 |
| PTPN14 | 9.1E-01 | 2.9E-01 | -1.3E+00 | 2.4E-01 | -2.2E+00 | 1.3E-02 | 3 |
| FTSJ3 | 1.1E+00 | 7.4E-03 | -5.7E-02 | 1.0E+00 | -1.1E+00 | 1.4E-02 | 3 |
| TFRC | 4.0E-01 | 3.2E-01 | -4.0E-01 | 4.5E-01 | -8.0E-01 | 3.2E-02 | 3 |
| RCAN3 | 3.1E-01 | 5.0E-01 | -4.5E-01 | 3.4E-01 | -7.6E-01 | 4.1E-02 | 3 |
| EPHB3 | 1.3E+00 | 1.7E-05 | -1.5E-01 | 9.7E-01 | -1.4E+00 | 5.0E-06 | 3 |
| ATF7IP | 2.6E-01 | 6.8E-01 | -6.3E-01 | 1.9E-01 | -8.9E-01 | 3.3E-02 | 3 |
| MFAP1 | 8.6E-01 | 3.6E-05 | -1.2E-01 | 9.5E-01 | -9.7E-01 | 8.2E-06 | 3 |
| CREB3L2 | 1.1E+00 | 3.3E-02 | -8.8E-02 | 1.0E+00 | -1.2E+00 | 5.1E-02 | 3 |
| ST6GAL1 | 1.2E+00 | 4.3E-02 | -1.8E-01 | 1.0E+00 | -1.4E+00 | 4.4E-02 | 3 |
| NMT1 | 8.7E-01 | 1.2E-03 | -9.0E-03 | 1.0E+00 | -8.8E-01 | 3.6E-03 | 3 |
| NODAL | 6.5E-01 | 4.9E-01 | -2.2E+00 | 2.9E-02 | -2.9E+00 | 2.9E-03 | 3 |
| PALD1 | 1.4E+00 | 4.9E-10 | -1.9E-02 | 1.0E+00 | -1.4E+00 | 3.5E-10 | 3 |
| DDX3X | 1.3E+00 | 2.5E-03 | -3.5E-01 | 8.1E-01 | -1.7E+00 | 4.6E-04 | 3 |
| UBTF | 2.2E-01 | 2.9E-01 | -4.1E-01 | 2.9E-02 | -6.3E-01 | 1.7E-04 | 3 |
| SUGP1 | 3.8E-01 | 3.5E-01 | -4.0E-01 | 4.4E-01 | -7.8E-01 | 3.6E-02 | 3 |
| MEPCE | 3.4E-01 | 2.5E-01 | -6.9E-01 | 1.5E-02 | -1.0E+00 | 3.1E-05 | 3 |
| MICU1 | 1.9E-01 | 6.8E-01 | -5.7E-01 | 7.1E-02 | -7.6E-01 | 8.4E-03 | 3 |
| FUNDC1 | 1.2E+00 | 1.3E-01 | -5.6E-01 | 7.9E-01 | -1.7E+00 | 4.5E-02 | 3 |
| TPI1 | 5.1E-01 | 8.7E-02 | -1.6E-01 | 8.9E-01 | -6.7E-01 | 3.7E-02 | 3 |
| MARK2 | 4.8E-01 | 7.5E-02 | -6.5E-01 | 2.9E-02 | -1.1E+00 | 8.2E-06 | 3 |
| NA | 4.2E-01 | 3.7E-01 | -4.4E-01 | 4.7E-01 | -8.7E-01 | 4.3E-02 | 3 |
| DLG5 | 5.3E-01 | 3.7E-01 | -6.4E-01 | 3.9E-01 | -1.2E+00 | 3.4E-02 | 3 |

| | | | | | | | |
|---|---|---|---|---|---|---|---|
| MINK | 9.9E-01 | 2.3E-02 | -1.5E-01 | 1.0E+00 | -1.1E+00 | 2.2E-02 | 3 |
| TNS4 | 8.5E-01 | 1.7E-04 | -9.9E-01 | 8.1E-05 | -1.8E+00 | 6.1E-18 | 3 |
| CASC3 | 2.7E-01 | 2.0E-01 | -6.9E-01 | 4.0E-04 | -9.6E-01 | 4.1E-08 | 3 |
| CASC3 | 2.7E-01 | 2.0E-01 | -6.9E-01 | 4.0E-04 | -9.6E-01 | 4.1E-08 | 3 |
| BCL2L13 | 9.3E-01 | 1.9E-03 | -1.2E-01 | 1.0E+00 | -1.1E+00 | 1.5E-03 | 3 |
| FAM120C | 1.2E+00 | 5.0E-02 | -5.0E-01 | 8.1E-01 | -1.7E+00 | 1.8E-02 | 3 |
| POLR2A | 5.9E-01 | 4.3E-02 | -6.0E-02 | 1.0E+00 | -6.6E-01 | 5.5E-02 | 3 |
| TARDBP | 1.3E+00 | 5.4E-02 | -2.8E-01 | 1.0E+00 | -1.6E+00 | 4.6E-02 | 3 |
| GPATCH4 | 3.1E-01 | 5.6E-01 | -5.2E-01 | 3.1E-01 | -8.3E-01 | 4.6E-02 | 3 |
| SPOP | 1.5E-01 | 7.1E-01 | -4.6E-01 | 6.0E-02 | -6.1E-01 | 7.5E-03 | 3 |
| ASH1L | 1.4E-01 | 7.0E-01 | -4.8E-01 | 3.5E-02 | -6.3E-01 | 3.1E-03 | 3 |
| WWC1 | 5.6E-01 | 2.4E-01 | -7.4E-01 | 1.6E-01 | -1.3E+00 | 2.4E-03 | 3 |
| IGDCC3 | 7.7E-01 | 5.3E-01 | -2.4E+00 | 1.0E-01 | -3.1E+00 | 2.0E-02 | 3 |
| GTF3C1 | 7.0E-01 | 2.0E-03 | -3.1E-01 | 4.1E-01 | -1.0E+00 | 1.3E-05 | 3 |
| SLC27A3 | 7.7E-01 | 3.3E-01 | -1.5E+00 | 1.3E-01 | -2.3E+00 | 8.5E-03 | 3 |
| LDLR | 4.8E-01 | 5.9E-01 | -1.3E+00 | 4.1E-02 | -1.8E+00 | 2.3E-03 | 3 |
| DDX24 | 2.1E-01 | 5.6E-01 | -4.5E-01 | 1.1E-01 | -6.6E-01 | 9.7E-03 | 3 |
| UBFD1 | 2.0E-01 | 6.7E-01 | -4.6E-01 | 1.9E-01 | -6.6E-01 | 3.0E-02 | 3 |
| LHX1 | 1.0E+00 | 4.8E-03 | -3.3E-01 | 6.9E-01 | -1.3E+00 | 3.8E-04 | 3 |
| RAVER1 | 7.8E-01 | 7.0E-03 | -2.8E-01 | 6.8E-01 | -1.1E+00 | 6.4E-04 | 3 |
| HNF1B | 1.3E+00 | 5.7E-09 | -3.6E-01 | 3.3E-01 | -1.6E+00 | 7.5E-14 | 3 |
| LRRC8E | 9.1E-01 | 7.7E-03 | -7.4E-01 | 1.2E-01 | -1.6E+00 | 4.9E-06 | 3 |
| JAG2 | 1.2E+00 | 1.2E-01 | -7.7E-01 | 6.1E-01 | -2.0E+00 | 1.8E-02 | 3 |
| MIER1 | 4.4E-01 | 4.0E-01 | -5.7E-01 | 3.3E-01 | -1.0E+00 | 2.7E-02 | 3 |
| NA | 9.1E-01 | 2.0E-01 | -7.3E-01 | 5.2E-01 | -1.6E+00 | 2.4E-02 | 3 |
| NLK | 6.5E-01 | 2.2E-02 | -3.9E-02 | 1.0E+00 | -6.8E-01 | 3.7E-02 | 3 |
| BCORL1 | 7.2E-01 | 1.9E-01 | -5.0E-01 | 6.0E-01 | -1.2E+00 | 3.1E-02 | 3 |
| SUPT6H | 6.0E-01 | 2.1E-01 | -4.7E-01 | 5.5E-01 | -1.1E+00 | 2.7E-02 | 3 |
| SAFB | 4.9E-01 | 5.4E-01 | -1.1E+00 | 1.3E-01 | -1.6E+00 | 1.5E-02 | 3 |
| SAFB2 | 3.2E-01 | 5.1E-01 | -4.6E-01 | 3.7E-01 | -7.8E-01 | 4.9E-02 | 3 |
| SNX29 | 9.2E-01 | 2.2E-01 | -1.3E+00 | 1.6E-01 | -2.3E+00 | 4.0E-03 | 3 |
| SNX29 | 9.2E-01 | 2.2E-01 | -1.3E+00 | 1.6E-01 | -2.3E+00 | 4.0E-03 | 3 |
| MYO18A | 1.4E+00 | 1.8E-02 | -3.6E-01 | 9.0E-01 | -1.7E+00 | 9.0E-03 | 3 |
| CXXC1 | 7.4E-01 | 1.6E-02 | -1.6E-02 | 1.0E+00 | -7.6E-01 | 3.6E-02 | 3 |
| PPL | 1.0E+00 | 1.3E-01 | -9.0E-01 | 4.5E-01 | -1.9E+00 | 1.3E-02 | 3 |
| ISG15 | 1.0E+00 | 1.1E-02 | -8.1E-01 | 9.7E-02 | -1.8E+00 | 3.0E-06 | 3 |
| TBL3 | 4.8E-01 | 1.6E-01 | -2.3E-01 | 7.8E-01 | -7.1E-01 | 4.9E-02 | 3 |
| MED16 | 6.4E-01 | 1.3E-02 | 8.0E-03 | 1.0E+00 | -6.4E-01 | 3.7E-02 | 3 |
| NUP214 | 1.4E+00 | 5.3E-03 | -4.4E-01 | 7.7E-01 | -1.8E+00 | 1.1E-03 | 3 |
| RFWD3 | 7.0E-01 | 8.9E-02 | -2.5E-01 | 8.7E-01 | -9.5E-01 | 3.6E-02 | 3 |
| MTF2 | 4.1E-01 | 1.4E-01 | -4.7E-01 | 1.4E-01 | -8.8E-01 | 6.4E-04 | 3 |
| AP1G1 | 1.1E+00 | 1.9E-02 | -5.9E-01 | 4.6E-01 | -1.7E+00 | 6.0E-04 | 3 |
| PRMT7 | 5.3E-01 | 2.1E-01 | -4.4E-01 | 4.9E-01 | -9.8E-01 | 2.1E-02 | 3 |
| NA | 8.7E-01 | 1.8E-01 | -6.4E-01 | 5.6E-01 | -1.5E+00 | 2.4E-02 | 3 |
| NA | 1.5E+00 | 2.7E-01 | -2.7E+00 | 2.7E-01 | -4.2E+00 | 3.6E-02 | 3 |
| MT-ND2 | 2.9E-01 | 5.3E-01 | -6.1E-01 | 9.8E-02 | -9.0E-01 | 6.5E-03 | 3 |
| NA | 2.3E+00 | 3.2E-03 | -9.0E-01 | 7.0E-01 | -3.2E+00 | 1.2E-03 | 3 |
| NA | 1.9E+00 | 1.7E-02 | -5.9E-01 | 8.8E-01 | -2.5E+00 | 1.0E-02 | 3 |
| NA | 7.4E-01 | 1.6E-01 | -4.3E-01 | 6.8E-01 | -1.2E+00 | 3.2E-02 | 3 |
| LEMD3 | 4.2E-01 | 2.5E-01 | -5.5E-01 | 1.7E-01 | -9.7E-01 | 3.0E-03 | 3 |
| GEMIN4 | 5.3E-01 | 7.1E-02 | -5.1E-01 | 1.8E-01 | -1.0E+00 | 3.3E-04 | 3 |
| ABCA8 | 7.8E-01 | 1.0E-01 | -3.4E-01 | 8.1E-01 | -1.1E+00 | 3.3E-02 | 3 |
| METTL14 | 5.3E-01 | 4.2E-01 | -7.9E-01 | 2.9E-01 | -1.3E+00 | 2.4E-02 | 3 |
| NA | 3.7E+00 | 2.3E-02 | 7.0E-01 | 1.0E+00 | -3.0E+00 | 8.2E-02 | 3 |
| NA | 9.7E-01 | 5.8E-01 | -3.9E+00 | 5.5E-02 | -4.8E+00 | 1.3E-02 | 3 |
| NA | 2.1E+00 | 1.6E-02 | -9.7E-01 | 6.1E-01 | -3.0E+00 | 2.2E-03 | 3 |
| ANKRD22 | 1.1E+00 | 4.2E-02 | -5.7E-02 | 1.0E+00 | -1.1E+00 | 7.2E-02 | 3 |
| BASP1 | 8.4E-01 | 2.0E-01 | -5.8E-01 | 6.4E-01 | -1.4E+00 | 4.1E-02 | 3 |
| NA | 2.4E+00 | 2.9E-02 | -1.2E-01 | 1.0E+00 | -2.5E+00 | 5.6E-02 | 3 |
| NACC2 | 5.1E-01 | 5.6E-02 | -1.6E-01 | 8.8E-01 | -6.7E-01 | 2.2E-02 | 3 |
| YIF1A | 2.0E-01 | 7.3E-01 | -5.3E-01 | 1.4E-01 | -7.2E-01 | 2.5E-02 | 3 |
| NA | 5.2E-01 | 1.9E-02 | -2.6E-01 | 5.1E-01 | -7.9E-01 | 7.3E-04 | 3 |
| NA | 6.5E-01 | 2.4E-01 | -7.5E-01 | 3.0E-01 | -1.4E+00 | 1.0E-02 | 3 |
| ZBTB47 | 8.9E-01 | 1.6E-02 | -7.6E-02 | 1.0E+00 | -9.6E-01 | 2.2E-02 | 3 |
| ZFP36L2 | 1.6E-01 | 7.4E-01 | -5.1E-01 | 6.6E-02 | -6.7E-01 | 9.8E-03 | 3 |
| GID4 | 2.1E-01 | 7.0E-01 | -6.4E-01 | 5.6E-02 | -8.4E-01 | 6.4E-03 | 3 |
| ZCCHC10 | 1.2E+00 | 2.6E-02 | -3.3E-01 | 8.8E-01 | -1.5E+00 | 1.0E-02 | 3 |
| NA | 8.1E-01 | 1.8E-01 | -5.6E-01 | 6.4E-01 | -1.4E+00 | 3.6E-02 | 3 |
| NA | 1.1E+00 | 2.6E-02 | -2.0E-01 | 1.0E+00 | -1.3E+00 | 2.3E-02 | 3 |
| BCL7A | 1.0E+00 | 1.9E-04 | -2.1E-01 | 8.4E-01 | -1.2E+00 | 2.1E-05 | 3 |
| LURAP1L | 5.6E-01 | 1.2E-01 | -1.2E+00 | 2.7E-04 | -1.8E+00 | 4.2E-09 | 3 |
| NA | 9.9E-01 | 2.6E-02 | -2.8E-01 | 8.8E-01 | -1.3E+00 | 1.1E-02 | 3 |
| NA | 9.9E-01 | 2.6E-02 | -2.8E-01 | 8.8E-01 | -1.3E+00 | 1.1E-02 | 3 |
| NA | 2.6E-01 | 7.1E-01 | -8.2E-01 | 5.1E-02 | -1.1E+00 | 5.8E-03 | 3 |
| NA | 2.3E+00 | 2.1E-02 | 1.1E-01 | 1.0E+00 | -2.2E+00 | 5.8E-02 | 3 |
| NA | 5.7E-01 | 3.8E-01 | -1.2E+00 | 5.0E-02 | -1.8E+00 | 1.2E-03 | 3 |
| RTTN | -7.2E-01 | 5.4E-02 | 1.5E-01 | 9.9E-01 | 8.7E-01 | 3.6E-02 | 2 |
| SEC11C | -8.2E-01 | 6.6E-04 | 3.6E-01 | 3.2E-01 | 1.2E+00 | 1.1E-06 | 2 |
| TXNL1 | -6.1E-01 | 7.7E-03 | -1.5E-01 | 8.5E-01 | 4.6E-01 | 1.3E-01 | 2 |
| RAB27B | -2.2E+00 | 9.5E-03 | 3.9E-02 | 1.0E+00 | 2.2E+00 | 2.2E-02 | 2 |
| STX7 | -6.9E-01 | 4.5E-02 | 3.1E-02 | 1.0E+00 | 7.2E-01 | 8.0E-02 | 2 |
| PIP4K2C | -8.1E-01 | 7.9E-03 | 2.7E-02 | 1.0E+00 | 8.4E-01 | 1.7E-02 | 2 |
| MAML1 | -1.2E+00 | 6.9E-03 | -3.7E-01 | 7.6E-01 | 8.7E-01 | 1.7E-01 | 2 |
| NA | -9.3E-01 | 5.7E-03 | -8.5E-02 | 1.0E+00 | 8.5E-01 | 3.6E-02 | 2 |
| PCMT1 | -5.9E-01 | 1.2E-02 | -1.1E-01 | 9.7E-01 | 4.8E-01 | 1.2E-01 | 2 |
| NA | -1.2E+00 | 4.0E-04 | -3.7E-01 | 5.7E-01 | 8.2E-01 | 6.0E-02 | 2 |
| SOD2 | -7.7E-01 | 3.9E-02 | -1.1E-01 | 1.0E+00 | 6.6E-01 | 1.9E-01 | 2 |
| SOD2 | -7.7E-01 | 3.9E-02 | -1.1E-01 | 1.0E+00 | 6.6E-01 | 1.9E-01 | 2 |
| ACAT2 | -1.3E+00 | 1.5E-04 | -4.0E-01 | 5.2E-01 | 8.6E-01 | 3.7E-02 | 2 |
| ISOC1 | -6.8E-01 | 2.9E-02 | 3.6E-02 | 1.0E+00 | 7.2E-01 | 4.9E-02 | 2 |
| SFT2D1 | -7.1E-01 | 2.6E-03 | -2.1E-02 | 1.0E+00 | 6.9E-01 | 1.2E-02 | 2 |
| MAN1A1 | -1.3E+00 | 4.6E-02 | -7.5E-02 | 1.0E+00 | 1.2E+00 | 1.4E-01 | 2 |
| SMDT1 | -7.9E-01 | 9.9E-04 | -6.2E-02 | 1.0E+00 | 7.3E-01 | 9.2E-03 | 2 |
| GOLM1 | -8.1E-01 | 1.2E-02 | -2.0E-01 | 8.7E-01 | 6.1E-01 | 1.7E-01 | 2 |

| | | | | | | | |
|---|---|---|---|---|---|---|---|
| NA | -6.5E-01 | 4.3E-02 | 2.3E-01 | 7.8E-01 | 8.9E-01 | 1.0E-02 | 2 |
| ZFAND3 | -8.5E-01 | 2.4E-03 | -1.7E-01 | 9.0E-01 | 6.8E-01 | 5.1E-02 | 2 |
| NA | -1.6E+00 | 1.6E-04 | -4.9E-01 | 5.3E-01 | 1.1E+00 | 3.9E-02 | 2 |
| NA | -1.6E+00 | 1.6E-04 | -4.9E-01 | 5.3E-01 | 1.1E+00 | 3.9E-02 | 2 |
| NA | -1.6E+00 | 1.6E-04 | -4.9E-01 | 5.3E-01 | 1.1E+00 | 3.9E-02 | 2 |
| NA | -1.6E+00 | 1.6E-04 | -4.9E-01 | 5.3E-01 | 1.1E+00 | 3.9E-02 | 2 |
| NA | -1.6E+00 | 1.6E-04 | -4.9E-01 | 5.3E-01 | 1.1E+00 | 3.9E-02 | 2 |
| NA | -1.6E+00 | 1.6E-04 | -4.9E-01 | 5.3E-01 | 1.1E+00 | 3.9E-02 | 2 |
| NA | -1.6E+00 | 1.6E-04 | -4.9E-01 | 5.3E-01 | 1.1E+00 | 3.9E-02 | 2 |
| NA | -1.6E+00 | 1.6E-04 | -4.9E-01 | 5.3E-01 | 1.1E+00 | 3.9E-02 | 2 |
| NA | -1.6E+00 | 1.6E-04 | -4.9E-01 | 5.3E-01 | 1.1E+00 | 3.9E-02 | 2 |
| NA | -1.6E+00 | 1.6E-04 | -4.9E-01 | 5.3E-01 | 1.1E+00 | 3.9E-02 | 2 |
| NA | -1.6E+00 | 1.6E-04 | -4.9E-01 | 5.3E-01 | 1.1E+00 | 3.9E-02 | 2 |
| NA | -1.6E+00 | 1.6E-04 | -4.9E-01 | 5.3E-01 | 1.1E+00 | 3.9E-02 | 2 |
| NA | -1.6E+00 | 1.6E-04 | -4.9E-01 | 5.3E-01 | 1.1E+00 | 3.9E-02 | 2 |
| NA | -1.6E+00 | 1.6E-04 | -4.9E-01 | 5.3E-01 | 1.1E+00 | 3.9E-02 | 2 |
| NA | -1.6E+00 | 1.6E-04 | -4.9E-01 | 5.3E-01 | 1.1E+00 | 3.9E-02 | 2 |
| NA | -1.6E+00 | 1.6E-04 | -4.9E-01 | 5.3E-01 | 1.1E+00 | 3.9E-02 | 2 |
| NA | -1.6E+00 | 1.6E-04 | -4.9E-01 | 5.3E-01 | 1.1E+00 | 3.9E-02 | 2 |
| NA | -1.6E+00 | 1.6E-04 | -4.9E-01 | 5.3E-01 | 1.1E+00 | 3.9E-02 | 2 |
| NA | -1.6E+00 | 1.6E-04 | -4.9E-01 | 5.3E-01 | 1.1E+00 | 3.9E-02 | 2 |
| NA | -1.6E+00 | 1.6E-04 | -4.9E-01 | 5.3E-01 | 1.1E+00 | 3.9E-02 | 2 |
| NA | -1.6E+00 | 1.6E-04 | -4.9E-01 | 5.3E-01 | 1.1E+00 | 3.9E-02 | 2 |
| NA | -1.6E+00 | 1.6E-04 | -4.9E-01 | 5.3E-01 | 1.1E+00 | 3.9E-02 | 2 |
| NA | -1.6E+00 | 1.6E-04 | -4.9E-01 | 5.3E-01 | 1.1E+00 | 3.9E-02 | 2 |
| NA | -1.6E+00 | 1.6E-04 | -4.9E-01 | 5.3E-01 | 1.1E+00 | 3.9E-02 | 2 |
| NA | -1.6E+00 | 1.6E-04 | -4.9E-01 | 5.3E-01 | 1.1E+00 | 3.9E-02 | 2 |
| NA | -1.6E+00 | 1.6E-04 | -4.9E-01 | 5.3E-01 | 1.1E+00 | 3.9E-02 | 2 |
| NA | -1.6E+00 | 1.6E-04 | -4.9E-01 | 5.3E-01 | 1.1E+00 | 3.9E-02 | 2 |
| NA | -1.6E+00 | 1.6E-04 | -4.9E-01 | 5.3E-01 | 1.1E+00 | 3.9E-02 | 2 |
| NA | -1.6E+00 | 1.6E-04 | -4.9E-01 | 5.3E-01 | 1.1E+00 | 3.9E-02 | 2 |
| NA | -1.6E+00 | 1.6E-04 | -4.9E-01 | 5.3E-01 | 1.1E+00 | 3.9E-02 | 2 |
| FAM32A | -1.2E+00 | 2.3E-02 | -1.2E-01 | 1.0E+00 | 1.1E+00 | 1.1E-01 | 2 |
| NA | -1.3E+00 | 2.0E-02 | -4.2E-01 | 7.7E-01 | 9.2E-01 | 3.0E-01 | 2 |
| ACN9 | -7.2E-01 | 2.9E-02 | 3.1E-02 | 1.0E+00 | 7.6E-01 | 5.4E-02 | 2 |
| ANKRD39 | -1.2E+00 | 2.6E-02 | -2.0E-01 | 1.0E+00 | 9.7E-01 | 1.7E-01 | 2 |
| TSPAN13 | -8.5E-01 | 3.4E-02 | -4.6E-02 | 1.0E+00 | 8.1E-01 | 1.1E-01 | 2 |
| NA | -9.5E-01 | 2.4E-03 | -9.3E-02 | 1.0E+00 | 8.6E-01 | 2.1E-02 | 2 |
| TEX10 | -9.4E-01 | 1.1E-04 | -1.5E-01 | 9.0E-01 | 7.9E-01 | 4.9E-03 | 2 |
| SMAP1 | -6.7E-01 | 2.6E-04 | 1.2E-01 | 8.8E-01 | 7.9E-01 | 3.5E-05 | 2 |
| TMEM30A | -7.3E-01 | 1.3E-02 | -5.5E-02 | 1.0E+00 | 6.7E-01 | 6.1E-02 | 2 |
| NA | -9.8E-01 | 2.5E-02 | 2.2E-01 | 9.3E-01 | 1.2E+00 | 1.2E-02 | 2 |
| CTNNAL1 | -1.3E+00 | 2.9E-03 | -2.3E-01 | 9.4E-01 | 1.0E+00 | 4.7E-02 | 2 |
| UTP3 | -8.8E-01 | 9.1E-03 | -1.9E-01 | 9.2E-01 | 6.9E-01 | 1.2E-01 | 2 |
| BBS9 | -1.5E+00 | 3.1E-04 | -3.3E-01 | 7.6E-01 | 1.2E+00 | 2.2E-02 | 2 |
| APLF | -7.1E-01 | 1.0E-02 | -1.5E-01 | 9.2E-01 | 5.5E-01 | 1.3E-01 | 2 |
| CAPZA2 | -7.6E-01 | 4.8E-05 | 9.1E-02 | 9.7E-01 | 8.6E-01 | 1.3E-05 | 2 |
| MRPL32 | -1.6E+00 | 5.3E-03 | -4.9E-01 | 7.4E-01 | 1.2E+00 | 1.5E-01 | 2 |
| STARD3NL | -8.0E-01 | 4.1E-04 | -2.3E-01 | 6.2E-01 | 5.6E-01 | 4.9E-02 | 2 |
| MATN3 | -4.7E+00 | 1.5E-02 | -1.0E+00 | 8.3E-01 | 3.6E+00 | 1.8E-01 | 2 |
| FIG4 | -7.3E-01 | 1.0E-02 | -1.4E-01 | 9.5E-01 | 5.9E-01 | 1.2E-01 | 2 |
| NA | -8.3E-01 | 8.6E-02 | 2.2E-01 | 9.5E-01 | 1.0E+00 | 5.0E-02 | 2 |
| SYPL1 | -1.1E+00 | 1.3E-05 | -3.1E-01 | 4.7E-01 | 7.4E-01 | 1.0E-02 | 2 |
| NT5DC1 | -1.3E+00 | 5.9E-03 | -3.6E-01 | 7.9E-01 | 9.4E-01 | 1.4E-01 | 2 |
| SLC36A4 | -9.2E-01 | 1.8E-04 | -2.3E-01 | 6.7E-01 | 6.8E-01 | 2.2E-02 | 2 |
| THNSL1 | -1.2E+00 | 3.7E-02 | -1.4E-01 | 1.0E+00 | 1.1E+00 | 1.6E-01 | 2 |
| APMAP | -6.7E-01 | 1.7E-01 | 3.4E-01 | 7.4E-01 | 1.0E+00 | 4.2E-02 | 2 |
| PICALM | -6.6E-01 | 5.5E-04 | -9.8E-02 | 9.6E-01 | 5.6E-01 | 1.3E-02 | 2 |
| PTER | -1.9E+00 | 4.2E-02 | -6.2E-01 | 7.9E-01 | 1.3E+00 | 4.2E-01 | 2 |
| VWA8 | -1.0E+00 | 2.7E-02 | 2.6E-01 | 8.8E-01 | 1.3E+00 | 9.8E-03 | 2 |
| IFT122 | -8.8E-01 | 1.4E-02 | 2.8E-01 | 7.5E-01 | 1.2E+00 | 2.1E-03 | 2 |
| IFT122 | -8.8E-01 | 1.4E-02 | 2.8E-01 | 7.5E-01 | 1.2E+00 | 2.1E-03 | 2 |
| NA | -1.0E+00 | 8.3E-02 | 2.8E-01 | 9.3E-01 | 1.3E+00 | 4.3E-02 | 2 |
| NA | -1.1E+00 | 4.9E-05 | -7.6E-02 | 1.0E+00 | 1.1E+00 | 5.8E-04 | 2 |
| CLN5 | -1.0E+00 | 8.9E-06 | -2.0E-01 | 7.4E-01 | 8.3E-01 | 1.6E-03 | 2 |
| NA | -1.1E+00 | 5.4E-03 | -3.0E-01 | 7.9E-01 | 8.3E-01 | 1.3E-01 | 2 |
| MED8 | -1.1E+00 | 3.2E-03 | 1.5E-01 | 1.0E+00 | 1.2E+00 | 2.0E-03 | 2 |
| UVRAG | -9.8E-01 | 7.9E-04 | -1.4E-01 | 9.6E-01 | 8.3E-01 | 1.7E-02 | 2 |
| RPS3 | -4.5E-01 | 2.8E-03 | 1.6E-01 | 5.8E-01 | 6.1E-01 | 7.7E-05 | 2 |
| THUMPD3 | -6.7E-01 | 3.7E-02 | 1.5E-01 | 9.5E-01 | 8.2E-01 | 1.9E-02 | 2 |
| NA | -8.6E-01 | 1.5E-04 | -2.2E-01 | 6.6E-01 | 6.4E-01 | 2.1E-02 | 2 |
| METTL3 | -7.9E-01 | 2.6E-03 | 9.6E-02 | 1.0E+00 | 8.8E-01 | 1.9E-03 | 2 |
| STAM2 | -7.1E-01 | 3.8E-02 | -1.3E-01 | 1.0E+00 | 5.8E-01 | 2.1E-01 | 2 |
| SUMF1 | -9.7E-01 | 1.9E-04 | -2.5E-01 | 6.8E-01 | 7.3E-01 | 2.2E-02 | 2 |
| NA | -1.1E+00 | 2.7E-04 | -1.6E-01 | 9.3E-01 | 9.0E-01 | 8.8E-03 | 2 |
| DUSP6 | -1.8E+00 | 3.4E-06 | -1.1E-01 | 1.0E+00 | 1.7E+00 | 3.8E-05 | 2 |
| QPCT | -1.2E+00 | 5.1E-05 | -3.2E-01 | 6.1E-01 | 8.9E-01 | 1.3E-02 | 2 |
| GTF2E2 | -8.8E-01 | 3.5E-02 | -8.7E-02 | 1.0E+00 | 8.0E-01 | 1.4E-01 | 2 |
| RBPMS | -7.3E-01 | 4.2E-02 | -6.5E-02 | 1.0E+00 | 6.6E-01 | 1.5E-01 | 2 |
| NEDD1 | -1.0E+00 | 6.1E-04 | -1.1E-01 | 1.0E+00 | 9.1E-01 | 9.4E-03 | 2 |
| USPL1 | -6.7E-01 | 2.4E-01 | 5.3E-01 | 5.0E-01 | 1.2E+00 | 2.4E-02 | 2 |
| GTF3A | -5.2E-01 | 5.5E-02 | 3.0E-01 | 5.1E-01 | 8.2E-01 | 3.0E-03 | 2 |
| NA | -1.1E+00 | 1.3E-02 | 1.3E-02 | 1.0E+00 | 1.1E+00 | 3.1E-02 | 2 |
| TTL | -3.6E-01 | 3.0E-01 | 3.2E-01 | 5.0E-01 | 6.8E-01 | 3.5E-02 | 2 |
| GGH | -6.5E-01 | 2.8E-03 | -1.7E-03 | 1.0E+00 | 6.5E-01 | 9.4E-03 | 2 |
| EIF3M | -9.2E-01 | 3.6E-02 | 2.4E-03 | 1.0E+00 | 9.2E-01 | 8.2E-02 | 2 |
| PEPD | -5.2E-01 | 5.2E-02 | 2.0E-01 | 7.7E-01 | 7.1E-01 | 1.2E-02 | 2 |
| NA | -1.1E+00 | 2.9E-03 | -1.6E-01 | 9.7E-01 | 9.1E-01 | 4.1E-02 | 2 |
| MRPL35 | -8.5E-01 | 6.7E-03 | -2.1E-01 | 8.6E-01 | 6.5E-01 | 1.2E-01 | 2 |
| DPY19L3 | -4.8E-01 | 8.2E-02 | 3.2E-01 | 4.3E-01 | 8.0E-01 | 3.3E-03 | 2 |
| CLCN3 | -6.5E-01 | 3.6E-02 | -7.5E-02 | 1.0E+00 | 5.8E-01 | 1.5E-01 | 2 |
| CCDC62 | -5.0E-01 | 2.2E-01 | 3.5E-01 | 5.9E-01 | 8.6E-01 | 3.1E-02 | 2 |
| NA | -1.2E+00 | 7.9E-04 | -2.9E-01 | 7.5E-01 | 8.7E-01 | 4.4E-02 | 2 |
| CCNH | -1.1E+00 | 1.2E-03 | -1.7E-01 | 9.4E-01 | 8.8E-01 | 2.6E-02 | 2 |

| | | | | | | | |
|---|---|---|---|---|---|---|---|
| USO1 | -5.1E-01 | 5.1E-02 | 4.2E-01 | 2.1E-01 | 9.3E-01 | 2.3E-04 | 2 |
| CCT8 | -3.8E-01 | 6.9E-02 | 3.0E-01 | 2.8E-01 | 6.8E-01 | 8.5E-04 | 2 |
| TMEM106C | -9.0E-01 | 2.1E-04 | -1.0E-01 | 1.0E+00 | 8.0E-01 | 4.0E-03 | 2 |
| CPOX | -1.1E+00 | 7.4E-04 | -1.0E-01 | 1.0E+00 | 9.5E-01 | 8.8E-03 | 2 |
| AP3B1 | -5.5E-01 | 1.5E-02 | 3.0E-01 | 4.0E-01 | 8.4E-01 | 2.2E-04 | 2 |
| DPY19L4 | -1.1E+00 | 2.6E-02 | -3.1E-01 | 8.8E-01 | 8.3E-01 | 2.6E-01 | 2 |
| PMS1 | -7.0E-01 | 1.8E-02 | 2.2E-01 | 7.8E-01 | 9.2E-01 | 3.5E-03 | 2 |
| TXNDC5 | -6.4E-01 | 3.4E-03 | 7.7E-03 | 1.0E+00 | 6.5E-01 | 9.9E-03 | 2 |
| PCNP | -8.0E-01 | 1.9E-03 | -1.8E-02 | 1.0E+00 | 7.8E-01 | 8.4E-03 | 2 |
| CD47 | -8.5E-01 | 7.5E-05 | -1.7E-01 | 7.8E-01 | 6.8E-01 | 7.0E-03 | 2 |
| NPM3 | -6.8E-01 | 2.9E-03 | -5.5E-02 | 1.0E+00 | 6.2E-01 | 2.1E-02 | 2 |
| PSMD14 | -7.0E-01 | 1.6E-03 | -6.7E-02 | 1.0E+00 | 6.4E-01 | 1.6E-02 | 2 |
| PSPH | -9.1E-01 | 7.7E-05 | 1.8E-01 | 8.0E-01 | 1.1E+00 | 4.9E-06 | 2 |
| APLP2 | -8.7E-01 | 2.7E-04 | -1.5E-01 | 9.0E-01 | 7.2E-01 | 1.1E-02 | 2 |
| ITFG1 | -3.5E-01 | 1.4E-01 | 2.9E-01 | 3.7E-01 | 6.4E-01 | 5.9E-03 | 2 |
| METTL15 | -5.9E-01 | 1.7E-01 | 4.1E-01 | 5.4E-01 | 1.0E+00 | 1.8E-02 | 2 |
| AMN1 | -1.0E+00 | 1.8E-05 | -1.2E-01 | 9.6E-01 | 8.9E-01 | 6.6E-04 | 2 |
| ASB7 | -9.9E-01 | 3.8E-03 | -2.9E-01 | 7.3E-01 | 7.1E-01 | 1.3E-01 | 2 |
| MANBA | -4.4E-01 | 1.2E-02 | 3.9E-01 | 5.8E-02 | 8.2E-01 | 8.5E-07 | 2 |
| SF3B1 | -5.7E-01 | 6.5E-02 | 1.2E-01 | 9.9E-01 | 6.9E-01 | 4.7E-02 | 2 |
| INTS12 | -8.9E-01 | 4.5E-02 | -2.1E-01 | 9.6E-01 | 6.8E-01 | 3.0E-01 | 2 |
| NPNT | -7.4E-01 | 1.1E-02 | -1.5E-01 | 9.5E-01 | 5.9E-01 | 1.2E-01 | 2 |
| MRPS35 | -7.5E-01 | 2.2E-02 | 3.2E-01 | 6.2E-01 | 1.1E+00 | 1.8E-03 | 2 |
| PAPSS1 | -7.0E-01 | 2.1E-03 | -4.9E-02 | 1.0E+00 | 6.5E-01 | 1.5E-02 | 2 |
| OSTC | -1.4E+00 | 4.3E-02 | 1.3E-01 | 1.0E+00 | 1.6E+00 | 5.7E-02 | 2 |
| OSTC | -1.4E+00 | 4.3E-02 | 1.3E-01 | 1.0E+00 | 1.6E+00 | 5.7E-02 | 2 |
| RBM34 | -7.6E-01 | 1.1E-02 | -1.6E-01 | 9.3E-01 | 6.0E-01 | 1.3E-01 | 2 |
| ETNK1 | -1.0E+00 | 8.6E-03 | -1.5E-01 | 1.0E+00 | 8.6E-01 | 7.9E-02 | 2 |
| RAB9A | -9.3E-01 | 2.0E-02 | -2.6E-01 | 8.5E-01 | 6.7E-01 | 2.5E-01 | 2 |
| ZFAND2A | -1.1E+00 | 2.4E-03 | -1.5E-01 | 9.9E-01 | 9.2E-01 | 3.1E-02 | 2 |
| HSCB | -1.1E+00 | 1.3E-03 | -2.3E-01 | 8.3E-01 | 8.3E-01 | 4.3E-02 | 2 |
| FAM204A | -7.5E-01 | 8.5E-03 | 6.7E-02 | 1.0E+00 | 8.2E-01 | 1.1E-02 | 2 |
| ABCB10 | -1.2E+00 | 4.4E-02 | -1.9E-01 | 1.0E+00 | 1.0E+00 | 2.1E-01 | 2 |
| RAB4A | -1.2E+00 | 1.3E-02 | 5.2E-03 | 1.0E+00 | 1.2E+00 | 3.3E-02 | 2 |
| RAB4A | -1.2E+00 | 1.3E-02 | 5.2E-03 | 1.0E+00 | 1.2E+00 | 3.3E-02 | 2 |
| PYROXD1 | -1.3E+00 | 2.6E-03 | 1.7E-02 | 1.0E+00 | 1.3E+00 | 7.6E-03 | 2 |
| WDR11 | -7.0E-01 | 2.2E-03 | 1.7E-01 | 8.1E-01 | 8.7E-01 | 3.2E-04 | 2 |
| ASCC2 | -6.0E-01 | 8.8E-03 | 2.7E-01 | 5.0E-01 | 8.7E-01 | 2.1E-04 | 2 |
| PSMA7 | -6.0E-01 | 1.6E-04 | 1.3E-01 | 7.7E-01 | 7.3E-01 | 8.5E-06 | 2 |
| NA | -5.1E-01 | 3.0E-03 | 1.2E-01 | 8.3E-01 | 6.3E-01 | 5.4E-04 | 2 |
| HEATR5A | -9.2E-01 | 3.9E-02 | -8.1E-02 | 1.0E+00 | 8.4E-01 | 1.4E-01 | 2 |
| FUCA1 | -5.5E-01 | 6.7E-02 | 1.5E-01 | 9.3E-01 | 7.0E-01 | 3.4E-02 | 2 |
| OLA1 | -6.9E-01 | 6.1E-03 | -2.2E-01 | 7.2E-01 | 4.8E-01 | 1.8E-01 | 2 |
| SMG7 | -7.2E-01 | 4.7E-03 | -1.9E-01 | 8.0E-01 | 5.3E-01 | 1.2E-01 | 2 |
| NA | -4.9E-01 | 8.9E-02 | 3.7E-01 | 3.5E-01 | 8.6E-01 | 2.4E-03 | 2 |
| PRDX4 | -7.7E-01 | 4.6E-07 | 7.7E-02 | 9.6E-01 | 8.5E-01 | 4.1E-08 | 2 |
| EFTUD1 | -1.0E+00 | 4.6E-03 | -1.5E-01 | 9.9E-01 | 8.9E-01 | 5.1E-02 | 2 |
| DYNLT3 | -6.1E-01 | 4.9E-03 | -1.4E-02 | 1.0E+00 | 6.0E-01 | 1.8E-02 | 2 |
| KLHDC2 | -6.8E-01 | 6.1E-03 | -2.2E-03 | 1.0E+00 | 6.8E-01 | 1.9E-02 | 2 |
| COMT | -8.2E-01 | 3.6E-02 | -1.2E-01 | 1.0E+00 | 7.0E-01 | 1.8E-01 | 2 |
| ATIC | -8.1E-01 | 7.5E-04 | 1.7E-01 | 8.5E-01 | 9.8E-01 | 1.1E-04 | 2 |
| NCKAP1 | -8.9E-01 | 2.2E-05 | -1.1E-01 | 9.6E-01 | 7.8E-01 | 7.6E-04 | 2 |
| NA | -4.4E-01 | 8.8E-02 | 1.5E-01 | 8.8E-01 | 5.9E-01 | 3.6E-02 | 2 |
| ERO1L | -7.9E-01 | 9.1E-03 | -6.6E-02 | 1.0E+00 | 7.2E-01 | 4.8E-02 | 2 |
| GORAB | -1.2E+00 | 1.8E-02 | 3.9E-01 | 7.7E-01 | 1.6E+00 | 3.3E-03 | 2 |
| ARRDC2 | -1.4E+00 | 2.3E-02 | -2.7E-01 | 9.7E-01 | 1.1E+00 | 1.8E-01 | 2 |
| QDPR | -1.0E+00 | 1.6E-03 | -5.3E-02 | 1.0E+00 | 9.5E-01 | 9.8E-03 | 2 |
| STAMBPL1 | -7.4E-01 | 7.7E-03 | 3.8E-02 | 1.0E+00 | 7.8E-01 | 1.5E-02 | 2 |
| SCCPDH | -1.3E+00 | 8.6E-03 | -2.8E-01 | 9.1E-01 | 1.0E+00 | 1.2E-01 | 2 |
| CHURC1 | -9.6E-01 | 4.4E-02 | -2.8E-01 | 8.8E-01 | 6.8E-01 | 3.6E-01 | 2 |
| NA | -8.4E-01 | 3.2E-02 | -1.5E-01 | 1.0E+00 | 6.9E-01 | 1.9E-01 | 2 |
| NA | -8.4E-01 | 3.2E-02 | -1.5E-01 | 1.0E+00 | 6.9E-01 | 1.9E-01 | 2 |
| NA | -8.4E-01 | 3.2E-02 | -1.5E-01 | 1.0E+00 | 6.9E-01 | 1.9E-01 | 2 |
| PRR14 | -4.4E-01 | 9.0E-02 | 1.8E-01 | 7.8E-01 | 6.2E-01 | 2.3E-02 | 2 |
| HEPH | -8.5E-01 | 2.7E-02 | -2.5E-01 | 8.5E-01 | 6.0E-01 | 2.9E-01 | 2 |
| TMEM107 | -1.6E+00 | 2.1E-02 | -3.4E-01 | 9.4E-01 | 1.2E+00 | 1.9E-01 | 2 |
| NA | -1.3E+00 | 1.5E-02 | -3.7E-01 | 8.1E-01 | 9.4E-01 | 2.3E-01 | 2 |
| TRIP4 | -9.1E-01 | 2.8E-02 | -1.7E-01 | 1.0E+00 | 7.5E-01 | 1.8E-01 | 2 |
| NUPR1 | -1.9E+00 | 1.6E-04 | -4.8E-01 | 7.0E-01 | 1.5E+00 | 1.8E-02 | 2 |
| SPATA7 | -9.4E-01 | 1.8E-02 | -1.7E-01 | 9.8E-01 | 7.7E-01 | 1.5E-01 | 2 |
| RAD51C | -5.1E-01 | 1.9E-01 | 3.0E-01 | 6.7E-01 | 8.1E-01 | 3.6E-02 | 2 |
| HDHD2 | -1.1E+00 | 2.6E-04 | -2.1E-01 | 8.5E-01 | 9.1E-01 | 1.3E-02 | 2 |
| TIMM8A | -9.8E-01 | 2.3E-02 | -2.8E-01 | 8.5E-01 | 7.0E-01 | 2.8E-01 | 2 |
| NA | -4.9E-01 | 7.9E-02 | 2.7E-01 | 5.8E-01 | 7.5E-01 | 8.0E-03 | 2 |
| ACSL4 | -1.2E+00 | 1.7E-04 | -3.6E-01 | 5.5E-01 | 8.0E-01 | 3.6E-02 | 2 |
| ANKRD29 | -6.8E-01 | 1.9E-02 | -9.8E-02 | 1.0E+00 | 5.8E-01 | 1.2E-01 | 2 |
| MARK3 | -1.1E+00 | 8.9E-04 | -2.8E-01 | 7.5E-01 | 8.5E-01 | 4.6E-02 | 2 |
| COPS3 | -5.2E-01 | 6.0E-02 | 3.3E-01 | 4.3E-01 | 8.5E-01 | 2.1E-03 | 2 |
| MRPS30 | -9.0E-01 | 1.3E-04 | 2.1E-01 | 7.2E-01 | 1.1E+00 | 4.9E-06 | 2 |
| WSB1 | -8.7E-01 | 4.2E-03 | -2.1E-01 | 8.3E-01 | 6.6E-01 | 1.0E-01 | 2 |
| BFAR | -5.7E-01 | 4.1E-03 | 3.2E-02 | 1.0E+00 | 6.1E-01 | 7.6E-03 | 2 |
| PARN | -8.6E-01 | 1.4E-03 | -1.5E-01 | 9.3E-01 | 7.1E-01 | 3.0E-02 | 2 |
| BRIX1 | -9.3E-01 | 6.5E-03 | -3.6E-02 | 1.0E+00 | 8.9E-01 | 2.6E-02 | 2 |
| MPPE1 | -7.6E-01 | 3.3E-02 | 1.5E-01 | 9.8E-01 | 9.1E-01 | 2.1E-02 | 2 |
| ACAA2 | -7.6E-01 | 7.7E-05 | -1.5E-01 | 8.1E-01 | 6.1E-01 | 6.3E-03 | 2 |
| CETN2 | -4.2E-01 | 1.9E-01 | 3.7E-01 | 3.8E-01 | 7.9E-01 | 1.0E-02 | 2 |
| ZNF185 | -8.7E-01 | 4.7E-02 | -5.2E-02 | 1.0E+00 | 8.2E-01 | 1.4E-01 | 2 |
| EMC6 | -4.8E-01 | 1.2E-01 | 1.9E-01 | 8.3E-01 | 6.7E-01 | 4.1E-02 | 2 |
| CAMKK1 | -8.5E-01 | 7.2E-03 | -2.2E-01 | 8.3E-01 | 6.3E-01 | 1.4E-01 | 2 |
| NA | -1.2E+00 | 1.1E-04 | -3.1E-01 | 6.3E-01 | 8.6E-01 | 1.9E-02 | 2 |
| GPR153 | -1.4E+00 | 2.8E-03 | -2.2E-01 | 9.6E-01 | 1.1E+00 | 4.2E-02 | 2 |
| PER3 | -1.6E+00 | 2.4E-02 | 1.7E-01 | 1.0E+00 | 1.7E+00 | 2.7E-02 | 2 |
| H6PD | -9.0E-01 | 2.3E-03 | -3.0E-01 | 6.3E-01 | 6.0E-01 | 1.3E-01 | 2 |
| STXBP3 | -5.3E-01 | 8.1E-02 | 4.1E-01 | 3.2E-01 | 9.4E-01 | 1.6E-03 | 2 |

| | | | | | | | |
|---|---|---|---|---|---|---|---|
| SH3GLB1 | -7.9E-01 | 1.1E-02 | -1.6E-01 | 9.5E-01 | 6.4E-01 | 1.2E-01 | 2 |
| NAE1 | -6.7E-01 | 9.8E-03 | -1.9E-01 | 8.0E-01 | 4.8E-01 | 1.9E-01 | 2 |
| ANKRD13C | -6.1E-01 | 7.9E-03 | -1.5E-01 | 8.6E-01 | 4.6E-01 | 1.3E-01 | 2 |
| DUSP2 | -1.7E+00 | 9.5E-05 | -8.9E-03 | 1.0E+00 | 1.7E+00 | 3.4E-04 | 2 |
| NA | -1.2E+00 | 3.0E-02 | 9.8E-02 | 1.0E+00 | 1.3E+00 | 4.1E-02 | 2 |
| PTRHD1 | -1.0E+00 | 4.6E-03 | -9.8E-02 | 1.0E+00 | 9.4E-01 | 3.3E-02 | 2 |
| PHOSPHO2 | -8.5E-01 | 9.2E-04 | 3.1E-01 | 4.9E-01 | 1.2E+00 | 8.2E-06 | 2 |
| NA | -2.0E+00 | 9.7E-03 | -5.7E-01 | 7.7E-01 | 1.4E+00 | 2.0E-01 | 2 |
| NA | -1.7E+00 | 1.3E-01 | 1.1E+00 | 3.9E-01 | 2.8E+00 | 7.7E-03 | 2 |
| IGJ | -6.9E-01 | 2.4E-02 | 4.4E-01 | 3.0E-01 | 1.1E+00 | 1.8E-04 | 2 |
| TMPRSS11E | -8.4E-01 | 1.5E-02 | -1.1E-01 | 1.0E+00 | 7.3E-01 | 1.0E-01 | 2 |
| NDUFC1 | -1.9E+00 | 1.5E-02 | -3.8E-01 | 9.5E-01 | 1.6E+00 | 1.5E-01 | 2 |
| TBCA | -8.3E-01 | 4.5E-04 | 5.6E-02 | 1.0E+00 | 8.9E-01 | 5.5E-04 | 2 |
| UBE2V2 | -5.8E-01 | 1.7E-02 | 9.2E-02 | 1.0E+00 | 6.7E-01 | 1.4E-02 | 2 |
| NA | -8.4E-01 | 2.4E-02 | 1.0E-01 | 1.0E+00 | 9.4E-01 | 2.6E-02 | 2 |
| MTPN | -4.5E-01 | 7.8E-03 | 1.9E-01 | 5.3E-01 | 6.4E-01 | 2.2E-04 | 2 |
| MTPN | -4.5E-01 | 7.8E-03 | 1.9E-01 | 5.3E-01 | 6.4E-01 | 2.2E-04 | 2 |
| UCHL3 | -7.5E-01 | 1.1E-02 | 9.9E-02 | 1.0E+00 | 8.5E-01 | 1.0E-02 | 2 |
| MLX | -8.3E-01 | 8.5E-03 | -1.8E-01 | 9.1E-01 | 6.5E-01 | 1.2E-01 | 2 |
| ZNF25 | -1.0E+00 | 1.8E-02 | -1.9E-01 | 9.7E-01 | 8.1E-01 | 1.5E-01 | 2 |
| FOXN2 | -1.2E+00 | 6.3E-04 | 4.0E-01 | 4.7E-01 | 1.6E+00 | 4.9E-06 | 2 |
| SAAL1 | -9.0E-01 | 4.8E-04 | 1.7E-02 | 1.0E+00 | 9.2E-01 | 1.2E-03 | 2 |
| NA | -5.5E-01 | 2.5E-03 | 1.1E-01 | 9.1E-01 | 6.6E-01 | 7.3E-04 | 2 |
| NA | -5.5E-01 | 2.5E-03 | 1.1E-01 | 9.1E-01 | 6.6E-01 | 7.3E-04 | 2 |
| NA | -1.4E+00 | 9.7E-04 | -2.9E-01 | 8.5E-01 | 1.1E+00 | 3.3E-02 | 2 |
| NA | -1.1E+00 | 1.1E-03 | -1.6E-01 | 9.7E-01 | 9.7E-01 | 2.0E-02 | 2 |
| NA | -1.1E+00 | 1.1E-03 | -1.6E-01 | 9.7E-01 | 9.7E-01 | 2.0E-02 | 2 |
| COMMD2 | -7.3E-01 | 5.2E-03 | -2.2E-01 | 7.2E-01 | 5.1E-01 | 1.6E-01 | 2 |
| NA | -1.2E+00 | 1.4E-02 | 1.8E-01 | 1.0E+00 | 1.4E+00 | 1.2E-02 | 2 |
| ZNF133 | -1.2E+00 | 3.4E-03 | -1.6E-01 | 1.0E+00 | 1.1E+00 | 3.7E-02 | 2 |
| GOLPH3L | -7.1E-01 | 4.7E-02 | -2.3E-01 | 8.5E-01 | 4.8E-01 | 4.0E-01 | 2 |
| RGS19 | -1.3E+00 | 1.5E-03 | -1.4E-01 | 1.0E+00 | 1.1E+00 | 1.6E-02 | 2 |
| ZNF250 | -5.2E-01 | 1.0E-01 | 2.9E-01 | 6.1E-01 | 8.1E-01 | 1.3E-02 | 2 |
| TMEM126A | -9.1E-01 | 1.1E-03 | -2.1E-01 | 8.1E-01 | 7.0E-01 | 4.3E-02 | 2 |
| EMP1 | -7.2E-01 | 9.1E-02 | 3.5E-01 | 6.9E-01 | 1.1E+00 | 1.7E-02 | 2 |
| COA1 | -9.6E-01 | 3.8E-02 | 9.6E-03 | 1.0E+00 | 9.7E-01 | 8.2E-02 | 2 |
| SERTAD4 | -1.1E+00 | 2.7E-02 | -1.8E-01 | 1.0E+00 | 8.8E-01 | 1.7E-01 | 2 |
| NA | -1.2E+00 | 2.3E-05 | -3.4E-01 | 5.3E-01 | 8.8E-01 | 1.1E-02 | 2 |
| TMEM66 | -4.5E-01 | 1.3E-02 | 1.6E-01 | 7.1E-01 | 6.0E-01 | 1.5E-03 | 2 |
| GABARAPL2 | -8.3E-01 | 4.2E-05 | -1.4E-01 | 8.4E-01 | 6.8E-01 | 3.1E-03 | 2 |
| CCDC115 | -5.0E-01 | 5.2E-02 | 1.5E-01 | 8.9E-01 | 6.5E-01 | 2.2E-02 | 2 |
| RNF125 | -1.0E+00 | 1.3E-01 | 4.4E-01 | 7.9E-01 | 1.5E+00 | 3.9E-02 | 2 |
| NA | -9.8E-01 | 8.4E-06 | -2.6E-01 | 5.5E-01 | 7.2E-01 | 4.7E-03 | 2 |
| NA | -6.4E-01 | 1.8E-02 | 1.6E-01 | 9.0E-01 | 7.9E-01 | 7.3E-03 | 2 |
| NA | -6.8E-01 | 5.9E-03 | -7.0E-02 | 1.0E+00 | 6.1E-01 | 4.1E-02 | 2 |
| METTL7A | -1.2E+00 | 1.1E-03 | 3.9E-02 | 1.0E+00 | 1.3E+00 | 2.5E-03 | 2 |
| MBD2 | -4.1E-01 | 1.4E-02 | 2.0E-01 | 5.0E-01 | 6.1E-01 | 4.0E-04 | 2 |
| NA | -6.7E-01 | 1.1E-01 | 2.4E-01 | 8.6E-01 | 9.1E-01 | 4.1E-02 | 2 |
| NA | -7.7E-01 | 2.7E-02 | -3.2E-02 | 1.0E+00 | 7.4E-01 | 8.5E-02 | 2 |
| NACA | 1.8E-01 | 9.8E-01 | 2.3E+00 | 1.0E-02 | 2.1E+00 | 1.1E-02 | 6 |
| MARS | -3.4E-01 | 3.1E-01 | 3.2E-01 | 4.6E-01 | 6.6E-01 | 3.1E-02 | 6 |
| MARS | -3.4E-01 | 3.1E-01 | 3.2E-01 | 4.6E-01 | 6.6E-01 | 3.1E-02 | 6 |
| TNFAIP3 | -4.6E-01 | 7.4E-01 | 2.8E+00 | 6.1E-06 | 3.3E+00 | 1.3E-08 | 6 |
| ABRACL | 3.3E-01 | 3.4E-01 | 1.0E+00 | 2.1E-04 | 6.8E-01 | 2.5E-02 | 6 |
| AIG1 | 7.5E-03 | 9.9E-01 | 7.7E-01 | 1.5E-02 | 7.6E-01 | 1.8E-02 | 6 |
| MTHFD1L | 1.7E-01 | 5.9E-01 | 6.0E-01 | 2.8E-03 | 4.3E-01 | 5.8E-02 | 6 |
| MTHFD1L | 1.7E-01 | 5.9E-01 | 6.0E-01 | 2.8E-03 | 4.3E-01 | 5.8E-02 | 6 |
| MTHFD1L | 1.7E-01 | 5.9E-01 | 6.0E-01 | 2.8E-03 | 4.3E-01 | 5.8E-02 | 6 |
| NIPAL2 | -1.3E-01 | 9.5E-01 | 7.4E-01 | 5.6E-02 | 8.7E-01 | 2.1E-02 | 6 |
| NA | 3.4E-01 | 3.8E-01 | 8.8E-01 | 3.9E-03 | 5.4E-01 | 1.3E-01 | 6 |
| SNX2 | -3.0E-01 | 3.3E-01 | 3.2E-01 | 3.6E-01 | 6.2E-01 | 2.1E-02 | 6 |
| PHAX | 6.4E-01 | 6.7E-02 | 1.4E+00 | 9.0E-06 | 8.0E-01 | 2.3E-02 | 6 |
| GPANK1 | 4.9E-01 | 8.1E-01 | 2.3E+00 | 4.1E-03 | 1.8E+00 | 1.7E-02 | 6 |
| FZD6 | 3.6E-01 | 5.0E-01 | 8.5E-01 | 4.0E-02 | 4.9E-01 | 3.9E-01 | 6 |
| NA | 1.1E+00 | 4.2E-01 | 2.4E+00 | 1.7E-02 | 1.3E+00 | 2.1E-01 | 6 |
| DCAF13 | 3.8E-01 | 3.0E-01 | 1.0E+00 | 5.2E-04 | 6.5E-01 | 4.3E-02 | 6 |
| UQCRQ | -1.8E-01 | 6.7E-01 | 4.9E-01 | 6.2E-02 | 6.7E-01 | 6.4E-03 | 6 |
| RPS10 | 1.1E-01 | 9.8E-01 | 9.6E-01 | 4.4E-02 | 8.5E-01 | 8.7E-02 | 6 |
| PTK2 | 1.9E-01 | 9.5E-01 | 1.5E+00 | 2.8E-03 | 1.3E+00 | 9.8E-03 | 6 |
| PTCH1 | -9.1E-02 | 9.8E-01 | 1.1E+00 | 3.1E-02 | 1.2E+00 | 1.6E-02 | 6 |
| FBP1 | 2.1E-01 | 7.5E-01 | 1.3E+00 | 2.1E-05 | 1.1E+00 | 2.6E-04 | 6 |
| ATF4 | 6.4E-01 | 2.1E-01 | 1.5E+00 | 9.0E-04 | 8.7E-01 | 1.3E-01 | 6 |
| AGTPBP1 | 2.8E-01 | 8.0E-01 | 1.5E+00 | 1.8E-03 | 1.2E+00 | 1.7E-02 | 6 |
| NA | 3.4E-01 | 5.0E-01 | 8.1E-01 | 3.9E-02 | 4.6E-01 | 4.0E-01 | 6 |
| PSAT1 | 5.3E-01 | 1.8E-01 | 1.3E+00 | 3.3E-04 | 7.3E-01 | 7.2E-02 | 6 |
| OARD1 | 9.5E-01 | 1.0E-01 | 2.0E+00 | 1.3E-04 | 1.1E+00 | 7.3E-02 | 6 |
| CTACK | 5.4E-01 | 5.2E-01 | 1.3E+00 | 4.6E-02 | 7.3E-01 | 4.1E-01 | 6 |
| SGCB | -2.5E-01 | 5.8E-01 | 8.8E-01 | 2.9E-03 | 1.1E+00 | 3.1E-05 | 6 |
| FIP1L1 | -4.5E-01 | 6.9E-02 | 5.7E-01 | 2.9E-02 | 1.0E+00 | 5.4E-06 | 6 |
| PLA2G7 | -3.1E-01 | 1.5E-01 | 3.7E-01 | 1.2E-01 | 6.8E-01 | 5.3E-04 | 6 |
| GNG11 | 2.0E-01 | 6.6E-01 | 7.2E-01 | 8.7E-03 | 5.2E-01 | 1.0E-01 | 6 |
| NA | -6.8E-01 | 6.3E-02 | 1.1E+00 | 8.2E-04 | 1.8E+00 | 2.6E-09 | 6 |
| COL15A1 | 2.9E-01 | 4.5E-01 | 8.8E-01 | 2.0E-03 | 5.9E-01 | 7.2E-02 | 6 |
| NA | 7.1E-02 | 9.7E-01 | 1.1E+00 | 1.9E-07 | 9.8E-01 | 5.6E-07 | 6 |
| ZNF331 | 2.7E-01 | 5.7E-01 | 1.0E+00 | 9.1E-04 | 7.6E-01 | 2.2E-02 | 6 |
| UBE3D | -5.9E-01 | 4.1E-01 | 8.3E-01 | 2.3E-01 | 1.4E+00 | 1.4E-02 | 6 |
| VRK2 | -2.1E-01 | 8.0E-01 | 6.5E-01 | 1.4E-01 | 8.6E-01 | 3.1E-02 | 6 |
| VRK2 | -2.1E-01 | 8.0E-01 | 6.5E-01 | 1.4E-01 | 8.6E-01 | 3.1E-02 | 6 |
| ME1 | -4.5E-01 | 5.2E-02 | 5.9E-01 | 1.6E-02 | 1.0E+00 | 9.8E-07 | 6 |
| CYB5R4 | -4.1E-01 | 3.3E-01 | 5.2E-01 | 2.3E-01 | 9.3E-01 | 8.4E-03 | 6 |
| FAM71E1 | 1.5E-01 | 9.8E-01 | 1.4E+00 | 1.5E-02 | 1.2E+00 | 3.9E-02 | 6 |
| SNX14 | -4.2E-01 | 4.0E-01 | 4.6E-01 | 4.2E-01 | 8.8E-01 | 3.9E-02 | 6 |
| GARS | 1.2E-01 | 8.6E-01 | 6.7E-01 | 4.5E-03 | 5.6E-01 | 3.0E-02 | 6 |
| RNGTT | 4.6E-01 | 1.7E-01 | 1.0E+00 | 5.6E-04 | 5.7E-01 | 1.0E-01 | 6 |

| | | | | | | | |
|---|---|---|---|---|---|---|---|
| ANKRD6 | -5.2E-02 | 9.8E-01 | 9.0E-01 | 6.6E-02 | 9.5E-01 | 4.9E-02 | 6 |
| RPS7 | 1.5E-01 | 5.1E-01 | 8.5E-01 | 2.1E-08 | 6.9E-01 | 4.5E-06 | 6 |
| IAH1 | -4.3E-01 | 4.1E-01 | 6.3E-01 | 2.0E-01 | 1.1E+00 | 1.0E-02 | 6 |
| TES | -2.7E-01 | 6.2E-02 | 3.2E-01 | 4.4E-02 | 5.9E-01 | 9.5E-06 | 6 |
| MTPAP | -2.8E-01 | 4.7E-01 | 4.7E-01 | 1.6E-01 | 7.4E-01 | 9.4E-03 | 6 |
| MTMR2 | 2.7E-01 | 5.3E-01 | 7.6E-01 | 1.2E-02 | 4.9E-01 | 1.7E-01 | 6 |
| PSMB1 | 2.3E-01 | 2.5E-01 | 7.8E-01 | 1.2E-06 | 5.5E-01 | 9.5E-04 | 6 |
| PRTFDC1 | -2.0E-01 | 7.9E-01 | 5.4E-01 | 1.9E-01 | 7.5E-01 | 4.6E-02 | 6 |
| COG1 | 4.0E-01 | 8.2E-02 | 8.9E-01 | 4.8E-05 | 4.8E-01 | 6.1E-02 | 6 |
| C17orf80 | 2.3E-01 | 6.3E-01 | 7.3E-01 | 1.8E-02 | 5.0E-01 | 1.8E-01 | 6 |
| TPT1 | -1.5E-01 | 7.3E-01 | 7.3E-01 | 7.4E-04 | 8.7E-01 | 1.5E-05 | 6 |
| TPT1 | -1.5E-01 | 7.3E-01 | 7.3E-01 | 7.4E-04 | 8.7E-01 | 1.5E-05 | 6 |
| NA | 2.8E-01 | 4.0E-01 | 1.0E+00 | 2.5E-05 | 7.4E-01 | 3.2E-03 | 6 |
| ICT1 | 2.2E-01 | 5.9E-01 | 9.3E-01 | 3.0E-04 | 7.1E-01 | 9.4E-03 | 6 |
| HN1 | 2.4E-01 | 5.8E-01 | 6.5E-01 | 3.1E-02 | 4.1E-01 | 3.0E-01 | 6 |
| NA | 2.3E-01 | 3.1E-01 | 6.6E-01 | 5.4E-04 | 4.2E-01 | 5.5E-02 | 6 |
| NA | 2.3E-01 | 3.1E-01 | 6.6E-01 | 5.4E-04 | 4.2E-01 | 5.5E-02 | 6 |
| NA | 2.3E-01 | 3.1E-01 | 6.6E-01 | 5.4E-04 | 4.2E-01 | 5.5E-02 | 6 |
| CDC123 | -2.1E-02 | 9.8E-01 | 9.6E-01 | 2.4E-02 | 9.8E-01 | 2.3E-02 | 6 |
| LRRFIP2 | 3.5E-01 | 1.6E-01 | 7.5E-01 | 9.9E-04 | 4.0E-01 | 1.7E-01 | 6 |
| WBP4 | -1.8E-01 | 6.0E-01 | 4.7E-01 | 5.3E-02 | 6.6E-01 | 3.2E-03 | 6 |
| NA | 6.9E-01 | 1.3E-01 | 1.7E+00 | 3.2E-05 | 1.0E+00 | 2.7E-02 | 6 |
| GDPD4 | 3.2E-01 | 7.9E-01 | 1.3E+00 | 1.4E-02 | 1.0E+00 | 9.2E-02 | 6 |
| R3HDM1 | -1.3E-01 | 8.9E-01 | 7.8E-01 | 8.0E-03 | 9.2E-01 | 1.0E-03 | 6 |
| NAPB | 1.4E-01 | 8.8E-01 | 7.3E-01 | 2.4E-02 | 5.8E-01 | 1.1E-01 | 6 |
| FOXA2 | 1.5E-01 | 6.2E-01 | 7.1E-01 | 1.6E-04 | 5.6E-01 | 5.2E-03 | 6 |
| SCEL | 1.1E-02 | 9.8E-01 | 8.3E-01 | 8.4E-04 | 8.2E-01 | 8.7E-04 | 6 |
| NA | 5.2E-02 | 9.8E-01 | 6.5E-01 | 7.0E-03 | 6.0E-01 | 1.7E-02 | 6 |
| ANKRD16 | -2.9E-01 | 2.3E-01 | 3.6E-01 | 1.7E-01 | 6.5E-01 | 2.2E-03 | 6 |
| PGS1 | 1.4E-01 | 8.6E-01 | 6.5E-01 | 3.6E-02 | 5.1E-01 | 1.6E-01 | 6 |
| AKR1E2 | 2.7E-01 | 7.0E-01 | 1.0E+00 | 8.0E-03 | 7.5E-01 | 8.2E-02 | 6 |
| NA | -2.6E-01 | 3.8E-01 | 4.7E-01 | 8.0E-02 | 7.3E-01 | 1.8E-03 | 6 |
| SETD5 | 3.5E-01 | 4.3E-01 | 1.4E+00 | 2.0E-05 | 1.0E+00 | 2.5E-03 | 6 |
| SNRPB2 | 1.3E-01 | 8.7E-01 | 1.1E+00 | 1.5E-05 | 9.6E-01 | 1.3E-04 | 6 |
| SGSH | 3.7E-01 | 4.3E-01 | 9.0E-01 | 1.5E-02 | 5.2E-01 | 2.6E-01 | 6 |
| MKKS | 6.4E-02 | 9.8E-01 | 7.5E-01 | 1.3E-02 | 6.9E-01 | 3.0E-02 | 6 |
| RPL15 | 1.4E-01 | 7.6E-01 | 7.5E-01 | 8.0E-04 | 6.1E-01 | 1.1E-02 | 6 |
| WHSC1L1 | 3.5E-01 | 2.5E-01 | 1.1E+00 | 8.7E-06 | 7.6E-01 | 3.9E-03 | 6 |
| CCDC41 | -2.2E-01 | 7.3E-01 | 5.9E-01 | 1.3E-01 | 8.1E-01 | 2.0E-02 | 6 |
| NDFIP1 | -2.4E-01 | 5.7E-01 | 5.2E-01 | 1.1E-01 | 7.7E-01 | 9.4E-03 | 6 |
| DDRGK1 | 3.0E-01 | 1.9E-01 | 7.0E-01 | 7.4E-04 | 4.0E-01 | 1.2E-01 | 6 |
| RPL31 | 5.0E-01 | 3.1E-01 | 1.2E+00 | 1.4E-03 | 7.5E-01 | 8.0E-02 | 6 |
| PUS1 | -2.6E-01 | 3.5E-01 | 6.0E-01 | 8.4E-03 | 8.5E-01 | 2.8E-05 | 6 |
| LTA4H | -2.7E-01 | 2.4E-01 | 4.2E-01 | 6.1E-02 | 7.0E-01 | 3.0E-04 | 6 |
| LTA4H | -2.7E-01 | 2.4E-01 | 4.2E-01 | 6.1E-02 | 7.0E-01 | 3.0E-04 | 6 |
| NA | 6.7E-02 | 9.8E-01 | 6.8E-01 | 3.9E-03 | 6.2E-01 | 1.0E-02 | 6 |
| MAPK8 | 1.7E-01 | 7.5E-01 | 6.3E-01 | 2.5E-02 | 4.6E-01 | 1.6E-01 | 6 |
| DPYSL3 | 2.8E-01 | 3.4E-01 | 9.5E-01 | 3.7E-05 | 6.7E-01 | 8.1E-03 | 6 |
| NA | 7.2E-01 | 3.4E-01 | 1.7E+00 | 6.3E-03 | 9.4E-01 | 1.7E-01 | 6 |
| UHRF1BP1L | 3.6E-01 | 7.9E-01 | 1.3E+00 | 3.5E-02 | 9.8E-01 | 1.8E-01 | 6 |
| IL6ST | -2.4E-01 | 7.0E-01 | 6.2E-01 | 1.2E-01 | 8.5E-01 | 1.8E-02 | 6 |
| ATP6V1H | -1.4E-01 | 7.6E-01 | 5.3E-01 | 2.7E-02 | 6.7E-01 | 3.0E-03 | 6 |
| IMMP1L | -3.0E-01 | 3.7E-01 | 6.5E-01 | 1.8E-02 | 9.5E-01 | 1.1E-04 | 6 |
| ADM | 6.5E-01 | 2.3E-01 | 1.8E+00 | 5.4E-05 | 1.2E+00 | 2.0E-02 | 6 |
| ACPL2 | 4.1E-01 | 4.0E-01 | 1.3E+00 | 4.2E-04 | 8.9E-01 | 2.6E-02 | 6 |
| IRF2 | 7.9E-01 | 1.5E-01 | 2.1E+00 | 5.3E-06 | 1.3E+00 | 3.1E-03 | 6 |
| CGGBP1 | -7.6E-02 | 9.8E-01 | 7.3E-01 | 3.5E-02 | 8.1E-01 | 1.8E-02 | 6 |
| KCMF1 | -2.3E-01 | 4.9E-01 | 6.6E-01 | 5.9E-03 | 8.9E-01 | 4.3E-05 | 6 |
| MSC | -5.8E-01 | 5.1E-01 | 8.2E-01 | 3.4E-01 | 1.4E+00 | 4.1E-02 | 6 |
| MTMR9 | -3.1E-02 | 9.8E-01 | 7.4E-01 | 4.3E-03 | 7.7E-01 | 2.5E-03 | 6 |
| SETD1B | 1.1E+00 | 2.4E-01 | 2.4E+00 | 2.8E-03 | 1.3E+00 | 1.5E-01 | 6 |
| NA | 6.1E-01 | 3.7E-01 | 1.7E+00 | 7.9E-04 | 1.1E+00 | 3.8E-02 | 6 |
| FAR1 | -2.7E-01 | 6.0E-01 | 7.5E-01 | 2.3E-02 | 1.0E+00 | 6.6E-04 | 6 |
| CDKL2 | 2.0E-01 | 7.3E-01 | 8.9E-01 | 3.8E-03 | 6.9E-01 | 3.9E-02 | 6 |
| TCTN1 | -2.1E-01 | 8.8E-01 | 8.8E-01 | 8.1E-02 | 1.1E+00 | 2.1E-02 | 6 |
| MYL9 | 4.5E-01 | 3.7E-01 | 1.0E+00 | 1.2E-02 | 6.0E-01 | 2.9E-01 | 6 |
| DGUOK | 3.8E-01 | 3.9E-01 | 1.1E+00 | 7.0E-04 | 7.6E-01 | 4.3E-02 | 6 |
| CPNE3 | -2.1E-01 | 3.6E-01 | 3.8E-01 | 7.4E-02 | 5.9E-01 | 1.4E-03 | 6 |
| DECR1 | -3.8E-01 | 3.2E-01 | 7.4E-01 | 2.6E-02 | 1.1E+00 | 1.7E-04 | 6 |
| TFG | -4.0E-02 | 9.8E-01 | 7.3E-01 | 2.7E-03 | 7.7E-01 | 1.3E-03 | 6 |
| DSP | -3.8E-02 | 9.8E-01 | 5.7E-01 | 7.6E-03 | 6.1E-01 | 4.0E-03 | 6 |
| CHID1 | 3.8E-01 | 3.2E-01 | 9.5E-01 | 2.6E-03 | 5.7E-01 | 1.5E-01 | 6 |
| ZBTB11 | -4.7E-02 | 9.8E-01 | 9.5E-01 | 5.8E-04 | 9.9E-01 | 1.4E-04 | 6 |
| ALCAM | -3.0E-01 | 2.1E-01 | 3.4E-01 | 2.2E-01 | 6.4E-01 | 3.4E-03 | 6 |
| TTC3 | 2.8E-01 | 3.9E-01 | 7.3E-01 | 6.3E-03 | 4.4E-01 | 1.9E-01 | 6 |
| TTC3 | 2.8E-01 | 3.9E-01 | 7.3E-01 | 6.3E-03 | 4.4E-01 | 1.9E-01 | 6 |
| PSMG1 | -2.6E-01 | 3.4E-01 | 3.9E-01 | 1.4E-01 | 6.5E-01 | 3.6E-03 | 6 |
| NA | -1.4E-01 | 8.7E-01 | 5.6E-01 | 7.1E-02 | 7.0E-01 | 1.9E-02 | 6 |
| PDLIM5 | -3.5E-01 | 1.5E-01 | 5.6E-01 | 2.4E-02 | 9.2E-01 | 2.1E-05 | 6 |
| SNRPN | 1.0E-01 | 9.6E-01 | 8.1E-01 | 7.1E-03 | 7.1E-01 | 2.6E-02 | 6 |
| SNRPN | 1.0E-01 | 9.6E-01 | 8.1E-01 | 7.1E-03 | 7.1E-01 | 2.6E-02 | 6 |
| CCDC34 | 2.8E-01 | 2.7E-01 | 6.5E-01 | 3.4E-03 | 3.7E-01 | 2.0E-01 | 6 |
| PHC2 | 7.9E-02 | 9.7E-01 | 6.2E-01 | 1.5E-02 | 5.4E-01 | 4.7E-02 | 6 |
| FAN1 | 2.0E-01 | 7.7E-01 | 7.1E-01 | 3.8E-02 | 5.2E-01 | 2.1E-01 | 6 |
| MRPL36 | 3.3E-01 | 1.5E-01 | 8.4E-01 | 5.5E-05 | 5.1E-01 | 3.3E-02 | 6 |
| SYNC | 1.4E-01 | 9.8E-01 | 1.0E+00 | 4.4E-02 | 8.7E-01 | 1.0E-01 | 6 |
| LRRK1 | -1.6E-01 | 7.2E-01 | 4.6E-01 | 8.5E-02 | 6.2E-01 | 1.2E-02 | 6 |
| RGS2 | -1.7E-01 | 9.8E-01 | 1.3E+00 | 3.5E-02 | 1.5E+00 | 1.6E-02 | 6 |
| TRIP12 | 3.4E-01 | 9.5E-02 | 8.5E-01 | 4.2E-06 | 5.2E-01 | 1.2E-02 | 6 |
| EIF3I | 1.6E-01 | 7.3E-01 | 6.3E-01 | 1.1E-02 | 4.7E-01 | 9.8E-02 | 6 |
| OXA1L | 2.4E-01 | 5.9E-01 | 6.5E-01 | 4.0E-02 | 4.1E-01 | 3.5E-01 | 6 |
| DAG1 | -1.3E-02 | 9.8E-01 | 7.1E-01 | 7.0E-05 | 7.2E-01 | 2.8E-05 | 6 |
| TBCE | 1.8E-01 | 6.8E-01 | 7.9E-01 | 9.8E-04 | 6.1E-01 | 1.8E-02 | 6 |
| CFI | -3.2E-01 | 5.6E-01 | 5.5E-01 | 2.3E-01 | 8.7E-01 | 2.6E-02 | 6 |

| | | | | | | | |
|---|---|---|---|---|---|---|---|
| SNAI1 | 3.7E-01 | 6.7E-01 | 1.8E+00 | 1.4E-04 | 1.5E+00 | 3.6E-03 | 6 |
| SSH3 | 1.8E-01 | 7.5E-01 | 6.8E-01 | 2.0E-02 | 5.0E-01 | 1.4E-01 | 6 |
| B3GNT5 | 1.4E-01 | 9.5E-01 | 1.0E+00 | 7.3E-03 | 8.8E-01 | 2.3E-02 | 6 |
| B3GNT5 | 1.4E-01 | 9.5E-01 | 1.0E+00 | 7.3E-03 | 8.8E-01 | 2.3E-02 | 6 |
| TBCEL | 2.0E-01 | 7.4E-01 | 6.7E-01 | 4.4E-02 | 4.7E-01 | 2.4E-01 | 6 |
| PCK2 | 1.6E-01 | 8.9E-01 | 8.4E-01 | 2.5E-02 | 6.8E-01 | 1.1E-01 | 6 |
| DALRD3 | -2.7E-01 | 7.6E-01 | 6.8E-01 | 1.9E-01 | 9.5E-01 | 3.6E-02 | 6 |
| PDIA5 | -3.1E-01 | 2.1E-01 | 3.1E-01 | 3.0E-01 | 6.2E-01 | 7.3E-03 | 6 |
| PDIA5 | -3.1E-01 | 2.1E-01 | 3.1E-01 | 3.0E-01 | 6.2E-01 | 7.3E-03 | 6 |
| CAMK2D | -3.6E-02 | 9.8E-01 | 6.0E-01 | 1.9E-03 | 6.3E-01 | 7.3E-04 | 6 |
| LPGAT1 | -1.4E-01 | 8.7E-01 | 5.5E-01 | 8.6E-02 | 6.9E-01 | 2.2E-02 | 6 |
| AP1S2 | 3.9E-01 | 7.1E-02 | 8.2E-01 | 5.5E-05 | 4.3E-01 | 7.9E-02 | 6 |
| PRMT2 | 2.8E-01 | 5.9E-01 | 1.1E+00 | 5.9E-04 | 8.3E-01 | 1.7E-02 | 6 |
| ATF3 | -5.9E-01 | 4.3E-01 | 1.3E+00 | 2.9E-02 | 1.9E+00 | 4.6E-04 | 6 |
| SCLY | 7.9E-02 | 9.8E-01 | 8.3E-01 | 1.4E-02 | 7.5E-01 | 3.2E-02 | 6 |
| CCDC173 | 9.8E-02 | 9.8E-01 | 1.0E+00 | 1.5E-02 | 9.3E-01 | 3.3E-02 | 6 |
| NA | -2.4E+00 | 3.8E-01 | 2.5E+00 | 2.7E-01 | 4.9E+00 | 2.3E-02 | 6 |
| TMEM25 | 2.2E-02 | 9.8E-01 | 9.0E-01 | 1.3E-02 | 8.8E-01 | 1.5E-02 | 6 |
| NA | 3.2E-01 | 3.7E-01 | 7.0E-01 | 2.1E-02 | 3.8E-01 | 3.9E-01 | 6 |
| RPS17 | -2.8E-01 | 2.3E-01 | 5.4E-01 | 1.3E-02 | 8.2E-01 | 2.1E-05 | 6 |
| RPS17 | -2.8E-01 | 2.3E-01 | 5.4E-01 | 1.3E-02 | 8.2E-01 | 2.1E-05 | 6 |
| UCKL1 | 2.6E-01 | 5.4E-01 | 7.2E-01 | 2.1E-02 | 4.6E-01 | 2.6E-01 | 6 |
| GLRX3 | 1.1E-01 | 8.6E-01 | 8.0E-01 | 6.6E-05 | 7.0E-01 | 6.4E-04 | 6 |
| IGF2BP2 | 1.9E-01 | 6.3E-01 | 6.4E-01 | 9.7E-03 | 4.5E-01 | 1.2E-01 | 6 |
| KIF9 | 3.7E-01 | 9.8E-01 | 4.0E+00 | 2.0E-03 | 3.6E+00 | 1.2E-03 | 6 |
| NFKBIA | -1.9E-01 | 7.4E-01 | 1.6E+00 | 4.0E-09 | 1.8E+00 | 5.4E-13 | 6 |
| DNAJB11 | -2.4E-01 | 7.2E-01 | 5.6E-01 | 2.0E-01 | 8.0E-01 | 3.9E-02 | 6 |
| OSBPL6 | 6.6E-01 | 3.6E-02 | 1.4E+00 | 2.1E-06 | 7.7E-01 | 2.4E-02 | 6 |
| TOR3A | -6.4E-03 | 9.9E-01 | 7.1E-01 | 2.2E-03 | 7.2E-01 | 1.8E-03 | 6 |
| CDCP1 | -1.6E-01 | 6.9E-01 | 5.2E-01 | 3.1E-02 | 6.8E-01 | 2.4E-03 | 6 |
| PCBD1 | 3.9E-01 | 4.8E-01 | 8.6E-01 | 4.4E-02 | 4.8E-01 | 4.4E-01 | 6 |
| SMARCB1 | 3.9E-01 | 1.5E-01 | 8.5E-01 | 5.9E-04 | 4.6E-01 | 1.4E-01 | 6 |
| NMD3 | 1.5E-01 | 8.9E-01 | 9.5E-01 | 3.6E-03 | 8.0E-01 | 1.8E-02 | 6 |
| MCU | 2.0E-01 | 6.1E-01 | 1.0E+00 | 1.5E-05 | 8.1E-01 | 6.6E-04 | 6 |
| TRMT44 | 8.0E-01 | 4.9E-01 | 1.9E+00 | 2.5E-02 | 1.1E+00 | 2.7E-01 | 6 |
| P4HA1 | -3.6E-01 | 4.3E-01 | 4.7E-01 | 3.3E-01 | 8.3E-01 | 2.9E-02 | 6 |
| DNAJC9 | 7.1E-01 | 2.3E-01 | 2.4E+00 | 1.3E-07 | 1.7E+00 | 3.1E-05 | 6 |
| FERMT2 | -4.1E-01 | 3.5E-01 | 6.8E-01 | 1.0E-01 | 1.1E+00 | 2.3E-03 | 6 |
| NCAPD2 | -3.4E-01 | 5.9E-01 | 9.4E-01 | 3.5E-02 | 1.3E+00 | 1.7E-03 | 6 |
| FNDC3B | 3.0E-01 | 6.9E-01 | 9.1E-01 | 4.2E-02 | 6.1E-01 | 2.6E-01 | 6 |
| FNDC3B | 3.0E-01 | 6.9E-01 | 9.1E-01 | 4.2E-02 | 6.1E-01 | 2.6E-01 | 6 |
| MAPK1 | -1.0E-01 | 9.1E-01 | 6.0E-01 | 1.2E-02 | 7.0E-01 | 2.3E-03 | 6 |
| NA | -4.3E-01 | 5.8E-02 | 5.1E-01 | 4.2E-02 | 9.4E-01 | 8.2E-06 | 6 |
| NA | -4.3E-01 | 5.8E-02 | 5.1E-01 | 4.2E-02 | 9.4E-01 | 8.2E-06 | 6 |
| DAGLB | 3.0E-01 | 3.4E-01 | 7.5E-01 | 4.2E-03 | 4.5E-01 | 1.7E-01 | 6 |
| NA | 5.2E-01 | 6.6E-01 | 1.7E+00 | 1.2E-02 | 1.2E+00 | 9.5E-02 | 6 |
| CDC42BPA | 1.6E-01 | 7.3E-01 | 6.3E-01 | 1.2E-02 | 4.7E-01 | 1.1E-01 | 6 |
| PARP1 | 4.6E-01 | 7.3E-02 | 1.1E+00 | 7.1E-06 | 6.2E-01 | 2.2E-02 | 6 |
| PLEKHM2 | 2.8E-01 | 7.2E-01 | 1.6E+00 | 4.6E-05 | 1.3E+00 | 7.6E-04 | 6 |
| WDFY1 | 3.4E-02 | 9.8E-01 | 3.8E+00 | 3.7E-07 | 3.8E+00 | 1.2E-08 | 6 |
| ETV3 | -6.8E-03 | 9.9E-01 | 2.3E+00 | 5.5E-09 | 2.4E+00 | 8.2E-11 | 6 |
| BNIP2 | 3.9E-01 | 7.0E-01 | 1.6E+00 | 2.0E-03 | 1.2E+00 | 1.8E-02 | 6 |
| NPC2 | -2.5E-01 | 3.9E-01 | 4.3E-01 | 1.1E-01 | 6.8E-01 | 3.6E-03 | 6 |
| FAM96A | 3.4E-01 | 3.9E-01 | 7.5E-01 | 2.5E-02 | 4.0E-01 | 4.1E-01 | 6 |
| SPG21 | 3.7E-01 | 3.3E-01 | 1.0E+00 | 6.2E-04 | 6.6E-01 | 5.1E-02 | 6 |
| RAB11A | -2.3E-01 | 4.2E-01 | 3.9E-01 | 1.2E-01 | 6.2E-01 | 5.2E-03 | 6 |
| XPO6 | -9.3E-02 | 9.5E-01 | 7.3E-01 | 3.8E-03 | 8.2E-01 | 6.3E-04 | 6 |
| SCPEP1 | -3.6E-01 | 3.7E-01 | 3.8E-01 | 4.3E-01 | 7.4E-01 | 3.7E-02 | 6 |
| ZC3H14 | -2.9E-01 | 5.8E-01 | 7.0E-01 | 5.0E-02 | 9.9E-01 | 2.3E-03 | 6 |
| S100A6 | 1.9E-01 | 6.1E-01 | 6.6E-01 | 5.5E-03 | 4.7E-01 | 8.9E-02 | 6 |
| UACA | 4.4E-01 | 3.1E-01 | 1.0E+00 | 5.9E-03 | 5.7E-01 | 2.4E-01 | 6 |
| NA | -1.0E-01 | 8.4E-01 | 5.2E-01 | 1.1E-02 | 6.2E-01 | 1.4E-03 | 6 |
| CENPV | 6.0E-01 | 3.4E-01 | 1.5E+00 | 4.1E-03 | 8.6E-01 | 1.5E-01 | 6 |
| FBXO22 | -2.0E-01 | 3.6E-01 | 4.1E-01 | 3.5E-02 | 6.1E-01 | 3.9E-04 | 6 |
| NA | -1.7E-01 | 7.9E-01 | 7.6E-01 | 9.3E-03 | 9.2E-01 | 6.6E-04 | 6 |
| IL13RA1 | 4.2E-01 | 4.1E-01 | 9.7E-01 | 1.8E-02 | 5.5E-01 | 3.0E-01 | 6 |
| MYO1D | 2.0E-01 | 6.1E-01 | 7.4E-01 | 3.6E-03 | 5.4E-01 | 6.3E-02 | 6 |
| METTL4 | -5.4E-01 | 3.1E-01 | 5.4E-01 | 3.8E-01 | 1.1E+00 | 2.1E-02 | 6 |
| MOCS2 | 7.3E-02 | 9.8E-01 | 6.8E-01 | 6.2E-03 | 6.1E-01 | 1.7E-02 | 6 |
| LPIN2 | 3.9E-01 | 4.7E-01 | 8.6E-01 | 4.7E-02 | 4.7E-01 | 4.7E-01 | 6 |
| RHOT1 | 1.9E-01 | 5.5E-01 | 6.5E-01 | 3.3E-03 | 4.6E-01 | 6.9E-02 | 6 |
| COPRS | -2.2E-01 | 6.8E-01 | 4.8E-01 | 1.9E-01 | 7.0E-01 | 3.0E-02 | 6 |
| CUL4B | -4.7E-01 | 3.3E-01 | 5.6E-01 | 2.9E-01 | 1.0E+00 | 1.5E-02 | 6 |
| ANKRD12 | -2.1E-01 | 6.5E-01 | 4.2E-01 | 2.6E-01 | 6.3E-01 | 4.4E-02 | 6 |
| PIGS | -1.3E-01 | 7.6E-01 | 5.9E-01 | 4.1E-03 | 7.2E-01 | 1.9E-04 | 6 |
| ATG4C | 2.3E-01 | 7.7E-01 | 8.5E-01 | 4.2E-02 | 6.1E-01 | 2.3E-01 | 6 |
| TNFRSF17 | -2.0E-02 | 9.8E-01 | 7.2E-01 | 2.5E-02 | 7.4E-01 | 2.0E-02 | 6 |
| ZFR | 2.6E-01 | 5.2E-01 | 6.1E-01 | 4.9E-02 | 3.5E-01 | 4.4E-01 | 6 |
| NUFIP2 | 7.1E-01 | 5.7E-01 | 1.9E+00 | 2.4E-02 | 1.2E+00 | 2.2E-01 | 6 |
| IDS | -2.1E-01 | 6.2E-01 | 5.1E-01 | 9.5E-02 | 7.3E-01 | 9.2E-03 | 6 |
| NA | 2.8E-01 | 2.6E-01 | 8.2E-01 | 6.7E-05 | 5.4E-01 | 1.8E-02 | 6 |
| DNAJA3 | -2.1E-01 | 4.3E-01 | 5.8E-01 | 4.6E-03 | 8.0E-01 | 2.0E-05 | 6 |
| GNB1 | 3.0E-01 | 2.8E-01 | 8.2E-01 | 3.4E-04 | 5.2E-01 | 4.7E-02 | 6 |
| PGP | 3.0E-01 | 4.3E-01 | 7.9E-01 | 8.5E-03 | 4.9E-01 | 2.0E-01 | 6 |
| CIRBP | -6.3E-01 | 3.6E-01 | 9.3E-01 | 1.3E-01 | 1.6E+00 | 3.5E-03 | 6 |
| LUC7L | 1.7E-02 | 9.8E-01 | 8.1E-01 | 5.2E-03 | 7.9E-01 | 6.7E-03 | 6 |
| WDR5 | 2.5E-01 | 3.8E-01 | 7.2E-01 | 1.4E-03 | 4.7E-01 | 7.7E-02 | 6 |
| GTF3C4 | -5.3E-02 | 9.8E-01 | 1.3E+00 | 2.1E-02 | 1.3E+00 | 1.3E-02 | 6 |
| C16orf95 | -8.3E-01 | 2.3E-01 | 7.7E-01 | 3.3E-01 | 1.6E+00 | 9.4E-03 | 6 |
| FAM102B | 1.1E-01 | 9.8E-01 | 1.2E+00 | 8.6E-03 | 1.1E+00 | 1.8E-02 | 6 |
| EXOSC2 | 2.5E-01 | 6.6E-01 | 8.3E-01 | 1.7E-02 | 5.8E-01 | 1.5E-01 | 6 |
| RTCA | -4.6E-02 | 9.8E-01 | 6.8E-01 | 2.5E-02 | 7.2E-01 | 1.7E-02 | 6 |
| PBX3 | 8.3E-02 | 9.8E-01 | 6.9E-01 | 5.0E-02 | 6.0E-01 | 1.2E-01 | 6 |
| NOB1 | 4.0E-01 | 1.7E-01 | 1.1E+00 | 2.5E-05 | 6.8E-01 | 1.7E-02 | 6 |

| | | | | | | | |
|---|---|---|---|---|---|---|---|
| DAB2IP | 2.7E-01 | 7.6E-01 | 1.2E+00 | 4.1E-03 | 9.5E-01 | 3.5E-02 | 6 |
| NA | -1.1E+00 | 4.1E-02 | 1.1E+00 | 6.1E-02 | 2.2E+00 | 8.5E-06 | 6 |
| NA | 5.3E-01 | 4.4E-01 | 1.6E+00 | 1.1E-03 | 1.1E+00 | 3.9E-02 | 6 |
| MT-ATP6 | 1.7E-01 | 5.3E-01 | 7.5E-01 | 1.4E-05 | 5.8E-01 | 1.1E-03 | 6 |
| UBB | 5.1E-01 | 5.8E-01 | 1.8E+00 | 2.2E-03 | 1.3E+00 | 3.6E-02 | 6 |
| RPS27A | 4.6E-01 | 1.2E-01 | 1.2E+00 | 7.3E-06 | 7.3E-01 | 1.4E-02 | 6 |
| SURF1 | -2.1E-01 | 5.5E-01 | 4.7E-01 | 7.3E-02 | 6.8E-01 | 4.0E-03 | 6 |
| MT2A | -3.4E-02 | 9.8E-01 | 7.4E-01 | 5.0E-02 | 7.8E-01 | 4.3E-02 | 6 |
| MT2A | -3.4E-02 | 9.8E-01 | 7.4E-01 | 5.0E-02 | 7.8E-01 | 4.3E-02 | 6 |
| ARGLU1 | 1.4E-01 | 8.5E-01 | 1.3E+00 | 2.4E-07 | 1.1E+00 | 3.0E-06 | 6 |
| NA | -5.0E-02 | 9.8E-01 | 8.0E-01 | 4.4E-05 | 8.5E-01 | 5.1E-06 | 6 |
| KLHL23 | 3.8E-01 | 7.1E-01 | 1.4E+00 | 1.6E-02 | 9.8E-01 | 1.4E-01 | 6 |
| KLHL23 | 3.8E-01 | 7.1E-01 | 1.4E+00 | 1.6E-02 | 9.8E-01 | 1.4E-01 | 6 |
| NA | 5.3E-01 | 2.0E-01 | 1.2E+00 | 6.8E-04 | 7.1E-01 | 1.1E-01 | 6 |
| NA | 3.0E-01 | 2.6E-01 | 7.5E-01 | 9.8E-04 | 4.5E-01 | 1.1E-01 | 6 |
| CD59 | 3.8E-01 | 2.4E-01 | 9.5E-01 | 5.1E-04 | 5.8E-01 | 7.4E-02 | 6 |
| NA | -4.2E-01 | 8.8E-01 | 2.3E+00 | 5.0E-03 | 2.7E+00 | 4.3E-04 | 6 |
| GEMIN8 | 1.6E-01 | 8.2E-01 | 7.9E-01 | 7.4E-03 | 6.3E-01 | 5.1E-02 | 6 |
| NA | 5.1E-01 | 5.3E-03 | 1.2E+00 | 3.4E-12 | 7.3E-01 | 7.3E-05 | 6 |
| NA | 5.1E-01 | 5.3E-03 | 1.2E+00 | 3.4E-12 | 7.3E-01 | 7.3E-05 | 6 |
| NA | 5.1E-01 | 5.3E-03 | 1.2E+00 | 3.4E-12 | 7.3E-01 | 7.3E-05 | 6 |
| NA | 5.1E-01 | 5.3E-03 | 1.2E+00 | 3.4E-12 | 7.3E-01 | 7.3E-05 | 6 |
| NA | 5.1E-01 | 5.3E-03 | 1.2E+00 | 3.4E-12 | 7.3E-01 | 7.3E-05 | 6 |
| ARL14EP | 1.7E-01 | 8.3E-01 | 9.2E-01 | 3.3E-03 | 7.5E-01 | 2.2E-02 | 6 |
| CD164 | -9.4E-02 | 9.2E-01 | 5.8E-01 | 1.6E-02 | 6.7E-01 | 3.6E-03 | 6 |
| SCAMP4 | 3.2E-01 | 4.9E-01 | 7.4E-01 | 4.4E-02 | 4.2E-01 | 4.5E-01 | 6 |
| SAT1 | 4.1E-01 | 1.3E-01 | 1.6E+00 | 1.0E-11 | 1.2E+00 | 5.8E-07 | 6 |
| C18orf21 | -2.8E-01 | 5.9E-01 | 7.7E-01 | 2.6E-02 | 1.1E+00 | 8.2E-04 | 6 |
| SUI1ISO1 | -2.6E-01 | 3.9E-01 | 5.6E-01 | 2.6E-02 | 8.2E-01 | 3.0E-04 | 6 |
| NA | 4.7E-01 | 1.1E-01 | 1.1E+00 | 4.1E-05 | 6.3E-01 | 4.4E-02 | 6 |
| NA | 4.7E-01 | 1.1E-01 | 1.1E+00 | 4.1E-05 | 6.3E-01 | 4.4E-02 | 6 |
| PRPF38B | 2.3E-01 | 5.3E-01 | 8.5E-01 | 6.9E-04 | 6.2E-01 | 2.4E-02 | 6 |
| EFHD1 | -2.5E-01 | 3.3E-01 | 4.9E-01 | 2.6E-02 | 7.4E-01 | 1.9E-04 | 6 |
| ERP29 | -3.6E-01 | 1.8E-01 | 4.6E-01 | 1.1E-01 | 8.2E-01 | 5.7E-04 | 6 |
| RBM18 | 1.0E-01 | 8.5E-01 | 8.1E-01 | 8.9E-06 | 7.1E-01 | 7.7E-05 | 6 |
| NA | 6.0E-02 | 9.8E-01 | 1.5E+00 | 1.3E-02 | 1.4E+00 | 1.5E-02 | 6 |
| NA | 5.0E-01 | 4.2E-02 | 1.1E+00 | 1.6E-06 | 6.1E-01 | 1.4E-02 | 6 |
| C12orf73 | -5.3E-01 | 3.7E-01 | 7.5E-01 | 1.8E-01 | 1.3E+00 | 6.9E-03 | 6 |
| DUSP5 | -1.1E+00 | 6.9E-02 | 2.1E+00 | 2.1E-04 | 3.2E+00 | 3.5E-10 | 6 |
| MZT1 | 2.0E-01 | 6.7E-01 | 7.2E-01 | 8.1E-03 | 5.3E-01 | 8.6E-02 | 6 |
| DSTN | -1.3E-01 | 9.5E-01 | 9.1E-01 | 1.1E-02 | 1.0E+00 | 2.7E-03 | 6 |
| RPS23 | -7.3E-02 | 9.1E-01 | 7.3E-01 | 5.1E-06 | 8.0E-01 | 1.1E-07 | 6 |
| RALA | -4.4E-01 | 3.4E-01 | 5.3E-01 | 3.0E-01 | 9.7E-01 | 1.6E-02 | 6 |
| UBE2D2 | 2.9E-01 | 1.5E-01 | 6.8E-01 | 2.5E-04 | 3.8E-01 | 8.7E-02 | 6 |
| TMEM109 | 2.6E-02 | 9.8E-01 | 6.9E-01 | 2.4E-02 | 6.6E-01 | 3.6E-02 | 6 |
| PAIP2 | -1.2E-01 | 8.8E-01 | 5.3E-01 | 5.4E-02 | 6.4E-01 | 1.4E-02 | 6 |
| RHOU | 3.2E-01 | 4.4E-01 | 1.4E+00 | 1.6E-06 | 1.1E+00 | 2.3E-04 | 6 |
| RWDD1 | -2.6E-01 | 3.3E-01 | 7.7E-01 | 3.5E-04 | 1.0E+00 | 1.8E-07 | 6 |
| NA | 2.0E-01 | 6.3E-01 | 7.0E-01 | 4.8E-03 | 5.1E-01 | 7.1E-02 | 6 |
| MRPL21 | -8.1E-02 | 9.0E-01 | 5.7E-01 | 2.0E-03 | 6.5E-01 | 2.1E-04 | 6 |
| NUDT2 | 2.8E-01 | 5.3E-01 | 1.3E+00 | 2.6E-06 | 1.1E+00 | 1.6E-04 | 6 |
| NA | 2.4E-01 | 6.0E-01 | 7.5E-01 | 1.3E-02 | 5.1E-01 | 1.5E-01 | 6 |
| NA | 2.0E-01 | 7.7E-01 | 9.7E-01 | 2.0E-03 | 7.7E-01 | 2.2E-02 | 6 |
| POLR2M | -3.8E-02 | 9.8E-01 | 6.2E-01 | 3.1E-02 | 6.6E-01 | 2.0E-02 | 6 |
| POLR2M | -3.8E-02 | 9.8E-01 | 6.2E-01 | 3.1E-02 | 6.6E-01 | 2.0E-02 | 6 |
| RMDN1 | 2.9E-01 | 3.8E-01 | 9.9E-01 | 4.9E-05 | 7.0E-01 | 6.5E-03 | 6 |
| PPIC | -2.0E-01 | 5.5E-01 | 5.7E-01 | 1.7E-02 | 7.6E-01 | 4.5E-04 | 6 |
| NA | -1.1E+00 | 5.0E-01 | 2.7E+00 | 8.1E-03 | 3.8E+00 | 1.7E-04 | 6 |
| SMIM19 | -2.1E-01 | 5.0E-01 | 4.2E-01 | 9.6E-02 | 6.2E-01 | 5.2E-03 | 6 |
| ZSCAN31 | 2.2E-01 | 9.0E-01 | 1.2E+00 | 2.0E-02 | 9.8E-01 | 6.7E-02 | 6 |
| IRX3 | 4.4E-01 | 4.0E-01 | 1.2E+00 | 4.7E-03 | 7.3E-01 | 1.5E-01 | 6 |
| ZNF780B | -1.3E-01 | 9.5E-01 | 9.8E-01 | 7.7E-03 | 1.1E+00 | 1.6E-03 | 6 |
| MRRF | 2.2E-01 | 7.4E-01 | 8.1E-01 | 2.7E-02 | 5.9E-01 | 1.8E-01 | 6 |
| PCBD2 | -4.9E-02 | 9.8E-01 | 5.6E-01 | 2.7E-02 | 6.1E-01 | 1.4E-02 | 6 |
| NA | 4.7E-01 | 3.5E-01 | 1.4E+00 | 5.6E-04 | 8.9E-01 | 3.9E-02 | 6 |
| NA | -6.7E-01 | 9.0E-01 | 2.6E+00 | 8.3E-02 | 3.3E+00 | 2.4E-02 | 6 |
| NA | 3.1E-01 | 6.2E-01 | 8.6E-01 | 4.2E-02 | 5.5E-01 | 3.3E-01 | 6 |
| NA | 5.6E-01 | 6.2E-02 | 1.3E+00 | 1.9E-06 | 7.7E-01 | 1.1E-02 | 6 |
| NA | 8.5E-01 | 2.9E-01 | 2.5E+00 | 5.6E-05 | 1.6E+00 | 6.8E-03 | 6 |
| NA | 5.9E-01 | 6.3E-01 | 1.7E+00 | 3.5E-02 | 1.1E+00 | 2.7E-01 | 6 |
| NA | 4.7E-01 | 4.8E-01 | 1.5E+00 | 1.8E-03 | 9.8E-01 | 4.4E-02 | 6 |
| NA | 4.8E-01 | 9.7E-01 | 3.5E+00 | 7.4E-03 | 3.0E+00 | 8.3E-03 | 6 |
| NA | 1.2E+00 | 7.5E-01 | 3.6E+00 | 3.9E-02 | 2.4E+00 | 8.2E-02 | 6 |
| NA | 2.1E-01 | 9.8E-01 | 4.4E+00 | 2.6E-03 | 4.2E+00 | 1.1E-03 | 6 |
| NA | 1.2E+00 | 9.0E-01 | 6.1E+00 | 7.2E-03 | 4.9E+00 | 3.2E-02 | 6 |
| ZCCHC2 | 1.2E+00 | 1.4E-02 | 1.2E-01 | 1.0E+00 | -1.1E+00 | 6.5E-02 | 4 |
| WIBG | 1.1E+00 | 7.8E-06 | 6.9E-01 | 2.1E-02 | -4.1E-01 | 2.8E-01 | 4 |
| SGK | 8.1E-01 | 1.2E-02 | 3.0E-01 | 6.7E-01 | -5.1E-01 | 3.0E-01 | 4 |
| AHI1 | 1.2E+00 | 1.4E-04 | 7.0E-01 | 8.8E-02 | -4.9E-01 | 3.2E-01 | 4 |
| SQSTM1 | 2.2E+00 | 1.5E-05 | 1.3E+00 | 4.3E-02 | -8.9E-01 | 1.8E-01 | 4 |
| FRS2 | 2.3E+00 | 1.5E-03 | 4.6E-01 | 9.0E-01 | -1.8E+00 | 2.8E-02 | 4 |
| NA | 1.6E+00 | 1.2E-02 | 7.0E-01 | 5.6E-01 | -8.9E-01 | 3.7E-01 | 4 |
| NA | 1.6E+00 | 1.2E-02 | 7.0E-01 | 5.6E-01 | -8.9E-01 | 3.7E-01 | 4 |
| NA | 1.6E+00 | 1.2E-02 | 7.0E-01 | 5.6E-01 | -8.9E-01 | 3.7E-01 | 4 |
| NA | 1.6E+00 | 1.2E-02 | 7.0E-01 | 5.6E-01 | -8.9E-01 | 3.7E-01 | 4 |
| EZR | 1.3E+00 | 5.6E-04 | 3.2E-01 | 7.5E-01 | -9.7E-01 | 3.3E-02 | 4 |
| ALG12 | 1.1E+00 | 1.2E-02 | 7.5E-01 | 2.3E-01 | -3.8E-01 | 7.7E-01 | 4 |
| SLC39A7 | 6.9E-01 | 1.2E-02 | 3.5E-03 | 1.0E+00 | -6.8E-01 | 3.3E-02 | 4 |
| PFDN6 | 2.0E+00 | 3.7E-09 | 1.4E+00 | 5.4E-04 | -6.6E-01 | 1.6E-01 | 4 |
| PFDN6 | 2.0E+00 | 3.7E-09 | 1.4E+00 | 5.4E-04 | -6.6E-01 | 1.6E-01 | 4 |
| EP300 | 1.4E+00 | 3.3E-07 | 7.7E-01 | 2.7E-02 | -6.6E-01 | 6.6E-02 | 4 |
| ST13 | 9.7E-01 | 8.2E-03 | 4.3E-01 | 5.2E-01 | -5.4E-01 | 3.6E-01 | 4 |
| TRAPPC9 | 1.7E+00 | 1.2E-05 | 1.1E+00 | 1.6E-02 | -5.7E-01 | 3.6E-01 | 4 |
| TAF11 | 1.5E+00 | 1.9E-05 | 6.4E-01 | 2.1E-01 | -8.4E-01 | 4.9E-02 | 4 |

| | | | | | | | |
|---|---|---|---|---|---|---|---|
| GRAP2 | 1.3E+00 | 2.2E-02 | 5.9E-01 | 5.6E-01 | -6.8E-01 | 4.9E-01 | 4 |
| CTNNA1 | 1.2E+00 | 2.3E-03 | 3.0E-01 | 8.0E-01 | -8.8E-01 | 7.1E-02 | 4 |
| KIAA2026 | 8.6E-01 | 4.9E-02 | 2.4E-01 | 9.2E-01 | -6.2E-01 | 3.4E-01 | 4 |
| SRPK1 | 1.0E+00 | 3.9E-03 | 2.9E-01 | 7.6E-01 | -7.1E-01 | 1.2E-01 | 4 |
| UHRF2 | 1.1E+00 | 4.6E-02 | 7.3E-01 | 3.9E-01 | -3.9E-01 | 8.2E-01 | 4 |
| STK38 | 1.9E+00 | 9.8E-03 | 1.1E+00 | 2.8E-01 | -7.3E-01 | 5.8E-01 | 4 |
| CSNK1E | 1.5E+00 | 1.5E-02 | 4.9E-01 | 7.7E-01 | -9.8E-01 | 2.4E-01 | 4 |
| PICK1 | 9.9E-01 | 1.8E-03 | 6.5E-01 | 1.2E-01 | -3.4E-01 | 6.3E-01 | 4 |
| GRINA | 1.2E+00 | 4.7E-05 | 4.0E-01 | 4.4E-01 | -8.1E-01 | 2.5E-02 | 4 |
| TRIOBP | 9.5E-01 | 2.3E-03 | 3.4E-01 | 5.9E-01 | -6.1E-01 | 1.5E-01 | 4 |
| OPLAH | 8.8E-01 | 8.1E-03 | 2.2E-01 | 8.6E-01 | -6.5E-01 | 1.3E-01 | 4 |
| NA | 7.8E-01 | 3.7E-03 | 1.5E-01 | 9.4E-01 | -6.3E-01 | 5.7E-02 | 4 |
| NA | 7.8E-01 | 3.7E-03 | 1.5E-01 | 9.4E-01 | -6.3E-01 | 5.7E-02 | 4 |
| PLIN2 | 6.4E-01 | 8.3E-03 | 4.1E-01 | 2.2E-01 | -2.3E-01 | 7.2E-01 | 4 |
| ZNF318 | 8.7E-01 | 4.5E-02 | 3.4E-01 | 7.5E-01 | -5.3E-01 | 4.7E-01 | 4 |
| ALDH1L2 | 1.3E+00 | 5.9E-04 | 6.4E-01 | 2.5E-01 | -6.4E-01 | 2.3E-01 | 4 |
| 10-Sep | 1.0E+00 | 7.5E-03 | 5.2E-01 | 3.9E-01 | -4.8E-01 | 4.6E-01 | 4 |
| SLAIN2 | 1.8E+00 | 9.6E-04 | 9.6E-01 | 2.4E-01 | -8.6E-01 | 2.5E-01 | 4 |
| SMARCA2 | 8.0E-01 | 1.5E-02 | 9.2E-02 | 1.0E+00 | -7.1E-01 | 8.2E-02 | 4 |
| USP46 | 1.2E+00 | 1.2E-02 | 7.8E-02 | 1.0E+00 | -1.2E+00 | 4.6E-02 | 4 |
| CD2AP | 1.0E+00 | 2.0E-03 | 6.7E-01 | 1.3E-01 | -3.7E-01 | 6.3E-01 | 4 |
| UXS1 | 8.4E-01 | 2.5E-03 | 5.4E-02 | 1.0E+00 | -7.9E-01 | 1.6E-02 | 4 |
| IL1R1 | 1.8E+00 | 1.1E-02 | 6.5E-01 | 6.8E-01 | -1.1E+00 | 2.5E-01 | 4 |
| TLN1 | 1.2E+00 | 1.5E-03 | 3.7E-01 | 6.7E-01 | -8.5E-01 | 8.5E-02 | 4 |
| EXOC1 | 1.2E+00 | 3.4E-03 | 4.6E-01 | 5.7E-01 | -7.5E-01 | 1.9E-01 | 4 |
| ASNS | 1.0E+00 | 4.5E-04 | 6.7E-01 | 7.5E-02 | -3.8E-01 | 5.4E-01 | 4 |
| ZNF579 | 7.6E-01 | 5.7E-03 | 1.3E-01 | 9.8E-01 | -6.3E-01 | 6.5E-02 | 4 |
| RHOQ | 1.1E+00 | 2.1E-03 | 4.7E-01 | 4.7E-01 | -6.6E-01 | 2.0E-01 | 4 |
| ERLEC1 | 6.5E-01 | 4.7E-02 | 9.7E-02 | 1.0E+00 | -5.5E-01 | 2.1E-01 | 4 |
| RAPGEF5 | 1.8E+00 | 4.7E-02 | 7.6E-01 | 7.4E-01 | -1.1E+00 | 4.1E-01 | 4 |
| IGF2BP3 | 1.3E+00 | 1.5E-02 | 2.8E-01 | 9.6E-01 | -1.1E+00 | 1.3E-01 | 4 |
| PALM2 | 1.1E+00 | 7.3E-04 | 4.1E-01 | 4.8E-01 | -6.7E-01 | 1.1E-01 | 4 |
| PALM2 | 1.1E+00 | 7.3E-04 | 4.1E-01 | 4.8E-01 | -6.7E-01 | 1.1E-01 | 4 |
| PALM2 | 1.1E+00 | 7.3E-04 | 4.1E-01 | 4.8E-01 | -6.7E-01 | 1.1E-01 | 4 |
| PALM2 | 1.1E+00 | 7.3E-04 | 4.1E-01 | 4.8E-01 | -6.7E-01 | 1.1E-01 | 4 |
| PALM2 | 1.1E+00 | 7.3E-04 | 4.1E-01 | 4.8E-01 | -6.7E-01 | 1.1E-01 | 4 |
| HOXA7 | 1.3E+00 | 1.5E-03 | 6.8E-01 | 2.9E-01 | -6.6E-01 | 3.1E-01 | 4 |
| MDN1 | 8.9E-01 | 1.3E-02 | 5.4E-01 | 3.0E-01 | -3.5E-01 | 6.9E-01 | 4 |
| MEIS1 | 2.6E+00 | 4.2E-06 | 1.4E+00 | 4.0E-02 | -1.1E+00 | 9.5E-02 | 4 |
| GMCL1 | 1.1E+00 | 4.5E-02 | 6.7E-01 | 4.6E-01 | -4.5E-01 | 7.6E-01 | 4 |
| KIDINS220 | 9.0E-01 | 4.2E-02 | 4.5E-01 | 5.8E-01 | -4.5E-01 | 6.3E-01 | 4 |
| STK17A | 1.7E+00 | 1.6E-06 | 5.9E-01 | 3.0E-01 | -1.1E+00 | 5.9E-03 | 4 |
| INHBA | 1.5E+00 | 3.6E-03 | 5.6E-01 | 5.9E-01 | -9.8E-01 | 1.9E-01 | 4 |
| CADPS2 | 8.2E-01 | 2.6E-02 | 8.8E-02 | 1.0E+00 | -7.3E-01 | 1.2E-01 | 4 |
| IRF3 | 6.5E-01 | 1.5E-02 | 1.2E-01 | 9.8E-01 | -5.3E-01 | 1.3E-01 | 4 |
| ELF2 | 1.3E+00 | 1.4E-03 | 3.2E-01 | 7.9E-01 | -9.4E-01 | 5.1E-02 | 4 |
| SMPD2 | 7.7E-01 | 4.1E-03 | 4.4E-01 | 2.5E-01 | -3.3E-01 | 5.3E-01 | 4 |
| LAPTM4A | 8.8E-01 | 3.8E-02 | 2.2E-01 | 9.4E-01 | -6.6E-01 | 2.8E-01 | 4 |
| KIAA1109 | 8.1E-01 | 4.9E-02 | 2.5E-01 | 8.8E-01 | -5.6E-01 | 3.7E-01 | 4 |
| SRPK2 | 1.2E+00 | 2.9E-05 | 3.5E-01 | 5.4E-01 | -8.6E-01 | 1.2E-02 | 4 |
| DNMT3A | 1.0E+00 | 1.8E-02 | 3.6E-01 | 7.4E-01 | -6.6E-01 | 3.0E-01 | 4 |
| TMEM160 | 1.9E+00 | 3.2E-02 | 5.6E-02 | 1.0E+00 | -1.8E+00 | 9.1E-02 | 4 |
| ARHGAP35 | 2.4E+00 | 1.4E-05 | 1.1E+00 | 1.8E-01 | -1.3E+00 | 2.6E-02 | 4 |
| AP2S1 | 1.4E+00 | 1.7E-07 | 8.8E-01 | 6.2E-03 | -5.4E-01 | 1.6E-01 | 4 |
| PRKD2 | 1.2E+00 | 2.9E-03 | 2.1E-01 | 9.6E-01 | -1.0E+00 | 4.0E-02 | 4 |
| FNDC3A | 1.1E+00 | 2.5E-02 | 7.6E-01 | 2.6E-01 | -3.4E-01 | 8.7E-01 | 4 |
| CAB39L | 1.2E+00 | 8.4E-06 | 6.3E-02 | 1.0E+00 | -1.1E+00 | 7.2E-05 | 4 |
| GPBP1L1 | 1.6E+00 | 2.3E-03 | 3.7E-01 | 8.6E-01 | -1.2E+00 | 4.8E-02 | 4 |
| KLC3 | 1.1E+00 | 1.5E-03 | 7.0E-01 | 1.0E-01 | -3.6E-01 | 6.5E-01 | 4 |
| STAM | 1.1E+00 | 4.5E-02 | 2.1E-01 | 1.0E+00 | -8.6E-01 | 2.3E-01 | 4 |
| TESK2 | 2.4E+00 | 4.2E-06 | 1.0E+00 | 1.7E-01 | -1.4E+00 | 1.7E-02 | 4 |
| PRR2 | 1.5E+00 | 2.0E-06 | 7.2E-02 | 1.0E+00 | -1.4E+00 | 1.5E-05 | 4 |
| CAD | 1.1E+00 | 2.2E-08 | 1.5E-01 | 8.4E-01 | -9.8E-01 | 5.1E-06 | 4 |
| TFCP2L1 | 1.3E+00 | 7.9E-03 | 5.7E-01 | 5.1E-01 | -7.1E-01 | 3.4E-01 | 4 |
| SLC5A6 | 2.5E+00 | 9.4E-03 | 1.2E+00 | 4.6E-01 | -1.3E+00 | 2.9E-01 | 4 |
| PAFAH1B3 | 7.2E-01 | 1.1E-02 | 3.5E-01 | 4.6E-01 | -3.7E-01 | 4.8E-01 | 4 |
| DEDD2 | 1.2E+00 | 2.1E-04 | 7.0E-01 | 9.0E-02 | -4.8E-01 | 3.7E-01 | 4 |
| KMT2C | 9.1E-01 | 1.8E-03 | 4.5E-02 | 1.0E+00 | -8.7E-01 | 1.0E-02 | 4 |
| NA | 1.2E+00 | 2.6E-02 | -5.9E-03 | 1.0E+00 | -1.2E+00 | 6.1E-02 | 4 |
| NA | 1.2E+00 | 2.6E-02 | -5.9E-03 | 1.0E+00 | -1.2E+00 | 6.1E-02 | 4 |
| TAF3 | 8.4E-01 | 2.9E-03 | 3.1E-01 | 5.7E-01 | -5.3E-01 | 1.8E-01 | 4 |
| EXOC7 | 1.0E+00 | 2.0E-05 | 3.9E-01 | 2.9E-01 | -6.2E-01 | 3.3E-02 | 4 |
| LMO7 | 6.7E-01 | 4.1E-04 | 1.9E-01 | 6.4E-01 | -4.7E-01 | 4.3E-02 | 4 |
| WDR48 | 1.3E+00 | 1.3E-02 | 6.7E-01 | 4.8E-01 | -6.8E-01 | 4.4E-01 | 4 |
| BLVRB | 6.9E-01 | 1.3E-03 | 4.5E-01 | 1.0E-01 | -2.4E-01 | 6.3E-01 | 4 |
| RRBP1 | 1.0E+00 | 3.4E-04 | 7.2E-02 | 1.0E+00 | -9.7E-01 | 3.1E-03 | 4 |
| PPME1 | 1.7E+00 | 3.7E-04 | 7.9E-01 | 2.6E-01 | -8.6E-01 | 1.6E-01 | 4 |
| SUPT16H | 1.1E+00 | 1.6E-02 | 3.7E-01 | 7.3E-01 | -6.8E-01 | 2.8E-01 | 4 |
| TOX4 | 1.3E+00 | 8.9E-04 | 7.2E-02 | 1.0E+00 | -1.2E+00 | 5.2E-03 | 4 |
| DNAJC18 | 1.1E+00 | 1.6E-03 | 1.1E-01 | 1.0E+00 | -1.0E+00 | 1.4E-02 | 4 |
| BIRC6 | 1.1E+00 | 2.3E-03 | 6.1E-01 | 2.3E-01 | -4.6E-01 | 4.5E-01 | 4 |
| TMX4 | 1.3E+00 | 2.1E-02 | 6.3E-01 | 5.6E-01 | -6.8E-01 | 4.6E-01 | 4 |
| DCXR | 7.4E-01 | 1.2E-03 | 1.7E-01 | 8.3E-01 | -5.8E-01 | 4.0E-02 | 4 |
| C20orf196 | 1.2E+00 | 3.2E-03 | 1.3E-01 | 1.0E+00 | -1.0E+00 | 2.6E-02 | 4 |
| GOT1L1 | 6.3E-01 | 3.6E-02 | 2.6E-01 | 6.9E-01 | -3.7E-01 | 4.8E-01 | 4 |
| ZNF703 | 2.4E+00 | 1.4E-10 | 1.5E+00 | 5.0E-04 | -9.0E-01 | 6.1E-02 | 4 |
| GNPDA1 | 2.3E+00 | 9.3E-04 | 9.3E-01 | 4.7E-01 | -1.4E+00 | 7.4E-02 | 4 |
| SEMA3E | 1.0E+00 | 1.3E-02 | 3.4E-01 | 7.5E-01 | -6.9E-01 | 2.4E-01 | 4 |
| EP400 | 1.1E+00 | 1.2E-02 | 2.4E-01 | 9.4E-01 | -8.6E-01 | 1.2E-01 | 4 |
| UBOX5 | 1.4E+00 | 1.5E-03 | 1.8E-01 | 1.0E+00 | -1.2E+00 | 1.8E-02 | 4 |
| RRP8 | 1.3E+00 | 1.1E-02 | 4.6E-01 | 7.1E-01 | -8.2E-01 | 2.4E-01 | 4 |
| PKP3 | 1.7E+00 | 1.5E-09 | 1.1E+00 | 4.9E-04 | -5.9E-01 | 1.5E-01 | 4 |
| NA | 1.9E+00 | 2.6E-06 | 7.5E-01 | 2.1E-01 | -1.2E+00 | 1.6E-02 | 4 |
| NCOA4 | 9.9E-01 | 1.1E-02 | 3.1E-01 | 7.7E-01 | -6.8E-01 | 2.1E-01 | 4 |

| | | | | | | | |
|---|---|---|---|---|---|---|---|
| NCOR2 | 1.5E+00 | 1.4E-04 | 7.5E-01 | 1.9E-01 | -8.0E-01 | 1.5E-01 | 4 |
| RASSF4 | 8.9E-01 | 4.0E-02 | 3.0E-01 | 8.4E-01 | -6.0E-01 | 3.7E-01 | 4 |
| ZNF740 | 1.8E+00 | 1.7E-07 | 6.7E-01 | 1.6E-01 | -1.1E+00 | 5.0E-03 | 4 |
| ASAH1 | 1.6E+00 | 2.1E-04 | 6.8E-01 | 3.0E-01 | -8.9E-01 | 1.1E-01 | 4 |
| SMIM15 | 1.3E+00 | 1.3E-03 | 6.7E-01 | 2.3E-01 | -6.0E-01 | 3.6E-01 | 4 |
| KIF3B | 1.1E+00 | 4.5E-02 | 7.4E-02 | 1.0E+00 | -1.0E+00 | 1.4E-01 | 4 |
| FAT1 | 1.3E+00 | 1.2E-04 | 2.0E-01 | 9.3E-01 | -1.1E+00 | 4.0E-03 | 4 |
| PAX8 | 2.0E+00 | 5.5E-12 | 1.1E+00 | 7.3E-04 | -8.5E-01 | 1.6E-02 | 4 |
| SETD8 | 1.6E+00 | 1.4E-02 | 2.9E-01 | 1.0E+00 | -1.4E+00 | 1.0E-01 | 4 |
| HHEX | 2.1E+00 | 3.5E-03 | 3.0E-01 | 1.0E+00 | -1.8E+00 | 3.2E-02 | 4 |
| ITCH | 1.5E+00 | 2.3E-07 | 3.8E-01 | 4.9E-01 | -1.1E+00 | 3.9E-04 | 4 |
| MRAS | 2.1E+00 | 3.7E-02 | 1.1E+00 | 5.4E-01 | -1.0E+00 | 5.4E-01 | 4 |
| OSBP | 1.7E+00 | 7.4E-05 | 6.5E-01 | 3.8E-01 | -1.1E+00 | 3.2E-02 | 4 |
| SH3RF1 | 1.7E+00 | 4.3E-04 | 8.2E-01 | 2.4E-01 | -8.6E-01 | 2.0E-01 | 4 |
| ZBTB38 | 1.3E+00 | 1.5E-04 | 1.9E-01 | 9.5E-01 | -1.1E+00 | 4.4E-03 | 4 |
| EIF4G2 | 1.7E+00 | 2.1E-04 | 5.9E-01 | 4.9E-01 | -1.1E+00 | 4.3E-02 | 4 |
| ERAP1 | 1.3E+00 | 6.5E-05 | 8.0E-01 | 4.3E-02 | -4.6E-01 | 3.6E-01 | 4 |
| CAST | 1.3E+00 | 7.7E-05 | 5.6E-01 | 2.2E-01 | -7.0E-01 | 1.1E-01 | 4 |
| SSRP1 | 7.8E-01 | 2.5E-02 | 2.5E-01 | 8.0E-01 | -5.3E-01 | 3.2E-01 | 4 |
| LARP4 | 1.3E+00 | 5.0E-03 | 4.2E-01 | 7.3E-01 | -9.2E-01 | 1.4E-01 | 4 |
| LYSMD3 | 1.4E+00 | 2.2E-02 | 8.7E-01 | 3.6E-01 | -5.7E-01 | 6.9E-01 | 4 |
| CXADR | 1.1E+00 | 1.8E-02 | 2.8E-01 | 9.1E-01 | -8.6E-01 | 1.8E-01 | 4 |
| SMARCD1 | 1.8E+00 | 3.9E-02 | 3.1E-01 | 1.0E+00 | -1.5E+00 | 1.8E-01 | 4 |
| G3BP2 | 1.1E+00 | 3.4E-03 | 5.7E-02 | 1.0E+00 | -1.1E+00 | 1.6E-02 | 4 |
| ZBTB10 | 1.1E+00 | 3.6E-02 | 6.6E-01 | 4.4E-01 | -4.6E-01 | 7.4E-01 | 4 |
| EDIL3 | 1.6E+00 | 1.3E-02 | 3.7E-01 | 9.2E-01 | -1.2E+00 | 1.4E-01 | 4 |
| INO80B | 1.7E+00 | 2.3E-05 | 9.7E-01 | 5.7E-02 | -7.1E-01 | 1.9E-01 | 4 |
| 11-Sep | 1.2E+00 | 4.1E-03 | 2.5E-01 | 8.9E-01 | -9.0E-01 | 7.4E-02 | 4 |
| RHEBL1 | 1.5E+00 | 5.9E-03 | 2.8E-01 | 9.6E-01 | -1.2E+00 | 6.8E-02 | 4 |
| RAPGEF2 | 1.1E+00 | 7.2E-03 | 3.3E-01 | 7.8E-01 | -7.8E-01 | 1.5E-01 | 4 |
| ANTXR2 | 9.3E-01 | 5.7E-04 | 2.9E-01 | 6.1E-01 | -6.4E-01 | 6.0E-02 | 4 |
| PTPN11 | 8.7E-01 | 3.3E-03 | 4.0E-01 | 4.2E-01 | -4.7E-01 | 3.0E-01 | 4 |
| RSRC1 | 8.6E-01 | 2.5E-03 | 1.8E-01 | 8.9E-01 | -6.8E-01 | 5.4E-02 | 4 |
| CKAP5 | 1.2E+00 | 4.4E-02 | 1.6E-01 | 1.0E+00 | -1.0E+00 | 1.8E-01 | 4 |
| TOP | 1.2E+00 | 8.8E-04 | 1.5E-01 | 1.0E+00 | -1.1E+00 | 1.3E-02 | 4 |
| STC1 | 9.1E-01 | 1.1E-02 | 2.9E-01 | 7.5E-01 | -6.2E-01 | 2.3E-01 | 4 |
| RBM19 | 9.5E-01 | 1.5E-03 | 4.5E-01 | 3.4E-01 | -5.0E-01 | 2.5E-01 | 4 |
| RSBN1 | 9.5E-01 | 1.1E-02 | 1.9E-01 | 9.5E-01 | -7.6E-01 | 1.2E-01 | 4 |
| AGGF1 | 9.3E-01 | 7.5E-03 | 2.5E-01 | 8.3E-01 | -6.8E-01 | 1.4E-01 | 4 |
| R3HCC1L | 1.2E+00 | 3.8E-02 | 3.5E-01 | 9.0E-01 | -9.0E-01 | 2.9E-01 | 4 |
| SPTY2D1 | 8.2E-01 | 4.6E-02 | 5.1E-01 | 4.3E-01 | -3.1E-01 | 8.2E-01 | 4 |
| PEX16 | 1.4E+00 | 3.0E-05 | 3.0E-01 | 7.5E-01 | -1.1E+00 | 3.9E-03 | 4 |
| NISCH | 7.1E-01 | 2.3E-02 | 1.3E-01 | 9.9E-01 | -5.8E-01 | 1.6E-01 | 4 |
| PNPLA2 | 6.9E-01 | 2.6E-02 | 3.6E-01 | 4.8E-01 | -3.2E-01 | 6.3E-01 | 4 |
| CEP97 | 2.0E+00 | 4.2E-02 | 2.6E-02 | 1.0E+00 | -2.0E+00 | 9.6E-02 | 4 |
| SIM2 | 2.1E+00 | 4.7E-02 | 1.3E+00 | 4.4E-01 | -8.1E-01 | 7.1E-01 | 4 |
| YAF2 | 1.3E+00 | 2.1E-03 | 7.4E-01 | 2.3E-01 | -5.7E-01 | 4.2E-01 | 4 |
| WDR3 | 9.2E-01 | 1.7E-02 | -1.4E-02 | 1.0E+00 | -9.4E-01 | 3.9E-02 | 4 |
| HMGN1 | 2.6E+00 | 2.1E-05 | 1.6E+00 | 4.7E-02 | -1.1E+00 | 1.3E-01 | 4 |
| HMGN1 | 2.6E+00 | 2.1E-05 | 1.6E+00 | 4.7E-02 | -1.1E+00 | 1.3E-01 | 4 |
| NOLC1 | 1.6E+00 | 8.8E-04 | 5.8E-01 | 5.2E-01 | -1.0E+00 | 1.0E-01 | 4 |
| GCN1L1 | 6.0E-01 | 1.1E-02 | 2.2E-01 | 6.6E-01 | -3.8E-01 | 2.9E-01 | 4 |
| ACTR1A | 6.5E-01 | 1.7E-02 | 1.6E-01 | 9.1E-01 | -4.9E-01 | 1.9E-01 | 4 |
| TSPAN5 | 7.5E-01 | 1.2E-03 | 3.6E-01 | 3.0E-01 | -3.9E-01 | 2.8E-01 | 4 |
| KIAA0947 | 1.3E+00 | 2.1E-02 | 2.4E-01 | 9.9E-01 | -1.0E+00 | 1.4E-01 | 4 |
| KDM5B | 1.4E+00 | 6.6E-06 | 4.0E-01 | 4.8E-01 | -9.8E-01 | 5.6E-03 | 4 |
| OASL1 | 2.2E+00 | 4.6E-02 | 1.1E+00 | 5.8E-01 | -1.1E+00 | 5.9E-01 | 4 |
| LGR6 | 2.4E+00 | 4.6E-02 | 1.1E+00 | 7.0E-01 | -1.4E+00 | 4.2E-01 | 4 |
| HRPT2 | 1.3E+00 | 1.1E-02 | 6.2E-01 | 5.0E-01 | -6.9E-01 | 4.0E-01 | 4 |
| COL17A1 | 1.6E+00 | 6.6E-06 | 4.3E-01 | 5.3E-01 | -1.1E+00 | 3.5E-03 | 4 |
| HDAC1 | 1.3E+00 | 3.7E-02 | 2.5E-01 | 1.0E+00 | -1.0E+00 | 2.0E-01 | 4 |
| RRP1B | 5.9E-01 | 2.2E-02 | 2.5E-01 | 6.2E-01 | -3.4E-01 | 4.4E-01 | 4 |
| SP110 | 1.3E+00 | 3.5E-04 | 5.7E-01 | 3.5E-01 | -7.5E-01 | 1.1E-01 | 4 |
| KIAA2018 | 1.3E+00 | 2.9E-03 | 8.6E-01 | 1.4E-01 | -4.4E-01 | 6.5E-01 | 4 |
| ICOSLG | 1.3E+00 | 2.3E-03 | 7.3E-01 | 2.4E-01 | -5.7E-01 | 4.1E-01 | 4 |
| SMNDC1 | 9.0E-01 | 1.7E-02 | 5.4E-01 | 3.4E-01 | -3.6E-01 | 6.9E-01 | 4 |
| MVK | 7.0E-01 | 6.0E-03 | 4.4E-01 | 2.2E-01 | -2.7E-01 | 6.6E-01 | 4 |
| BRD9 | 1.2E+00 | 3.2E-06 | 7.0E-01 | 3.4E-02 | -5.4E-01 | 1.4E-01 | 4 |
| NA | 9.2E-01 | 1.2E-02 | 3.0E-01 | 7.6E-01 | -6.2E-01 | 2.3E-01 | 4 |
| RBM5 | 1.0E+00 | 2.7E-02 | 2.2E-01 | 9.7E-01 | -8.1E-01 | 1.9E-01 | 4 |
| EI24 | 1.0E+00 | 8.7E-05 | 6.5E-02 | 1.0E+00 | -9.4E-01 | 8.7E-04 | 4 |
| C21orf2 | 2.2E+00 | 2.5E-02 | 1.3E+00 | 3.6E-01 | -8.2E-01 | 6.6E-01 | 4 |
| ADARB1 | 1.4E+00 | 1.2E-02 | 7.5E-01 | 4.3E-01 | -6.8E-01 | 4.8E-01 | 4 |
| NDUFS8 | 6.9E-01 | 1.1E-02 | 2.3E-01 | 7.3E-01 | -4.6E-01 | 2.4E-01 | 4 |
| NDUFS8 | 6.9E-01 | 1.1E-02 | 2.3E-01 | 7.3E-01 | -4.6E-01 | 2.4E-01 | 4 |
| NDUFS8 | 6.9E-01 | 1.1E-02 | 2.3E-01 | 7.3E-01 | -4.6E-01 | 2.4E-01 | 4 |
| MMP-14 | 1.5E+00 | 2.4E-06 | 8.6E-01 | 3.4E-02 | -6.9E-01 | 1.3E-01 | 4 |
| ITPR2 | 7.7E-01 | 4.2E-02 | 4.9E-01 | 3.9E-01 | -2.8E-01 | 8.4E-01 | 4 |
| AP1G2 | 6.7E-01 | 8.4E-03 | 3.8E-01 | 3.1E-01 | -2.9E-01 | 5.7E-01 | 4 |
| ELOVL6 | 1.3E+00 | 4.1E-02 | 7.7E-02 | 1.0E+00 | -1.2E+00 | 1.2E-01 | 4 |
| RAB22A | 1.1E+00 | 2.8E-02 | 3.4E-01 | 8.3E-01 | -7.4E-01 | 3.1E-01 | 4 |
| PRKAR2A | 1.5E+00 | 2.7E-02 | 5.6E-01 | 7.3E-01 | -9.0E-01 | 3.6E-01 | 4 |
| UNC45A | 8.9E-01 | 1.2E-03 | 1.4E-01 | 9.7E-01 | -7.5E-01 | 2.1E-02 | 4 |
| LRRFIP1 | 1.2E+00 | 7.1E-03 | 5.7E-01 | 4.7E-01 | -6.5E-01 | 3.5E-01 | 4 |
| PPP2R5A | 1.0E+00 | 4.2E-03 | 3.3E-01 | 7.1E-01 | -6.9E-01 | 1.4E-01 | 4 |
| OATPA | 1.2E+00 | 2.6E-02 | 7.9E-01 | 3.2E-01 | -4.3E-01 | 8.2E-01 | 4 |
| NFATC4 | 7.1E-01 | 1.4E-03 | 6.3E-02 | 1.0E+00 | -6.5E-01 | 1.3E-02 | 4 |
| RPS6KA1 | 8.7E-01 | 2.5E-03 | 2.3E-01 | 7.8E-01 | -6.4E-01 | 8.3E-02 | 4 |
| CCDC6 | 2.1E+00 | 4.8E-02 | 1.0E+00 | 6.1E-01 | -1.0E+00 | 5.3E-01 | 4 |
| GATSL3 | 1.4E+00 | 1.2E-04 | 1.7E-01 | 9.9E-01 | -1.3E+00 | 2.5E-03 | 4 |
| ZNF148 | 1.2E+00 | 6.5E-03 | 4.4E-01 | 6.3E-01 | -7.5E-01 | 2.3E-01 | 4 |
| RNPEPL1 | 1.7E+00 | 1.1E-03 | 3.4E-01 | 8.9E-01 | -1.4E+00 | 2.7E-02 | 4 |
| EIF4G1 | 9.7E-01 | 8.2E-03 | 1.1E-01 | 1.0E+00 | -8.6E-01 | 5.4E-02 | 4 |
| KMT2A | 9.7E-01 | 9.7E-04 | 2.7E-01 | 7.0E-01 | -7.0E-01 | 5.8E-02 | 4 |

| | | | | | | | |
|---|---|---|---|---|---|---|---|
| CHST15 | 7.3E-01 | 3.8E-02 | 6.2E-02 | 1.0E+00 | -6.7E-01 | 1.4E-01 | 4 |
| FAM53B | 1.2E+00 | 4.2E-05 | 6.7E-01 | 7.6E-02 | -5.1E-01 | 2.2E-01 | 4 |
| ARFGAP1 | 2.3E+00 | 1.6E-06 | 1.3E+00 | 2.1E-02 | -9.4E-01 | 1.1E-01 | 4 |
| STRN3 | 1.2E+00 | 4.3E-04 | 5.5E-01 | 3.1E-01 | -6.7E-01 | 1.6E-01 | 4 |
| TMPRSS4 | 1.0E+00 | 1.7E-03 | 5.8E-01 | 1.9E-01 | -4.2E-01 | 4.8E-01 | 4 |
| CCDC47 | 1.0E+00 | 5.7E-03 | 1.5E-01 | 1.0E+00 | -8.7E-01 | 5.3E-02 | 4 |
| NA | 6.2E-01 | 1.1E-02 | 3.1E-01 | 4.5E-01 | -3.1E-01 | 4.8E-01 | 4 |
| THEM4 | 1.5E+00 | 3.3E-07 | 8.2E-01 | 2.8E-02 | -7.1E-01 | 5.7E-02 | 4 |
| DHX32 | 3.8E+00 | 1.5E-02 | 8.3E-01 | 9.6E-01 | -3.0E+00 | 5.7E-02 | 4 |
| PCYT1A | 1.5E+00 | 1.2E-03 | 4.1E-01 | 7.3E-01 | -1.1E+00 | 5.9E-02 | 4 |
| C10orf90 | 2.8E+00 | 9.2E-03 | 1.8E+00 | 2.3E-01 | -9.9E-01 | 4.3E-01 | 4 |
| CEP164 | 7.3E-01 | 4.0E-02 | 1.8E-01 | 9.5E-01 | -5.5E-01 | 2.7E-01 | 4 |
| DOCK1 | 7.3E-01 | 1.9E-02 | 8.3E-02 | 1.0E+00 | -6.5E-01 | 9.8E-02 | 4 |
| SMARCC1 | 7.9E-01 | 1.4E-03 | 5.0E-01 | 1.2E-01 | -2.9E-01 | 5.9E-01 | 4 |
| BAZ1B | 1.3E+00 | 1.3E-04 | 6.2E-01 | 2.2E-01 | -7.2E-01 | 1.3E-01 | 4 |
| NA | 9.4E-01 | 1.2E-02 | 6.2E-01 | 2.2E-01 | -3.2E-01 | 7.9E-01 | 4 |
| CDKN1B | 7.6E-01 | 5.4E-04 | 1.0E-01 | 9.9E-01 | -6.6E-01 | 1.0E-02 | 4 |
| FAM177A1 | 1.1E+00 | 1.4E-03 | 5.7E-01 | 2.9E-01 | -5.6E-01 | 3.1E-01 | 4 |
| FAM177A1 | 1.1E+00 | 1.4E-03 | 5.7E-01 | 2.9E-01 | -5.6E-01 | 3.1E-01 | 4 |
| TMCO1 | 1.0E+00 | 3.2E-04 | 6.9E-01 | 5.5E-02 | -3.5E-01 | 5.6E-01 | 4 |
| TMEM120A | 6.5E-01 | 2.2E-03 | 2.6E-01 | 5.1E-01 | -3.9E-01 | 1.9E-01 | 4 |
| DTX2 | 8.5E-01 | 9.3E-04 | 2.1E-01 | 7.8E-01 | -6.4E-01 | 4.2E-02 | 4 |
| CUX1 | 1.1E+00 | 1.8E-02 | 4.5E-01 | 6.1E-01 | -6.0E-01 | 4.2E-01 | 4 |
| SOAT1 | 4.5E+00 | 1.4E-02 | 2.2E+00 | 5.3E-01 | -2.3E+00 | 3.6E-01 | 4 |
| AP1S1 | 2.2E+00 | 3.9E-03 | 1.0E+00 | 4.1E-01 | -1.1E+00 | 2.3E-01 | 4 |
| SLC2A3 | 1.7E+00 | 3.1E-02 | 7.7E-01 | 6.6E-01 | -9.6E-01 | 4.1E-01 | 4 |
| SLC2A3 | 1.7E+00 | 3.1E-02 | 7.7E-01 | 6.6E-01 | -9.6E-01 | 4.1E-01 | 4 |
| LPP | 1.2E+00 | 2.0E-04 | 4.0E-01 | 5.1E-01 | -8.1E-01 | 4.6E-02 | 4 |
| RALGPS2 | 9.9E-01 | 1.8E-03 | 1.7E-01 | 9.5E-01 | -8.2E-01 | 3.1E-02 | 4 |
| EFTUD2 | 1.7E+00 | 2.7E-02 | 1.3E-01 | 1.0E+00 | -1.6E+00 | 8.8E-02 | 4 |
| ATG2A | 1.3E+00 | 5.2E-05 | 3.5E-01 | 6.1E-01 | -9.4E-01 | 1.2E-02 | 4 |
| NA | 1.4E+00 | 7.3E-04 | 5.7E-01 | 4.6E-01 | -8.7E-01 | 1.1E-01 | 4 |
| ATXN7L3 | 1.7E+00 | 2.7E-03 | 1.1E-01 | 1.0E+00 | -1.6E+00 | 1.3E-02 | 4 |
| RDX | 6.9E-01 | 1.2E-02 | 2.9E-01 | 5.7E-01 | -4.0E-01 | 3.7E-01 | 4 |
| LRCH4 | 1.5E+00 | 4.1E-09 | 4.4E-01 | 3.1E-01 | -1.1E+00 | 6.9E-05 | 4 |
| G6PC3 | 8.5E-01 | 1.1E-02 | 9.9E-02 | 1.0E+00 | -7.5E-01 | 6.4E-02 | 4 |
| ATM | 1.2E+00 | 1.1E-02 | 7.0E-01 | 3.1E-01 | -4.6E-01 | 6.1E-01 | 4 |
| PEA3 | 1.6E+00 | 2.3E-05 | 6.5E-01 | 2.4E-01 | -9.2E-01 | 4.8E-02 | 4 |
| HIRA | 2.4E+00 | 2.0E-02 | 8.5E-01 | 7.5E-01 | -1.5E+00 | 2.2E-01 | 4 |
| PTMS | 1.3E+00 | 1.5E-03 | 4.6E-01 | 5.7E-01 | -8.4E-01 | 1.3E-01 | 4 |
| CCDC56 | 6.1E-01 | 4.0E-02 | 2.6E-01 | 6.7E-01 | -3.5E-01 | 5.2E-01 | 4 |
| SH3BP2 | 9.2E-01 | 1.7E-03 | 5.4E-01 | 1.7E-01 | -3.8E-01 | 5.0E-01 | 4 |
| FAM134C | 1.4E+00 | 5.0E-05 | -6.2E-02 | 1.0E+00 | -1.5E+00 | 7.2E-05 | 4 |
| CHD4 | 1.0E+00 | 1.8E-04 | 5.2E-01 | 1.8E-01 | -5.3E-01 | 1.9E-01 | 4 |
| UBA1 | 9.0E-01 | 2.2E-04 | 2.3E-01 | 7.0E-01 | -6.7E-01 | 2.2E-02 | 4 |
| NA | 9.0E-01 | 1.9E-03 | 5.4E-01 | 1.8E-01 | -3.7E-01 | 5.0E-01 | 4 |
| NA | 7.0E-01 | 2.3E-02 | 4.1E-01 | 3.7E-01 | -2.9E-01 | 7.1E-01 | 4 |
| NA | 7.0E-01 | 2.3E-02 | 4.1E-01 | 3.7E-01 | -2.9E-01 | 7.1E-01 | 4 |
| NA | 7.0E-01 | 2.3E-02 | 4.1E-01 | 3.7E-01 | -2.9E-01 | 7.1E-01 | 4 |
| NA | 7.0E-01 | 2.3E-02 | 4.1E-01 | 3.7E-01 | -2.9E-01 | 7.1E-01 | 4 |
| NA | 7.0E-01 | 2.3E-02 | 4.1E-01 | 3.7E-01 | -2.9E-01 | 7.1E-01 | 4 |
| HP1BP3 | 8.1E-01 | 1.1E-02 | 4.8E-01 | 3.1E-01 | -3.4E-01 | 6.3E-01 | 4 |
| SLBP | 2.5E+00 | 4.9E-02 | 6.5E-01 | 9.5E-01 | -1.8E+00 | 2.3E-01 | 4 |
| FBXO34 | 1.3E+00 | 6.2E-03 | 4.3E-01 | 6.9E-01 | -8.3E-01 | 1.8E-01 | 4 |
| PDAP1 | 1.5E+00 | 6.4E-03 | 1.1E-01 | 1.0E+00 | -1.4E+00 | 3.2E-02 | 4 |
| SELP | 1.1E+00 | 3.2E-02 | 5.0E-01 | 6.1E-01 | -5.9E-01 | 5.2E-01 | 4 |
| KAT6B | 8.8E-01 | 4.2E-02 | 3.1E-01 | 8.0E-01 | -5.7E-01 | 4.2E-01 | 4 |
| CCND2 | 7.4E-01 | 5.9E-03 | 1.6E-01 | 9.0E-01 | -5.7E-01 | 9.6E-02 | 4 |
| KIAA0753 | 1.6E+00 | 1.1E-07 | 9.5E-01 | 6.9E-03 | -6.3E-01 | 1.3E-01 | 4 |
| ZNF503 | 1.1E+00 | 1.6E-03 | 2.1E-01 | 9.3E-01 | -9.0E-01 | 3.0E-02 | 4 |
| ARL6IP1 | 2.1E+00 | 2.6E-04 | 1.1E+00 | 1.5E-01 | -9.8E-01 | 2.3E-01 | 4 |
| OTUD5 | 7.8E-01 | 1.5E-02 | 4.9E-01 | 2.9E-01 | -2.9E-01 | 7.5E-01 | 4 |
| GRIPAP1 | 1.2E+00 | 5.8E-05 | 3.9E-01 | 4.6E-01 | -7.8E-01 | 2.6E-02 | 4 |
| MYO5A | 1.3E+00 | 6.5E-05 | 7.7E-01 | 4.8E-02 | -4.8E-01 | 3.6E-01 | 4 |
| DNAJB6 | 2.6E+00 | 3.9E-02 | 1.8E+00 | 3.1E-01 | -7.8E-01 | 7.9E-01 | 4 |
| LEPREL4 | 6.7E-01 | 2.7E-02 | 3.9E-01 | 4.1E-01 | -2.8E-01 | 7.1E-01 | 4 |
| MICAL3 | 1.7E+00 | 2.9E-03 | 8.4E-01 | 3.4E-01 | -8.3E-01 | 3.1E-01 | 4 |
| MED24 | 8.6E-01 | 6.5E-03 | 4.3E-01 | 4.0E-01 | -4.3E-01 | 4.3E-01 | 4 |
| MED24 | 8.6E-01 | 6.5E-03 | 4.3E-01 | 4.0E-01 | -4.3E-01 | 4.3E-01 | 4 |
| PSMD3 | 1.3E+00 | 3.4E-06 | 8.3E-01 | 1.1E-02 | -4.6E-01 | 3.2E-01 | 4 |
| HUWE1 | 1.1E+00 | 2.0E-05 | 6.7E-01 | 2.6E-02 | -3.8E-01 | 3.6E-01 | 4 |
| KAZN | 2.0E+00 | 4.0E-02 | 9.9E-01 | 6.2E-01 | -1.1E+00 | 4.7E-01 | 4 |
| ERBB2 | 1.0E+00 | 1.5E-05 | 6.2E-01 | 3.3E-02 | -4.0E-01 | 2.8E-01 | 4 |
| NVL | 1.3E+00 | 1.3E-02 | 7.4E-01 | 3.4E-01 | -5.2E-01 | 5.9E-01 | 4 |
| RBPJ | 9.7E-01 | 5.2E-03 | 7.4E-03 | 1.0E+00 | -9.6E-01 | 1.7E-02 | 4 |
| CDK12 | 1.3E+00 | 3.0E-04 | 5.7E-02 | 1.0E+00 | -1.2E+00 | 1.8E-03 | 4 |
| LAS1L | 8.2E-01 | 4.3E-02 | 5.4E-01 | 3.6E-01 | -2.8E-01 | 8.7E-01 | 4 |
| MAD2L2 | 1.1E+00 | 5.8E-03 | 4.0E-01 | 6.1E-01 | -6.6E-01 | 2.4E-01 | 4 |
| EXOSC10 | 6.6E-01 | 1.9E-02 | 4.2E-01 | 3.1E-01 | -2.4E-01 | 7.7E-01 | 4 |
| HSD3B7 | 6.0E-01 | 1.1E-02 | 2.1E-01 | 6.9E-01 | -3.9E-01 | 2.7E-01 | 4 |
| RBM25 | 8.8E-01 | 1.2E-02 | 4.1E-01 | 5.2E-01 | -4.7E-01 | 4.2E-01 | 4 |
| HOXB7 | 8.2E-01 | 3.5E-02 | 2.5E-01 | 8.6E-01 | -5.7E-01 | 3.2E-01 | 4 |
| HOXB8 | 1.0E+00 | 2.6E-02 | 3.8E-02 | 1.0E+00 | -1.0E+00 | 7.8E-02 | 4 |
| ELMSAN1 | 3.9E+00 | 4.7E-02 | 1.7E+00 | 7.4E-01 | -2.2E+00 | 4.9E-01 | 4 |
| ALDH6A1 | 1.9E+00 | 8.2E-03 | 9.6E-01 | 4.0E-01 | -8.9E-01 | 3.9E-01 | 4 |
| NA | 1.6E+00 | 1.5E-02 | 5.9E-01 | 7.0E-01 | -9.9E-01 | 2.6E-01 | 4 |
| PER1 | 2.5E+00 | 7.1E-23 | 1.4E+00 | 2.6E-06 | -1.1E+00 | 1.9E-04 | 4 |
| PER1 | 2.5E+00 | 7.1E-23 | 1.4E+00 | 2.6E-06 | -1.1E+00 | 1.9E-04 | 4 |
| HERC1 | 8.5E-01 | 9.1E-03 | 1.3E-01 | 1.0E+00 | -7.2E-01 | 7.5E-02 | 4 |
| TMED10 | 1.6E+00 | 1.8E-03 | 1.1E+00 | 1.2E-01 | -5.7E-01 | 5.7E-01 | 4 |
| TRMT1 | 9.7E-01 | 2.3E-02 | 6.5E-01 | 2.9E-01 | -3.2E-01 | 8.2E-01 | 4 |
| ZNF609 | 1.4E+00 | 1.8E-05 | 7.1E-01 | 1.0E-01 | -6.9E-01 | 1.1E-01 | 4 |
| TOB1 | 1.6E+00 | 6.4E-09 | 9.3E-01 | 3.5E-03 | -6.3E-01 | 7.4E-02 | 4 |
| NA | 1.9E+00 | 3.7E-02 | 1.3E+00 | 3.1E-01 | -5.7E-01 | 8.5E-01 | 4 |

| | | | | | | | |
|---|---|---|---|---|---|---|---|
| MYH10 | 1.4E+00 | 2.8E-03 | 4.1E-01 | 7.2E-01 | -9.5E-01 | 9.6E-02 | 4 |
| SPAG9 | 1.2E+00 | 1.5E-03 | 3.6E-01 | 6.6E-01 | -8.1E-01 | 9.3E-02 | 4 |
| SPNS1 | 1.3E+00 | 1.2E-05 | 7.3E-01 | 4.5E-02 | -5.3E-01 | 1.9E-01 | 4 |
| SEL1L | 1.3E+00 | 5.9E-03 | 5.9E-01 | 4.6E-01 | -7.0E-01 | 3.5E-01 | 4 |
| GATAD2B | 3.4E+00 | 2.7E-02 | 1.6E+00 | 6.1E-01 | -1.8E+00 | 2.4E-01 | 4 |
| NA | 2.1E+00 | 5.3E-03 | 1.2E+00 | 3.1E-01 | -9.4E-01 | 3.8E-01 | 4 |
| PSMC1 | 1.3E+00 | 6.1E-06 | 3.9E-01 | 4.5E-01 | -9.3E-01 | 6.6E-03 | 4 |
| CCDC88C | 1.5E+00 | 3.1E-03 | 2.7E-01 | 9.6E-01 | -1.3E+00 | 4.0E-02 | 4 |
| BTBD7 | 2.0E+00 | 3.5E-02 | 8.1E-01 | 7.2E-01 | -1.2E+00 | 3.8E-01 | 4 |
| SLC14A1 | 2.8E+00 | 2.7E-05 | 1.3E+00 | 1.7E-01 | -1.5E+00 | 3.6E-02 | 4 |
| RPS6KB1 | 8.4E-01 | 6.2E-03 | 5.2E-01 | 2.3E-01 | -3.3E-01 | 6.4E-01 | 4 |
| HNRNPH2 | 1.9E+00 | 4.5E-02 | 1.8E-01 | 1.0E+00 | -1.7E+00 | 1.4E-01 | 4 |
| HNRNPH2 | 1.9E+00 | 4.5E-02 | 1.8E-01 | 1.0E+00 | -1.7E+00 | 1.4E-01 | 4 |
| CDR2 | 1.8E+00 | 3.1E-02 | 1.1E+00 | 3.9E-01 | -6.7E-01 | 7.1E-01 | 4 |
| CDR2 | 1.8E+00 | 3.1E-02 | 1.1E+00 | 3.9E-01 | -6.7E-01 | 7.1E-01 | 4 |
| BCAS3 | 1.7E+00 | 6.2E-03 | 8.1E-01 | 4.5E-01 | -8.9E-01 | 3.4E-01 | 4 |
| MORF4L2 | 2.0E+00 | 8.2E-03 | 1.1E+00 | 3.7E-01 | -9.4E-01 | 4.4E-01 | 4 |
| SLC39A6 | 1.1E+00 | 4.7E-04 | 6.4E-01 | 1.2E-01 | -4.4E-01 | 4.1E-01 | 4 |
| DYNC1H1 | 1.2E+00 | 1.1E-08 | 6.8E-01 | 6.1E-03 | -5.1E-01 | 6.5E-02 | 4 |
| SIN3A | 1.9E+00 | 1.5E-05 | 9.1E-01 | 1.5E-01 | -1.0E+00 | 5.5E-02 | 4 |
| KNOP1 | 1.3E+00 | 5.4E-03 | 1.3E-01 | 1.0E+00 | -1.1E+00 | 3.6E-02 | 4 |
| ANKRD9 | 1.3E+00 | 1.4E-07 | 7.0E-01 | 2.2E-02 | -6.1E-01 | 5.6E-02 | 4 |
| RCOR1 | 1.4E+00 | 3.4E-03 | 8.6E-01 | 1.9E-01 | -5.3E-01 | 5.7E-01 | 4 |
| DSC3 | 1.3E+00 | 1.8E-03 | 6.7E-01 | 2.5E-01 | -5.8E-01 | 3.7E-01 | 4 |
| SMG1 | 9.1E-01 | 4.4E-02 | 3.7E-01 | 7.2E-01 | -5.4E-01 | 5.0E-01 | 4 |
| SMG1 | 9.1E-01 | 4.4E-02 | 3.7E-01 | 7.2E-01 | -5.4E-01 | 5.0E-01 | 4 |
| SMG1 | 9.1E-01 | 4.4E-02 | 3.7E-01 | 7.2E-01 | -5.4E-01 | 5.0E-01 | 4 |
| SMG1 | 9.1E-01 | 4.4E-02 | 3.7E-01 | 7.2E-01 | -5.4E-01 | 5.0E-01 | 4 |
| MRP1 | 2.1E+00 | 4.3E-02 | 1.2E+00 | 4.6E-01 | -8.4E-01 | 6.8E-01 | 4 |
| AP2B1 | 7.7E-01 | 2.2E-02 | 4.3E-01 | 4.2E-01 | -3.3E-01 | 6.6E-01 | 4 |
| USP14 | 7.7E-01 | 1.4E-02 | 3.0E-01 | 6.6E-01 | -4.8E-01 | 3.3E-01 | 4 |
| BRF1 | 6.4E-01 | 5.0E-02 | 3.2E-01 | 6.1E-01 | -3.3E-01 | 6.4E-01 | 4 |
| ITGA2 | 9.8E-01 | 5.9E-03 | 5.2E-01 | 3.2E-01 | -4.5E-01 | 5.0E-01 | 4 |
| ANGPTL4 | 1.6E+00 | 9.0E-09 | 1.2E-01 | 1.0E+00 | -1.5E+00 | 3.0E-07 | 4 |
| JAK1 | 7.5E-01 | 9.1E-03 | 1.9E-01 | 8.7E-01 | -5.7E-01 | 1.4E-01 | 4 |
| GTF2F1 | 7.7E-01 | 1.2E-02 | 4.1E-01 | 3.9E-01 | -3.6E-01 | 5.6E-01 | 4 |
| RALBP1 | 6.0E-01 | 3.8E-02 | 3.4E-01 | 4.6E-01 | -2.6E-01 | 7.3E-01 | 4 |
| ALDOC | 8.2E-01 | 8.7E-04 | 4.0E-01 | 2.7E-01 | -4.2E-01 | 2.7E-01 | 4 |
| PDXDC1 | 1.5E+00 | 1.4E-02 | 7.7E-02 | 1.0E+00 | -1.4E+00 | 4.9E-02 | 4 |
| RAI14 | 1.3E+00 | 1.6E-02 | 4.5E-01 | 7.6E-01 | -8.8E-01 | 2.8E-01 | 4 |
| ERCC4 | 1.6E+00 | 3.0E-02 | 2.5E-01 | 1.0E+00 | -1.3E+00 | 1.5E-01 | 4 |
| GSPT1 | 1.3E+00 | 2.3E-05 | 3.6E-01 | 5.3E-01 | -8.9E-01 | 1.0E-02 | 4 |
| MOSPD1 | 8.2E-01 | 6.8E-03 | 1.8E-01 | 9.1E-01 | -6.4E-01 | 1.0E-01 | 4 |
| KDM4B | 8.6E-01 | 5.3E-04 | 2.6E-01 | 6.2E-01 | -6.0E-01 | 5.5E-02 | 4 |
| CCDC11 | 1.2E+00 | 2.3E-02 | 3.0E-01 | 9.3E-01 | -9.1E-01 | 1.8E-01 | 4 |
| PLIN3 | 1.3E+00 | 1.1E-03 | 2.8E-01 | 8.6E-01 | -1.0E+00 | 3.3E-02 | 4 |
| STAP2 | 1.0E+00 | 1.5E-03 | 6.2E-01 | 1.6E-01 | -4.1E-01 | 5.1E-01 | 4 |
| GLYR1 | 8.7E-01 | 4.4E-03 | 1.0E-01 | 1.0E+00 | -7.7E-01 | 3.7E-02 | 4 |
| BGN | 1.3E+00 | 3.7E-03 | 7.6E-01 | 2.0E-01 | -4.9E-01 | 5.9E-01 | 4 |
| HMG20B | 1.6E+00 | 4.8E-02 | 4.8E-01 | 9.1E-01 | -1.2E+00 | 3.1E-01 | 4 |
| NA | 1.2E+00 | 1.1E-04 | 7.5E-01 | 6.6E-02 | -4.9E-01 | 3.6E-01 | 4 |
| NA | 1.2E+00 | 1.1E-04 | 7.5E-01 | 6.6E-02 | -4.9E-01 | 3.6E-01 | 4 |
| NA | 1.2E+00 | 1.1E-04 | 7.5E-01 | 6.6E-02 | -4.9E-01 | 3.6E-01 | 4 |
| NA | 1.2E+00 | 1.1E-04 | 7.5E-01 | 6.6E-02 | -4.9E-01 | 3.6E-01 | 4 |
| BCAP31 | 1.2E+00 | 2.3E-05 | 7.8E-01 | 2.0E-02 | -4.0E-01 | 4.3E-01 | 4 |
| SGTA | 6.5E-01 | 3.3E-02 | 3.5E-01 | 4.9E-01 | -3.0E-01 | 6.7E-01 | 4 |
| RNPS1 | 1.5E+00 | 3.4E-03 | 7.7E-01 | 3.4E-01 | -7.3E-01 | 3.3E-01 | 4 |
| AP3D1 | 1.7E+00 | 2.0E-08 | 7.6E-01 | 4.4E-02 | -8.9E-01 | 1.1E-02 | 4 |
| TUBB4B | 8.5E-01 | 4.9E-02 | 2.9E-01 | 8.2E-01 | -5.6E-01 | 4.3E-01 | 4 |
| CEP104 | 1.5E+00 | 3.3E-02 | 6.6E-01 | 6.8E-01 | -8.9E-01 | 4.4E-01 | 4 |
| NA | 1.1E+00 | 4.3E-06 | 3.9E-01 | 2.8E-01 | -6.9E-01 | 1.4E-02 | 4 |
| MAPK8IP3 | 2.0E+00 | 9.0E-09 | 9.8E-01 | 2.4E-02 | -1.0E+00 | 1.1E-02 | 4 |
| TSR3 | 6.1E-01 | 6.6E-03 | 3.8E-01 | 2.2E-01 | -2.3E-01 | 6.8E-01 | 4 |
| WDR24 | 9.0E-01 | 2.6E-02 | 1.1E-01 | 1.0E+00 | -7.9E-01 | 1.2E-01 | 4 |
| PIGQ | 1.3E+00 | 2.3E-03 | 2.1E-01 | 9.7E-01 | -1.1E+00 | 3.1E-02 | 4 |
| RERE | 1.4E+00 | 6.1E-06 | 5.6E-01 | 2.3E-01 | -8.8E-01 | 2.3E-02 | 4 |
| REXO4 | 1.6E+00 | 1.7E-02 | 2.2E-01 | 1.0E+00 | -1.4E+00 | 9.4E-02 | 4 |
| KIF1B | 7.3E-01 | 7.9E-04 | 2.6E-01 | 5.2E-01 | -4.7E-01 | 1.0E-01 | 4 |
| TSC1 | 2.2E+00 | 3.0E-02 | 7.0E-01 | 8.4E-01 | -1.5E+00 | 2.5E-01 | 4 |
| TTF1 | 1.2E+00 | 1.7E-03 | 2.2E-01 | 9.2E-01 | -9.7E-01 | 3.5E-02 | 4 |
| SETX | 1.3E+00 | 1.3E-03 | 5.6E-01 | 3.9E-01 | -7.2E-01 | 2.1E-01 | 4 |
| ZCCHC14 | 3.2E+00 | 1.1E-02 | 1.5E+00 | 5.2E-01 | -1.7E+00 | 1.8E-01 | 4 |
| WWOX | 1.6E+00 | 4.5E-02 | 8.6E-01 | 5.4E-01 | -7.3E-01 | 6.5E-01 | 4 |
| C9orf114 | 1.9E+00 | 7.2E-06 | 5.2E-01 | 5.4E-01 | -1.4E+00 | 3.3E-03 | 4 |
| NA | 1.6E+00 | 1.7E-03 | 7.8E-01 | 3.3E-01 | -8.4E-01 | 2.8E-01 | 4 |
| NA | 1.6E+00 | 1.7E-03 | 7.8E-01 | 3.3E-01 | -8.4E-01 | 2.8E-01 | 4 |
| NA | 1.6E+00 | 1.7E-03 | 7.8E-01 | 3.3E-01 | -8.4E-01 | 2.8E-01 | 4 |
| NA | 1.6E+00 | 1.7E-03 | 7.8E-01 | 3.3E-01 | -8.4E-01 | 2.8E-01 | 4 |
| NA | 1.6E+00 | 1.7E-03 | 7.8E-01 | 3.3E-01 | -8.4E-01 | 2.8E-01 | 4 |
| NA | 1.6E+00 | 1.7E-03 | 7.8E-01 | 3.3E-01 | -8.4E-01 | 2.8E-01 | 4 |
| NA | 1.6E+00 | 1.7E-03 | 7.8E-01 | 3.3E-01 | -8.4E-01 | 2.8E-01 | 4 |
| NA | 1.6E+00 | 1.7E-03 | 7.8E-01 | 3.3E-01 | -8.4E-01 | 2.8E-01 | 4 |
| NA | 1.6E+00 | 1.7E-03 | 7.8E-01 | 3.3E-01 | -8.4E-01 | 2.8E-01 | 4 |
| NA | 1.6E+00 | 1.7E-03 | 7.8E-01 | 3.3E-01 | -8.4E-01 | 2.8E-01 | 4 |
| NA | 1.6E+00 | 1.7E-03 | 7.8E-01 | 3.3E-01 | -8.4E-01 | 2.8E-01 | 4 |
| NA | 1.6E+00 | 1.7E-03 | 7.8E-01 | 3.3E-01 | -8.4E-01 | 2.8E-01 | 4 |
| NA | 1.6E+00 | 1.7E-03 | 7.8E-01 | 3.3E-01 | -8.4E-01 | 2.8E-01 | 4 |
| NA | 1.6E+00 | 1.7E-03 | 7.8E-01 | 3.3E-01 | -8.4E-01 | 2.8E-01 | 4 |
| NA | 1.6E+00 | 1.7E-03 | 7.8E-01 | 3.3E-01 | -8.4E-01 | 2.8E-01 | 4 |
| NA | 1.6E+00 | 1.7E-03 | 7.8E-01 | 3.3E-01 | -8.4E-01 | 2.8E-01 | 4 |
| NA | 1.6E+00 | 1.7E-03 | 7.8E-01 | 3.3E-01 | -8.4E-01 | 2.8E-01 | 4 |
| NA | 1.6E+00 | 1.7E-03 | 7.8E-01 | 3.3E-01 | -8.4E-01 | 2.8E-01 | 4 |
| NA | 1.6E+00 | 1.7E-03 | 7.8E-01 | 3.3E-01 | -8.4E-01 | 2.8E-01 | 4 |
| NA | 1.6E+00 | 1.7E-03 | 7.8E-01 | 3.3E-01 | -8.4E-01 | 2.8E-01 | 4 |

| | | | | | | | |
|---|---|---|---|---|---|---|---|
| NA | 1.6E+00 | 1.7E-03 | 7.8E-01 | 3.3E-01 | -8.4E-01 | 2.8E-01 | 4 |
| NA | 1.6E+00 | 1.7E-03 | 7.8E-01 | 3.3E-01 | -8.4E-01 | 2.8E-01 | 4 |
| NA | 1.6E+00 | 1.7E-03 | 7.8E-01 | 3.3E-01 | -8.4E-01 | 2.8E-01 | 4 |
| NA | 1.6E+00 | 1.7E-03 | 7.8E-01 | 3.3E-01 | -8.4E-01 | 2.8E-01 | 4 |
| NA | 1.6E+00 | 1.7E-03 | 7.8E-01 | 3.3E-01 | -8.4E-01 | 2.8E-01 | 4 |
| NA | 1.6E+00 | 1.7E-03 | 7.8E-01 | 3.3E-01 | -8.4E-01 | 2.8E-01 | 4 |
| NA | 1.6E+00 | 1.7E-03 | 7.8E-01 | 3.3E-01 | -8.4E-01 | 2.8E-01 | 4 |
| NA | 1.6E+00 | 1.7E-03 | 7.8E-01 | 3.3E-01 | -8.4E-01 | 2.8E-01 | 4 |
| NA | 1.6E+00 | 1.7E-03 | 7.8E-01 | 3.3E-01 | -8.4E-01 | 2.8E-01 | 4 |
| NA | 1.6E+00 | 1.7E-03 | 7.8E-01 | 3.3E-01 | -8.4E-01 | 2.8E-01 | 4 |
| NA | 1.6E+00 | 1.7E-03 | 7.8E-01 | 3.3E-01 | -8.4E-01 | 2.8E-01 | 4 |
| NA | 1.6E+00 | 1.7E-03 | 7.8E-01 | 3.3E-01 | -8.4E-01 | 2.8E-01 | 4 |
| NA | 1.6E+00 | 1.7E-03 | 7.8E-01 | 3.3E-01 | -8.4E-01 | 2.8E-01 | 4 |
| NA | 1.6E+00 | 1.7E-03 | 7.8E-01 | 3.3E-01 | -8.4E-01 | 2.8E-01 | 4 |
| NA | 1.6E+00 | 1.7E-03 | 7.8E-01 | 3.3E-01 | -8.4E-01 | 2.8E-01 | 4 |
| NA | 1.6E+00 | 1.7E-03 | 7.8E-01 | 3.3E-01 | -8.4E-01 | 2.8E-01 | 4 |
| NA | 1.6E+00 | 1.7E-03 | 7.8E-01 | 3.3E-01 | -8.4E-01 | 2.8E-01 | 4 |
| NA | 1.6E+00 | 1.7E-03 | 7.8E-01 | 3.3E-01 | -8.4E-01 | 2.8E-01 | 4 |
| NA | 1.6E+00 | 1.7E-03 | 7.8E-01 | 3.3E-01 | -8.4E-01 | 2.8E-01 | 4 |
| NA | 1.6E+00 | 1.7E-03 | 7.8E-01 | 3.3E-01 | -8.4E-01 | 2.8E-01 | 4 |
| NA | 1.6E+00 | 1.7E-03 | 7.8E-01 | 3.3E-01 | -8.4E-01 | 2.8E-01 | 4 |
| NA | 1.6E+00 | 1.7E-03 | 7.8E-01 | 3.3E-01 | -8.4E-01 | 2.8E-01 | 4 |
| NA | 1.6E+00 | 1.7E-03 | 7.8E-01 | 3.3E-01 | -8.4E-01 | 2.8E-01 | 4 |
| NA | 1.6E+00 | 1.7E-03 | 7.8E-01 | 3.3E-01 | -8.4E-01 | 2.8E-01 | 4 |
| NA | 1.6E+00 | 1.7E-03 | 7.8E-01 | 3.3E-01 | -8.4E-01 | 2.8E-01 | 4 |
| NA | 1.6E+00 | 1.7E-03 | 7.8E-01 | 3.3E-01 | -8.4E-01 | 2.8E-01 | 4 |
| NA | 1.6E+00 | 1.7E-03 | 7.8E-01 | 3.3E-01 | -8.4E-01 | 2.8E-01 | 4 |
| NA | 1.6E+00 | 1.7E-03 | 7.8E-01 | 3.3E-01 | -8.4E-01 | 2.8E-01 | 4 |
| NA | 1.6E+00 | 1.7E-03 | 7.8E-01 | 3.3E-01 | -8.4E-01 | 2.8E-01 | 4 |
| NA | 1.6E+00 | 1.7E-03 | 7.8E-01 | 3.3E-01 | -8.4E-01 | 2.8E-01 | 4 |
| NA | 1.6E+00 | 1.7E-03 | 7.8E-01 | 3.3E-01 | -8.4E-01 | 2.8E-01 | 4 |
| NA | 1.6E+00 | 1.7E-03 | 7.8E-01 | 3.3E-01 | -8.4E-01 | 2.8E-01 | 4 |
| NA | 1.6E+00 | 1.7E-03 | 7.8E-01 | 3.3E-01 | -8.4E-01 | 2.8E-01 | 4 |
| NA | 1.6E+00 | 1.7E-03 | 7.8E-01 | 3.3E-01 | -8.4E-01 | 2.8E-01 | 4 |
| NA | 1.6E+00 | 1.7E-03 | 7.8E-01 | 3.3E-01 | -8.4E-01 | 2.8E-01 | 4 |
| NA | 1.6E+00 | 1.7E-03 | 7.8E-01 | 3.3E-01 | -8.4E-01 | 2.8E-01 | 4 |
| NA | 1.6E+00 | 1.7E-03 | 7.8E-01 | 3.3E-01 | -8.4E-01 | 2.8E-01 | 4 |
| NA | 1.6E+00 | 1.7E-03 | 7.8E-01 | 3.3E-01 | -8.4E-01 | 2.8E-01 | 4 |
| GOLGA1 | 1.6E+00 | 2.6E-02 | 2.9E-01 | 1.0E+00 | -1.3E+00 | 1.5E-01 | 4 |
| PKN2 | 8.1E-01 | 5.4E-03 | 3.4E-01 | 5.1E-01 | -4.6E-01 | 2.9E-01 | 4 |
| NQO1 | 1.4E+00 | 1.1E-02 | 8.4E-01 | 2.9E-01 | -5.4E-01 | 6.5E-01 | 4 |
| CTCF | 1.2E+00 | 6.4E-03 | 3.8E-01 | 7.6E-01 | -8.7E-01 | 1.5E-01 | 4 |
| NA | 1.8E+00 | 3.9E-02 | 6.4E-01 | 7.8E-01 | -1.1E+00 | 3.9E-01 | 4 |
| NA | 4.5E+00 | 1.4E-02 | 3.2E+00 | 1.9E-01 | -1.3E+00 | 7.0E-01 | 4 |
| NA | 3.7E+00 | 4.4E-02 | 1.7E+00 | 6.6E-01 | -2.0E+00 | 3.6E-01 | 4 |
| NA | 1.2E+00 | 7.0E-03 | 7.3E-01 | 2.4E-01 | -4.4E-01 | 6.4E-01 | 4 |
| MT-ND4 | 7.7E-01 | 3.2E-03 | 2.2E-01 | 7.4E-01 | -5.5E-01 | 1.1E-01 | 4 |
| NA | 1.7E+00 | 4.4E-02 | 5.0E-01 | 9.0E-01 | -1.2E+00 | 3.0E-01 | 4 |
| NFE2L1 | 1.9E+00 | 1.6E-06 | 5.2E-01 | 4.8E-01 | -1.4E+00 | 1.6E-03 | 4 |
| NA | 1.1E+00 | 1.7E-04 | 4.9E-01 | 2.5E-01 | -5.8E-01 | 1.3E-01 | 4 |
| PRSS22 | 7.8E-01 | 2.3E-02 | 4.0E-01 | 4.8E-01 | -3.7E-01 | 6.0E-01 | 4 |
| TGM3 | 4.1E+00 | 4.8E-02 | 2.3E+00 | 5.1E-01 | -1.8E+00 | 5.6E-01 | 4 |
| ZBTB7A | 1.6E+00 | 3.5E-05 | 5.2E-02 | 1.0E+00 | -1.6E+00 | 1.9E-04 | 4 |
| MPHOSPH6 | 1.4E+00 | 3.1E-03 | 7.3E-02 | 1.0E+00 | -1.3E+00 | 1.7E-02 | 4 |
| NA | 1.4E+00 | 3.1E-02 | 6.8E-01 | 5.9E-01 | -7.3E-01 | 5.0E-01 | 4 |
| NA | 1.4E+00 | 3.1E-02 | 6.8E-01 | 5.9E-01 | -7.3E-01 | 5.0E-01 | 4 |
| NA | 1.4E+00 | 3.1E-02 | 6.8E-01 | 5.9E-01 | -7.3E-01 | 5.0E-01 | 4 |
| NA | 1.4E+00 | 3.1E-02 | 6.8E-01 | 5.9E-01 | -7.3E-01 | 5.0E-01 | 4 |
| NA | 1.4E+00 | 3.1E-02 | 6.8E-01 | 5.9E-01 | -7.3E-01 | 5.0E-01 | 4 |
| NA | 1.4E+00 | 3.1E-02 | 6.8E-01 | 5.9E-01 | -7.3E-01 | 5.0E-01 | 4 |
| KPNA4 | 1.2E+00 | 1.9E-03 | 4.4E-01 | 5.8E-01 | -7.9E-01 | 1.3E-01 | 4 |
| C5orf24 | 1.4E+00 | 4.9E-03 | 6.1E-01 | 5.0E-01 | -7.8E-01 | 2.8E-01 | 4 |
| IGFBP6 | 9.5E-01 | 1.7E-02 | 3.9E-01 | 6.3E-01 | -5.6E-01 | 3.8E-01 | 4 |
| MRPS25 | 6.8E-01 | 1.4E-02 | 2.8E-01 | 6.1E-01 | -4.0E-01 | 3.8E-01 | 4 |
| JAGN1 | 1.2E+00 | 1.2E-04 | 9.6E-02 | 1.0E+00 | -1.1E+00 | 1.4E-03 | 4 |
| NA | 1.9E+00 | 2.1E-09 | 9.8E-01 | 9.2E-03 | -8.8E-01 | 1.8E-02 | 4 |
| PTRF | 1.4E+00 | 3.6E-05 | 3.3E-01 | 6.7E-01 | -1.1E+00 | 7.3E-03 | 4 |
| NA | 2.3E+00 | 3.3E-07 | 1.3E+00 | 2.0E-02 | -1.0E+00 | 4.4E-02 | 4 |
| NA | 2.1E+00 | 1.7E-06 | 9.0E-01 | 1.6E-01 | -1.2E+00 | 1.3E-02 | 4 |
| RHOV | 1.7E+00 | 4.0E-02 | 9.5E-01 | 4.8E-01 | -7.2E-01 | 7.0E-01 | 4 |
| UBE2S | 8.5E-01 | 5.2E-03 | 3.5E-02 | 1.0E+00 | -8.2E-01 | 2.2E-02 | 4 |
| NA | 1.4E+00 | 4.0E-04 | 4.2E-01 | 6.2E-01 | -9.6E-01 | 4.0E-02 | 4 |
| NOM1 | 1.2E+00 | 3.3E-02 | 5.8E-01 | 6.0E-01 | -6.4E-01 | 5.3E-01 | 4 |
| NA | 1.1E+00 | 8.2E-03 | 6.9E-01 | 2.8E-01 | -4.6E-01 | 6.1E-01 | 4 |
| NA | 4.4E+00 | 2.2E-02 | 2.2E+00 | 5.5E-01 | -2.1E+00 | 4.4E-01 | 4 |
| TGIF2 | 1.0E+00 | 6.1E-03 | 1.3E-01 | 1.0E+00 | -8.8E-01 | 4.8E-02 | 4 |
| ABCF1 | 1.3E+00 | 3.3E-06 | 7.8E-01 | 2.8E-02 | -5.6E-01 | 1.7E-01 | 4 |
| HSPB8 | 1.1E+00 | 5.3E-04 | 2.8E-01 | 6.9E-01 | -7.8E-01 | 4.1E-02 | 4 |
| TTC33 | 3.9E+00 | 4.1E-02 | 2.9E+00 | 2.5E-01 | -9.9E-01 | 8.2E-01 | 4 |
| NA | 1.6E+00 | 6.7E-06 | 3.7E-01 | 6.6E-01 | -1.2E+00 | 1.7E-03 | 4 |
| NA | 7.8E-01 | 4.7E-02 | 2.2E-01 | 9.1E-01 | -5.6E-01 | 3.4E-01 | 4 |
| NA | 1.1E+00 | 2.5E-04 | 4.8E-01 | 3.3E-01 | -6.6E-01 | 1.1E-01 | 4 |
| YWHAH | 1.8E+00 | 8.7E-06 | 5.8E-01 | 3.9E-01 | -1.2E+00 | 1.0E-02 | 4 |
| K-RAS | 2.3E+00 | 1.2E-04 | 1.4E+00 | 8.8E-02 | -9.7E-01 | 2.2E-01 | 4 |
| BHLHE41 | 1.3E+00 | 3.0E-05 | 5.3E-01 | 2.8E-01 | -7.9E-01 | 4.3E-02 | 4 |
| FAM222B | 1.3E+00 | 2.9E-05 | 4.2E-01 | 4.2E-01 | -8.4E-01 | 2.1E-02 | 4 |
| NA | 1.5E+00 | 3.3E-02 | 9.2E-01 | 3.9E-01 | -5.6E-01 | 7.5E-01 | 4 |
| PIGA | 1.2E+00 | 3.1E-03 | 5.2E-01 | 4.9E-01 | -7.1E-01 | 2.3E-01 | 4 |
| C11orf68 | 1.5E+00 | 2.1E-06 | 3.5E-01 | 6.1E-01 | -1.2E+00 | 9.3E-04 | 4 |
| PMEPA1 | 6.7E-01 | 2.0E-02 | 3.5E-01 | 4.5E-01 | -3.2E-01 | 6.1E-01 | 4 |
| EHF | 7.7E-01 | 6.8E-03 | 2.8E-01 | 6.2E-01 | -4.8E-01 | 2.5E-01 | 4 |
| VGLL4 | 1.5E+00 | 3.9E-07 | 2.4E-01 | 8.0E-01 | -1.2E+00 | 6.4E-05 | 4 |
| NA | 1.6E+00 | 1.9E-02 | 2.8E-01 | 1.0E+00 | -1.3E+00 | 1.2E-01 | 4 |
| FAM46A | 6.4E-01 | 3.7E-03 | 2.9E-02 | 1.0E+00 | -6.1E-01 | 1.8E-02 | 4 |

| | | | | | | | |
|---|---|---|---|---|---|---|---|
| IRGQ | 1.3E+00 | 2.3E-06 | 3.0E-01 | 6.2E-01 | -9.9E-01 | 9.7E-04 | 4 |
| OGFR | 1.6E+00 | 7.4E-08 | 7.9E-01 | 3.0E-02 | -7.8E-01 | 3.0E-02 | 4 |
| MESDC2 | 1.3E+00 | 2.0E-03 | 1.7E-01 | 1.0E+00 | -1.1E+00 | 2.1E-02 | 4 |
| REEP3 | 6.2E-01 | 3.3E-02 | 3.3E-01 | 5.0E-01 | -2.9E-01 | 6.6E-01 | 4 |
| ZNF775 | 1.2E+00 | 3.3E-05 | 9.7E-02 | 1.0E+00 | -1.1E+00 | 4.7E-04 | 4 |
| NA | 1.9E+00 | 3.3E-06 | 1.1E+00 | 2.7E-02 | -8.0E-01 | 1.5E-01 | 4 |
| NA | 9.9E-01 | 3.3E-03 | 1.9E-01 | 9.4E-01 | -8.0E-01 | 5.3E-02 | 4 |
| NA | 2.0E+00 | 5.1E-03 | 3.7E-01 | 9.6E-01 | -1.6E+00 | 5.1E-02 | 4 |
| TXNL4A | 4.9E-01 | 1.8E-01 | 8.0E-01 | 2.8E-02 | 3.1E-01 | 7.1E-01 | 5 |
| NA | 1.5E+00 | 2.2E-04 | 1.0E+00 | 4.0E-02 | -4.6E-01 | 5.4E-01 | 5 |
| NA | 1.5E+00 | 2.2E-04 | 1.0E+00 | 4.0E-02 | -4.6E-01 | 5.4E-01 | 5 |
| NA | 1.5E+00 | 2.2E-04 | 1.0E+00 | 4.0E-02 | -4.6E-01 | 5.4E-01 | 5 |
| NA | 1.5E+00 | 2.2E-04 | 1.0E+00 | 4.0E-02 | -4.6E-01 | 5.4E-01 | 5 |
| KDSR | 3.4E-01 | 3.2E-01 | 6.2E-01 | 4.6E-02 | 2.8E-01 | 6.4E-01 | 5 |
| SMARCC2 | 2.3E+00 | 1.6E-04 | 2.1E+00 | 1.7E-03 | -1.9E-01 | 9.9E-01 | 5 |
| NARS | 5.3E-01 | 1.7E-03 | 6.3E-01 | 5.8E-04 | 9.8E-02 | 9.7E-01 | 5 |
| RPS12 | 7.3E-01 | 2.7E-04 | 1.3E+00 | 7.7E-10 | 5.3E-01 | 3.4E-02 | 5 |
| CCDC112 | 1.3E+00 | 1.0E-04 | 1.4E+00 | 6.4E-05 | 1.3E-01 | 9.9E-01 | 5 |
| R3HDM2 | 1.3E+00 | 1.3E-03 | 1.1E+00 | 2.6E-02 | -2.4E-01 | 9.5E-01 | 5 |
| TMED9 | 1.1E+00 | 1.5E-05 | 8.8E-01 | 2.3E-03 | -2.2E-01 | 8.2E-01 | 5 |
| N4BP3 | 7.5E-01 | 9.6E-02 | 1.0E+00 | 2.9E-02 | 2.7E-01 | 9.0E-01 | 5 |
| OS9 | 8.2E-01 | 1.1E-04 | 9.8E-01 | 1.5E-05 | 1.6E-01 | 8.7E-01 | 5 |
| PHACTR2 | 7.5E-01 | 6.2E-02 | 1.1E+00 | 1.2E-02 | 3.0E-01 | 8.2E-01 | 5 |
| ADAMTS2 | 1.4E+00 | 1.4E-02 | 1.9E+00 | 6.9E-04 | 5.6E-01 | 5.6E-01 | 5 |
| STXBP5 | 1.3E+00 | 2.1E-03 | 8.5E-01 | 1.1E-01 | -4.1E-01 | 6.4E-01 | 5 |
| MGAT4B | 5.2E-01 | 4.5E-02 | 9.1E-01 | 4.5E-04 | 3.9E-01 | 3.1E-01 | 5 |
| DDR1 | 6.2E-01 | 1.7E-03 | 9.0E-01 | 1.0E-05 | 2.9E-01 | 4.0E-01 | 5 |
| DDR1 | 6.2E-01 | 1.7E-03 | 9.0E-01 | 1.0E-05 | 2.9E-01 | 4.0E-01 | 5 |
| CLIC1 | 1.4E+00 | 6.0E-07 | 1.1E+00 | 8.2E-04 | -3.5E-01 | 5.8E-01 | 5 |
| SNX9 | 1.4E+00 | 6.1E-04 | 1.2E+00 | 1.0E-02 | -1.8E-01 | 9.9E-01 | 5 |
| DENND6B | 1.1E+00 | 9.5E-04 | 1.3E+00 | 3.0E-04 | 2.0E-01 | 9.5E-01 | 5 |
| VARS | 8.6E-01 | 3.5E-04 | 8.5E-01 | 1.5E-03 | -1.0E-02 | 9.9E-01 | 5 |
| EHMT2 | 8.4E-01 | 2.5E-03 | 5.8E-01 | 1.1E-01 | -2.6E-01 | 7.3E-01 | 5 |
| TRABD | 8.3E-01 | 1.5E-03 | 7.3E-01 | 1.7E-02 | -1.1E-01 | 9.9E-01 | 5 |
| CERK | 7.5E-01 | 4.1E-04 | 6.9E-01 | 3.9E-03 | -5.6E-02 | 9.9E-01 | 5 |
| CERK | 7.5E-01 | 4.1E-04 | 6.9E-01 | 3.9E-03 | -5.6E-02 | 9.9E-01 | 5 |
| ATF6B | 8.3E-01 | 3.1E-03 | 8.4E-01 | 7.3E-03 | 1.5E-02 | 9.9E-01 | 5 |
| NA | 8.8E-01 | 3.3E-04 | 9.2E-01 | 6.7E-04 | 3.7E-02 | 9.9E-01 | 5 |
| RXRB | 6.6E-01 | 2.9E-02 | 4.9E-01 | 2.3E-01 | -1.8E-01 | 9.5E-01 | 5 |
| CYB5R3 | 5.7E-01 | 3.6E-03 | 8.2E-01 | 6.1E-05 | 2.5E-01 | 5.0E-01 | 5 |
| RGL2 | 8.9E-01 | 5.5E-04 | 1.2E+00 | 8.7E-06 | 3.2E-01 | 5.4E-01 | 5 |
| SREBF2 | 9.0E-01 | 5.7E-04 | 6.5E-01 | 4.1E-02 | -2.5E-01 | 7.3E-01 | 5 |
| NCOA7 | 1.2E+00 | 4.2E-03 | 8.4E-01 | 1.0E-01 | -3.2E-01 | 8.2E-01 | 5 |
| FAM49B | 8.3E-01 | 2.2E-02 | 8.9E-01 | 2.7E-02 | 6.8E-02 | 9.9E-01 | 5 |
| L3MBTL3 | 2.7E+00 | 8.7E-03 | 2.0E+00 | 1.3E-01 | -7.1E-01 | 7.3E-01 | 5 |
| TRIM11 | 9.4E-01 | 9.6E-03 | 9.7E-01 | 1.9E-02 | 2.6E-02 | 9.9E-01 | 5 |
| MRPL55 | 8.2E-01 | 9.9E-04 | 9.9E-01 | 2.1E-04 | 1.7E-01 | 9.1E-01 | 5 |
| LEMD2 | 8.3E-01 | 3.3E-03 | 7.9E-01 | 1.6E-02 | -4.4E-02 | 9.9E-01 | 5 |
| C6orf106 | 9.6E-01 | 2.7E-06 | 7.7E-01 | 7.1E-04 | -1.8E-01 | 7.9E-01 | 5 |
| EEF1D | 7.5E-01 | 1.0E-03 | 9.6E-01 | 7.3E-05 | 2.1E-01 | 7.6E-01 | 5 |
| RPL10A | 1.5E+00 | 1.3E-07 | 1.4E+00 | 2.8E-06 | -6.2E-02 | 9.9E-01 | 5 |
| MAPK13 | 9.4E-01 | 1.0E-04 | 8.9E-01 | 8.2E-04 | -4.1E-02 | 9.9E-01 | 5 |
| POLR2F | 4.0E-01 | 2.3E-01 | 6.5E-01 | 4.3E-02 | 2.5E-01 | 7.4E-01 | 5 |
| MPST | 1.0E+00 | 1.5E-04 | 1.1E+00 | 4.9E-04 | 1.0E-02 | 9.9E-01 | 5 |
| GPAA1 | 7.0E-01 | 6.1E-03 | 8.4E-01 | 2.6E-03 | 1.3E-01 | 9.9E-01 | 5 |
| AHCYL2 | 8.4E-01 | 2.3E-01 | 1.4E+00 | 2.5E-02 | 6.0E-01 | 5.4E-01 | 5 |
| TNPO3 | 5.5E-01 | 9.6E-03 | 6.5E-01 | 5.4E-03 | 9.7E-02 | 9.9E-01 | 5 |
| EIF3D | 7.7E-01 | 2.2E-04 | 9.9E-01 | 8.1E-06 | 2.2E-01 | 6.9E-01 | 5 |
| HSF1 | 6.9E-01 | 2.7E-04 | 6.1E-01 | 4.6E-03 | -7.7E-02 | 9.9E-01 | 5 |
| TFEB | 8.2E-01 | 4.3E-02 | 6.4E-01 | 2.3E-01 | -1.8E-01 | 9.9E-01 | 5 |
| CYHR1 | 1.7E+00 | 3.7E-03 | 1.4E+00 | 3.5E-02 | -2.3E-01 | 9.9E-01 | 5 |
| IMPDH1 | 8.2E-01 | 2.3E-04 | 8.6E-01 | 4.5E-04 | 3.8E-02 | 9.9E-01 | 5 |
| RBM28 | 7.7E-01 | 1.1E-03 | 8.6E-01 | 8.0E-04 | 9.3E-02 | 9.9E-01 | 5 |
| ARF5 | 6.5E-01 | 6.6E-03 | 7.5E-01 | 4.3E-03 | 9.9E-02 | 9.9E-01 | 5 |
| WFS1 | 1.4E+00 | 7.6E-08 | 1.0E+00 | 8.7E-04 | -4.2E-01 | 3.5E-01 | 5 |
| TOPORS | 8.9E-01 | 1.4E-02 | 8.3E-01 | 5.0E-02 | -5.9E-02 | 9.9E-01 | 5 |
| B4GALT1 | 6.8E-01 | 7.8E-04 | 6.6E-01 | 3.3E-03 | -1.5E-02 | 9.9E-01 | 5 |
| DCTN3 | 6.4E-01 | 3.8E-02 | 7.1E-01 | 3.9E-02 | 7.1E-02 | 9.9E-01 | 5 |
| SLC29A1 | 7.3E-01 | 2.6E-02 | 4.9E-01 | 2.8E-01 | -2.4E-01 | 8.7E-01 | 5 |
| SUPT3H | 6.5E-01 | 6.7E-03 | 6.1E-01 | 2.7E-02 | -3.7E-02 | 9.9E-01 | 5 |
| KIAA0020 | 5.5E-01 | 2.5E-02 | 6.4E-01 | 1.8E-02 | 9.1E-02 | 9.9E-01 | 5 |
| TESK1 | 1.1E+00 | 6.0E-03 | 9.9E-01 | 2.7E-02 | -7.2E-02 | 9.9E-01 | 5 |
| CCDC107 | 6.8E-01 | 5.2E-02 | 9.4E-01 | 1.0E-02 | 2.6E-01 | 8.2E-01 | 5 |
| FAM120A | 8.9E-01 | 4.1E-05 | 8.7E-01 | 2.3E-04 | -1.3E-02 | 9.9E-01 | 5 |
| CHMP2A | 5.2E-01 | 3.7E-02 | 6.2E-01 | 2.2E-02 | 1.0E-01 | 9.9E-01 | 5 |
| NA | 1.8E+00 | 8.2E-04 | 1.9E+00 | 1.8E-03 | 3.6E-02 | 9.9E-01 | 5 |
| XPA | 1.6E+00 | 1.2E-05 | 1.4E+00 | 1.1E-03 | -2.6E-01 | 8.7E-01 | 5 |
| PHF3 | 8.2E-01 | 1.3E-02 | 7.9E-01 | 3.9E-02 | -3.2E-02 | 9.9E-01 | 5 |
| TMEM190 | 1.4E+00 | 7.6E-02 | 1.7E+00 | 4.2E-02 | 3.1E-01 | 9.9E-01 | 5 |
| CALM2 | 1.3E+00 | 9.7E-04 | 9.3E-01 | 6.1E-02 | -3.8E-01 | 7.0E-01 | 5 |
| CALM2 | 1.3E+00 | 9.7E-04 | 9.3E-01 | 6.1E-02 | -3.8E-01 | 7.0E-01 | 5 |
| CALM2 | 1.3E+00 | 9.7E-04 | 9.3E-01 | 6.1E-02 | -3.8E-01 | 7.0E-01 | 5 |
| RPS9 | 5.0E-01 | 3.3E-02 | 7.0E-01 | 4.1E-03 | 2.0E-01 | 7.6E-01 | 5 |
| LENG1 | 8.2E-01 | 1.9E-03 | 7.1E-01 | 2.1E-02 | -1.1E-01 | 9.9E-01 | 5 |
| PRPF31 | 5.1E-01 | 2.0E-02 | 7.8E-01 | 5.2E-04 | 2.7E-01 | 4.9E-01 | 5 |
| RAD23B | 2.5E+00 | 2.0E-03 | 2.2E+00 | 2.2E-02 | -3.7E-01 | 9.5E-01 | 5 |
| NA | 5.8E-01 | 1.2E-01 | 1.2E+00 | 7.1E-04 | 6.0E-01 | 1.7E-01 | 5 |
| TAX1BP1 | 8.4E-01 | 3.9E-02 | 8.6E-01 | 6.4E-02 | 2.4E-02 | 9.9E-01 | 5 |
| RLF | 7.9E-01 | 2.0E-03 | 5.9E-01 | 6.2E-02 | -1.9E-01 | 8.4E-01 | 5 |
| UBE2J1 | 6.5E-01 | 5.7E-02 | 7.5E-01 | 4.8E-02 | 9.9E-02 | 9.9E-01 | 5 |
| SH3BP5L | 1.3E+00 | 4.9E-04 | 1.1E+00 | 1.2E-02 | -2.1E-01 | 9.8E-01 | 5 |
| SLC1A4 | 3.9E-01 | 2.2E-01 | 8.2E-01 | 5.2E-03 | 4.2E-01 | 3.0E-01 | 5 |
| C7orf49 | 1.4E+00 | 1.4E-05 | 1.1E+00 | 4.1E-03 | -3.3E-01 | 6.9E-01 | 5 |
| NA | 1.8E+00 | 6.5E-02 | 2.2E+00 | 2.8E-02 | 4.3E-01 | 9.4E-01 | 5 |

| | | | | | | | |
|---|---|---|---|---|---|---|---|
| TSSC1 | 7.9E-01 | 9.3E-04 | 1.4E+00 | 1.1E-08 | 5.9E-01 | 4.2E-02 | 5 |
| STK40 | 1.4E+00 | 7.3E-02 | 2.0E+00 | 8.0E-03 | 6.4E-01 | 6.1E-01 | 5 |
| MED25 | 8.3E-01 | 7.9E-03 | 1.0E+00 | 2.6E-03 | 1.8E-01 | 9.5E-01 | 5 |
| MED25 | 8.3E-01 | 7.9E-03 | 1.0E+00 | 2.6E-03 | 1.8E-01 | 9.5E-01 | 5 |
| PDIA4 | 1.1E+00 | 2.1E-04 | 2.0E+00 | 8.3E-11 | 8.7E-01 | 1.5E-02 | 5 |
| ATP6V1B1 | 9.8E-01 | 1.4E-04 | 1.2E+00 | 6.2E-06 | 2.5E-01 | 7.1E-01 | 5 |
| TBC1D17 | 5.7E-01 | 1.0E-01 | 8.8E-01 | 1.1E-02 | 3.1E-01 | 6.8E-01 | 5 |
| PTPRZ1 | 1.2E+00 | 2.9E-02 | 8.8E-01 | 2.3E-01 | -3.1E-01 | 9.4E-01 | 5 |
| CDK5RAP2 | 9.9E-01 | 2.3E-03 | 9.0E-01 | 1.6E-02 | -8.4E-02 | 9.9E-01 | 5 |
| RPL3 | 5.4E-01 | 2.0E-01 | 1.1E+00 | 5.5E-03 | 5.3E-01 | 3.2E-01 | 5 |
| RPL3 | 5.4E-01 | 2.0E-01 | 1.1E+00 | 5.5E-03 | 5.3E-01 | 3.2E-01 | 5 |
| RPL3 | 5.4E-01 | 2.0E-01 | 1.1E+00 | 5.5E-03 | 5.3E-01 | 3.2E-01 | 5 |
| ZMYM4 | 1.3E+00 | 2.5E-03 | 1.4E+00 | 2.8E-03 | 1.0E-01 | 9.9E-01 | 5 |
| PRMT1 | 9.8E-01 | 1.2E-05 | 6.6E-01 | 1.3E-02 | -3.2E-01 | 4.3E-01 | 5 |
| NA | 5.5E-01 | 1.9E-01 | 9.6E-01 | 1.5E-02 | 4.1E-01 | 5.3E-01 | 5 |
| GSN | 6.3E-01 | 2.2E-02 | 8.7E-01 | 2.4E-03 | 2.5E-01 | 7.4E-01 | 5 |
| STOM | 1.2E+00 | 4.4E-07 | 1.2E+00 | 2.5E-06 | 6.6E-03 | 9.9E-01 | 5 |
| TEAD2 | 4.8E-01 | 9.1E-02 | 6.8E-01 | 2.1E-02 | 2.0E-01 | 8.4E-01 | 5 |
| NA | 8.7E-01 | 9.2E-05 | 1.0E+00 | 1.5E-05 | 1.6E-01 | 9.0E-01 | 5 |
| NA | 8.5E-01 | 1.3E-03 | 1.0E+00 | 3.0E-04 | 1.7E-01 | 9.2E-01 | 5 |
| TRAF3IP2 | 1.1E+00 | 1.4E-04 | 8.8E-01 | 1.3E-02 | -2.6E-01 | 7.9E-01 | 5 |
| C19orf68 | 9.7E-01 | 1.1E-01 | 1.3E+00 | 3.8E-02 | 3.3E-01 | 9.1E-01 | 5 |
| SEPW1 | 1.0E+00 | 9.5E-04 | 1.5E+00 | 3.7E-06 | 4.7E-01 | 3.6E-01 | 5 |
| GLTSCR2 | 4.8E-01 | 5.8E-02 | 9.9E-01 | 5.0E-05 | 5.1E-01 | 9.4E-02 | 5 |
| GLTSCR2 | 4.8E-01 | 5.8E-02 | 9.9E-01 | 5.0E-05 | 5.1E-01 | 9.4E-02 | 5 |
| JRKL | 1.6E+00 | 1.2E-02 | 1.2E+00 | 1.3E-01 | -3.5E-01 | 9.3E-01 | 5 |
| CEP57 | 7.4E-01 | 2.1E-02 | 1.1E+00 | 8.4E-04 | 3.6E-01 | 5.4E-01 | 5 |
| SEC61A1 | 1.1E+00 | 2.4E-05 | 1.0E+00 | 1.4E-04 | -1.3E-02 | 9.9E-01 | 5 |
| FAM76B | 1.8E+00 | 6.3E-03 | 1.4E+00 | 7.0E-02 | -3.3E-01 | 9.5E-01 | 5 |
| RUVBL1 | 9.2E-01 | 1.8E-04 | 8.9E-01 | 1.0E-03 | -2.5E-02 | 9.9E-01 | 5 |
| ACAD9 | 6.8E-01 | 4.5E-03 | 1.2E+00 | 1.0E-06 | 5.1E-01 | 9.8E-02 | 5 |
| EML2 | 1.6E+00 | 2.5E-07 | 1.4E+00 | 2.1E-05 | -1.6E-01 | 9.9E-01 | 5 |
| RTN2 | 8.4E-01 | 1.9E-02 | 6.6E-01 | 1.5E-01 | -1.8E-01 | 9.8E-01 | 5 |
| XPC | 4.0E+00 | 3.4E-02 | 3.5E+00 | 1.1E-01 | -4.8E-01 | 9.9E-01 | 5 |
| CCDC174 | 1.1E+00 | 5.7E-03 | 1.0E+00 | 3.5E-02 | -1.2E-01 | 9.9E-01 | 5 |
| ZNF862 | 6.9E-01 | 1.9E-01 | 1.1E+00 | 3.6E-02 | 3.8E-01 | 7.4E-01 | 5 |
| TSC22D1 | 4.2E-01 | 5.1E-02 | 6.8E-01 | 1.5E-03 | 2.7E-01 | 4.6E-01 | 5 |
| PRCP | 8.1E-01 | 2.6E-02 | 1.5E+00 | 3.2E-05 | 6.8E-01 | 1.3E-01 | 5 |
| BCAM | 4.2E-01 | 8.9E-02 | 6.6E-01 | 7.3E-03 | 2.4E-01 | 6.1E-01 | 5 |
| TMEM214 | 6.3E-01 | 2.5E-02 | 4.7E-01 | 2.1E-01 | -1.6E-01 | 9.6E-01 | 5 |
| FAM107B | 1.5E+00 | 1.1E-04 | 1.1E+00 | 1.5E-02 | -3.8E-01 | 7.0E-01 | 5 |
| CDK5 | 6.6E-01 | 3.1E-02 | 4.9E-01 | 2.3E-01 | -1.6E-01 | 9.7E-01 | 5 |
| PRPF18 | 1.8E+00 | 1.2E-04 | 1.2E+00 | 2.8E-02 | -5.4E-01 | 4.9E-01 | 5 |
| FASTK | 1.8E+00 | 3.4E-09 | 1.2E+00 | 5.5E-04 | -5.8E-01 | 1.8E-01 | 5 |
| PHLDB3 | 1.6E+00 | 5.2E-03 | 1.5E+00 | 2.1E-02 | -8.9E-02 | 9.9E-01 | 5 |
| GAB2 | 9.7E-01 | 6.5E-03 | 1.5E+00 | 2.8E-05 | 5.6E-01 | 3.0E-01 | 5 |
| TMEM53 | 1.7E+00 | 9.4E-02 | 2.2E+00 | 3.9E-02 | 4.8E-01 | 9.4E-01 | 5 |
| DMAP1 | 8.6E-01 | 3.2E-03 | 6.2E-01 | 9.6E-02 | -2.5E-01 | 8.1E-01 | 5 |
| ELF1 | 7.3E-01 | 4.9E-02 | 5.6E-01 | 2.7E-01 | -1.7E-01 | 9.9E-01 | 5 |
| SLC6A9 | 1.4E+00 | 3.6E-04 | 1.6E+00 | 2.1E-04 | 1.8E-01 | 9.9E-01 | 5 |
| B4GALT2 | 1.2E+00 | 2.9E-05 | 1.2E+00 | 1.2E-04 | 4.9E-03 | 9.9E-01 | 5 |
| RPL32 | 8.9E-01 | 2.7E-04 | 1.5E+00 | 4.3E-10 | 6.6E-01 | 2.6E-02 | 5 |
| RSF1 | 6.4E-01 | 3.1E-02 | 5.2E-01 | 1.6E-01 | -1.1E-01 | 9.9E-01 | 5 |
| NA | 1.5E+00 | 9.0E-09 | 1.5E+00 | 5.7E-08 | 1.9E-02 | 9.9E-01 | 5 |
| TGFB1 | 7.8E-01 | 6.7E-03 | 8.2E-01 | 1.1E-02 | 3.8E-02 | 9.9E-01 | 5 |
| HNRNPUL1 | 6.6E-01 | 2.2E-03 | 5.4E-01 | 3.5E-02 | -1.2E-01 | 9.8E-01 | 5 |
| SRP68 | 3.9E-01 | 1.6E-01 | 6.0E-01 | 3.2E-02 | 2.1E-01 | 7.7E-01 | 5 |
| NA | 6.7E-01 | 4.8E-02 | 1.1E+00 | 1.1E-03 | 4.3E-01 | 4.0E-01 | 5 |
| UBXN4 | 6.5E-01 | 3.0E-02 | 8.5E-01 | 7.3E-03 | 2.0E-01 | 8.9E-01 | 5 |
| NUMBL | 1.0E+00 | 1.5E-02 | 1.3E+00 | 4.0E-03 | 2.7E-01 | 9.1E-01 | 5 |
| LTBP4 | 2.4E+00 | 4.6E-02 | 2.3E+00 | 1.1E-01 | -1.6E-01 | 9.9E-01 | 5 |
| BRE | 9.3E-01 | 1.1E-04 | 9.6E-01 | 2.4E-04 | 3.6E-02 | 9.9E-01 | 5 |
| IL17RC | 1.6E+00 | 1.9E-03 | 1.7E+00 | 3.8E-03 | 4.7E-02 | 9.9E-01 | 5 |
| NKTR | 7.6E-01 | 1.2E-02 | 5.4E-01 | 1.8E-01 | -2.2E-01 | 8.7E-01 | 5 |
| PNP | 7.8E-01 | 4.9E-06 | 5.9E-01 | 2.5E-03 | -1.9E-01 | 6.4E-01 | 5 |
| STT3B | 5.3E-01 | 3.4E-02 | 8.2E-01 | 1.4E-03 | 2.9E-01 | 5.4E-01 | 5 |
| NDRG2 | 2.1E+00 | 3.5E-08 | 1.6E+00 | 1.9E-04 | -5.0E-01 | 4.8E-01 | 5 |
| PFKP | 9.0E-01 | 9.4E-05 | 7.9E-01 | 2.3E-03 | -1.1E-01 | 9.9E-01 | 5 |
| ZNF219 | 1.4E+00 | 8.8E-06 | 9.9E-01 | 7.4E-03 | -4.1E-01 | 5.0E-01 | 5 |
| POLB | 5.1E-01 | 8.9E-02 | 8.7E-01 | 3.2E-03 | 3.6E-01 | 4.3E-01 | 5 |
| CBX4 | 1.0E+00 | 4.4E-02 | 1.3E+00 | 1.3E-02 | 3.1E-01 | 9.2E-01 | 5 |
| GAA | 5.3E-01 | 1.0E-01 | 7.7E-01 | 1.8E-02 | 2.5E-01 | 7.9E-01 | 5 |
| EDEM1 | 4.2E-01 | 8.7E-02 | 6.3E-01 | 1.2E-02 | 2.1E-01 | 7.1E-01 | 5 |
| ARHGEF17 | 1.0E+00 | 4.4E-04 | 9.1E-01 | 5.8E-03 | -1.1E-01 | 9.9E-01 | 5 |
| FOLR2 | 6.1E-01 | 3.4E-03 | 7.7E-01 | 5.4E-04 | 1.6E-01 | 8.5E-01 | 5 |
| NA | 4.0E-01 | 5.3E-02 | 8.3E-01 | 2.6E-05 | 4.4E-01 | 6.9E-02 | 5 |
| EIF3K | 5.0E-01 | 5.1E-03 | 1.0E+00 | 5.5E-09 | 5.3E-01 | 7.7E-03 | 5 |
| FEZ2 | 1.5E+00 | 5.1E-04 | 1.2E+00 | 2.5E-02 | -3.5E-01 | 8.0E-01 | 5 |
| GPS1 | 4.8E-01 | 6.5E-02 | 6.1E-01 | 2.9E-02 | 1.3E-01 | 9.8E-01 | 5 |
| CHFR | 1.3E+00 | 1.8E-05 | 1.2E+00 | 2.9E-04 | -8.6E-02 | 9.9E-01 | 5 |
| KITLG | 1.2E+00 | 1.5E-03 | 9.5E-01 | 3.6E-02 | -2.5E-01 | 9.1E-01 | 5 |
| HDAC3 | 6.7E-01 | 1.0E-03 | 9.4E-01 | 1.3E-05 | 2.7E-01 | 5.0E-01 | 5 |
| PPP2CB | 6.1E-01 | 6.7E-03 | 8.6E-01 | 2.4E-04 | 2.5E-01 | 5.8E-01 | 5 |
| PSMD13 | 7.6E-01 | 7.4E-03 | 9.4E-01 | 2.0E-03 | 1.8E-01 | 9.3E-01 | 5 |
| ILK | 6.7E-01 | 9.6E-03 | 6.8E-01 | 2.0E-02 | 1.4E-02 | 9.9E-01 | 5 |
| B4GALNT4 | 1.7E+00 | 1.6E-01 | 2.5E+00 | 3.1E-02 | 8.2E-01 | 6.9E-01 | 5 |
| PTDSS2 | 9.9E-01 | 6.8E-03 | 1.4E+00 | 1.2E-04 | 4.6E-01 | 4.7E-01 | 5 |
| MRPL17 | 6.7E-01 | 4.3E-02 | 7.8E-01 | 3.2E-03 | 1.1E-01 | 9.9E-01 | 5 |
| POLR2I | 6.5E-01 | 1.9E-02 | 8.1E-01 | 6.7E-03 | 1.6E-01 | 9.5E-01 | 5 |
| SUCLG2 | 6.9E-01 | 1.6E-03 | 1.0E+00 | 4.4E-06 | 3.5E-01 | 3.0E-01 | 5 |
| HOXC10 | 1.5E+00 | 4.1E-07 | 1.8E+00 | 5.5E-09 | 3.0E-01 | 6.7E-01 | 5 |
| SPINK5 | 4.2E-01 | 1.1E-01 | 6.1E-01 | 2.3E-02 | 1.9E-01 | 8.2E-01 | 5 |
| NA | 8.1E-01 | 1.0E-02 | 9.7E-01 | 4.5E-03 | 1.6E-01 | 9.9E-01 | 5 |
| CALCOCO1 | 9.9E-01 | 6.3E-03 | 9.1E-01 | 3.3E-02 | -8.6E-02 | 9.9E-01 | 5 |

| | | | | | | | |
|---|---|---|---|---|---|---|---|
| HSD17B12 | 4.3E-01 | 1.2E-01 | 6.9E-01 | 1.2E-02 | 2.6E-01 | 6.5E-01 | 5 |
| PSMF1 | 7.4E-01 | 1.4E-02 | 1.2E+00 | 1.3E-04 | 4.4E-01 | 3.4E-01 | 5 |
| NA | 1.2E+00 | 5.9E-07 | 8.2E-01 | 4.5E-03 | -4.1E-01 | 2.9E-01 | 5 |
| HM13 | 1.3E+00 | 2.8E-05 | 1.3E+00 | 7.3E-05 | 3.5E-02 | 9.9E-01 | 5 |
| USF2 | 9.5E-01 | 4.0E-06 | 8.5E-01 | 1.7E-04 | -9.7E-02 | 9.9E-01 | 5 |
| PPP2R3A | 1.6E+00 | 1.6E-01 | 2.3E+00 | 3.9E-02 | 6.9E-01 | 7.6E-01 | 5 |
| PDE4D | 3.5E+00 | 2.9E-02 | 3.3E+00 | 6.9E-02 | -2.0E-01 | 9.9E-01 | 5 |
| PCM1 | 9.7E-01 | 2.6E-02 | 9.7E-01 | 5.3E-02 | 5.0E-03 | 1.0E+00 | 5 |
| HPN | 8.4E-01 | 1.4E-04 | 1.3E+00 | 7.2E-09 | 4.7E-01 | 1.1E-01 | 5 |
| FBLN7 | 5.6E-01 | 6.1E-02 | 9.6E-01 | 9.6E-04 | 4.1E-01 | 3.4E-01 | 5 |
| GRAMD1A | 1.3E+00 | 2.8E-06 | 1.0E+00 | 8.3E-04 | -2.5E-01 | 7.6E-01 | 5 |
| NA | 7.8E-01 | 4.4E-02 | 1.3E+00 | 5.8E-04 | 5.5E-01 | 3.4E-01 | 5 |
| KIF2A | 1.1E+00 | 6.6E-04 | 8.4E-01 | 2.9E-02 | -2.6E-01 | 8.4E-01 | 5 |
| ASPH | 8.6E-01 | 2.0E-02 | 7.3E-01 | 1.1E-01 | -1.3E-01 | 9.9E-01 | 5 |
| SLC20A1 | 4.6E-01 | 1.2E-01 | 9.1E-01 | 1.4E-03 | 4.5E-01 | 2.6E-01 | 5 |
| CWC27 | 5.9E-01 | 3.7E-02 | 6.1E-01 | 5.7E-02 | 2.6E-02 | 9.9E-01 | 5 |
| 5-Mar | 1.0E+00 | 3.0E-05 | 1.2E+00 | 3.1E-06 | 2.0E-01 | 8.5E-01 | 5 |
| RMND5A | 6.1E-01 | 1.4E-01 | 9.8E-01 | 1.6E-02 | 3.7E-01 | 6.2E-01 | 5 |
| KRCC1 | 1.3E+00 | 4.1E-03 | 1.5E+00 | 2.0E-03 | 2.0E-01 | 9.9E-01 | 5 |
| NA | 1.5E+00 | 3.4E-07 | 1.1E+00 | 1.7E-03 | -4.6E-01 | 3.9E-01 | 5 |
| NA | 1.5E+00 | 3.4E-07 | 1.1E+00 | 1.7E-03 | -4.6E-01 | 3.9E-01 | 5 |
| NA | 1.5E+00 | 3.4E-07 | 1.1E+00 | 1.7E-03 | -4.6E-01 | 3.9E-01 | 5 |
| NA | 1.5E+00 | 3.4E-07 | 1.1E+00 | 1.7E-03 | -4.6E-01 | 3.9E-01 | 5 |
| NA | 1.5E+00 | 3.4E-07 | 1.1E+00 | 1.7E-03 | -4.6E-01 | 3.9E-01 | 5 |
| CEP89 | 1.2E+00 | 3.8E-03 | 9.4E-01 | 6.5E-02 | -2.5E-01 | 9.3E-01 | 5 |
| ARFGEF1 | 6.8E-01 | 1.5E-02 | 8.7E-01 | 3.6E-03 | 1.9E-01 | 8.8E-01 | 5 |
| PPIP5K2 | 5.5E-01 | 5.8E-02 | 7.3E-01 | 1.7E-02 | 1.8E-01 | 9.0E-01 | 5 |
| FLNB | 5.2E-01 | 4.9E-02 | 6.9E-01 | 1.4E-02 | 1.7E-01 | 9.1E-01 | 5 |
| SULF1 | 1.6E+00 | 3.6E-05 | 1.2E+00 | 5.9E-03 | -3.7E-01 | 7.6E-01 | 5 |
| TFDP2 | 1.2E+00 | 1.3E-03 | 1.6E+00 | 4.1E-05 | 4.0E-01 | 5.9E-01 | 5 |
| CAPG | 7.3E-01 | 2.4E-03 | 6.7E-01 | 1.6E-02 | -6.0E-02 | 9.9E-01 | 5 |
| OCLN | 4.6E-01 | 1.3E-01 | 6.7E-01 | 2.9E-02 | 2.1E-01 | 8.3E-01 | 5 |
| UBE2L6 | 8.1E-01 | 8.2E-04 | 1.1E+00 | 9.1E-06 | 3.2E-01 | 4.9E-01 | 5 |
| CLIP1 | 1.3E+00 | 4.8E-04 | 1.2E+00 | 4.3E-03 | -9.3E-02 | 9.9E-01 | 5 |
| NEIL2 | 8.2E-01 | 1.8E-02 | 9.5E-01 | 1.1E-02 | 1.4E-01 | 9.9E-01 | 5 |
| STAU2 | 1.1E+00 | 1.3E-02 | 8.8E-01 | 1.4E-01 | -2.6E-01 | 9.4E-01 | 5 |
| HLTF | 4.2E-01 | 2.2E-01 | 7.4E-01 | 2.2E-02 | 3.3E-01 | 5.7E-01 | 5 |
| POLR3G | 1.0E+00 | 2.4E-03 | 8.4E-01 | 3.9E-02 | -1.9E-01 | 9.6E-01 | 5 |
| SEMA4F | 9.3E-01 | 1.4E-04 | 6.2E-01 | 3.9E-02 | -3.1E-01 | 5.1E-01 | 5 |
| NA | 7.2E-01 | 7.2E-03 | 1.4E+00 | 1.3E-07 | 6.9E-01 | 2.7E-02 | 5 |
| CLU | 8.4E-01 | 1.1E-05 | 6.9E-01 | 1.4E-03 | -1.6E-01 | 8.5E-01 | 5 |
| RASA1 | 8.7E-01 | 2.8E-02 | 6.2E-01 | 2.6E-01 | -2.6E-01 | 9.1E-01 | 5 |
| MADD | 1.6E+00 | 2.5E-07 | 1.3E+00 | 1.0E-04 | -2.7E-01 | 8.1E-01 | 5 |
| CCDC102A | 8.2E-01 | 3.0E-05 | 6.9E-01 | 1.8E-03 | -1.3E-01 | 9.4E-01 | 5 |
| NA | 1.9E+00 | 2.4E-02 | 2.3E+00 | 1.4E-02 | 3.3E-01 | 9.9E-01 | 5 |
| TMA16 | 9.2E-01 | 6.5E-03 | 8.8E-01 | 2.4E-02 | -4.2E-02 | 9.9E-01 | 5 |
| ACP2 | 1.1E+00 | 6.3E-06 | 9.4E-01 | 7.4E-04 | -1.9E-01 | 8.9E-01 | 5 |
| PPP2R2A | 5.9E-01 | 5.2E-02 | 8.9E-01 | 4.8E-03 | 2.9E-01 | 6.8E-01 | 5 |
| TIPARP | 2.2E+00 | 2.3E-02 | 3.5E+00 | 1.7E-04 | 1.3E+00 | 1.5E-01 | 5 |
| NA | 8.7E-01 | 1.6E-01 | 1.5E+00 | 1.3E-02 | 5.9E-01 | 5.2E-01 | 5 |
| RAB11FIP5 | 5.5E-01 | 3.2E-02 | 8.0E-01 | 2.7E-03 | 2.5E-01 | 6.8E-01 | 5 |
| SMIM4 | 6.3E-01 | 6.0E-02 | 1.2E+00 | 2.1E-04 | 5.9E-01 | 1.8E-01 | 5 |
| CDS1 | 3.8E-01 | 2.7E-01 | 7.3E-01 | 2.0E-02 | 3.5E-01 | 4.9E-01 | 5 |
| NR4A2 | 2.6E+00 | 7.3E-10 | 4.0E+00 | 7.2E-21 | 1.4E+00 | 5.0E-03 | 5 |
| CREB3L1 | 1.2E+00 | 2.0E-06 | 8.6E-01 | 5.1E-03 | -3.9E-01 | 3.9E-01 | 5 |
| RIPK1 | 9.1E-01 | 1.1E-02 | 7.5E-01 | 9.5E-02 | -1.6E-01 | 9.9E-01 | 5 |
| SLC22A23 | 8.8E-01 | 5.8E-03 | 6.4E-01 | 1.2E-01 | -2.4E-01 | 8.5E-01 | 5 |
| ZFYVE19 | 1.1E+00 | 4.5E-03 | 8.3E-01 | 9.2E-02 | -2.8E-01 | 8.8E-01 | 5 |
| PRPF4B | 1.3E+00 | 8.2E-04 | 8.9E-01 | 5.8E-02 | -3.8E-01 | 7.0E-01 | 5 |
| CD82 | 1.6E+00 | 3.2E-11 | 1.4E+00 | 6.9E-08 | -1.9E-01 | 8.7E-01 | 5 |
| IRX5 | 5.4E-01 | 1.9E-01 | 1.0E+00 | 7.5E-03 | 4.9E-01 | 3.9E-01 | 5 |
| PTDSS1 | 6.0E-01 | 2.9E-02 | 7.7E-01 | 8.6E-03 | 1.7E-01 | 9.2E-01 | 5 |
| MTDH | 5.9E-01 | 5.1E-03 | 4.6E-01 | 7.0E-02 | -1.2E-01 | 9.6E-01 | 5 |
| NA | 4.0E-01 | 1.2E-01 | 7.1E-01 | 4.3E-03 | 3.1E-01 | 4.3E-01 | 5 |
| NA | 4.0E-01 | 1.2E-01 | 7.1E-01 | 4.3E-03 | 3.1E-01 | 4.3E-01 | 5 |
| TYRO3 | 6.5E-01 | 3.9E-03 | 6.9E-01 | 5.8E-03 | 4.2E-02 | 9.9E-01 | 5 |
| GCNT2 | 1.1E+00 | 4.3E-02 | 8.8E-01 | 2.0E-01 | -1.9E-01 | 9.9E-01 | 5 |
| AP2A2 | 7.0E-01 | 6.0E-03 | 1.1E+00 | 1.5E-05 | 4.3E-01 | 2.5E-01 | 5 |
| AP2A2 | 7.0E-01 | 6.0E-03 | 1.1E+00 | 1.5E-05 | 4.3E-01 | 2.5E-01 | 5 |
| WFDC2 | 5.4E-01 | 1.1E-01 | 1.1E+00 | 9.1E-04 | 5.2E-01 | 2.4E-01 | 5 |
| PRICKLE1 | 1.4E+00 | 1.5E-05 | 9.5E-01 | 1.2E-02 | -4.3E-01 | 4.7E-01 | 5 |
| CTSA | 9.2E-01 | 7.7E-05 | 1.1E+00 | 4.1E-06 | 2.2E-01 | 7.5E-01 | 5 |
| TTF2 | 6.9E-01 | 1.8E-02 | 9.3E-01 | 2.4E-03 | 2.4E-01 | 7.7E-01 | 5 |
| BBX | 7.7E-01 | 1.5E-02 | 8.6E-01 | 1.3E-02 | 9.8E-02 | 9.9E-01 | 5 |
| MAN1A2 | 8.9E-01 | 7.2E-04 | 7.9E-01 | 8.7E-03 | -9.8E-02 | 9.9E-01 | 5 |
| TOLLIP | 5.3E-01 | 1.9E-02 | 8.4E-01 | 3.1E-04 | 3.1E-01 | 4.2E-01 | 5 |
| ETS2 | 1.7E+00 | 2.8E-09 | 2.0E+00 | 3.9E-12 | 3.6E-01 | 5.4E-01 | 5 |
| ST14 | 1.3E+00 | 2.1E-04 | 1.0E+00 | 8.1E-03 | -2.2E-01 | 9.4E-01 | 5 |
| PARP3 | 4.9E-01 | 1.3E-01 | 8.0E-01 | 1.1E-02 | 3.1E-01 | 6.1E-01 | 5 |
| CTSD | 1.7E+00 | 3.7E-05 | 1.2E+00 | 1.1E-02 | -4.7E-01 | 6.3E-01 | 5 |
| CMBL | 4.7E-01 | 1.7E-01 | 8.3E-01 | 1.3E-02 | 3.6E-01 | 5.4E-01 | 5 |
| IGSF5 | 7.7E-01 | 2.0E-03 | 7.2E-01 | 1.3E-02 | -5.8E-02 | 9.9E-01 | 5 |
| IGF2 | 1.6E+00 | 2.2E-04 | 1.8E+00 | 1.7E-04 | 1.7E-01 | 9.9E-01 | 5 |
| RAB35 | 2.8E-01 | 4.2E-01 | 6.0E-01 | 3.7E-02 | 3.2E-01 | 4.8E-01 | 5 |
| GUSB | 5.0E-01 | 4.2E-02 | 7.8E-01 | 1.7E-03 | 2.8E-01 | 5.2E-01 | 5 |
| SFXN2 | 1.2E+00 | 3.3E-05 | 1.0E+00 | 1.1E-03 | -1.3E-01 | 9.9E-01 | 5 |
| SLC35C2 | 8.0E-01 | 1.1E-02 | 8.3E-01 | 2.0E-02 | 2.8E-02 | 9.9E-01 | 5 |
| ADIPOR1 | 6.6E-01 | 1.1E-02 | 5.8E-01 | 6.5E-02 | -8.5E-02 | 9.9E-01 | 5 |
| SLK | 8.9E-01 | 2.5E-02 | 8.5E-01 | 7.1E-02 | -4.4E-02 | 9.9E-01 | 5 |
| VIMP | 1.2E+00 | 6.1E-03 | 1.1E+00 | 4.1E-02 | -1.4E-01 | 9.9E-01 | 5 |
| FGD4 | 7.6E-01 | 3.9E-02 | 7.0E-01 | 1.2E-01 | -6.7E-02 | 9.9E-01 | 5 |
| NPRL2 | 5.1E-01 | 4.3E-02 | 6.2E-01 | 2.2E-02 | 1.1E-01 | 9.9E-01 | 5 |
| GSTO1 | 7.1E-01 | 4.6E-05 | 9.9E-01 | 5.6E-08 | 2.8E-01 | 3.4E-01 | 5 |
| SIK1 | 2.4E+00 | 1.5E-06 | 3.5E+00 | 3.9E-12 | 1.1E+00 | 8.4E-02 | 5 |

| | | | | | | | |
|---|---|---|---|---|---|---|---|
| HYAL1 | 8.4E-01 | 6.9E-03 | 1.4E+00 | 2.2E-05 | 5.1E-01 | 2.7E-01 | 5 |
| ELF3 | 7.5E-01 | 1.0E-02 | 1.1E+00 | 2.4E-04 | 3.6E-01 | 5.0E-01 | 5 |
| COBLL1 | 1.0E+00 | 5.6E-03 | 7.3E-01 | 1.4E-01 | -3.2E-01 | 7.7E-01 | 5 |
| XPNPEP1 | 5.7E-01 | 2.6E-02 | 8.2E-01 | 2.0E-03 | 2.5E-01 | 6.7E-01 | 5 |
| ZNF496 | 6.0E-01 | 5.9E-02 | 7.3E-01 | 3.4E-02 | 1.3E-01 | 9.9E-01 | 5 |
| ADD3 | 9.5E-01 | 4.7E-02 | 1.2E+00 | 1.3E-02 | 3.0E-01 | 8.9E-01 | 5 |
| MXI1 | 4.1E-01 | 1.4E-01 | 7.1E-01 | 1.0E-02 | 2.9E-01 | 5.4E-01 | 5 |
| GNAI2 | 7.1E-01 | 1.5E-03 | 8.7E-01 | 2.7E-04 | 1.6E-01 | 8.8E-01 | 5 |
| PTP4A2 | 5.6E-01 | 2.0E-02 | 7.3E-01 | 4.7E-03 | 1.7E-01 | 8.8E-01 | 5 |
| PFKL | 1.5E+00 | 5.4E-05 | 1.3E+00 | 1.4E-03 | -1.7E-01 | 9.9E-01 | 5 |
| LRP5 | 1.6E+00 | 1.1E-02 | 1.2E+00 | 1.4E-01 | -4.1E-01 | 8.7E-01 | 5 |
| LRP5 | 1.6E+00 | 1.1E-02 | 1.2E+00 | 1.4E-01 | -4.1E-01 | 8.7E-01 | 5 |
| PPP2R2D | 1.0E+00 | 7.9E-04 | 7.8E-01 | 2.9E-02 | -2.2E-01 | 8.6E-01 | 5 |
| SPATS2L | 1.1E+00 | 2.4E-03 | 1.1E+00 | 3.6E-03 | 6.8E-02 | 9.9E-01 | 5 |
| KCTD18 | 9.9E-01 | 3.3E-02 | 1.2E+00 | 1.8E-02 | 2.0E-01 | 9.9E-01 | 5 |
| GSK3B | 5.2E-01 | 9.9E-02 | 1.0E+00 | 5.4E-04 | 5.2E-01 | 1.8E-01 | 5 |
| INPP5A | 7.3E-01 | 2.4E-02 | 6.4E-01 | 1.1E-01 | -9.2E-02 | 9.9E-01 | 5 |
| MSANTD2 | 9.3E-01 | 5.6E-04 | 8.7E-01 | 4.9E-03 | -6.5E-02 | 9.9E-01 | 5 |
| FAM20C | 6.4E-01 | 8.8E-03 | 7.1E-01 | 9.4E-03 | 6.4E-02 | 9.9E-01 | 5 |
| NA | 8.5E-01 | 2.2E-02 | 1.0E+00 | 1.3E-02 | 1.5E-01 | 9.9E-01 | 5 |
| STAU1 | 4.6E-01 | 3.3E-03 | 5.9E-01 | 4.6E-04 | 1.3E-01 | 8.2E-01 | 5 |
| DENND1B | 1.3E+00 | 4.0E-02 | 1.1E+00 | 1.4E-01 | -1.5E-01 | 9.9E-01 | 5 |
| DISP1 | 1.4E+00 | 1.4E-05 | 1.2E+00 | 1.3E-03 | -2.2E-01 | 8.9E-01 | 5 |
| TMEM134 | 4.8E-01 | 8.2E-02 | 7.2E-01 | 1.1E-02 | 2.4E-01 | 7.1E-01 | 5 |
| USP19 | 1.8E+00 | 3.0E-02 | 1.7E+00 | 6.6E-02 | -3.6E-02 | 9.9E-01 | 5 |
| C3orf62 | 1.1E+00 | 3.0E-02 | 1.4E+00 | 1.7E-02 | 2.2E-01 | 9.9E-01 | 5 |
| C3orf62 | 1.1E+00 | 3.0E-02 | 1.4E+00 | 1.7E-02 | 2.2E-01 | 9.9E-01 | 5 |
| DHRS4 | 9.5E-01 | 1.1E-04 | 1.0E+00 | 1.8E-04 | 5.7E-02 | 9.9E-01 | 5 |
| DHRS4 | 9.5E-01 | 1.1E-04 | 1.0E+00 | 1.8E-04 | 5.7E-02 | 9.9E-01 | 5 |
| DHRS4 | 9.5E-01 | 1.1E-04 | 1.0E+00 | 1.8E-04 | 5.7E-02 | 9.9E-01 | 5 |
| ZFP64 | 4.1E-01 | 2.2E-01 | 8.0E-01 | 8.8E-03 | 3.9E-01 | 3.9E-01 | 5 |
| DGKD | 7.3E-01 | 1.5E-02 | 1.3E+00 | 1.3E-05 | 5.8E-01 | 1.2E-01 | 5 |
| PCBP3 | 1.6E+00 | 8.1E-03 | 1.4E+00 | 5.0E-02 | -2.0E-01 | 9.9E-01 | 5 |
| REC8 | 1.2E+00 | 6.5E-04 | 9.1E-01 | 4.0E-02 | -3.2E-01 | 7.4E-01 | 5 |
| ORC2 | 1.2E+00 | 1.9E-03 | 1.1E+00 | 1.0E-02 | -6.4E-02 | 9.9E-01 | 5 |
| PEPCK | 3.6E+00 | 1.3E-02 | 2.9E+00 | 1.1E-01 | -7.5E-01 | 9.4E-01 | 5 |
| RHBDD3 | 9.3E-01 | 3.3E-03 | 8.8E-01 | 1.6E-02 | -5.2E-02 | 9.9E-01 | 5 |
| HYOU1 | 1.2E+00 | 1.3E-02 | 1.6E+00 | 1.9E-03 | 3.9E-01 | 8.2E-01 | 5 |
| GMPR2 | 5.9E-01 | 5.9E-02 | 1.2E+00 | 7.3E-05 | 5.9E-01 | 1.1E-01 | 5 |
| NPEPL1 | 7.4E-01 | 1.4E-02 | 9.0E-01 | 6.3E-03 | 1.6E-01 | 9.8E-01 | 5 |
| NPEPL1 | 7.4E-01 | 1.4E-02 | 9.0E-01 | 6.3E-03 | 1.6E-01 | 9.8E-01 | 5 |
| MAP3K6 | 1.1E+00 | 8.3E-03 | 1.0E+00 | 2.9E-02 | -5.1E-02 | 9.9E-01 | 5 |
| ARSJ | 1.8E+00 | 1.0E-09 | 1.5E+00 | 3.3E-06 | -3.0E-01 | 6.8E-01 | 5 |
| RABGGTA | 8.2E-01 | 1.4E-02 | 7.9E-01 | 4.0E-02 | -2.9E-02 | 9.9E-01 | 5 |
| WDTC1 | 1.7E+00 | 5.3E-08 | 1.1E+00 | 2.0E-03 | -5.6E-01 | 2.1E-01 | 5 |
| GAS2L1 | 1.2E+00 | 1.1E-05 | 8.8E-01 | 4.1E-03 | -2.9E-01 | 6.6E-01 | 5 |
| NA | 8.3E-01 | 1.2E-02 | 1.2E+00 | 9.1E-04 | 3.2E-01 | 6.8E-01 | 5 |
| C2CD2L | 9.3E-01 | 2.8E-03 | 9.1E-01 | 1.0E-02 | -2.3E-02 | 9.9E-01 | 5 |
| TMEM206 | 5.6E-01 | 1.4E-01 | 1.1E+00 | 3.3E-03 | 5.0E-01 | 3.2E-01 | 5 |
| ILKAP | 1.0E+00 | 5.3E-05 | 1.7E+00 | 1.0E-11 | 6.9E-01 | 1.3E-02 | 5 |
| CCS | 7.0E-01 | 2.6E-02 | 5.3E-01 | 1.9E-01 | -1.6E-01 | 9.8E-01 | 5 |
| TRAF3IP1 | 1.4E+00 | 9.5E-04 | 1.2E+00 | 2.0E-02 | -2.4E-01 | 9.5E-01 | 5 |
| SMYD2 | 6.6E-01 | 5.5E-02 | 9.2E-01 | 1.1E-02 | 2.6E-01 | 8.1E-01 | 5 |
| ITGB5 | 7.1E-01 | 1.1E-03 | 8.7E-01 | 2.2E-04 | 1.6E-01 | 9.0E-01 | 5 |
| SCNM1 | 7.6E-01 | 1.4E-01 | 1.3E+00 | 5.2E-03 | 5.9E-01 | 3.9E-01 | 5 |
| SCNM1 | 7.6E-01 | 1.4E-01 | 1.3E+00 | 5.2E-03 | 5.9E-01 | 3.9E-01 | 5 |
| SF3A1 | 8.3E-01 | 6.2E-03 | 1.1E+00 | 5.5E-04 | 2.8E-01 | 7.4E-01 | 5 |
| PSMD4 | 1.7E+00 | 2.3E-10 | 1.2E+00 | 4.9E-05 | -4.9E-01 | 2.5E-01 | 5 |
| SYAP1 | 1.4E+00 | 1.9E-03 | 9.9E-01 | 7.2E-02 | -3.8E-01 | 7.9E-01 | 5 |
| IFT46 | 4.7E-01 | 1.1E-01 | 7.2E-01 | 1.6E-02 | 2.5E-01 | 7.3E-01 | 5 |
| NBEAL1 | 6.1E-01 | 6.1E-02 | 1.0E+00 | 2.0E-03 | 4.0E-01 | 4.6E-01 | 5 |
| SELENBP1 | 6.0E-01 | 2.1E-01 | 1.0E+00 | 2.0E-02 | 4.4E-01 | 5.5E-01 | 5 |
| RPS21 | 5.7E-01 | 1.4E-02 | 1.1E+00 | 6.4E-06 | 4.8E-01 | 1.1E-01 | 5 |
| CEP350 | 6.0E-01 | 4.0E-02 | 5.5E-01 | 1.2E-01 | -4.8E-02 | 9.9E-01 | 5 |
| BUB3 | 4.8E-01 | 3.8E-02 | 9.4E-01 | 4.2E-05 | 4.5E-01 | 1.2E-01 | 5 |
| MRPL11 | 8.3E-01 | 2.2E-04 | 1.2E+00 | 9.3E-07 | 3.3E-01 | 4.0E-01 | 5 |
| UFC1 | 1.8E+00 | 2.0E-12 | 1.5E+00 | 5.3E-07 | -3.9E-01 | 4.2E-01 | 5 |
| TMEM57 | 1.3E+00 | 2.9E-05 | 9.4E-01 | 1.0E-02 | -3.6E-01 | 5.9E-01 | 5 |
| MUC20 | 7.8E-01 | 1.6E-02 | 5.8E-01 | 1.8E-01 | -2.0E-01 | 9.2E-01 | 5 |
| NDUFS2 | 1.7E+00 | 1.6E-06 | 1.5E+00 | 1.4E-04 | -2.2E-01 | 9.6E-01 | 5 |
| TOMM40L | 7.4E-01 | 3.9E-02 | 6.1E-01 | 1.9E-01 | -1.3E-01 | 9.9E-01 | 5 |
| HELZ2 | 6.8E-01 | 1.3E-01 | 9.7E-01 | 3.6E-02 | 2.9E-01 | 8.7E-01 | 5 |
| SDHC | 1.2E+00 | 5.2E-05 | 9.2E-01 | 8.4E-03 | -2.9E-01 | 7.3E-01 | 5 |
| GMEB2 | 8.9E-01 | 5.9E-03 | 1.0E+00 | 4.2E-03 | 1.2E-01 | 9.9E-01 | 5 |
| RNF168 | 3.8E-01 | 3.3E-01 | 7.3E-01 | 3.7E-02 | 3.5E-01 | 5.7E-01 | 5 |
| CYB561 | 7.7E-01 | 1.9E-03 | 9.1E-01 | 7.2E-04 | 1.4E-01 | 9.8E-01 | 5 |
| DHX30 | 1.3E+00 | 3.2E-04 | 1.7E+00 | 1.9E-05 | 3.5E-01 | 7.4E-01 | 5 |
| RNF214 | 7.3E-01 | 1.1E-01 | 1.1E+00 | 1.6E-02 | 3.9E-01 | 7.1E-01 | 5 |
| PRPF6 | 1.2E+00 | 7.1E-07 | 1.0E+00 | 2.4E-04 | -2.2E-01 | 8.2E-01 | 5 |
| VPS8 | 1.1E+00 | 1.5E-04 | 8.7E-01 | 7.0E-03 | -1.8E-01 | 9.2E-01 | 5 |
| NA | 5.8E-01 | 4.3E-02 | 1.0E+00 | 2.0E-04 | 4.7E-01 | 2.1E-01 | 5 |
| SIDT2 | 7.7E-01 | 1.2E-03 | 8.6E-01 | 8.8E-04 | 9.1E-02 | 9.9E-01 | 5 |
| NA | 9.7E-01 | 1.3E-03 | 8.1E-01 | 2.5E-02 | -1.7E-01 | 9.7E-01 | 5 |
| HEBP1 | 6.6E-01 | 1.9E-03 | 6.4E-01 | 8.1E-03 | -2.2E-02 | 9.9E-01 | 5 |
| SMS | 1.1E+00 | 1.9E-02 | 8.0E-01 | 2.2E-01 | -3.2E-01 | 8.7E-01 | 5 |
| KAT5 | 1.5E+00 | 3.0E-05 | 1.3E+00 | 1.1E-03 | -1.8E-01 | 9.9E-01 | 5 |
| CSDA | 1.4E+00 | 6.7E-04 | 1.1E+00 | 3.0E-02 | -3.3E-01 | 8.2E-01 | 5 |
| PDK3 | 4.5E-01 | 3.6E-02 | 6.2E-01 | 5.6E-03 | 1.7E-01 | 8.0E-01 | 5 |
| STYXL1 | 1.6E+00 | 4.0E-09 | 1.3E+00 | 3.8E-05 | -3.6E-01 | 4.8E-01 | 5 |
| LTBP3 | 4.9E-01 | 1.1E-01 | 9.3E-01 | 1.7E-03 | 4.4E-01 | 3.0E-01 | 5 |
| HSPB1 | 8.3E-01 | 8.3E-03 | 9.5E-01 | 5.7E-03 | 1.2E-01 | 9.9E-01 | 5 |
| EIF3J | 1.3E+00 | 5.3E-07 | 1.0E+00 | 3.4E-04 | -2.6E-01 | 7.1E-01 | 5 |
| ARHGEF10 | 9.1E-01 | 2.0E-01 | 1.4E+00 | 3.9E-02 | 4.9E-01 | 7.3E-01 | 5 |
| NA | 2.3E+00 | 1.4E-03 | 2.0E+00 | 1.3E-02 | -2.5E-01 | 9.9E-01 | 5 |

| | | | | | | | |
|---|---|---|---|---|---|---|---|
| MAP2 | 4.5E-01 | 3.3E-02 | 7.2E-01 | 6.8E-04 | 2.8E-01 | 4.3E-01 | 5 |
| BCR | 1.8E+00 | 3.4E-03 | 1.7E+00 | 1.6E-02 | -1.1E-01 | 9.9E-01 | 5 |
| ABL2 | 1.0E+00 | 1.7E-01 | 1.7E+00 | 1.4E-02 | 7.0E-01 | 5.5E-01 | 5 |
| NA | 3.8E-01 | 1.1E-01 | 7.9E-01 | 3.3E-04 | 4.1E-01 | 1.4E-01 | 5 |
| TYSND1 | 7.1E-01 | 4.4E-02 | 7.5E-01 | 6.2E-02 | 4.1E-02 | 9.9E-01 | 5 |
| OSBPL1A | 6.3E-01 | 4.9E-02 | 7.7E-01 | 2.5E-02 | 1.5E-01 | 9.9E-01 | 5 |
| TANGO2 | 5.3E-01 | 1.9E-01 | 1.0E+00 | 9.7E-03 | 4.6E-01 | 4.5E-01 | 5 |
| SSFA2 | 1.3E+00 | 3.2E-02 | 1.4E+00 | 5.8E-02 | 1.7E-02 | 9.9E-01 | 5 |
| DNAJB12 | 1.4E+00 | 2.3E-05 | 1.0E+00 | 6.3E-03 | -3.4E-01 | 6.6E-01 | 5 |
| PPP1R35 | 8.4E-01 | 2.1E-03 | 1.2E+00 | 1.6E-05 | 3.8E-01 | 4.1E-01 | 5 |
| HOMER3 | 9.9E-01 | 1.3E-02 | 9.5E-01 | 3.9E-02 | -3.7E-02 | 9.9E-01 | 5 |
| ZNF445 | 7.8E-01 | 2.3E-02 | 7.2E-01 | 7.9E-02 | -6.6E-02 | 9.9E-01 | 5 |
| LRPAP1 | 1.1E+00 | 7.7E-06 | 1.4E+00 | 3.4E-07 | 2.4E-01 | 7.8E-01 | 5 |
| CTDSP1 | 1.3E+00 | 3.7E-08 | 8.9E-01 | 1.6E-03 | -4.5E-01 | 2.1E-01 | 5 |
| TMEM59L | 8.9E-01 | 1.6E-04 | 1.0E+00 | 3.4E-05 | 1.6E-01 | 9.2E-01 | 5 |
| UBA52 | 1.2E+00 | 2.9E-07 | 9.6E-01 | 3.9E-04 | -2.8E-01 | 6.2E-01 | 5 |
| NOP14 | 8.5E-01 | 5.7E-03 | 6.5E-01 | 9.6E-02 | -2.1E-01 | 9.0E-01 | 5 |
| RBM10 | 7.0E-01 | 6.2E-03 | 7.7E-01 | 6.9E-03 | 6.7E-02 | 9.9E-01 | 5 |
| TUBG1 | 1.2E+00 | 1.8E-05 | 8.5E-01 | 1.0E-02 | -3.5E-01 | 5.3E-01 | 5 |
| ECE1 | 1.7E+00 | 2.6E-05 | 1.8E+00 | 1.9E-05 | 1.6E-01 | 9.9E-01 | 5 |
| LETM1 | 8.6E-01 | 1.2E-02 | 9.2E-01 | 1.7E-02 | 6.1E-02 | 9.9E-01 | 5 |
| USP11 | 1.5E+00 | 1.1E-12 | 1.0E+00 | 2.5E-05 | -5.0E-01 | 8.4E-02 | 5 |
| DDOST | 1.5E+00 | 1.7E-10 | 1.3E+00 | 2.6E-07 | -1.9E-01 | 8.6E-01 | 5 |
| MUL1 | 9.6E-01 | 1.2E-02 | 1.1E+00 | 1.0E-02 | 1.3E-01 | 9.9E-01 | 5 |
| NA | 1.4E+00 | 1.6E-07 | 1.4E+00 | 1.6E-07 | 8.7E-02 | 9.9E-01 | 5 |
| ANKZF1 | 6.8E-01 | 1.3E-02 | 6.7E-01 | 3.5E-02 | -1.4E-02 | 9.9E-01 | 5 |
| SELL | 1.4E+00 | 9.8E-05 | 1.2E+00 | 5.9E-03 | -2.6E-01 | 8.7E-01 | 5 |
| SNAP29 | 8.0E-01 | 2.8E-03 | 1.0E+00 | 3.4E-04 | 2.2E-01 | 8.2E-01 | 5 |
| TUBA4A | 1.5E+00 | 1.3E-06 | 1.2E+00 | 6.5E-04 | -3.2E-01 | 7.0E-01 | 5 |
| MYBBP1A | 7.9E-01 | 1.2E-02 | 9.3E-01 | 6.3E-03 | 1.5E-01 | 9.9E-01 | 5 |
| SDF2L1 | 1.3E+00 | 1.9E-02 | 1.9E+00 | 1.1E-03 | 5.8E-01 | 6.3E-01 | 5 |
| UBR4 | 7.3E-01 | 2.1E-02 | 4.9E-01 | 2.7E-01 | -2.4E-01 | 8.4E-01 | 5 |
| MVB12A | 7.6E-01 | 3.8E-03 | 5.7E-01 | 8.1E-02 | -1.9E-01 | 8.8E-01 | 5 |
| TAS1R2 | 3.8E-01 | 1.6E-01 | 6.3E-01 | 1.8E-02 | 2.5E-01 | 6.4E-01 | 5 |
| TOP3B | 9.6E-01 | 3.9E-04 | 8.0E-01 | 1.1E-02 | -1.6E-01 | 9.7E-01 | 5 |
| WDR1 | 6.9E-01 | 8.4E-04 | 1.0E+00 | 1.5E-06 | 3.5E-01 | 2.8E-01 | 5 |
| DHX33 | 6.9E-01 | 2.2E-02 | 6.4E-01 | 7.1E-02 | -4.5E-02 | 9.9E-01 | 5 |
| RABEP1 | 2.4E+00 | 2.6E-03 | 1.8E+00 | 6.5E-02 | -5.8E-01 | 8.1E-01 | 5 |
| NA | 1.2E+00 | 6.7E-04 | 1.7E+00 | 5.2E-06 | 4.8E-01 | 4.5E-01 | 5 |
| RAB5C | 1.5E+00 | 1.8E-06 | 1.6E+00 | 2.4E-06 | 8.5E-02 | 9.9E-01 | 5 |
| ACTC | 1.3E+00 | 1.1E-01 | 2.2E+00 | 4.8E-03 | 9.4E-01 | 4.7E-01 | 5 |
| ARHGEF10L | 7.4E-01 | 7.3E-03 | 7.5E-01 | 1.7E-02 | 8.1E-03 | 9.9E-01 | 5 |
| EVC2 | 7.9E-01 | 5.6E-04 | 6.1E-01 | 2.6E-02 | -1.8E-01 | 8.5E-01 | 5 |
| TFE3 | 8.2E-01 | 1.3E-02 | 8.0E-01 | 3.6E-02 | -2.0E-02 | 9.9E-01 | 5 |
| WNK1 | 6.4E-01 | 4.1E-02 | 4.7E-01 | 2.7E-01 | -1.7E-01 | 9.7E-01 | 5 |
| SYNE2 | 8.3E-01 | 1.2E-02 | 6.9E-01 | 9.2E-02 | -1.4E-01 | 9.9E-01 | 5 |
| DYDC1 | 5.4E-01 | 2.9E-01 | 9.8E-01 | 3.6E-02 | 4.4E-01 | 6.2E-01 | 5 |
| ERC1 | 1.3E+00 | 9.7E-05 | 9.0E-01 | 2.7E-02 | -4.1E-01 | 5.5E-01 | 5 |
| FKBP10 | 1.3E+00 | 1.6E-06 | 1.5E+00 | 1.1E-07 | 2.1E-01 | 8.6E-01 | 5 |
| DVL2 | 1.4E+00 | 2.7E-05 | 1.4E+00 | 1.3E-04 | -4.0E-03 | 1.0E+00 | 5 |
| RFC1 | 7.6E-01 | 3.6E-02 | 5.4E-01 | 2.8E-01 | -2.2E-01 | 9.3E-01 | 5 |
| SNCG | 1.1E+00 | 8.2E-03 | 1.1E+00 | 1.3E-02 | 5.9E-02 | 9.9E-01 | 5 |
| AP5Z1 | 1.3E+00 | 3.9E-07 | 1.0E+00 | 3.4E-04 | -2.8E-01 | 6.8E-01 | 5 |
| AP5Z1 | 1.3E+00 | 3.9E-07 | 1.0E+00 | 3.4E-04 | -2.8E-01 | 6.8E-01 | 5 |
| SPEN | 1.5E+00 | 8.4E-06 | 1.1E+00 | 7.8E-03 | -4.6E-01 | 4.9E-01 | 5 |
| GPS2 | 6.6E-01 | 4.0E-02 | 4.9E-01 | 2.6E-01 | -1.7E-01 | 9.7E-01 | 5 |
| NA | 1.4E+00 | 3.4E-03 | 1.3E+00 | 2.8E-02 | -1.8E-01 | 9.9E-01 | 5 |
| ENAH | 1.5E+00 | 1.1E-07 | 1.2E+00 | 1.5E-04 | -3.2E-01 | 6.6E-01 | 5 |
| TBC1D1 | 6.7E-01 | 5.4E-03 | 6.0E-01 | 3.2E-02 | -6.6E-02 | 9.9E-01 | 5 |
| ACAP1 | 8.5E-01 | 2.6E-02 | 6.0E-01 | 2.6E-01 | -2.5E-01 | 8.9E-01 | 5 |
| CGNL1 | 1.3E+00 | 9.3E-04 | 1.3E+00 | 1.6E-03 | 6.0E-02 | 9.9E-01 | 5 |
| CGNL1 | 1.3E+00 | 9.3E-04 | 1.3E+00 | 1.6E-03 | 6.0E-02 | 9.9E-01 | 5 |
| CGNL1 | 1.3E+00 | 9.3E-04 | 1.3E+00 | 1.6E-03 | 6.0E-02 | 9.9E-01 | 5 |
| MYZAP | 1.7E+00 | 3.6E-03 | 2.6E+00 | 5.0E-06 | 9.4E-01 | 1.1E-01 | 5 |
| GIPC1 | 1.4E+00 | 3.2E-06 | 1.1E+00 | 1.3E-03 | -3.1E-01 | 7.0E-01 | 5 |
| ZBTB4 | 4.1E-01 | 9.8E-02 | 7.2E-01 | 3.2E-03 | 3.1E-01 | 4.3E-01 | 5 |
| BRAT1 | 6.0E-01 | 3.1E-02 | 6.0E-01 | 6.5E-02 | -2.0E-03 | 1.0E+00 | 5 |
| TCAP | 1.9E+00 | 1.9E-03 | 1.5E+00 | 4.4E-02 | -4.1E-01 | 8.4E-01 | 5 |
| STARD3 | 8.1E-01 | 1.5E-02 | 7.0E-01 | 8.0E-02 | -1.1E-01 | 9.9E-01 | 5 |
| SMIM20 | 5.8E-01 | 1.8E-02 | 1.1E+00 | 2.8E-06 | 5.5E-01 | 5.5E-02 | 5 |
| NSD1 | 1.7E+00 | 1.3E-05 | 1.1E+00 | 1.1E-02 | -5.1E-01 | 4.3E-01 | 5 |
| PRKACA | 1.1E+00 | 8.8E-06 | 1.0E+00 | 2.4E-04 | -1.0E-01 | 9.9E-01 | 5 |
| CCDC149 | 1.2E+00 | 6.3E-03 | 1.6E+00 | 6.5E-04 | 3.7E-01 | 7.5E-01 | 5 |
| ACTN1 | 7.8E-01 | 5.8E-02 | 1.3E+00 | 1.7E-03 | 5.1E-01 | 4.6E-01 | 5 |
| PFDN4 | 1.4E+00 | 1.6E-03 | 1.1E+00 | 2.4E-02 | -2.1E-01 | 9.8E-01 | 5 |
| MTOR | 4.4E-01 | 2.0E-02 | 8.0E-01 | 2.2E-05 | 3.6E-01 | 1.3E-01 | 5 |
| FGFRL1 | 1.6E+00 | 3.0E-03 | 1.9E+00 | 8.2E-04 | 3.1E-01 | 9.3E-01 | 5 |
| KIAA1191 | 5.1E-01 | 4.0E-02 | 9.6E-01 | 8.4E-05 | 4.5E-01 | 1.6E-01 | 5 |
| P53 | 8.3E-01 | 2.6E-02 | 1.2E+00 | 2.7E-03 | 3.3E-01 | 6.9E-01 | 5 |
| WRAP53 | 5.7E-01 | 1.4E-02 | 6.3E-01 | 1.6E-02 | 5.6E-02 | 9.9E-01 | 5 |
| HDGF | 8.6E-01 | 3.4E-03 | 7.3E-01 | 3.3E-02 | -1.2E-01 | 9.9E-01 | 5 |
| STX4 | 6.2E-01 | 3.8E-02 | 6.9E-01 | 3.7E-02 | 7.3E-02 | 9.9E-01 | 5 |
| APOA1BP | 6.2E-01 | 1.0E-02 | 6.4E-01 | 2.2E-02 | 1.2E-02 | 9.9E-01 | 5 |
| PGD | 1.1E+00 | 2.0E-06 | 1.2E+00 | 4.5E-06 | 4.3E-02 | 9.9E-01 | 5 |
| CCT3 | 8.7E-01 | 3.2E-03 | 1.1E+00 | 5.9E-04 | 2.2E-01 | 8.8E-01 | 5 |
| CCT3 | 8.7E-01 | 3.2E-03 | 1.1E+00 | 5.9E-04 | 2.2E-01 | 8.8E-01 | 5 |
| NA | 9.3E-01 | 3.7E-02 | 1.2E+00 | 1.1E-02 | 2.7E-01 | 9.1E-01 | 5 |
| FOXO4 | 1.5E+00 | 1.7E-03 | 1.3E+00 | 2.8E-02 | -2.6E-01 | 9.7E-01 | 5 |
| MSTO1 | 8.0E-01 | 4.7E-02 | 7.7E-01 | 1.1E-01 | -3.5E-02 | 9.9E-01 | 5 |
| LRP | 6.4E-01 | 6.9E-03 | 4.6E-01 | 1.3E-01 | -1.8E-01 | 8.5E-01 | 5 |
| GBA | 7.9E-01 | 5.7E-03 | 6.1E-01 | 8.7E-02 | -1.8E-01 | 9.3E-01 | 5 |
| COL1A1 | 9.9E-01 | 2.6E-02 | 8.8E-01 | 1.0E-01 | -1.1E-01 | 9.9E-01 | 5 |
| NA | 1.1E+00 | 9.4E-03 | 1.0E+00 | 3.9E-02 | -8.1E-02 | 9.9E-01 | 5 |
| RAD23A | 7.0E-01 | 1.2E-02 | 8.8E-01 | 2.9E-03 | 1.8E-01 | 9.1E-01 | 5 |

| | | | | | | | |
|---|---|---|---|---|---|---|---|
| CALR | 1.4E+00 | 1.2E-03 | 1.0E+00 | 5.0E-02 | -3.6E-01 | 8.0E-01 | 5 |
| CLN3 | 5.9E-01 | 2.2E-01 | 9.5E-01 | 4.2E-02 | 3.6E-01 | 7.5E-01 | 5 |
| PBXIP1 | 7.0E-01 | 5.4E-04 | 7.9E-01 | 2.8E-04 | 9.7E-02 | 9.9E-01 | 5 |
| POMT2 | 1.3E+00 | 1.9E-06 | 1.1E+00 | 2.1E-04 | -1.8E-01 | 9.3E-01 | 5 |
| ATP8B2 | 4.6E-01 | 1.9E-01 | 7.9E-01 | 1.6E-02 | 3.4E-01 | 5.6E-01 | 5 |
| NA | 1.2E+00 | 2.3E-02 | 9.2E-01 | 2.1E-01 | -3.2E-01 | 9.1E-01 | 5 |
| NA | 5.9E-01 | 9.3E-02 | 8.9E-01 | 1.2E-02 | 3.0E-01 | 6.9E-01 | 5 |
| PWWP2A | 1.3E+00 | 1.7E-05 | 1.2E+00 | 4.5E-04 | -1.2E-01 | 9.9E-01 | 5 |
| CD19 | 8.3E-01 | 9.0E-02 | 1.5E+00 | 1.3E-03 | 6.8E-01 | 2.9E-01 | 5 |
| SLC39A1 | 1.7E+00 | 9.6E-10 | 1.4E+00 | 5.8E-06 | -3.1E-01 | 6.5E-01 | 5 |
| CUEDC1 | 1.6E+00 | 6.3E-08 | 1.2E+00 | 2.0E-04 | -3.6E-01 | 5.5E-01 | 5 |
| MAP2K5 | 1.4E+00 | 3.1E-05 | 1.2E+00 | 3.1E-03 | -2.7E-01 | 8.5E-01 | 5 |
| NA | 5.4E-01 | 8.3E-02 | 9.4E-01 | 1.8E-03 | 4.0E-01 | 3.5E-01 | 5 |
| CLN6 | 9.6E-01 | 7.1E-03 | 1.1E+00 | 6.8E-03 | 1.1E-01 | 9.9E-01 | 5 |
| LCMT1 | 3.0E-01 | 2.5E-01 | 6.0E-01 | 1.2E-02 | 3.0E-01 | 4.0E-01 | 5 |
| S100A4 | 6.5E-01 | 1.1E-02 | 4.7E-01 | 1.5E-01 | -1.8E-01 | 9.0E-01 | 5 |
| NA | 3.7E-01 | 1.2E-01 | 6.8E-01 | 3.4E-03 | 3.1E-01 | 3.7E-01 | 5 |
| FHOD3 | 1.5E+00 | 2.7E-05 | 1.1E+00 | 8.5E-03 | -3.9E-01 | 6.3E-01 | 5 |
| NPTN | 7.4E-01 | 1.4E-03 | 1.2E+00 | 1.2E-06 | 4.3E-01 | 2.1E-01 | 5 |
| SETD3 | 1.2E+00 | 1.7E-03 | 8.2E-01 | 8.2E-02 | -3.5E-01 | 7.2E-01 | 5 |
| ZNF18 | 1.3E+00 | 3.3E-06 | 1.3E+00 | 1.5E-05 | 1.2E-02 | 9.9E-01 | 5 |
| AATF | 8.5E-01 | 2.1E-03 | 8.3E-01 | 7.9E-03 | -2.0E-02 | 9.9E-01 | 5 |
| EVL | 1.5E+00 | 2.4E-03 | 1.2E+00 | 3.6E-02 | -2.6E-01 | 9.8E-01 | 5 |
| YY1 | 4.4E-01 | 3.5E-03 | 7.8E-01 | 4.8E-07 | 3.3E-01 | 7.9E-02 | 5 |
| WARS | 1.1E+00 | 5.4E-05 | 7.7E-01 | 2.6E-02 | -3.7E-01 | 4.7E-01 | 5 |
| C19orf66 | 1.6E+00 | 1.5E-06 | 1.1E+00 | 4.1E-03 | -5.0E-01 | 3.5E-01 | 5 |
| SNUPN | 1.2E+00 | 7.6E-04 | 1.3E+00 | 1.2E-03 | 6.9E-02 | 9.9E-01 | 5 |
| SNX33 | 1.1E+00 | 1.6E-04 | 1.1E+00 | 2.1E-04 | 7.5E-02 | 9.9E-01 | 5 |
| DCTN4 | 4.3E-01 | 1.8E-01 | 6.6E-01 | 4.0E-02 | 2.3E-01 | 8.1E-01 | 5 |
| RCN2 | 9.3E-01 | 2.7E-04 | 9.6E-01 | 6.2E-04 | 3.1E-02 | 9.9E-01 | 5 |
| RPS14 | 8.3E-01 | 7.6E-04 | 1.3E+00 | 4.4E-07 | 4.6E-01 | 2.0E-01 | 5 |
| CDC42BPB | 7.4E-01 | 1.8E-03 | 6.0E-01 | 3.3E-02 | -1.4E-01 | 9.6E-01 | 5 |
| ZNF358 | 1.5E+00 | 5.3E-09 | 1.2E+00 | 2.8E-05 | -2.9E-01 | 6.0E-01 | 5 |
| SNRPD1 | 1.4E+00 | 8.0E-05 | 1.1E+00 | 7.3E-03 | -2.9E-01 | 8.3E-01 | 5 |
| AKT1 | 6.1E-01 | 4.4E-03 | 6.4E-01 | 7.4E-03 | 3.1E-02 | 9.9E-01 | 5 |
| DRG2 | 7.9E-01 | 1.8E-03 | 6.6E-01 | 2.7E-02 | -1.3E-01 | 9.9E-01 | 5 |
| TMEM11 | 1.5E+00 | 1.8E-06 | 1.2E+00 | 1.4E-03 | -3.7E-01 | 6.0E-01 | 5 |
| PDE4B | 1.0E+00 | 6.4E-07 | 8.8E-01 | 1.2E-04 | -1.5E-01 | 9.0E-01 | 5 |
| PTGER4 | 6.5E-01 | 3.5E-02 | 1.1E+00 | 2.3E-04 | 4.9E-01 | 2.7E-01 | 5 |
| KHSRP | 1.0E+00 | 8.1E-05 | 9.2E-01 | 1.9E-03 | -1.2E-01 | 9.9E-01 | 5 |
| CAPS | 1.1E+00 | 2.2E-04 | 1.1E+00 | 1.0E-03 | -1.6E-02 | 9.9E-01 | 5 |
| NA | 9.2E-01 | 1.8E-04 | 7.9E-01 | 5.4E-03 | -1.4E-01 | 9.8E-01 | 5 |
| LONP1 | 1.9E+00 | 1.9E-06 | 1.6E+00 | 2.2E-04 | -2.7E-01 | 9.0E-01 | 5 |
| MAP7D3 | 8.6E-01 | 1.1E-03 | 7.9E-01 | 9.4E-03 | -7.3E-02 | 9.9E-01 | 5 |
| CORO6 | 1.9E+00 | 7.4E-03 | 1.8E+00 | 3.6E-02 | -1.8E-01 | 9.9E-01 | 5 |
| BLMH | 8.1E-01 | 4.2E-03 | 7.1E-01 | 3.3E-02 | -1.0E-01 | 9.9E-01 | 5 |
| PMM2 | 7.8E-01 | 2.7E-02 | 7.6E-01 | 6.4E-02 | -1.7E-02 | 9.9E-01 | 5 |
| SSBP3 | 6.0E-01 | 2.3E-02 | 8.3E-01 | 3.1E-03 | 2.2E-01 | 7.8E-01 | 5 |
| NXN | 1.1E+00 | 4.4E-02 | 9.4E-01 | 1.6E-01 | -1.3E-01 | 9.9E-01 | 5 |
| NA | 1.6E+00 | 1.5E-05 | 1.4E+00 | 1.1E-03 | -2.6E-01 | 9.1E-01 | 5 |
| NCLN | 1.4E+00 | 5.3E-06 | 9.1E-01 | 9.1E-03 | -4.4E-01 | 4.0E-01 | 5 |
| DVL1 | 8.1E-01 | 3.3E-04 | 9.7E-01 | 7.2E-05 | 1.5E-01 | 9.3E-01 | 5 |
| CPSF3L | 7.9E-01 | 1.4E-03 | 8.8E-01 | 1.2E-03 | 8.5E-02 | 9.9E-01 | 5 |
| CPSF3L | 7.9E-01 | 1.4E-03 | 8.8E-01 | 1.2E-03 | 8.5E-02 | 9.9E-01 | 5 |
| SRR | 1.0E+00 | 9.6E-05 | 1.1E+00 | 8.0E-05 | 9.6E-02 | 9.9E-01 | 5 |
| CLUH | 4.0E-01 | 1.9E-01 | 6.3E-01 | 3.7E-02 | 2.3E-01 | 7.6E-01 | 5 |
| PLEKHN1 | 4.9E-01 | 3.5E-01 | 9.5E-01 | 3.6E-02 | 4.6E-01 | 5.1E-01 | 5 |
| L1CAM | 5.0E-01 | 8.7E-02 | 7.7E-01 | 8.5E-03 | 2.7E-01 | 6.6E-01 | 5 |
| SF3A2 | 1.1E+00 | 1.2E-04 | 7.6E-01 | 3.8E-02 | -3.9E-01 | 5.0E-01 | 5 |
| PANK4 | 1.1E+00 | 5.4E-03 | 1.0E+00 | 2.0E-02 | -4.2E-02 | 9.9E-01 | 5 |
| NOXA1 | 5.7E-01 | 4.3E-02 | 7.6E-01 | 1.1E-02 | 1.9E-01 | 8.8E-01 | 5 |
| TSC2 | 8.1E-01 | 9.8E-03 | 5.7E-01 | 1.7E-01 | -2.4E-01 | 8.4E-01 | 5 |
| ABHD17A | 6.5E-01 | 8.7E-02 | 8.9E-01 | 2.5E-02 | 2.4E-01 | 9.0E-01 | 5 |
| RNF151 | 2.4E+00 | 2.5E-02 | 2.0E+00 | 1.4E-01 | -4.6E-01 | 9.7E-01 | 5 |
| MRPS34 | 2.6E+00 | 5.1E-04 | 1.9E+00 | 2.9E-02 | -6.5E-01 | 6.2E-01 | 5 |
| NME3 | 1.6E+00 | 4.1E-02 | 1.2E+00 | 2.4E-01 | -3.7E-01 | 9.6E-01 | 5 |
| FBXW5 | 4.3E-01 | 1.1E-01 | 6.2E-01 | 2.3E-02 | 2.0E-01 | 8.1E-01 | 5 |
| ATP6AP1 | 6.4E-01 | 3.7E-03 | 7.2E-01 | 3.2E-03 | 7.5E-02 | 9.9E-01 | 5 |
| RHOT2 | 9.3E-01 | 4.9E-04 | 7.9E-01 | 1.1E-02 | -1.4E-01 | 9.9E-01 | 5 |
| WDR90 | 6.8E-01 | 1.9E-02 | 5.4E-01 | 1.4E-01 | -1.4E-01 | 9.9E-01 | 5 |
| TMEM8A | 8.6E-01 | 1.4E-03 | 9.5E-01 | 1.3E-03 | 8.8E-02 | 9.9E-01 | 5 |
| MPG | 6.6E-01 | 2.9E-02 | 5.0E-01 | 2.1E-01 | -1.6E-01 | 9.8E-01 | 5 |
| POLRMT | 6.9E-01 | 4.2E-02 | 7.1E-01 | 7.5E-02 | 1.2E-02 | 9.9E-01 | 5 |
| BSG | 1.1E+00 | 2.7E-05 | 1.0E+00 | 6.4E-04 | -1.1E-01 | 9.9E-01 | 5 |
| SURF2 | 4.7E-01 | 1.6E-01 | 9.0E-01 | 4.0E-03 | 4.3E-01 | 3.4E-01 | 5 |
| TCF25 | 6.4E-01 | 3.7E-02 | 6.3E-01 | 8.2E-02 | -1.5E-02 | 9.9E-01 | 5 |
| GBGT1 | 1.0E+00 | 9.5E-04 | 1.1E+00 | 2.4E-03 | 1.9E-02 | 9.9E-01 | 5 |
| GTF3C5 | 7.8E-01 | 1.4E-02 | 9.5E-01 | 5.4E-03 | 1.7E-01 | 9.7E-01 | 5 |
| RNF166 | 7.7E-01 | 7.5E-03 | 1.2E+00 | 1.1E-04 | 3.8E-01 | 4.4E-01 | 5 |
| CYBA | 5.8E-01 | 8.6E-03 | 6.7E-01 | 5.2E-03 | 9.4E-02 | 9.9E-01 | 5 |
| SLC25A24 | 7.8E-01 | 4.2E-02 | 5.6E-01 | 3.0E-01 | -2.2E-01 | 9.3E-01 | 5 |
| COQ4 | 1.4E+00 | 6.6E-05 | 1.1E+00 | 8.7E-03 | -3.1E-01 | 7.3E-01 | 5 |
| CIZ1 | 1.3E+00 | 3.0E-04 | 1.2E+00 | 2.2E-03 | -6.5E-02 | 9.9E-01 | 5 |
| TMED5 | 1.4E+00 | 6.4E-06 | 1.6E+00 | 5.0E-06 | 1.2E-01 | 9.9E-01 | 5 |
| EVI5 | 4.4E-01 | 5.8E-02 | 6.4E-01 | 6.7E-03 | 2.1E-01 | 6.9E-01 | 5 |
| VAC14 | 5.3E-01 | 1.5E-01 | 7.8E-01 | 3.4E-02 | 2.5E-01 | 8.2E-01 | 5 |
| RABEPK | 5.1E-01 | 1.3E-01 | 8.7E-01 | 8.9E-03 | 3.6E-01 | 5.4E-01 | 5 |
| RPL35 | 7.9E-01 | 6.1E-04 | 8.8E-01 | 4.4E-04 | 9.2E-02 | 9.9E-01 | 5 |
| NA | 9.2E-01 | 1.4E-02 | 1.3E+00 | 6.8E-04 | 3.9E-01 | 5.7E-01 | 5 |
| NA | 1.4E+00 | 6.5E-03 | 1.1E+00 | 9.5E-02 | -3.3E-01 | 9.1E-01 | 5 |
| NA | 8.7E-01 | 1.4E-01 | 1.5E+00 | 8.1E-03 | 6.3E-01 | 4.7E-01 | 5 |
| ACD | 1.4E+00 | 4.7E-03 | 1.2E+00 | 4.6E-02 | -2.2E-01 | 9.9E-01 | 5 |
| ATP6V0D1 | 1.0E+00 | 3.3E-06 | 7.1E-01 | 6.4E-03 | -3.3E-01 | 4.2E-01 | 5 |
| ZZZ3 | 8.6E-01 | 4.2E-02 | 7.2E-01 | 1.8E-01 | -1.4E-01 | 9.9E-01 | 5 |

| | | | | | | | |
|---|---|---|---|---|---|---|---|
| SRSF11 | 4.2E-01 | 2.3E-02 | 8.0E-01 | 1.5E-05 | 3.8E-01 | 1.0E-01 | 5 |
| NA | 1.2E+00 | 1.2E-03 | 1.3E+00 | 1.2E-03 | 1.2E-01 | 9.9E-01 | 5 |
| NA | 1.1E+00 | 1.7E-04 | 8.3E-01 | 1.9E-02 | -2.9E-01 | 7.4E-01 | 5 |
| NA | 7.5E-01 | 1.9E-02 | 5.8E-01 | 1.5E-01 | -1.7E-01 | 9.8E-01 | 5 |
| NA | 9.1E-01 | 4.0E-02 | 1.6E+00 | 2.5E-04 | 6.7E-01 | 2.3E-01 | 5 |
| POC1A | 9.1E-01 | 2.8E-02 | 7.6E-01 | 1.4E-01 | -1.5E-01 | 9.9E-01 | 5 |
| CILP2 | 1.3E+00 | 6.0E-04 | 1.0E+00 | 2.4E-02 | -2.7E-01 | 8.5E-01 | 5 |
| NA | 1.3E+00 | 1.1E-07 | 1.2E+00 | 1.5E-05 | -1.4E-01 | 9.8E-01 | 5 |
| FAM188A | 5.7E-01 | 2.4E-01 | 1.0E+00 | 2.0E-02 | 4.6E-01 | 5.0E-01 | 5 |
| NA | 1.5E+00 | 4.0E-02 | 1.1E+00 | 2.5E-01 | -3.5E-01 | 9.7E-01 | 5 |
| NA | 1.5E+00 | 4.0E-02 | 1.1E+00 | 2.5E-01 | -3.5E-01 | 9.7E-01 | 5 |
| NA | 1.5E+00 | 4.0E-02 | 1.1E+00 | 2.5E-01 | -3.5E-01 | 9.7E-01 | 5 |
| NA | 1.5E+00 | 4.0E-02 | 1.1E+00 | 2.5E-01 | -3.5E-01 | 9.7E-01 | 5 |
| NA | 1.5E+00 | 4.0E-02 | 1.1E+00 | 2.5E-01 | -3.5E-01 | 9.7E-01 | 5 |
| NA | 1.5E+00 | 4.0E-02 | 1.1E+00 | 2.5E-01 | -3.5E-01 | 9.7E-01 | 5 |
| LOXL1 | 9.2E-01 | 4.0E-04 | 9.9E-01 | 4.8E-04 | 7.2E-02 | 9.9E-01 | 5 |
| CD151 | 7.8E-01 | 2.4E-04 | 5.8E-01 | 2.0E-02 | -2.0E-01 | 7.7E-01 | 5 |
| ZNF444 | 6.7E-01 | 3.6E-02 | 8.8E-01 | 9.9E-03 | 2.1E-01 | 9.0E-01 | 5 |
| CENPP | 8.5E-01 | 4.7E-02 | 1.1E+00 | 1.8E-02 | 2.3E-01 | 9.6E-01 | 5 |
| NA | 1.1E+00 | 2.7E-02 | 1.7E+00 | 1.1E-03 | 5.4E-01 | 4.7E-01 | 5 |
| PRRG2 | 5.5E-01 | 1.6E-02 | 1.1E+00 | 1.2E-06 | 5.5E-01 | 3.7E-02 | 5 |
| NA | 1.7E+00 | 4.2E-03 | 1.2E+00 | 1.1E-01 | -4.9E-01 | 7.3E-01 | 5 |
| NA | 6.8E-01 | 1.3E-02 | 5.2E-01 | 1.4E-01 | -1.7E-01 | 9.4E-01 | 5 |
| CNTF | 1.1E+00 | 4.3E-02 | 1.0E+00 | 1.3E-01 | -1.0E-01 | 9.9E-01 | 5 |
| ZNF777 | 8.5E-01 | 3.5E-02 | 1.0E+00 | 2.4E-02 | 1.5E-01 | 9.9E-01 | 5 |
| NA | 1.9E+00 | 1.7E-02 | 3.1E+00 | 3.5E-05 | 1.3E+00 | 1.2E-01 | 5 |
| NA | 1.9E+00 | 1.7E-02 | 3.1E+00 | 3.5E-05 | 1.3E+00 | 1.2E-01 | 5 |
| NA | 1.9E+00 | 1.7E-02 | 3.1E+00 | 3.5E-05 | 1.3E+00 | 1.2E-01 | 5 |
| NA | 1.9E+00 | 1.7E-02 | 3.1E+00 | 3.5E-05 | 1.3E+00 | 1.2E-01 | 5 |
| NA | 1.3E+00 | 5.0E-03 | 1.6E+00 | 8.2E-04 | 3.4E-01 | 8.4E-01 | 5 |
| NA | 2.6E+00 | 1.2E-03 | 2.2E+00 | 1.2E-02 | -3.1E-01 | 9.9E-01 | 5 |
| NA | 6.4E-01 | 2.6E-02 | 4.6E-01 | 2.4E-01 | -1.8E-01 | 9.3E-01 | 5 |
| PPAP2B | 7.9E-01 | 2.1E-04 | 7.4E-01 | 2.2E-03 | -5.7E-02 | 9.9E-01 | 5 |
| RAB6C | 5.4E-01 | 1.5E-02 | 9.9E-01 | 9.8E-06 | 4.5E-01 | 1.2E-01 | 5 |
| RAB6C | 5.4E-01 | 1.5E-02 | 9.9E-01 | 9.8E-06 | 4.5E-01 | 1.2E-01 | 5 |
| NA | 1.3E+00 | 2.7E-05 | 1.5E+00 | 4.2E-06 | 2.1E-01 | 8.9E-01 | 5 |
| WDR83OS | 1.1E+00 | 8.0E-05 | 1.2E+00 | 2.7E-05 | 1.5E-01 | 9.8E-01 | 5 |
| ABHD17C | 9.1E-01 | 4.0E-04 | 8.9E-01 | 1.9E-03 | -2.0E-02 | 9.9E-01 | 5 |
| PTGES3 | 6.3E-01 | 1.4E-02 | 5.9E-01 | 4.8E-02 | -3.6E-02 | 9.9E-01 | 5 |
| COX7A2 | 1.0E+00 | 5.6E-05 | 1.2E+00 | 8.5E-06 | 1.8E-01 | 9.0E-01 | 5 |
| NCR3LG1 | 9.5E-01 | 7.8E-03 | 7.7E-01 | 7.5E-02 | -1.8E-01 | 9.9E-01 | 5 |
| CSRP1 | 1.2E+00 | 5.1E-05 | 1.3E+00 | 2.8E-05 | 1.4E-01 | 9.9E-01 | 5 |
| CDC42EP5 | 1.3E+00 | 1.0E-03 | 1.3E+00 | 1.4E-03 | 8.3E-02 | 9.9E-01 | 5 |
| FAM133B | 8.4E-01 | 9.0E-03 | 6.1E-01 | 1.5E-01 | -2.3E-01 | 8.7E-01 | 5 |
| ODF3B | 1.9E+00 | 1.7E-02 | 1.4E+00 | 1.8E-01 | -5.0E-01 | 9.0E-01 | 5 |
| NA | 8.0E-01 | 1.8E-04 | 6.6E-01 | 7.8E-03 | -1.4E-01 | 9.3E-01 | 5 |
| TNFRSF6B | 1.2E+00 | 4.9E-02 | 8.0E-01 | 3.4E-01 | -3.5E-01 | 9.1E-01 | 5 |
| NA | 1.7E+00 | 1.3E-01 | 2.5E+00 | 1.5E-02 | 8.9E-01 | 5.4E-01 | 5 |
| NA | 8.1E-01 | 3.1E-04 | 1.0E+00 | 2.3E-05 | 2.1E-01 | 7.8E-01 | 5 |
| SRPK3 | 9.5E-01 | 2.7E-02 | 1.2E+00 | 1.3E-02 | 2.1E-01 | 9.9E-01 | 5 |
| CDC34 | 1.1E+00 | 1.4E-04 | 1.1E+00 | 4.4E-04 | 1.7E-02 | 9.9E-01 | 5 |
| TP53RK | 9.8E-01 | 5.2E-04 | 1.3E+00 | 3.0E-05 | 2.7E-01 | 7.3E-01 | 5 |
| LDLRAP1 | 6.9E-01 | 1.0E-02 | 8.4E-01 | 3.4E-03 | 1.6E-01 | 9.5E-01 | 5 |
| ATF5 | 1.4E+00 | 7.8E-06 | 1.3E+00 | 9.2E-05 | -6.4E-02 | 9.9E-01 | 5 |
| METTL12 | 1.1E+00 | 4.4E-02 | 1.7E+00 | 1.1E-03 | 6.6E-01 | 4.2E-01 | 5 |
| METTL12 | 1.1E+00 | 4.4E-02 | 1.7E+00 | 1.1E-03 | 6.6E-01 | 4.2E-01 | 5 |
| NA | 1.9E+00 | 1.5E-02 | 2.1E+00 | 1.1E-02 | 2.3E-01 | 9.9E-01 | 5 |
| ZNF517 | 9.4E-01 | 7.2E-05 | 1.0E+00 | 5.0E-05 | 9.5E-02 | 9.9E-01 | 5 |
| LOH12CR1 | 9.8E-01 | 4.6E-05 | 1.1E+00 | 1.2E-05 | 1.5E-01 | 9.5E-01 | 5 |
| IFNE | 1.4E+00 | 2.2E-05 | 1.9E+00 | 1.6E-08 | 5.2E-01 | 2.9E-01 | 5 |
| RPL13A | 7.0E-01 | 7.5E-04 | 1.2E+00 | 2.6E-08 | 4.8E-01 | 7.2E-02 | 5 |
| RPL13A | 7.0E-01 | 7.5E-04 | 1.2E+00 | 2.6E-08 | 4.8E-01 | 7.2E-02 | 5 |
| RPL13A | 7.0E-01 | 7.5E-04 | 1.2E+00 | 2.6E-08 | 4.8E-01 | 7.2E-02 | 5 |
| RPL13A | 7.0E-01 | 7.5E-04 | 1.2E+00 | 2.6E-08 | 4.8E-01 | 7.2E-02 | 5 |
| RPL13A | 7.0E-01 | 7.5E-04 | 1.2E+00 | 2.6E-08 | 4.8E-01 | 7.2E-02 | 5 |
| CLIC3 | 6.9E-01 | 3.3E-02 | 4.9E-01 | 2.7E-01 | -2.0E-01 | 9.3E-01 | 5 |
| SIGMAR1 | 5.5E-01 | 5.3E-02 | 6.3E-01 | 4.6E-02 | 8.0E-02 | 9.9E-01 | 5 |
| ECI1 | 9.5E-01 | 1.2E-03 | 8.1E-01 | 1.8E-02 | -1.4E-01 | 9.9E-01 | 5 |
| NDUFA8 | 4.5E-01 | 5.0E-02 | 7.1E-01 | 2.5E-03 | 2.6E-01 | 5.5E-01 | 5 |
| CDR2L | 1.2E+00 | 1.3E-03 | 8.5E-01 | 5.2E-02 | -3.1E-01 | 8.1E-01 | 5 |
| COMMD7 | 7.4E-01 | 1.5E-04 | 9.1E-01 | 1.5E-05 | 1.7E-01 | 8.2E-01 | 5 |
| ZNHIT2 | 5.6E-01 | 3.1E-01 | 1.1E+00 | 2.4E-02 | 5.4E-01 | 4.8E-01 | 5 |
| PDCD2 | 3.3E-01 | 2.6E-01 | 5.9E-01 | 2.8E-02 | 2.6E-01 | 5.9E-01 | 5 |
| MBLAC2 | 5.3E-01 | 2.0E-01 | 8.4E-01 | 3.8E-02 | 3.1E-01 | 7.3E-01 | 5 |
| NFKBIL1 | 1.1E+00 | 4.8E-03 | 1.3E+00 | 2.5E-03 | 1.8E-01 | 9.9E-01 | 5 |
| NXPH3 | 1.4E+00 | 1.2E-03 | 1.9E+00 | 5.0E-05 | 4.4E-01 | 6.6E-01 | 5 |
| TUBGCP2 | 6.6E-01 | 1.4E-02 | 9.7E-01 | 5.2E-04 | 3.1E-01 | 5.6E-01 | 5 |
| BTBD19 | 8.4E-01 | 1.2E-02 | 1.2E+00 | 3.7E-04 | 4.0E-01 | 5.3E-01 | 5 |
| STARD4 | 5.9E-01 | 3.7E-02 | 4.1E-01 | 2.9E-01 | -1.7E-01 | 9.2E-01 | 5 |
| LAD1 | 7.6E-01 | 2.3E-03 | 7.5E-01 | 7.4E-03 | -6.5E-03 | 9.9E-01 | 5 |
| NUDT8 | 6.0E-01 | 6.9E-02 | 1.1E+00 | 6.4E-04 | 4.9E-01 | 2.6E-01 | 5 |
| GTDC2 | 1.2E+00 | 3.7E-04 | 1.2E+00 | 6.8E-04 | 5.7E-02 | 9.9E-01 | 5 |
| RDH14 | 6.9E-01 | 1.7E-03 | 7.8E-01 | 1.3E-03 | 8.4E-02 | 9.9E-01 | 5 |
| NA | 1.3E+00 | 4.9E-04 | 1.7E+00 | 9.4E-06 | 4.2E-01 | 5.5E-01 | 5 |
| NA | 8.3E-01 | 2.9E-03 | 9.2E-01 | 2.6E-03 | 9.1E-02 | 9.9E-01 | 5 |
| ERF | 1.6E+00 | 2.0E-09 | 1.4E+00 | 3.8E-07 | -1.3E-01 | 9.9E-01 | 5 |
| NA | 9.8E-01 | 3.1E-04 | 1.2E+00 | 1.5E-05 | 2.7E-01 | 7.1E-01 | 5 |
| GNG4 | 6.2E-01 | 1.2E-01 | 8.3E-01 | 4.9E-02 | 2.1E-01 | 9.5E-01 | 5 |
| GNG2 | 7.5E-01 | 3.6E-02 | 7.4E-01 | 7.9E-02 | -1.3E-02 | 9.9E-01 | 5 |
| ADH4 | 6.3E-01 | 1.9E-01 | 1.1E+00 | 1.6E-02 | 4.8E-01 | 5.5E-01 | 5 |
| C14orf79 | 7.8E-01 | 4.4E-02 | 9.4E-01 | 2.7E-02 | 1.6E-01 | 9.9E-01 | 5 |
| SNAPC2 | 1.2E+00 | 1.4E-05 | 1.1E+00 | 2.9E-04 | -9.5E-02 | 9.9E-01 | 5 |
| NA | 7.4E-01 | 1.5E-03 | 8.1E-01 | 1.6E-03 | 6.8E-02 | 9.9E-01 | 5 |
| TOMM7 | 4.2E-01 | 1.0E-01 | 7.2E-01 | 5.2E-03 | 3.0E-01 | 5.0E-01 | 5 |

| | | | | | | | |
|---|---|---|---|---|---|---|---|
| MZF1 | 8.9E-01 | 2.6E-02 | 7.4E-01 | 1.4E-01 | -1.5E-01 | 9.9E-01 | 5 |
| TSSC4 | 1.0E+00 | 3.3E-05 | 6.9E-01 | 2.5E-02 | -3.6E-01 | 4.3E-01 | 5 |
| NAA20 | 6.0E-01 | 1.0E-01 | 8.0E-01 | 3.9E-02 | 2.0E-01 | 9.5E-01 | 5 |
| DUSP7 | 1.6E+00 | 6.6E-02 | 2.2E+00 | 2.1E-02 | 5.4E-01 | 9.2E-01 | 5 |
| RAB11B | 6.0E-01 | 1.4E-02 | 6.3E-01 | 2.2E-02 | 3.4E-02 | 9.9E-01 | 5 |
| MAGOHB | 1.4E+00 | 5.2E-04 | 1.1E+00 | 1.7E-02 | -2.6E-01 | 9.1E-01 | 5 |
| OGFRL1 | 1.7E+00 | 3.6E-02 | 1.3E+00 | 2.2E-01 | -3.9E-01 | 9.5E-01 | 5 |
| DERL2 | 9.9E-01 | 4.3E-04 | 1.3E+00 | 1.8E-05 | 2.9E-01 | 6.9E-01 | 5 |
| NA | 1.4E+00 | 7.2E-05 | 1.8E+00 | 2.2E-06 | 3.6E-01 | 6.6E-01 | 5 |
| NA | 5.5E-01 | 1.0E-01 | 9.5E-01 | 4.1E-03 | 4.0E-01 | 4.4E-01 | 5 |
| NA | 1.9E+00 | 4.1E-02 | 1.6E+00 | 1.8E-01 | -3.1E-01 | 9.9E-01 | 5 |
| COL8A2 | 1.0E+00 | 2.8E-02 | 1.2E+00 | 1.1E-02 | 2.4E-01 | 9.6E-01 | 5 |
| COX7C | 1.8E+00 | 3.9E-07 | 1.8E+00 | 5.3E-06 | -5.4E-02 | 9.9E-01 | 5 |
| PTMA | 1.2E+00 | 3.1E-05 | 9.3E-01 | 5.2E-03 | -2.7E-01 | 7.6E-01 | 5 |
| PTMA | 1.2E+00 | 3.1E-05 | 9.3E-01 | 5.2E-03 | -2.7E-01 | 7.6E-01 | 5 |
| PTMA | 1.2E+00 | 3.1E-05 | 9.3E-01 | 5.2E-03 | -2.7E-01 | 7.6E-01 | 5 |
| PTMA | 1.2E+00 | 3.1E-05 | 9.3E-01 | 5.2E-03 | -2.7E-01 | 7.6E-01 | 5 |
| PTMA | 1.2E+00 | 3.1E-05 | 9.3E-01 | 5.2E-03 | -2.7E-01 | 7.6E-01 | 5 |
| PTMA | 1.2E+00 | 3.1E-05 | 9.3E-01 | 5.2E-03 | -2.7E-01 | 7.6E-01 | 5 |
| C4orf48 | 9.2E-01 | 1.4E-04 | 9.5E-01 | 3.0E-04 | 3.5E-02 | 9.9E-01 | 5 |
| SRM | 1.1E+00 | 3.0E-03 | 1.0E+00 | 1.6E-02 | -6.9E-02 | 9.9E-01 | 5 |
| AIF1 | 1.1E+00 | 1.6E-01 | 1.7E+00 | 3.3E-02 | 5.5E-01 | 7.5E-01 | 5 |
| NA | 1.0E+00 | 1.7E-02 | 7.9E-01 | 1.4E-01 | -2.1E-01 | 9.8E-01 | 5 |
| NA | 1.3E+00 | 4.3E-02 | 1.7E+00 | 1.0E-02 | 4.0E-01 | 8.4E-01 | 5 |
| NA | 4.9E+00 | 8.2E-03 | 4.0E+00 | 6.7E-02 | -8.4E-01 | 9.1E-01 | 5 |
| NA | 1.2E+00 | 2.1E-03 | 9.6E-01 | 3.3E-02 | -2.0E-01 | 9.8E-01 | 5 |
| NA | 1.2E+00 | 1.2E-01 | 1.6E+00 | 4.0E-02 | 4.2E-01 | 8.9E-01 | 5 |
| NA | 5.5E-01 | 8.8E-02 | 7.3E-01 | 3.3E-02 | 1.8E-01 | 9.4E-01 | 5 |
| NA | 2.3E+00 | 1.8E-07 | 2.1E+00 | 2.2E-05 | -2.7E-01 | 9.3E-01 | 5 |
| NA | 5.8E-01 | 3.6E-02 | 7.7E-01 | 9.0E-03 | 1.9E-01 | 8.8E-01 | 5 |
| NA | 7.5E-01 | 1.5E-02 | 7.1E-01 | 5.0E-02 | -4.1E-02 | 9.9E-01 | 5 |
| NA | 1.1E+00 | 1.4E-02 | 1.1E+00 | 3.9E-02 | -3.1E-02 | 9.9E-01 | 5 |
| NA | 1.2E+00 | 5.5E-02 | 1.4E+00 | 3.9E-02 | 2.0E-01 | 9.9E-01 | 5 |
| NA | 8.8E-01 | 9.4E-03 | 9.1E-01 | 1.9E-02 | 2.3E-02 | 9.9E-01 | 5 |
| NA | 5.2E+00 | 2.0E-04 | 4.6E+00 | 2.4E-03 | -6.3E-01 | 7.6E-01 | 5 |

| Gene | Primer | Sequence |
|---|---|---|
| NR4A2 | Fwd | AGGTTGCAATGCGTTCGT |
| NR4A2 | Rev | TTGAGCCCGTGTCTCTCTGT |
| PAX8 | Fwd | AGCGAATACTCTGGCAATGC |
| PAX8 | Rev | AACTCAGCAAGCTGGAGTTG |
| PER1 | Fwd | AGCGTGTCATGATGACCTACC |
| PER1 | Rev | TCTGAAAACCGAGGCTGCTG |
| GATA3 | Fwd | TGTGCGAACTGTCAAACCAC |
| GATA3 | Rev | TTCATGGTCAGGGGTCTGTTG |
| GUSB | Fwd | AGACAATTGCAGGCTTCCAC |
| GUSB | Rev | TTGCGTTTCTGATCCAGCAC |
